# GENETH’OFF: a flexible workflow for genome-wide profiling of CRISPR/Cas off-targets

**DOI:** 10.1101/2025.09.30.679427

**Authors:** Corre Guillaume, Rouillon Marine, Mombled Margaux, Amendola Mario

## Abstract

**Summary:** We developed GENETH’OFF, a flexible single-command versatile Snakemake workflow designed for the comprehensive analysis of CRISPR/Cas9 related OFF-targets genomic positions from GUIDE-Seq derived protocols. It efficiently processes multiplexed libraries from different organisms, PCR orientations, and Cas nucleases with varying PAM specificities in a single run, all based on a simple user-specified datasheet containing sample metadata.

**Availability and implementation:** GENETH’OFF is freely available at: https://github.com/gcorre/GENETHOFF

**Supplementary information:** Supplementary data are available at Bioinformatics online.

## INTRODUCTION

CRISPR/Cas genome-editing technologies have transformed genome engineering by enabling precise and programmable DNA modifications. These systems show major promise for the treatment of rare genetic diseases, as illustrated by the recent approval of Casgevy® for sickle cell disease (Frangoul *et al*. 2024). Editing relies on the induction of a site-specific double-strand break (DSB) by a Cas nuclease, guided by a sequence-specific guide RNA (gRNA) and a protospacer adjacent motif (PAM), followed by DNA repair through endogenous cellular pathways. Despite its precision, CRISPR/Cas editing is associated with a key safety concern: genotoxicity caused by off-target (OT) DNA cleavage. Off-target activity occurs when the Cas/gRNA complex cleaves genomic sites that partially match the intended target sequence. Such unintended DSBs can result in insertions or deletions, chromosomal rearrangements, or disruption of essential genes, potentially leading to genomic instability, cellular dysfunction, or oncogenic transformation (Amendola, Brusson, and Miccio 2022). Comprehensive detection and characterization of off-target activity is therefore essential to assess genotoxic risk and to guide the development of safer genome-editing tools as demonstrated by the recent release of FDA guidelines (<u>FDA-2026-D-1255</u>). This need has driven the emergence of genome-wide experimental methods combined with computational prediction approaches (Kalter *et al*. 2025). Among these, GUIDE-seq (Genome-wide Unbiased Identification of Double-strand breaks Enabled by Sequencing ) is a widely used cellular assay that enables unbiased, genome-wide identification of DSBs at frequencies as low as 0.1% (Tsai *et al*. 2015). The method relies on the integration of short double-stranded oligonucleotides (dsODN) at Cas-induced DSBs, followed by next-generation sequencing (NGS) to identify and quantify both on- and off-target sites. Multiple derived protocols have been developed to increase sensitivity (Nobles *et al*. 2019, Huang *et al*. 2021, Yang *et al*. 2024). Although multiple pipelines have been co-developed to analyze sequencing data, our pipeline focuses on flexibility and robustness with the possibility to process multiple samples from different organisms, using different CAS nuclease specificities and library structure in a single run. It is based on popular bioinformatics tools with versioning based on Conda, all processing steps being piped together using the Snakemake workflow management for reproducibility and ease of installation.

### IMPLEMENTATION

The pipeline is implemented using the Snakemake workflow management language (Mölder *et al*. 2021), which enables structured rule-based execution with efficient job scheduling and resource management. This framework ensures scalability, robustness, and reproducibility across computing environments. Whenever possible, we relied on well-established bioinformatics tools to maximize reliability and maintainability. Custom Python and R scripts were developed only when required to provide functionality not covered by existing software.

The entire workflow is executed through a single command-line instruction on a Linux system (tested on Debian and Ubuntu Linux distributions), and each step runs within a Conda-managed virtual environment, ensuring strict control of software versions and dependencies. This design guarantees reproducibility across users and platforms. The pipeline is fully documented and publicly available through our GitHub repository, which includes installation instructions, configuration examples, and a test dataset (https://github.com/gcorre/GENETHOFF; DOI: 10.5281/zenodo.17341915).

### WORKFLOW OVERVIEW AND STEPS

The proposed workflow consists of distinct rules executing sequential data processing (Figure1).Thanks to precise resources management, multiple samples can be processed in parallel, minimizing computing time and resources.

#### User inputs

Alt text: “Graphical representation of the workflow showing the 4 main steps of data processing starting with user inputs to final report generation.”

The pipeline requires three mandatory input elements (Figure 1).

**Figure 1:**
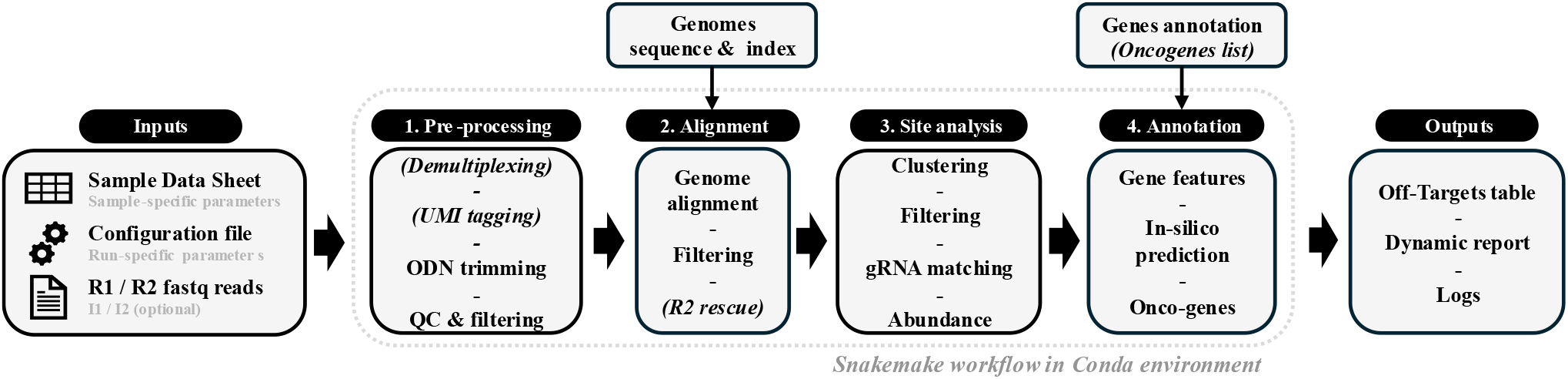
Overview of the analysis pipeline. Steps description is provided in the text. Optional steps are in parenthesis.

First, paired-end sequencing files (R1 and R2) and indexing files (I1 and I2) generated by the NGS platform are provided. If libraries have already been demultiplexed and index files are unavailable, the pipeline attempts to extract index sequences from the read headers, as I2 typically contains the UMI in addition to the sample barcode. If no index sequence can be recovered, double-strand break frequencies are estimated using fragment size deduplication or raw read counts rather than UMI counts, allowing quantification even when UMI information is unavailable.

Second, users must supply a Sample Data Sheet (SDS) in Excel or CSV format containing library-specific experimental metadata, including gRNA and PAM sequences, library name, reference genome, barcode sequences, and library structure. The SDS enables the pipeline to support heterogeneous experimental designs and to associate each sample with the correct reference genome and annotation resources. Libraries sharing identical names and metadata are automatically merged, facilitating the aggregation of technical or biological replicates.

Third, a configuration file defines processing parameters applied globally to all samples in a run, including paths to external tools, path to raw data, oligonucleotide sequences, thresholds applied at the different steps of the pipeline progression, the selected quantification method and whether to clean intermediate files. A checkpoint can be added to stop the pipeline before completion, just after OT identification and quantification.

For datasets generated using library designs not natively compatible with the pipeline (e.g., TagSeq or OliTag-seq), we provide conversion scripts on our GitHub repository to enable seamless integration.

#### Demultiplexing, trimming and UMI extraction

Raw reads are demultiplexed based on the unique i5/ i7 barcodes combination into separate individual R1 and R2 samples files. Paired-end reads containing the double-stranded oligonucleotide (dsODN) sequence within the R2 sequence are trimmed to remove both the dsODN and any Illumina adaptors (Figure1, step 1). Read pairs with no detectable ODN sequence and read pairs containing at least one read shorter than 25 bp were excluded from downstream analyses. When UMIs are used to estimate the frequency of ODN insertion events, the UMI sequence is extracted from the I2 index file. Demultiplexing, trimming, filtering and UMI extraction are performed using the *cutadapt* program (Martin 2011) with 6 threads, except demultiplexing that takes all user provided threads. Sequences of dsODN, adaptors and the pattern of the UMI are user-defined per-library in the sample data sheet allowing compatibility with alternative library designs and processing of libraries using different sequences in a single run. The default error rates used during demultiplexing (0), ODN trimming (0.1), UMI extraction (0), and the maximum distance allowed during UMI clustering (1) can be modified through the configuration file.

#### Genomic alignment

Trimmed paired-end reads are aligned to the reference genome (Figure1, step 2) specified in the Sample Data Sheet for each library using bowtie2 (Langmead *et al*. 2019). When paired-end alignment fails or R1 reads are too short, R2 reads are optionally rescued and aligned as single-end reads. This rescue strategy is disabled when fragment size–based quantification is selected, as accurate fragment length estimation requires both read ends.

Alongside uniquely aligned reads (MAPQ > 20), multi-mapping reads (MAPQ = 1 and identical AS/XS scores) are retained as discarding such reads would systematically underestimate off-target (OT) activity in repetitive or homologous regions and prevent detection of gRNAs intentionally designed to cleave multiple similar loci (Pavani *et al*. 2021). Reads with multiple valid alignments are assigned to one of the genomic positions at random to avoid counting the same read multiple times. This strategy reduces the resources as assigning reads to all its putative mapping positions would require extensive calculations and would significantly increase the processing time. As the number of analyzed reads increases, using a random assignment or an exhaustive assignment with weighted abundance would result in the same quantification. One limitation of these two approaches is that the abundance of the true OT site from where each read arises from may be under-estimated.

#### Insertion site calling and quantification

Putative cleavage sites are identified from the genomic start positions of R2 reads and clustered within a user-defined genomic window (Figure 1, step 3). For each cluster, forward and reverse ODN insertions are quantified either using fragment length or using the UMI counts using UMI-tools (Smith, Heger, and Sudbery 2017) to accommodate for sequencing errors. Off-target sites supported by both PCR orientations and both ODN insertion directions are assigned increased confidence. Quantification can alternatively be performed using fragment size information or raw read counts, depending on user-defined settings. *To avoid calling low-confidence off-target sites, UMIs supported by few reads and cleavage positions supported by few unique UMIs can be filtered out using user-defined thresholds (default values: 3 and 2, respectively)*.

#### gRNA detection and annotation

The presence of the expected gRNA sequence within each cluster is confirmed using Smith–Waterman local alignment, enabling accurate detection and quantification of mismatches, insertions, and deletions (bulges) between the gRNA and genomic target (Figure1, step 6). Incorporation of bulge-aware alignment is critical, as bulges have been shown to enhance nuclease activity in certain contexts (Lin *et al*. 2014, Corsi *et al*. 2022). User-defined thresholds control acceptable numbers of mismatches and bulges.

Identified OT sites are then annotated with genomic features using user-supplied GTF files (Figure1, step <u>4</u>). In addition, oncogene annotations can be performed using the OncoKB database (Chakravarty *et al*. 2017), with support for user-defined oncogenes annotation lists.

#### Off-targets prediction

An integrated prediction module runs in parallel with experimental processing. For each gRNA–PAM– genome combination, SWOffinder, a state-of-the-art prediction tools generate genome-wide theoretical OT sites, accounting for mismatch and bulge tolerance (Yaish *et al*. 2024). This prediction step employs established algorithms that consider sequence complementarity, PAM requirements, and bulge tolerance parameters. The module generates a genome-wide prediction of theoretical OT for each guide RNA, accounting for both perfect and imperfect matches within user-defined tolerance thresholds, and position-matched them with the experimentally identified OT. While experimental detection remains the primary evidence of off-target activity, computational predictions provide complementary information that can support the prioritization and interpretation of candidate sites, and sites identified by both approaches may warrant higher priority for follow-up analyses (Angelini Stewart *et al*. 2026, Kaufmann *et al*. 2026).

#### Output

Results are compiled into an interactive HTML report containing QC metrics, OT summaries, sortable and filterable tables, and graphical visualizations. Tables include sequence alignments highlighting mismatches and bulges, relative abundance, prediction status, and gene and oncogene annotations (Figure1, step 8). Additional visual outputs include rank-abundance curves to distinguish true OT signals from background noise, as well as chromosomal distribution plots. Complete result tables are also provided as Excel files for downstream analysis.

### USE CASE

A test dataset is publicly available in the Sequence Read Archive (SRA) under accession PRJNA1335208. This dataset can be analyzed using the configuration file and sample datasheet provided on our GitHub repository. A complete example of the resulting output is included as a supplementary file (supplementary_file1.zip).

#### BENCHMARKING

GENETHOFF was benchmarked against three established GUIDE-seq analysis workflows, iGUIDE-seq v1.2.0 and GUIDE-seq v1.0.2/v2, using the publicly available dataset PRJNA506241. This dataset comprises 17 GUIDE-seq libraries generated from four gRNAs (B2M, TRAC, VEGFAs2, and VEGFAs3) and a mock control, representing a total of 11.7 million sequencing reads. Benchmarking was performed on a Debian GNU/Linux 12 (Bookworm) server equipped with an Intel® Xeon® Gold 6246R CPU (3.40 GHz), 64 hardware threads, and 385 GB of RAM.

Runtime analysis demonstrated that GENETHOFF substantially outperformed the other workflows when using the same number of threads, reducing processing time by approximately 4-to 7-fold (Figure 2A). The complete analysis of the 17 libraries was completed in approximately 15 minutes. In addition, GENETHOFF exhibited efficient scalability as the number of available threads increased. Unlike the other pipelines, GENETHOFF effectively utilized nearly all allocated computational resources (Figure 2B), maintaining high CPU occupancy throughout the analysis.

**Figure 2:**
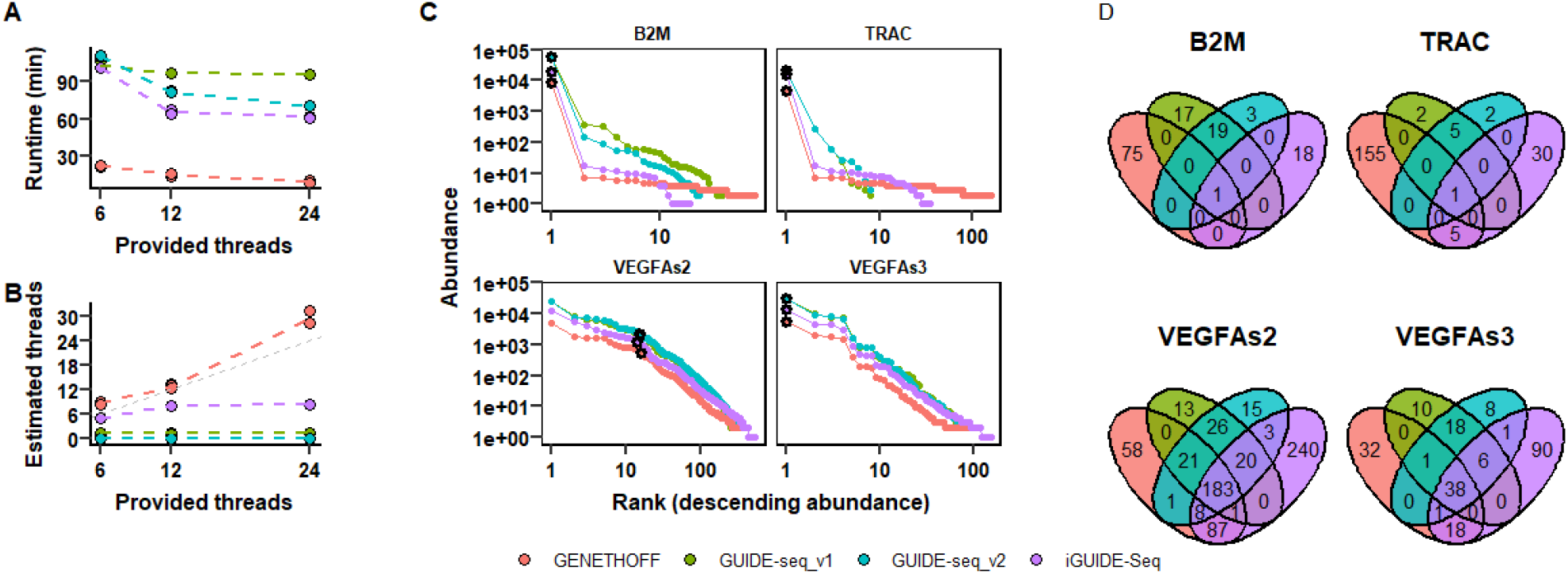
(A) Average runtime (n = 3) as a function of the total number of allocated threads. (B) Estimated thread utilization derived from wall-clock and CPU-time metrics. (C) Rank-abundance distributions of detected cleavage sites. Black dots indicate the on-target cleavage site. (D) Overlap of detected cleavage sites among pipelines for each gRNA target. All panels share the same color legend. Sites detected within a 10 bp window were considered identical across pipelines.

To compare cleavage-site calls across pipelines, sites located within a 10 bp window were merged and considered to represent the same cleavage event. As shown in Figure 2C, the four pipelines generated highly similar cleavage-site abundance profiles across all gRNAs. The on-target site was consistently identified by all methods and displayed comparable abundance estimates (black dots). Although each pipeline reported a small number of unique cleavage sites (Figure 2D), these were predominantly low-abundance events.

To quantitatively assess agreement in site prioritization, we computed the Kendall correlation and the Rank Biased Overlap index which evaluate the pairwise ranking order between two lists (Supplemental figure 1). The results suggest that not only the four workflows detect almost the same subset of dominant off-target sites but that they also agree on their ranking. As expected, the 2 versions of Guide-seq give more similar results. Of note the lower agreement observed for B2M and TRAC. It is driven by the large number of low-abundance sites for which small differences in read counts can determine whether a site is detected by a given workflow. Because these sites are supported by few UMIs, their ranking is not robust, leading to lower rank concordance. Such discrepancies among the low abundance sites are expected and reflect methodological differences in filtering, alignment and cleavage-site calling thresholds and methods.

## CONCLUSION

GUIDE-seq remains a gold-standard unbiased method for genome-wide detection of CRISPR/Cas9 off-target cleavage. Here, we address limitations of existing pipelines by implementing key improvements, including bulge-aware gRNA alignment, robust handling of multi-hit reads, integrated off-target prediction, oncogene annotation, and comprehensive interactive reporting. The pipeline is fast, reproducible and library design agnostic. It is configurable through simple input files, enabling analysis of multiplexed datasets across organisms, PAM variants, PCR orientations, and dsODN designs. By combining experimental and computational evidence, it improves confidence in OT detection. Overall, we believe this pipeline provides a robust and versatile framework for GUIDE-seq analysis and contributes to the development of safer and more reliable CRISPR/Cas-based genome-editing therapies.

## Acknowledgements

The authors thank Ayal Hendel, Nechama Kalter and the whole Amendola’s laboratory for fruitful discussion. The authors are Genopole’s members, first french biocluster dedicated to genetic, biotechnologies and biotherapies. We gratefully acknowledge the AFM-Telethon foundation, the Conseil Général de l’Essonne (ASTRES) and Genopole Research in Evry for the financial help to purchase equipment, the France Relance program; the INSERM; the University of Evry Val d’Essonne.

## Fundings

This work was supported by the AFM-Telethon foundation [BE-DREP]; the French National Research Agency [IRIS ANR-21-CE14-0063-03, GenoTher ANR-23-BIOC-0003]; National Recovery and Resilience Plan [CN00000041 – CUP UNIMORE E93C22001080001] and the European Union [MAGIC grant 101080690, EDISCD grant 101057659].

## Data availability

Example dataset to test the pipeline is available on SRA under ID: PRJNA1335208 and is used to reproduce results in supplementary_file1.zip. Code has been registered at DOI: 10.5281/zenodo.17341915.

---

~~~
<svg width=“2995” height=“747” xmlns=“http://www.w3.org/2000/svg” xmlns:xlink=“http://www.w3.org/1999/xlink” xml:space=“preserve” overflow=“hidden”><g transform=“translate(-710 -599)”><path d=“M713.5 944.001C713.5 911.692 739.692 885.5 772.001 885.5L1230 885.5C1262.31 885.5 1288.5 911.692 1288.5 944.001L1288.5 1178C1288.5 1210.31 1262.31 1236.5 1230 1236.5L772.001 1236.5C739.692 1236.5 713.5 1210.31 713.5
1178Z” stroke=“#000000” stroke-width=“6.875” stroke-miterlimit=“8” fill=“#F2F2F2” fill-rule=“evenodd”/><g><g><g><path d=“M88.8332 41.1666 67.1665 41.1666 67.1665 28.1666 88.8332 28.1666 88.8332 41.1666ZM88.8332
58.4999 67.1665 58.4999 67.1665 45.4999 88.8332 45.4999 88.8332
58.4999ZM88.8332 75.8332 67.1665 75.8332 67.1665 62.8332 88.8332 62.8332
88.8332 75.8332ZM41.1666 75.8332 41.1666 62.8332 62.8332 62.8332 62.8332
75.8332 41.1666 75.8332ZM15.1666 75.8332 15.1666 62.8332 36.8333 62.8332
36.8333 75.8332 15.1666 75.8332ZM15.1666 45.4999 36.8333 45.4999 36.8333
58.4999 15.1666 58.4999 15.1666 45.4999ZM15.1666 28.1666 36.8333 28.1666
36.8333 41.1666 15.1666 41.1666 15.1666 28.1666ZM62.8332 45.4999 62.8332
58.4999 41.1666 58.4999 41.1666 45.4999 62.8332 45.4999ZM62.8332 28.1666
62.8332 41.1666 41.1666 41.1666 41.1666 28.1666 62.8332 28.1666ZM8.66665
21.6666 8.66665 82.3332 95.3331 82.3332 95.3331 21.6666 8.66665 21.6666Z”
transform=“matrix(1.04808 0 0 1 744 904)”/></g></g></g><g><g><g><path
d=“M64.2418 40.4084C59.6918 40.4084 56.1168 36.7251 56.1168 32.2834
56.1168 27.8417 59.8001 24.1584 64.2418 24.1584 68.7918 24.1584 72.3668
27.8417 72.3668 32.2834 72.3668 36.7251 68.6834 40.4084 64.2418
40.4084ZM82.5501 27.1917C82.1168 25.675 81.5751 24.2667 80.8168
22.9667L82.5501 17.875 78.6501 13.975 73.5584 15.7084C72.2584 14.95
70.8501 14.4084 69.3334 13.975L66.9501 9.20835 61.5334 9.20835 59.1501
13.975C57.6334 14.4084 56.2251 14.95 54.9251 15.7084L49.8334 13.975
45.9334 17.875 47.6667 22.9667C46.9084 24.2667 46.3667 25.675 45.9334
27.1917L41.1667 29.575 41.1667 34.9917 45.9334 37.3751C46.3667 38.8917
46.9084 40.3001 47.6667 41.6001L45.9334 46.6917 49.7251 50.4834 54.8167
48.7501C56.1168 49.5084 57.5251 50.0501 59.0418 50.4834L61.4251 55.2501
66.8418 55.2501 69.2251 50.4834C70.7418 50.0501 72.1501 49.5084 73.4501
48.7501L78.5418 50.4834 82.4418 46.6917 80.7085 41.6001C81.4668 40.3001
82.1168 38.7834 82.5501 37.3751L87.3168 34.9917 87.3168 29.575 82.5501
27.1917Z” transform=“matrix(1.04807 0 0 1 744 1010)”/><path d=“M39.7584
79.8418C35.2084 79.8418 31.6334 76.1585 31.6334 71.7168 31.6334 67.1668
35.3167 63.5918 39.7584 63.5918 44.3084 63.5918 47.8834 67.2751 47.8834
71.7168 47.8834 76.1585 44.3084 79.8418 39.7584 79.8418L39.7584
79.8418ZM56.3334 62.4001 58.0668 57.3084 54.1668 53.4084 49.0751
55.1418C47.7751 54.3834 46.2584 53.8418 44.8501 53.4084L42.4667 48.6417
37.0501 48.6417 34.6667 53.4084C33.1501 53.8418 31.7417 54.3834 30.4417
55.1418L25.35 53.4084 21.5584 57.2001 23.1834 62.2918C22.425 63.5918
21.8834 65.1084 21.45 66.5168L16.6834 68.9001 16.6834 74.3168 21.45
76.7001C21.8834 78.2168 22.425 79.6251 23.1834 80.9251L21.5584 86.0168
25.35 89.8085 30.4417 88.1835C31.7417 88.9418 33.1501 89.4835 34.6667
89.9168L37.0501 94.6835 42.4667 94.6835 44.8501 89.9168C46.3667 89.4835
47.7751 88.9418 49.0751 88.1835L54.1668 89.9168 57.9584 86.0168 56.3334
81.0335C57.0918 79.7335 57.6334 78.3251 58.0668 76.8085L62.8334 74.4251
62.8334 69.0084 58.0668 66.6251C57.6334 65.1084 57.0918 63.7001 56.3334
62.4001Z” transform=“matrix(1.04807 0 0 1 744 1010)”/></g></g></g><text font-family=“Times New Roman,Times New Roman_MSFontService,sans-serif” font-weight=“700” font-size=“41” transform=“matrix(1 0 0 1 873.867 1055)”>Configuration file</text><text fill=“#A6A6A6” font-family=“Times New Roman,Times New Roman_MSFontService,sans-serif” font-weight=“700”
font-size=“32” transform=“matrix(1 0 0 1 873.867 1095)”>Run</text><text fill=“#A6A6A6” font-family=“Times New Roman,Times New Roman_MSFontService,sans-serif” font-weight=“700” font-size=“32” transform=“matrix(1 0 0 1 932.304 1095)”>-</text><text fill=“#A6A6A6” font-family=“Times New Roman,Times New Roman_MSFontService,sans-serif” font-weight=“700” font-size=“32” transform=“matrix(1 0 0 1 943.19 1095)”>specific parameters</text><g><g><g><path d=“M24.9166 88.8332
24.9166 15.1666 51.9999 15.1666 51.9999 37.9166 79.0832 37.9166 79.0832
88.8332 24.9166 88.8332ZM58.4999 17.875 72.0415 31.4166 58.4999 31.4166
58.4999 17.875ZM58.4999 8.66665 18.4166 8.66665 18.4166 95.3331 85.5832
95.3331 85.5832 32.4999 58.4999 8.66665Z” transform=“matrix(1.04808 0 0 1
744 1113)”/><path d=“M31.4166 48.7499 72.5832 48.7499 72.5832 53.0832
31.4166 53.0832Z” transform=“matrix(1.04808 0 0 1 744 1113)”/><path
d=“M31.4166 40.0833 45.4999 40.0833 45.4999 44.4166 31.4166 44.4166Z”
transform=“matrix(1.04808 0 0 1 744 1113)”/><path d=“M31.4166 57.4166
72.5832 57.4166 72.5832 61.7499 31.4166 61.7499Z”
transform=“matrix(1.04808 0 0 1 744 1113)”/><path d=“M31.4166 66.0832
72.5832 66.0832 72.5832 70.4165 31.4166 70.4165Z”
transform=“matrix(1.04808 0 0 1 744 1113)”/><path d=“M31.4166 74.7499
72.5832 74.7499 72.5832 79.0832 31.4166 79.0832Z”
transform=“matrix(1.04808 0 0 1 744 1113)”/></g></g></g><text font-family=“Times New Roman,Times New Roman_MSFontService,sans-serif” font-weight=“700” font-size=“41” transform=“matrix(1 0 0 1 873.867 1158)”>R1 / R2 </text><text font-family=“Times New Roman,Times New Roman_MSFontService,sans-serif” font-weight=“700” font-size=“41” transform=“matrix(1 0 0 1 1017.1 1158)”>fastq</text><text font-family=“Times New Roman,Times New Roman_MSFontService,sans-serif” font-weight=“700” font-size=“41” transform=“matrix(1 0 0 1 1114.49 1158)”>reads</text><text fill=“#A6A6A6” font-family=“Times New Roman,Times New Roman_MSFontService,sans-serif” font-weight=“700” font-size=“32” transform=“matrix(1 0 0 1 873.867 1198)”>I1 / I2
(optional)</text><path d=“M2310.5 941.001C2310.5 908.692 2336.69 882.5 2369 882.5L2642 882.5C2674.31 882.5 2700.5 908.692 2700.5 941.001L2700.5
1175C2700.5 1207.31 2674.31 1233.5 2642 1233.5L2369 1233.5C2336.69 1233.5
2310.5 1207.31 2310.5 1175Z” stroke=“#000000” stroke-width=“6.875”
stroke-miterlimit=“8” fill=“#F2F2F2” fill-rule=“evenodd”/><text font-family=“Times New Roman,Times New Roman_MSFontService,sans-serif” font-weight=“700” font-size=“41” transform=“matrix(1 0 0 1 2412.51 924)”>Clustering </text><text font-family=“Times New Roman,Times New Roman_MSFontService,sans-serif” font-weight=“700” font-size=“41” transform=“matrix(1 0 0 1 2498.44 973)”>-</text><text font-family=“Times New Roman,Times New Roman_MSFontService,sans-serif” font-weight=“700” font-size=“41” transform=“matrix(1 0 0 1 2428.55 1023)”>Filtering</text><text font-family=“Times New Roman,Times New Roman_MSFontService,sans-serif” font-weight=“700” font-size=“41” transform=“matrix(1 0 0 1 2498.44 1072)”>-</text><text font-family=“Times New Roman,Times New Roman_MSFontService,sans-serif” font-weight=“700” font-size=“41” transform=“matrix(1 0 0 1 2362.66 1122)”>gRNA matching</text><text font-family=“Times New Roman,Times New Roman_MSFontService,sans-serif” font-weight=“700” font-size=“41” transform=“matrix(1 0 0 1 2498.44 1171)”>-</text><text font-family=“Times New Roman,Times New Roman_MSFontService,sans-serif” font-weight=“700” font-size=“41” transform=“matrix(1 0 0 1 2404.49
1221)”>Abundance</text><path d=“M2791.5 941.001C2791.5 908.692 2817.69
882.5 2850 882.5L3091 882.5C3123.31 882.5 3149.5 908.692 3149.5
941.001L3149.5 1175C3149.5 1207.31 3123.31 1233.5 3091 1233.5L2850
1233.5C2817.69 1233.5 2791.5 1207.31 2791.5 1175Z” stroke=“#042433”
stroke-width=“6.875” stroke-miterlimit=“8” fill=“#F2F2F2” fill-rule=“evenodd”/><text font-family=“Times New Roman,Times New Roman_MSFontService,sans-serif” font-weight=“700” font-size=“41” transform=“matrix(1 0 0 1 2847.82 949)”>Gene features</text><text font-family=“Times New Roman,Times New Roman_MSFontService,sans-serif” font-weight=“700” font-size=“41” transform=“matrix(1 0 0 1 2962.98 998)”>-</text><text font-family=“Times New Roman,Times New Roman_MSFontService,sans-serif” font-weight=“700” font-size=“41” transform=“matrix(1 0 0 1 2898.81 1048)”>In</text><text font-family=“Times New Roman,Times New Roman_MSFontService,sans-serif” font-weight=“700” font-size=“41” transform=“matrix(1 0 0 1 2937.77 1048)”>-</text><text font-family=“Times New Roman,Times New Roman_MSFontService,sans-serif” font-weight=“700” font-size=“41” transform=“matrix(1 0 0 1 2951.52 1048)”>silico </text><text font-family=“Times New Roman,Times New Roman_MSFontService,sans-serif” font-weight=“700” font-size=“41” transform=“matrix(1 0 0 1 2879.33 1097)”>prediction</text><text font-family=“Times New Roman,Times New Roman_MSFontService,sans-serif” font-weight=“700” font-size=“41” transform=“matrix(1 0 0 1 2962.98 1147)”>-</text><text font-family=“Times New Roman,Times New Roman_MSFontService,sans-serif” font-style=“italic” font-weight=“700” font-size=“41” transform=“matrix(1 0 0 1 2855.27 1196)”>(Onco</text><text font-family=“Times New Roman,Times New Roman_MSFontService,sans-serif” font-style=“italic” font-weight=“700” font-size=“41” transform=“matrix(1 0 0 1 2960.69 1196)”>-</text><text font-family=“Times New Roman,Times New Roman_MSFontService,sans-serif” font-style=“italic” font-weight=“700” font-size=“41” transform=“matrix(1 0 0 1 2974.44 1196)”>genes)</text><pathd=“M1924.5 930.167C1924.5 903.289
1946.29 881.5 1973.17 881.5L2167.83 881.5C2194.71 881.5 2216.5 903.289
2216.5 930.167L2216.5 1185.83C2216.5 1212.71 2194.71 1234.5 2167.83
1234.5L1973.17 1234.5C1946.29 1234.5 1924.5 1212.71 1924.5 1185.83Z”
stroke=“#042433” stroke-width=“6.875” stroke-miterlimit=“8” fill=“#F2F2F2” fill-rule=“evenodd”/><text font-family=“Times New Roman,Times New Roman_MSFontService,sans-serif” font-weight=“700” font-size=“41” transform=“matrix(1 0 0 1 1997.02 949)”>Genome </text><text font-family=“Times New Roman,Times New Roman_MSFontService,sans-serif” font-weight=“700” font-size=“41” transform=“matrix(1 0 0 1 1982.12 998)”>alignment</text><text font-family=“Times New Roman,Times New Roman_MSFontService,sans-serif” font-weight=“700” font-size=“41” transform=“matrix(1 0 0 1 2063.48 1048)”>-</text><text font-family=“Times New Roman,Times New Roman_MSFontService,sans-serif” font-weight=“700” font-size=“41” transform=“matrix(1 0 0 1 1993.58 1097)”>Filtering</text><text font-family=“Times New Roman,Times New Roman_MSFontService,sans-serif” font-weight=“700” font-size=“41” transform=“matrix(1 0 0 1 2063.48 1147)”>-</text><text font-family=“Times New Roman,Times New Roman_MSFontService,sans-serif” font-style=“italic” font-weight=“700” font-size=“41” transform=“matrix(1 0 0 1 1972.38
1196)”>(R2 rescue)</text><path d=“M1416.5 940.334C1416.5 907.841 1442.84
881.5 1475.33 881.5L1761.67 881.5C1794.16 881.5 1820.5 907.841 1820.5
940.334L1820.5 1175.67C1820.5 1208.16 1794.16 1234.5 1761.67
1234.5L1475.33 1234.5C1442.84 1234.5 1416.5 1208.16 1416.5 1175.67Z”
stroke=“#000000” stroke-width=“6.875” stroke-miterlimit=“8”
fill=“#F2F2F2” fill-rule=“evenodd”/><text font-family=“Times New Roman,Times New Roman_MSFontService,sans-serif” font-style=“italic” font-weight=“700” font-size=“41” transform=“matrix(1 0 0 1 1472.42 924)”>(Demultiplexing)</text><text font-family=“Times New Roman,Times New Roman_MSFontService,sans-serif” font-style=“italic” font-weight=“700” font-size=“41” transform=“matrix(1 0 0 1 1611.07 973)”>-</text><text
font-family=“Times New Roman,Times New Roman_MSFontService,sans-serif” font-style=“italic” font-weight=“700” font-size=“41” transform=“matrix(10 0 1 1493.62 1023)”>(UMI tagging)</text><text font-family=“Times New Roman,Times New Roman_MSFontService,sans-serif” font-style=“italic” font-weight=“700” font-size=“41” transform=“matrix(1 0 0 1 1611.07 1072)”>-
</text><text font-family=“Times New Roman,Times New Roman_MSFontService,sans-serif” font-weight=“700” font-size=“41” transform=“matrix(1 0 0 1 1483.31 1122)”>ODN trimming</text><text font-family=“Times New Roman,Times New Roman_MSFontService,sans-serif” font-weight=“700” font-size=“41” transform=“matrix(1 0 0 1 1611.07 1171)”>-
</text><text font-family=“Times New Roman,Times New Roman_MSFontService,sans-serif” font-weight=“700” font-size=“41” transform=“matrix(1 0 0 1 1488.47 1221)”>QC & filtering</text><path
d=“M1382.5 848.668C1382.5 803.288 1419.29 766.5 1464.67 766.5L3102.33
766.5C3147.71 766.5 3184.5 803.288 3184.5 848.668L3184.5 1177.33C3184.5
1222.71 3147.71 1259.5 3102.33 1259.5L1464.67 1259.5C1419.29 1259.5
1382.5 1222.71 1382.5 1177.33Z” stroke=“#D1D1D1” stroke-width=“6.875”
stroke-miterlimit=“8” stroke-dasharray=“6.875 6.875” fill=“none” fill-rule=“evenodd”/><text fill=“#7F7F7F” font-family=“Times New Roman,Times New Roman_MSFontService,sans-serif” font-style=“italic” font-weight=“700” font-size=“46” transform=“matrix(1 0 0 1 2301.13 1307)”>Snakemake workflow in Conda environment</text><path d=“M1848.5 1036.75 1873.5 1036.75 1873.5 1015.5 1898.5 1058 1873.5 1100.5 1873.5 1079.25 1848.5
1079.25Z” stroke=“#000000” stroke-width=“6.875” stroke-miterlimit=“8” fill-rule=“evenodd”/><path d=“M2234.5 1036.75 2260 1036.75 2260 1015.5
2285.5 1058 2260 1100.5 2260 1079.25 2234.5 1079.25Z” stroke=“#000000”
stroke-width=“6.875” stroke-miterlimit=“8” fill-rule=“evenodd”/><path d=“M2719.5 1036.75 2745 1036.75 2745 1015.5 2770.5 1058 2745 1100.5 2745 1079.25 2719.5 1079.25Z” stroke=“#000000” stroke-width=“6.875” stroke-miterlimit=“8” fill-rule=“evenodd”/><path d=“M1311.5 1038.75 1336.51038.75 1336.5 1017.5 1361.5 1060 1336.5 1102.5 1336.5 1081.25 1311.5 1081.25Z” stroke=“#000000” stroke-width=“6.875” stroke-miterlimit=“8” fill-rule=“evenodd”/><path d=“M1870.5 622.501C1870.5 611.455 1879.45
602.5 1890.5 602.5L2250.5 602.5C2261.55 602.5 2270.5 611.455 2270.5
622.501L2270.5 702.5C2270.5 713.546 2261.55 722.5 2250.5 722.5L1890.5
722.5C1879.45 722.5 1870.5 713.546 1870.5 702.5Z” stroke=“#042433”
stroke-width=“6.875” stroke-miterlimit=“8” fill=“#F2F2F2” fill-rule=“evenodd”/><text font-family=“Times New Roman,Times New Roman_MSFontService,sans-serif” font-weight=“700” font-size=“41” transform=“matrix(1 0 0 1 1989 653)”>Genomes </text><text font-family=“Times New Roman,Times New Roman_MSFontService,sans-serif” font-weight=“700” font-size=“41” transform=“matrix(1 0 0 1 1910.51 702)”>sequence & index</text><path d=“M3.4375-0.000292938 3.44276 61.7812-3.43224 61.7818-3.4375 0.000292938ZM13.7549 57.197 0.0072178584.6982-13.7451 57.1994Z” transform=“matrix(-1 0 0 1 2070.51722.5)”/><path d=“M2771.5 623.5C2771.5 612.455 2780.45 603.5 2791.5603.5L3148.5 603.5C3159.55 603.5 3168.5 612.455 3168.5 623.5L3168.5 703.5C3168.5 714.546 3159.55 723.5 3148.5 723.5L2791.5 723.5C2780.45723.5 2771.5 714.546 2771.5 703.5Z” stroke=“#042433” stroke-width=“6.875”
stroke-miterlimit=“8” fill=“#F2F2F2” fill-rule=“evenodd”/><text font-family=“Times New Roman,Times New Roman_MSFontService,sans-serif” font-weight=“700” font-size=“41” transform=“matrix(1 0 0 1 2815.74 654)”>Genes annotation </text><text font-family=“Times New Roman,Times New Roman_MSFontService,sans-serif” font-style=“italic” font-weight=“700” font-size=“41” transform=“matrix(1 0 0 1 2831.78 703)”>(Oncogenes
list)</text><path d=“M2973.94 723.5 2973.94 784.047 2967.06 784.0472967.06 723.5ZM2984.25 779.464 2970.5 806.964 2956.75 779.464Z”/><path
d=“M1420.5 836C1420.5 820.26 1433.26 807.5 1449 807.5L1788 807.5C1803.74807.5 1816.5 820.26 1816.5 836L1816.5 836C1816.5 851.74 1803.74 864.51788 864.5L1449 864.5C1433.26 864.5 1420.5 851.74 1420.5 836Z”
stroke=“#000000” stroke-width=“6.875” stroke-miterlimit=“8” fill-rule=“evenodd”/><text fill=“#FFFFFF” font-family=“Times New Roman,Times New Roman_MSFontService,sans-serif” font-weight=“700” font-size=“37” transform=“matrix(1 0 0 1 1479.3 848)”>1. Pre </text><text fill=“#FFFFFF” font-family=“Times New Roman,Times New Roman_MSFontService,sans-serif” font-weight=“700” font-size=“37” transform=“matrix(1 0 0 1 1579.56848)”>-</text><text fill=“#FFFFFF” font-family=“Times New Roman,Times New Roman_MSFontService,sans-serif” font-weight=“700” font-size=“37” transform=“matrix(1 0 0 1 1591.59 848)”>processing</text><path d=“M1906.5 836C1906.5 820.26 1919.26 807.5 1935 807.5L2206 807.5C2221.74 807.52234.5 820.26 2234.5 836L2234.5 836C2234.5 851.74 2221.74 864.5 2206864.5L1935 864.5C1919.26 864.5 1906.5 851.74 1906.5 836Z”
stroke=“#000000” stroke-width=“6.875” stroke-miterlimit=“8” fill-rule=“evenodd”/><text fill=“#FFFFFF” font-family=“Times New Roman,Times New Roman_MSFontService,sans-serif” font-weight=“700” font-size=“37” transform=“matrix(1 0 0 1 1969.52 848)”>2. Alignment</text><pathd=“M2338.5 836C2338.5 820.26 2351.26 807.5 2367 807.5L2645 807.5C2660.74 807.5 2673.5 820.26 2673.5 836L2673.5 836C2673.5 851.74 2660.74 8642645 864.5L2367 864.5C2351.26 864.5 2338.5 851.74 2338.5 836Z”
stroke=“#000000” stroke-width=“6.875” stroke-miterlimit=“8” fill-rule=“evenodd”/><text fill=“#FFFFFF” font-family=“Times New Roman,Times New Roman_MSFontService,sans-serif” font-weight=“700” font-size=“37” transform=“matrix(1 0 0 1 2390.45 848)”>3. Site analysis</text><pathd=“M2816.5 836C2816.5 820.26 2829.26 807.5 2845 807.5L3095 807.5C3110.74807.5 3123.5 820.26 3123.5 836L3123.5 836C3123.5 851.74 3110.74 864.5
3095 864.5L2845 864.5C2829.26 864.5 2816.5 851.74 2816.5 836Z”
stroke=“#000000” stroke-width=“6.875” stroke-miterlimit=“8” fill-rule=“evenodd”/><text fill=“#FFFFFF” font-family=“Times New Roman,Times New Roman_MSFontService,sans-serif” font-weight=“700” font-size=“37” transform=“matrix(1 0 0 1 2862.72 848)”>4. Annotation</text><path
d=“M3295.5 941.001C3295.5 908.692 3321.69 882.5 3354 882.5L3643
882.5C3675.31 882.5 3701.5 908.692 3701.5 941.001L3701.5 1175C3701.5
1207.31 3675.31 1233.5 3643 1233.5L3354 1233.5C3321.69 1233.5 3295.5
1207.31 3295.5 1175Z” stroke=“#042433” stroke-width=“6.875” stroke-miterlimit=“8” fill=“#F2F2F2” fill-rule=“evenodd”/><text font-family=“Times New Roman,Times New Roman_MSFontService,sans-serif” font-weight=“700” font-size=“41” transform=“matrix(1 0 0 1 3344.93 974)”>Off</text><text font-family=“Times New Roman,Times New Roman_MSFontService,sans-serif” font-weight=“700” font-size=“41” transform=“matrix(1 0 0 1 3404.51 974)”>-</text><text font-family=“Times New Roman,Times New Roman_MSFontService,sans-serif” font-weight=“700” font-size=“41” transform=“matrix(1 0 0 1 3418.26 974)”>Targetstable</text><text font-family=“Times New Roman,Times New Roman_MSFontService,sans-serif” font-weight=“700” font-size=“41” transform=“matrix(1 0 0 1 3491.02 1023)”>-</text><text font-family=“Times New Roman,Times New Roman_MSFontService,sans-serif” font-weight=“700” font-size=“41” transform=“matrix(1 0 0 1 3357.53 1073)”>Dynamic report</text><text font-family=“Times New Roman,Times New Roman_MSFontService,sans-serif” font-weight=“700” font-size=“41” transform=“matrix(1 0 0 1 3491.02 1122)”>-</text><text font-family=“Times New Roman,Times New Roman_MSFontService,sans-serif” font-weight=“700” font-size=“41” transform=“matrix(1 0 0 1 3455.5 1172)”>Logs</text><pathd=“M3344.5 836C3344.5 820.26 3357.26 807.5 3373 807.5L3624 807.5C3639.74
807.5 3652.5 820.26 3652.5 836L3652.5 836C3652.5 851.74 3639.74 864.5
3624 864.5L3373 864.5C3357.26 864.5 3344.5 851.74 3344.5 836Z”
stroke=“#000000” stroke-width=“6.875” stroke-miterlimit=“8” fill-rule=“evenodd”/><text fill=“#FFFFFF” font-family=“Times New Roman,Times New Roman_MSFontService,sans-serif” font-weight=“700” font-size=“37” transform=“matrix(1 0 0 1 3433.44 848)”>Outputs</text><path d=“M858.5
837C858.5 821.26 871.26 808.5 887 808.5L1137 808.5C1152.74 808.5 1165.5
821.26 1165.5 837L1165.5 837C1165.5 852.74 1152.74 865.5 1137 865.5L887
865.5C871.26 865.5 858.5 852.74 858.5 837Z” stroke=“#000000” stroke-width=“6.875” stroke-miterlimit=“8” fill-rule=“evenodd”/><text fill=“#FFFFFF” font-family=“Times New Roman,Times New Roman_MSFontService,sans-serif” font-weight=“700” font-size=“37” transform=“matrix(1 0 0 1 960.201 849)”>Inputs</text><text font-family=“Times New Roman,Times New Roman_MSFontService,sans-serif” font-weight=“700” font-size=“41” transform=“matrix(1 0 0 1 873.867 952)”>Sample Data Sheet</text><text fill=“#A6A6A6” font-family=“Times New Roman,Times New Roman_MSFontService,sans-serif” font-weight=“700” font-size=“32” transform=“matrix(1 0 0 1 873.867 992)”>Sample</text><text fill=“#A6A6A6” font-family=“Times New Roman,Times New Roman_MSFontService,sans-serif” font-weight=“700” font-size=“32” transform=“matrix(1 0 0 1 975.846 992)”>-</text><text fill=“#A6A6A6”
font-family=“Times New Roman,Times New Roman_MSFontService,sans-serif” font-weight=“700” font-size=“32” transform=“matrix(1 0 0 1 986.731
992)”>specific parameters</text><path d=“M3216.5 1034.75 3242 1034.75 3242 1013.5 3267.5 1056 3242 1098.5 3242 1077.25 3216.5 1077.25Z”
stroke=“#000000” stroke-width=“6.875” stroke-miterlimit=“8” fill-rule=“evenodd”/></g></svg>
~~~

**Figure S1:**
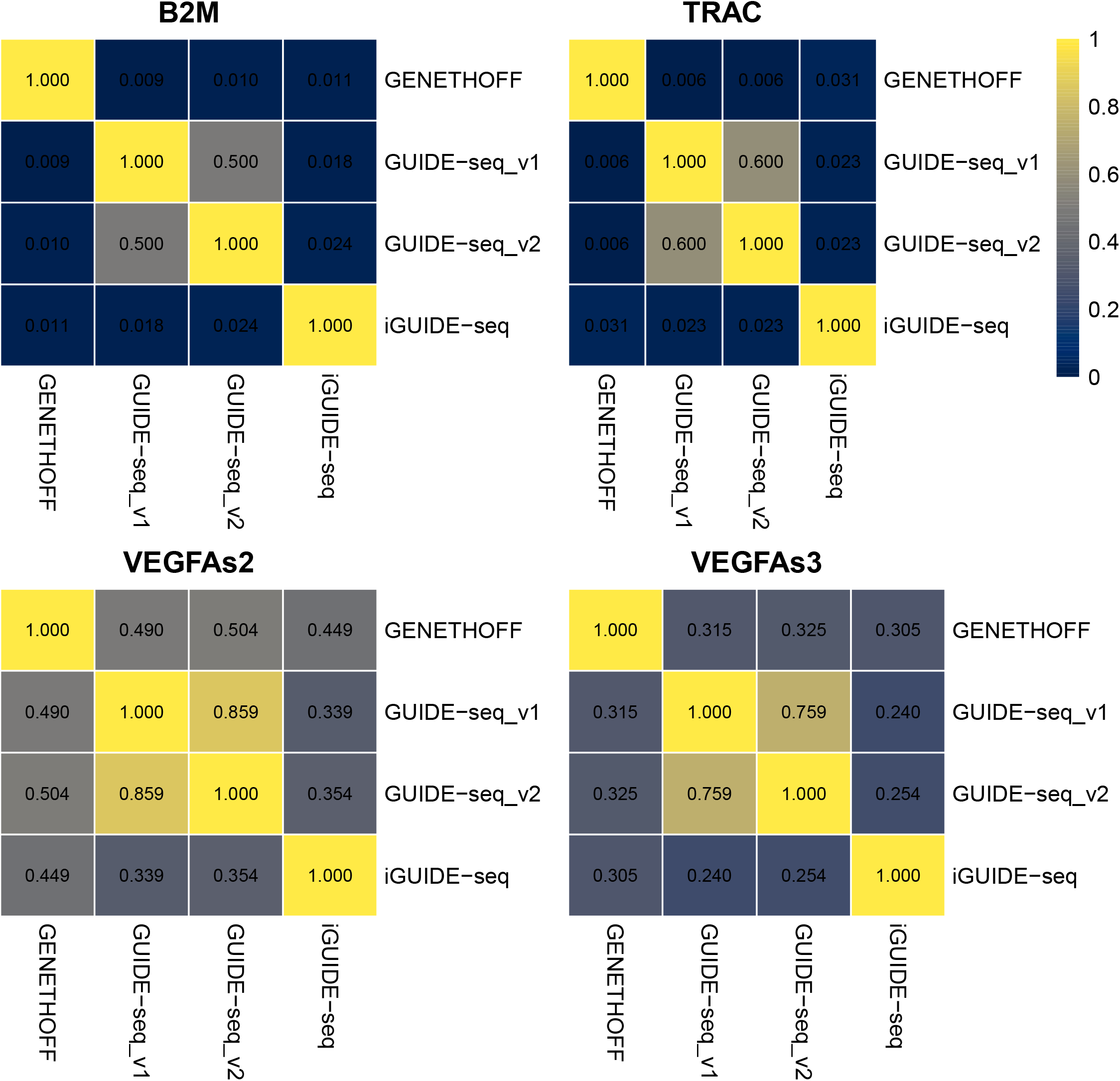
Jaccard coefficient matrix

**Figure S2:**
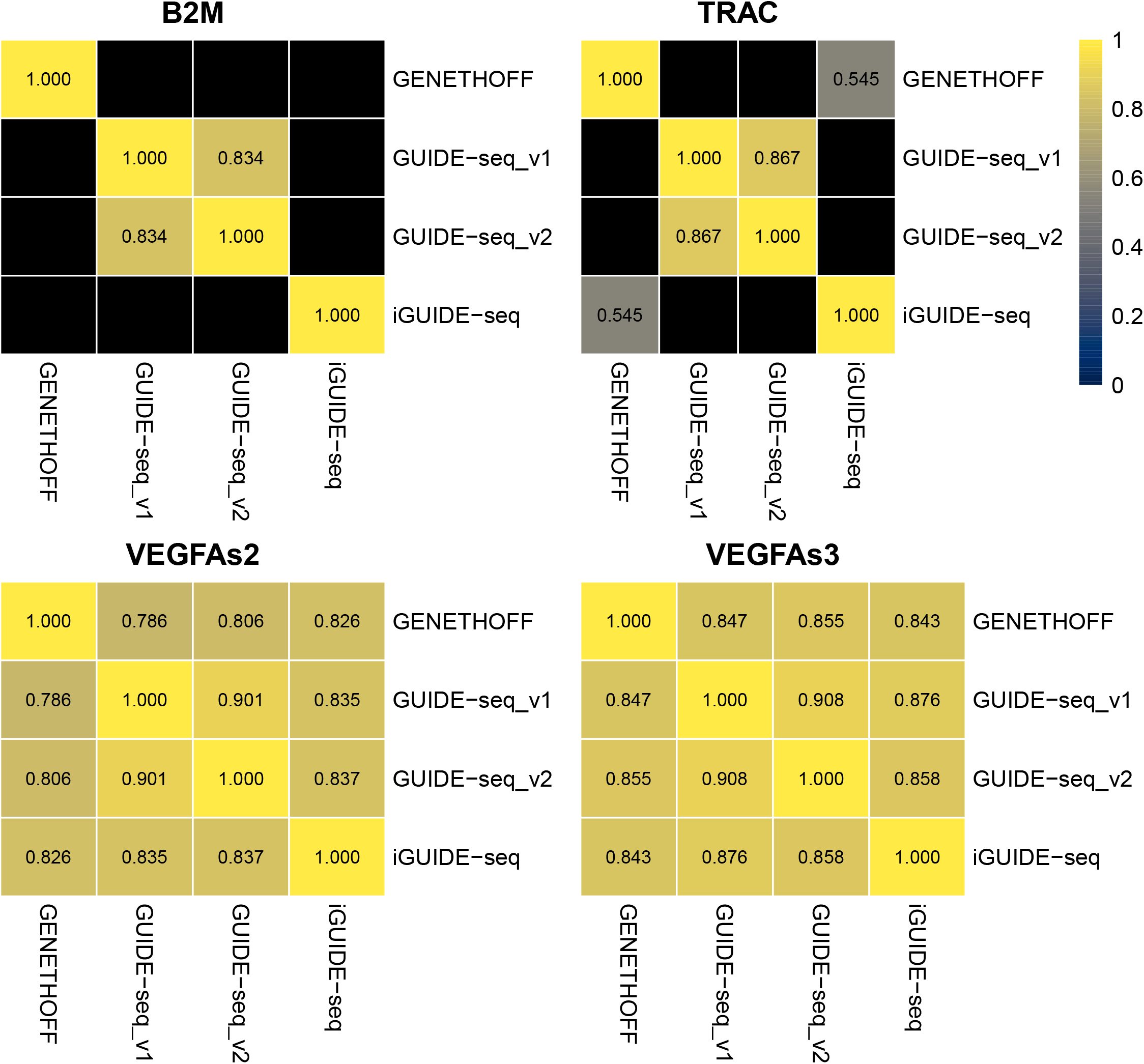
Kendall correlation matrix (shared sites only). When a single site is shared, no correlation can be calculated (black cells)

**Figure S3:**
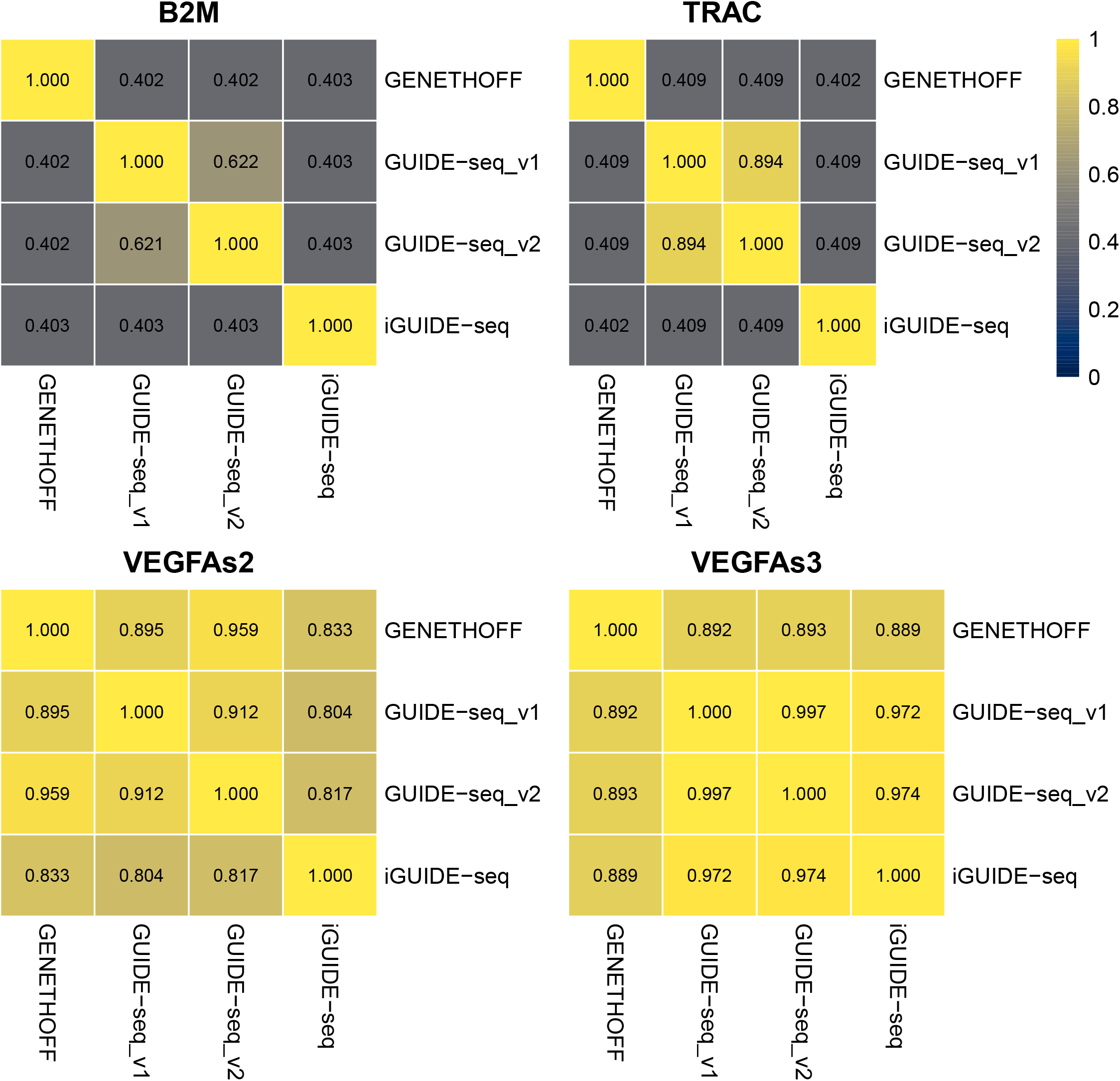
Rank Biased Overlap (RBO) matrix (Persistence parameter p=0.8)

## Notes

https://github.com/gcorre/GENETHOFF

### Competing Interest Statement

The authors have declared no competing interest.

### Summary of Updates

This version has been updated to comply with the new Bioinformatics journal format.

