## supplement_file1 for "GENETH’OFF: a flexible workflow for genome-wide profiling of CRISPR/Cas off-targets": VEGFA_s1_K562_neg_offtargets.html

table output


| Position | cut offset | Alignment | UMIs | UMIs (%) | Reads | Edits crRNA | Edits pam | Symbol | Feature | Predicted | PCR | Oncogene |
| --- | --- | --- | --- | --- | --- | --- | --- | --- | --- | --- | --- | --- |
| chr6:43769560 | -1 | gRNA: GGGTGGGGGGAGTTTGCTCC NGG  gDNA: .................... T.. | 445 | 25.7 | 10295 | 0 | 0 |  |  | yes | - |  |
| chr15:65345199 | -2 | gRNA: GGGTGGGGGGAGTTTGCTCC NGG  gDNA: ..A...A............. T.. | 251 | 14.5 | 4958 | 2 | 0 | IGDCC3 | intron | yes | - |  |
| chr17:41640076 | -2 | gRNA: GGGTGGGGGGAGTTTGCTCC NGG  gDNA: TA....A.....C....... T.. | 223 | 12.9 | 4652 | 4 | 0 | KRT42P | exon | yes | - |  |
| chr12:131205653 | -1 | gRNA: GGGTGGGGGGAGTTTGCTCC NGG  gDNA: ...A...T............ T.. | 73 | 4.2 | 1490 | 2 | 0 | LINC01257 | intron | yes | - |  |
| chr22:19710940 | -2 | gRNA: GGGTGGGGGGAGTTTGCTCC NGG  gDNA: .A.G...A.C.......... A.. | 33 | 1.9 | 363 | 4 | 0 | ENSG00000304096 | intron | yes | - |  |
| chr10:122971899 | -1 | gRNA: GGGTGGGGGGAGTTTGCTCC NGG  gDNA: A.C...A..........C.. A.. | 20 | 1.2 | 296 | 4 | 0 | PSTK | intron | yes | - |  |
| chr17:49240177 | -2 | gRNA: GGGTGGGGGGAGTTTGCTCC NGG  gDNA: T......-....C....... A.. | 18 | 1.0 | 569 | 3 | 0 | FLJ40194 | intron | yes | - |  |
| chr12:1878910 | 0 | gRNA: GGGTGGGGGGAGTTTGCTCC NGG  gDNA: C..G..A............. T.. | 15 | 0.9 | 231 | 3 | 0 | CACNA2D4 | exon | yes | - |  |
| chr1:98882095 | -1 | gRNA: GGGTGGGGGGAGTTTGCTCC NGG  gDNA: ...GA....A.......... T.. | 13 | 0.8 | 446 | 3 | 0 |  |  | yes | - |  |
| chrX:134220952 | 28 | gRNA: GGGTGGGGGGAGTTTGCTCC NGG  gDNA: .........T--.C.T...T ATT | 13 | 0.8 | 104 | 6 | 2 |  |  | no | - |  |
| chr10:28930074 | -15 | gRNA: GGGTGGGGGGAGTTTGCTCC NGG  gDNA: A.....AA--.......... ACT | 10 | 0.6 | 155 | 5 | 2 | ENSG00000298315   LINC01517 | intron  intron | no | - |  |
| chr11:34469490 | -25 | gRNA: GGGTGGGGGGAGTT-TGCTCC NGG  gDNA: AA...T.T......G....G. AT. | 10 | 0.6 | 603 | 6 | 1 | CAT | intron | no | - |  |
| chr18:78450126 | 1 | gRNA: GGGTGGGGGGAGTTTGCTCC NGG  gDNA: ...A.T.AT....CA..... CT. | 10 | 0.6 | 364 | 6 | 1 |  |  | no | - |  |
| chr2:2592289 | 34 | gRNA: GGGTGGGGGGAGTTTGCTCC NGG  gDNA: ......T.--.....C.C.A G.C | 9 | 0.5 | 110 | 6 | 1 | ENSG00000290112 | intron | yes | - |  |
| chr9:76420555 | 36 | gRNA: GGGTGGGGGGAGTTTGCTCC NGG  gDNA: .....T..T.-.C...TG.. T.C | 9 | 0.5 | 161 | 6 | 1 | GCNT1   H3P32 | intron  exon | no | - |  |
| chr2:38076679 | -18 | gRNA: GGGTGGGGGGAGTTTGCTCC NGG  gDNA: ...GC.......-CG...A. TTT | 8 | 0.5 | 93 | 6 | 2 | CYP1B1   CYP1B1-AS1 | intron  intron | no | - |  |
| chr20:32801821 | -1 | gRNA: GGGTGGGGGGAGTTTGCTCC NGG  gDNA: A......C...--.....TG A.C | 8 | 0.5 | 94 | 6 | 1 | DNMT3B | intron | no | - |  |
| chr5:113887358 | 11 | gRNA: GGGTGGGGGGAGTTTGCTCC NGG  gDNA: .....AAT..-..C.C.... A.T | 8 | 0.5 | 337 | 6 | 1 |  |  | no | - |  |
| chr5:139883438 | -1 | gRNA: GGGTGGGGGGAGTTTGCTCC NGG  gDNA: ...--....C.......... T.. | 8 | 0.5 | 231 | 3 | 0 | NRG2 | intron | yes | - |  |
| chr8:17045344 | 33 | gRNA: GGGTGGGGGG-AGTTTGCTCC NGG  gDNA: ....AT....C..GA...... CT. | 8 | 0.5 | 79 | 5 | 1 | MICU3 | intron | yes | - |  |
| chr10:33838832 | 4 | gRNA: GGGTGGGGGGAGTTTGCTCC NGG  gDNA: AA.CA....A.....-.... CTT | 7 | 0.4 | 161 | 6 | 2 |  |  | no | - |  |
| chr12:2820142 | -14 | gRNA: GGGTGGGGGGAGTTTGCTCC NGG  gDNA: .T....T.C...C..--... GCT | 7 | 0.4 | 37 | 6 | 2 | ITFG2 | exon | no | - |  |
| chr19:41898266 | -41 | gRNA: GGGTGGGGGGAGTTTGCTCC NGG  gDNA: A..A.....T..AA....T. TA. | 7 | 0.4 | 84 | 6 | 1 | ARHGEF1 | exon | no | - |  |
| chr3:9533004 | 1 | gRNA: GGGTGGG-GGGAGTTTGCTCC NGG  gDNA: .......CA..T.AAA..... TTT | 7 | 0.4 | 71 | 6 | 2 | LHFPL4 | intron | no | - |  |
| chr3:10193539 | -28 | gRNA: GGGTGGGGGGAGTTTGCTCC NGG  gDNA: A......A...--.....TG AAC | 7 | 0.4 | 30 | 6 | 2 | IRAK2 | intron | no | - |  |
| chr7:134074471 | 3 | gRNA: GGGTGGGGGGAGTTTGCTCC NGG  gDNA: ..........--..G..AT. AAT | 7 | 0.4 | 181 | 5 | 2 | ENSG00000294998 | intron | no | - |  |
| chr8:134800535 | 4 | gRNA: GGGTGGGGGGAGTTTGCTCC NGG  gDNA: ...A..T..AT....A..T. A.C | 7 | 0.4 | 195 | 6 | 1 | MIR30B   ENSG00000289405 | exon  intron | no | - |  |
| chr11:87544954 | -11 | gRNA: GGGTGGGGGGAGTTTGCTCC NGG  gDNA: C......A..T.....GGT. AT. | 6 | 0.3 | 36 | 6 | 1 | ENSG00000302190 | intron | yes | - |  |
| chr12:81500313 | -1 | gRNA: GGGTGGGGGGAGTTTGCTCC NGG  gDNA: ...A.....T...G.AG.A. TA. | 6 | 0.3 | 227 | 6 | 1 | ENSG00000258162   PPFIA2 | intron  intron | no | - |  |
| chr15:72211603 | 10 | gRNA: GGGTGGGGGGA--GTTTGCTCC NGG  gDNA: A......A...CA.....GG.. A.. | 6 | 0.3 | 40 | 6 | 0 | PKM | intron | yes | - |  |
| chr17:6396060 | 24 | gRNA: GGGTGGGGGGAGTTTGCTCC NGG  gDNA: .C....TCA.G.......T. CCC | 6 | 0.3 | 34 | 6 | 2 | AIPL1 | intron | no | - |  |
| chr2:151338765 | -32 | gRNA: GGGTGGGGGGAGTT-TGCTCC NGG  gDNA: ..T......A..AAG....G. A.A | 6 | 0.3 | 152 | 6 | 1 |  |  | no | - |  |
| chr5:31846195 | 13 | gRNA: GGGTGGGGGGAGTTTGCTCC NGG  gDNA: CA...A..C...A....G.. ACT | 6 | 0.3 | 44 | 6 | 2 | PDZD2 | intron | no | - |  |
| chr10:53943348 | 34 | gRNA: GGGTGGGGGGAGTTTGCTCC NGG  gDNA: ..T..T..T.-..G...... TAT | 5 | 0.3 | 26 | 5 | 2 | PCDH15 | intron | no | - |  |
| chr12:124347102 | -32 | gRNA: GGGTGGGGGGAGTTTGCTCC NGG  gDNA: ..C....T.C...A-....A C.C | 5 | 0.3 | 68 | 6 | 1 | NCOR2 | intron | no | - |  |
| chr14:42007698 | 35 | gRNA: GGGTGGGGGGAGTTTGCTCC NGG  gDNA: .......--T.T...A..T. AAA | 5 | 0.3 | 126 | 6 | 2 |  |  | yes | - |  |
| chr14:60652838 | -36 | gRNA: GGGTGGGGGGAGTTTGCTCC NGG  gDNA: ..A.TT.......---.... GCA | 5 | 0.3 | 240 | 6 | 2 | SIX1 | exon | no | - |  |
| chr15:99100068 | -1 | gRNA: GGGTGGGGGGAG-TTTGCTCC NGG  gDNA: TA...AA.....A.....A.. CCT | 5 | 0.3 | 40 | 6 | 2 | SYNM   SYNM-AS1 | intron  intron | no | - |  |
| chr16:15783300 | -26 | gRNA: GGGTGGGGGGAGTTTG-CTCC NGG  gDNA: ..A.TCT........TT.... TT. | 5 | 0.3 | 164 | 6 | 1 | MYH11 | exon | no | - |  |
| chr2:64971254 | -9 | gRNA: GGGTGGGGG-GAGTTTGCTCC NGG  gDNA: ..T......A...A.....TG A.C | 5 | 0.3 | 61 | 5 | 1 | LINC02245 | intron | no | - |  |
| chr22:22635670 | -23 | gRNA: GGGTGGGGGGAGTTTGCTCC NGG  gDNA: T..A.AT...-.......G. ATT | 5 | 0.3 | 160 | 6 | 2 | BCRP4   ENSG00000290990 | intron  exon | no | - |  |
| chr22:28802508 | 0 | gRNA: GGGTGGGGGGAGT-TTGCTCC NGG  gDNA: A.A..T......AC..A.... G.. | 5 | 0.3 | 185 | 6 | 0 | ENSG00000226471 | intron | yes | - |  |
| chr6:84789447 | 8 | gRNA: GGGTGGGGGGAGTTTGCTCC NGG  gDNA: ........A...A.G.TAG. TAA | 5 | 0.3 | 350 | 6 | 2 | ENSG00000307486 | intron | no | - |  |
| chr1:187031500 | 22 | gRNA: GGGTGGGGGGAGTTTGCTCC NGG  gDNA: .TT..TT........T..T. TTC | 4 | 0.2 | 131 | 6 | 2 |  |  | no | - |  |
| chr1:233021607 | -1 | gRNA: GGGTGG-GGGGAGTTTGCTCC NGG  gDNA: ...A..A.......C...... A.. | 4 | 0.2 | 194 | 3 | 0 | PCNX2 | intron | yes | - |  |
| chr10:21500431 | -28 | gRNA: GGGTGGGGGGAGTTTG-CTCC NGG  gDNA: .CTA.....C......G..T. CCC | 4 | 0.2 | 155 | 6 | 2 | ENSG00000308031 | intron | no | - |  |
| chr10:79603404 | -17 | gRNA: GGGTGGGGGGAGTTTGCTCC NGG  gDNA: .A....T.-...G.G....T CA. | 4 | 0.2 | 41 | 6 | 1 | ENSG00000295647 | exon | no | - |  |
| chr10:80435205 | 5 | gRNA: GGGTGGGGGGAGTTTG-CTCC NGG  gDNA: .....A-CA....G..A.... ATC | 4 | 0.2 | 148 | 6 | 2 | PRXL2A | exon | no | - |  |
| chr11:9714629 | -43 | gRNA: GGGTGGGG-GGAGTTTGCTCC NGG  gDNA: .......AA...T....GC.A ... | 4 | 0.2 | 58 | 6 | 3 | SWAP70 | intron | no | - |  |
| chr11:115362111 | -28 | gRNA: GGGTGGGGGGAGTTTGCTCC NGG  gDNA: ....AAA..T......T.T. ATT | 4 | 0.2 | 133 | 6 | 2 | CADM1 | intron | no | - |  |
| chr11:118366802 | 33 | gRNA: GGGTGGGGGGAGTTTGCTCC NGG  gDNA: ..C....C.C...G-....G C.C | 4 | 0.2 | 124 | 6 | 1 | UBE4A | intron | no | - |  |
| chr12:113465819 | 46 | gRNA: GGGTGGGGGGAGTTTGCTCC NGG  gDNA: ..A...T...C..G.CT... ... | 4 | 0.2 | 139 | 6 | 3 | LHX5 | intron | yes | - |  |
| chr13:80601083 | 25 | gRNA: GGGTGGGGGGAGT-TTGCTCC NGG  gDNA: A......A....GA.....TG A.C | 4 | 0.2 | 34 | 6 | 1 |  |  | no | - |  |
| chr14:29942785 | 20 | gRNA: GGGTGGGGGGAGTTTGCTCC NGG  gDNA: .T.AA.......G.G..A.. TCC | 4 | 0.2 | 31 | 6 | 2 | PRKD1   ENSG00000257904 | intron  intron | no | - |  |
| chr16:17839812 | -38 | gRNA: GGGTGGG--GGGAGTTT-GCTCC NGG  gDNA: .......TA..A...GAC..... ATT | 4 | 0.2 | 35 | 6 | 2 | ENSG00000259929 | intron | no | - |  |
| chr18:13572629 | -22 | gRNA: GGGTGGGG-GGAG-TTTGCTCC NGG  gDNA: ...A..A.A....A......T. TA. | 4 | 0.2 | 249 | 5 | 1 | LDLRAD4 | intron | no | - |  |
| chr19:36777726 | -23 | gRNA: GGGTGGGGGGAGTTTGCTCC NGG  gDNA: .....A..T.-.CA...A.. T.T | 4 | 0.2 | 156 | 6 | 1 | ZNF790-AS1 | intron | no | - |  |
| chr19:41971674 | 22 | gRNA: GGGTGGGGGGAGTTTGCTCC NGG  gDNA: .....--CT...CC...... A.T | 4 | 0.2 | 66 | 6 | 1 | ENSG00000285505   ATP1A3 | intron  intron | no | - |  |
| chr22:46672939 | -11 | gRNA: GGGTGGG--GGGAGTTTGCTCC NGG  gDNA: ....C..TG.....AC.C.... A.A | 4 | 0.2 | 63 | 6 | 1 | GRAMD4 | exon | no | - |  |
| chr3:15422067 | 23 | gRNA: GGGTGGGGGGAGTTTGCTCC NGG  gDNA: .A...CT....-..AA.... CAC | 4 | 0.2 | 31 | 6 | 2 | METTL6 | intron | no | - |  |
| chr3:44644753 | -46 | gRNA: GGGTGGGGGGAGTTTGCTCC NGG  gDNA: ..C....T.C...G-....A ... | 4 | 0.2 | 243 | 6 | 3 | ZNF197   ZKSCAN7-AS1 | exon  intron | no | - |  |
| chr4:62551937 | -5 | gRNA: GGGTGGGGGGAGTTTGCTCC NGG  gDNA: A..CCT.........C...A GAC | 4 | 0.2 | 32 | 6 | 2 |  |  | no | - |  |
| chr5:7067045 | -3 | gRNA: GGGTGGGGGGAGTTTGCTCC NGG  gDNA: ....--.........A.... T.. | 4 | 0.2 | 64 | 3 | 0 | LINC02196   ENSG00000287790 | intron  exon | yes | - |  |
| chr5:39091229 | 2 | gRNA: GGGTGGGG-GGAGTTTGCTCC NGG  gDNA: .......AA....GG..A..A A.A | 4 | 0.2 | 61 | 6 | 1 |  |  | no | - |  |
| chr6:14316142 | -2 | gRNA: GGGTGGGGG--GAGTTTGCTCC NGG  gDNA: A........TA........... A.. | 4 | 0.2 | 109 | 3 | 0 | ENSG00000286277 | intron | yes | - |  |
| chr6:21334699 | 39 | gRNA: GGGTGGGGGGAGTTTGCTCC NGG  gDNA: A.T.....A..-....T.T. GA. | 4 | 0.2 | 158 | 6 | 1 |  |  | no | - |  |
| chr6:99229976 | -46 | gRNA: GGGTGGGGGGAGTTTGCTCC NGG  gDNA: .T...A...-.....--.G. ... | 4 | 0.2 | 14 | 6 | 3 |  |  | no | - |  |
| chr6:108834808 | -27 | gRNA: GGGTGGGGGGAGTTTGCTCC NGG  gDNA: ...G.A.....--..T..T. ATT | 4 | 0.2 | 69 | 6 | 2 | LINC00222 | intron | no | - |  |
| chr6:123557797 | 1 | gRNA: GGGTGGGGGGAGT--TTGCTCC NGG  gDNA: ..........G..GC....C.. CCA | 4 | 0.2 | 48 | 4 | 2 | TRDN | intron | no | - |  |
| chr7:30668290 | 33 | gRNA: GGGTGGGGGGAGTTTGCTCC NGG  gDNA: A......A..-.C..C.... T.A | 4 | 0.2 | 73 | 5 | 1 | CRHR2   ENSG00000305335 | intron  intron | no | - |  |
| chr8:27896827 | 31 | gRNA: GGGTGGGGGGAGTTTGCTCC NGG  gDNA: ..C....C.C...G-....A C.C | 4 | 0.2 | 34 | 6 | 1 | SCARA5 | intron | no | - |  |
| chr9:6348281 | -12 | gRNA: GGGTGGGGGGAGTTTGCTCC NGG  gDNA: ..C....T.C...G-....A CAC | 4 | 0.2 | 37 | 6 | 2 |  |  | no | - |  |
| chrX:261577 | -3 | gRNA: GGGTGGGGGGAGTTTGCTCC NGG  gDNA: ...A..CC.......C..TA TTC | 4 | 0.2 | 88 | 6 | 2 |  |  | no | - |  |
| chrY:261577 | -3 | gRNA: GGGTGGGGGGAGTTTGCTCC NGG  gDNA: ...A..CC.......C..TA TTC | 4 | 0.2 | 78 | 6 | 2 |  |  | no | - |  |
| chrY:11208206 | -8 | gRNA: GGGTGGGGGGAGTTTGCTCC NGG  gDNA: ..C....T.C...G-....A T.T | 4 | 0.2 | 23 | 6 | 1 | ENSG00000291032 | intron | no | - |  |
| chr1:31492716 | -31 | gRNA: GGGTGGGGGGAGTTTGCTCC NGG  gDNA: ....T...CT....AT..G. A.C | 3 | 0.2 | 146 | 6 | 1 |  |  | no | - |  |
| chr1:44994950 | -25 | gRNA: GGGTGGGGGGAGTTTGCTCC NGG  gDNA: ..C....T.C...G-....A C.C | 3 | 0.2 | 108 | 6 | 1 | ENSG00000300507 | intron | no | - |  |
| chr1:53086541 | -6 | gRNA: GGGTGGGG-GGAGTTTGCTCC NGG  gDNA: ....CT..A..G.C....A.. AT. | 3 | 0.2 | 155 | 6 | 1 |  |  | no | - |  |
| chr1:208902624 | 39 | gRNA: GGGTGGGGGGAGTTTGCTCC NGG  gDNA: ..C....T.C...G-....A CCC | 3 | 0.2 | 24 | 6 | 2 |  |  | no | - |  |
| chr1:223339480 | 46 | gRNA: GGGTGGGGGGAGTTTGCTCC NGG  gDNA: ..T...A....--.G.G... ... | 3 | 0.2 | 14 | 6 | 3 | SUSD4 | intron | no | - |  |
| chr1:237947974 | 26 | gRNA: GGGTGGGGGGAGTTTGCTCC NGG  gDNA: ...A......----.T.... ATT | 3 | 0.2 | 15 | 6 | 2 | MTCYBP15   ENSG00000303841 | exon  intron | no | - |  |
| chr12:82645525 | 34 | gRNA: GGGTGGGGGGA--GTTTGCTCC NGG  gDNA: ....T...CC.AG.C....... TTT | 3 | 0.2 | 61 | 6 | 2 |  |  | no | - |  |
| chr15:31604799 | -40 | gRNA: GGGTGGGGGGAGTTTGCTCC NGG  gDNA: ........T...-AA....A ACA | 3 | 0.2 | 37 | 5 | 2 | OTUD7A | intron | no | - |  |
| chr15:39762879 | 24 | gRNA: GGGTGGGGGGAGTTTGCTCC NGG  gDNA: .T...T..-T......T.T. CAA | 3 | 0.2 | 18 | 6 | 2 | FSIP1 | intron | no | - |  |
| chr15:66070143 | -15 | gRNA: GGGTGGGGGGAGTTTGCTCC NGG  gDNA: ......T---...C.....A CCC | 3 | 0.2 | 129 | 6 | 2 | MEGF11 | intron | no | - |  |
| chr16:54608486 | 20 | gRNA: GGGTGGGGGGAGTTTGCTCC NGG  gDNA: ....CT..A..T...TT... C.A | 3 | 0.2 | 250 | 6 | 1 |  |  | no | - |  |
| chr17:18361521 | 13 | gRNA: GGGTGGGGGGAGTTTGCTCC NGG  gDNA: ..C....T.C...G-....A CAT | 3 | 0.2 | 16 | 6 | 2 | SHMT1 | intron | no | - |  |
| chr18:12394631 | -24 | gRNA: GGGTGGGGGGAGTTTGCTCC NGG  gDNA: ..CA.TT....-.....C.. TT. | 3 | 0.2 | 43 | 6 | 1 | ENSG00000296872 | intron | no | - |  |
| chr19:11736964 | -6 | gRNA: GGGTGGGGGGAGTTTGCTCC NGG  gDNA: .......A.......C..GG TAT | 3 | 0.2 | 15 | 4 | 2 | ZNF823 | intron | yes | - |  |
| chr2:20553435 | 11 | gRNA: GGGTGGGGGGAGTTTGCTCC NGG  gDNA: T.A...T..C...GG..... T.A | 3 | 0.2 | 73 | 6 | 1 |  |  | no | - |  |
| chr2:129623117 | 8 | gRNA: GGGTGGG-GGGAGTTTGCTCC NGG  gDNA: .....T.A.....CA--.... AA. | 3 | 0.2 | 29 | 6 | 1 |  |  | no | - |  |
| chr2:216303984 | 16 | gRNA: GGGTGGGG-GGAGTTTGCTCC NGG  gDNA: .......TA....G..--... A.T | 3 | 0.2 | 22 | 5 | 1 | ENSG00000233581   MARCHF4 | intron  intron | no | - |  |
| chr20:46758134 | 1 | gRNA: GGGTGGGGGGAGTTTGCTCC NGG  gDNA: ...AT...A....C..-..A T.A | 3 | 0.2 | 223 | 6 | 1 |  |  | no | - |  |
| chr20:47346158 | -15 | gRNA: GGGTGGGGGGAGTTTGCTCC NGG  gDNA: ........---..C.TG... CT. | 3 | 0.2 | 62 | 6 | 1 | ZMYND8 | intron | no | - |  |
| chr21:45061319 | -9 | gRNA: GGGTGGGGGGAGTTTG-CTCC NGG  gDNA: .....CT..T...G..G...A GTA | 3 | 0.2 | 17 | 6 | 2 | ENSG00000295969   SSR4P1 | intron  intron | no | - |  |
| chr3:13447304 | -37 | gRNA: GGGTGGG-GGGAGTTTGCTCC NGG  gDNA: .A.....T.....---....A G.C | 3 | 0.2 | 16 | 6 | 1 |  |  | no | - |  |
| chr3:136757416 | 20 | gRNA: GGGTGGGGGGAGTTTGCTCC NGG  gDNA: ..C....C.C...G-....A CAC | 3 | 0.2 | 20 | 6 | 2 | STAG1-DT | intron | no | - |  |
| chr4:121510508 | -41 | gRNA: GGGTGGGGGGAGTTTGCTCC NGG  gDNA: .....TA..---..A..... ACA | 3 | 0.2 | 261 | 6 | 2 | ENSG00000308636 | intron | no | - |  |
| chr4:133491051 | -31 | gRNA: GGGTGGGGGGAGTTTGCTCC NGG  gDNA: ............GGG..AGT GAT | 3 | 0.2 | 176 | 6 | 2 |  |  | no | - |  |
| chr5:32945168 | -1 | gRNA: GGGTGGGGGGAGTTTGCTCC NGG  gDNA: .C........T......... C.. | 3 | 0.2 | 69 | 2 | 0 | ENSG00000250697 | intron | yes | - |  |
