## supplement_file1 for "GENETH’OFF: a flexible workflow for genome-wide profiling of CRISPR/Cas off-targets": jquery.dataTables.min.js

```
/*! DataTables 1.13.6
 * ©2008-2023 SpryMedia Ltd - datatables.net/license
 */
!function(n){"use strict";var a;"function"==typeof define&&define.amd?define(["jquery"],function(t){return n(t,window,document)}):"object"==typeof exports?(a=require("jquery"),"undefined"==typeof window?module.exports=function(t,e){return t=t||window,e=e||a(t),n(e,t,t.document)}:n(a,window,window.document)):window.DataTable=n(jQuery,window,document)}(function(P,j,v,H){"use strict";function d(t){var e=parseInt(t,10);return!isNaN(e)&&isFinite(t)?e:null}function l(t,e,n){var a=typeof t,r="string"==a;return"number"==a||"bigint"==a||!!h(t)||(e&&r&&(t=$(t,e)),n&&r&&(t=t.replace(q,"")),!isNaN(parseFloat(t))&&isFinite(t))}function a(t,e,n){var a;return!!h(t)||(h(a=t)||"string"==typeof a)&&!!l(t.replace(V,"").replace(/<script/i,""),e,n)||null}function m(t,e,n,a){var r=[],o=0,i=e.length;if(a!==H)for(;o<i;o++)t[e[o]][n]&&r.push(t[e[o]][n][a]);else for(;o<i;o++)r.push(t[e[o]][n]);return r}function f(t,e){var n,a=[];e===H?(e=0,n=t):(n=e,e=t);for(var r=e;r<n;r++)a.push(r);return a}function _(t){for(var e=[],n=0,a=t.length;n<a;n++)t[n]&&e.push(t[n]);return e}function s(t,e){return-1!==this.indexOf(t,e=e===H?0:e)}var p,e,t,w=function(t,v){if(w.factory(t,v))return w;if(this instanceof w)return P(t).DataTable(v);v=t,this.$=function(t,e){return this.api(!0).$(t,e)},this._=function(t,e){return this.api(!0).rows(t,e).data()},this.api=function(t){return new B(t?ge(this[p.iApiIndex]):this)},this.fnAddData=function(t,e){var n=this.api(!0),t=(Array.isArray(t)&&(Array.isArray(t[0])||P.isPlainObject(t[0]))?n.rows:n.row).add(t);return e!==H&&!e||n.draw(),t.flatten().toArray()},this.fnAdjustColumnSizing=function(t){var e=this.api(!0).columns.adjust(),n=e.settings()[0],a=n.oScroll;t===H||t?e.draw(!1):""===a.sX&&""===a.sY||Qt(n)},this.fnClearTable=function(t){var e=this.api(!0).clear();t!==H&&!t||e.draw()},this.fnClose=function(t){this.api(!0).row(t).child.hide()},this.fnDeleteRow=function(t,e,n){var a=this.api(!0),t=a.rows(t),r=t.settings()[0],o=r.aoData[t[0][0]];return t.remove(),e&&e.call(this,r,o),n!==H&&!n||a.draw(),o},this.fnDestroy=function(t){this.api(!0).destroy(t)},this.fnDraw=function(t){this.api(!0).draw(t)},this.fnFilter=function(t,e,n,a,r,o){var i=this.api(!0);(null===e||e===H?i:i.column(e)).search(t,n,a,o),i.draw()},this.fnGetData=function(t,e){var n,a=this.api(!0);return t!==H?(n=t.nodeName?t.nodeName.toLowerCase():"",e!==H||"td"==n||"th"==n?a.cell(t,e).data():a.row(t).data()||null):a.data().toArray()},this.fnGetNodes=function(t){var e=this.api(!0);return t!==H?e.row(t).node():e.rows().nodes().flatten().toArray()},this.fnGetPosition=function(t){var e=this.api(!0),n=t.nodeName.toUpperCase();return"TR"==n?e.row(t).index():"TD"==n||"TH"==n?[(n=e.cell(t).index()).row,n.columnVisible,n.column]:null},this.fnIsOpen=function(t){return this.api(!0).row(t).child.isShown()},this.fnOpen=function(t,e,n){return this.api(!0).row(t).child(e,n).show().child()[0]},this.fnPageChange=function(t,e){t=this.api(!0).page(t);e!==H&&!e||t.draw(!1)},this.fnSetColumnVis=function(t,e,n){t=this.api(!0).column(t).visible(e);n!==H&&!n||t.columns.adjust().draw()},this.fnSettings=function(){return ge(this[p.iApiIndex])},this.fnSort=function(t){this.api(!0).order(t).draw()},this.fnSortListener=function(t,e,n){this.api(!0).order.listener(t,e,n)},this.fnUpdate=function(t,e,n,a,r){var o=this.api(!0);return(n===H||null===n?o.row(e):o.cell(e,n)).data(t),r!==H&&!r||o.columns.adjust(),a!==H&&!a||o.draw(),0},this.fnVersionCheck=p.fnVersionCheck;var e,y=this,D=v===H,_=this.length;for(e in D&&(v={}),this.oApi=this.internal=p.internal,w.ext.internal)e&&(this[e]=$e(e));return this.each(function(){var r=1<_?be({},v,!0):v,o=0,t=this.getAttribute("id"),i=!1,e=w.defaults,l=P(this);if("table"!=this.nodeName.toLowerCase())W(null,0,"Non-table node initialisation ("+this.nodeName+")",2);else{K(e),Q(e.column),C(e,e,!0),C(e.column,e.column,!0),C(e,P.extend(r,l.data()),!0);for(var n=w.settings,o=0,s=n.length;o<s;o++){var a=n[o];if(a.nTable==this||a.nTHead&&a.nTHead.parentNode==this||a.nTFoot&&a.nTFoot.parentNode==this){var u=(r.bRetrieve!==H?r:e).bRetrieve,c=(r.bDestroy!==H?r:e).bDestroy;if(D||u)return a.oInstance;if(c){a.oInstance.fnDestroy();break}return void W(a,0,"Cannot reinitialise DataTable",3)}if(a.sTableId==this.id){n.splice(o,1);break}}null!==t&&""!==t||(t="DataTables_Table_"+w.ext._unique++,this.id=t);var f,d,h=P.extend(!0,{},w.models.oSettings,{sDestroyWidth:l[0].style.width,sInstance:t,sTableId:t}),p=(h.nTable=this,h.oApi=y.internal,h.oInit=r,n.push(h),h.oInstance=1===y.length?y:l.dataTable(),K(r),Z(r.oLanguage),r.aLengthMenu&&!r.iDisplayLength&&(r.iDisplayLength=(Array.isArray(r.aLengthMenu[0])?r.aLengthMenu[0]:r.aLengthMenu)[0]),r=be(P.extend(!0,{},e),r),F(h.oFeatures,r,["bPaginate","bLengthChange","bFilter","bSort","bSortMulti","bInfo","bProcessing","bAutoWidth","bSortClasses","bServerSide","bDeferRender"]),F(h,r,["asStripeClasses","ajax","fnServerData","fnFormatNumber","sServerMethod","aaSorting","aaSortingFixed","aLengthMenu","sPaginationType","sAjaxSource","sAjaxDataProp","iStateDuration","sDom","bSortCellsTop","iTabIndex","fnStateLoadCallback","fnStateSaveCallback","renderer","searchDelay","rowId",["iCookieDuration","iStateDuration"],["oSearch","oPreviousSearch"],["aoSearchCols","aoPreSearchCols"],["iDisplayLength","_iDisplayLength"]]),F(h.oScroll,r,[["sScrollX","sX"],["sScrollXInner","sXInner"],["sScrollY","sY"],["bScrollCollapse","bCollapse"]]),F(h.oLanguage,r,"fnInfoCallback"),L(h,"aoDrawCallback",r.fnDrawCallback,"user"),L(h,"aoServerParams",r.fnServerParams,"user"),L(h,"aoStateSaveParams",r.fnStateSaveParams,"user"),L(h,"aoStateLoadParams",r.fnStateLoadParams,"user"),L(h,"aoStateLoaded",r.fnStateLoaded,"user"),L(h,"aoRowCallback",r.fnRowCallback,"user"),L(h,"aoRowCreatedCallback",r.fnCreatedRow,"user"),L(h,"aoHeaderCallback",r.fnHeaderCallback,"user"),L(h,"aoFooterCallback",r.fnFooterCallback,"user"),L(h,"aoInitComplete",r.fnInitComplete,"user"),L(h,"aoPreDrawCallback",r.fnPreDrawCallback,"user"),h.rowIdFn=A(r.rowId),tt(h),h.oClasses),g=(P.extend(p,w.ext.classes,r.oClasses),l.addClass(p.sTable),h.iInitDisplayStart===H&&(h.iInitDisplayStart=r.iDisplayStart,h._iDisplayStart=r.iDisplayStart),null!==r.iDeferLoading&&(h.bDeferLoading=!0,t=Array.isArray(r.iDeferLoading),h._iRecordsDisplay=t?r.iDeferLoading[0]:r.iDeferLoading,h._iRecordsTotal=t?r.iDeferLoading[1]:r.iDeferLoading),h.oLanguage),t=(P.extend(!0,g,r.oLanguage),g.sUrl?(P.ajax({dataType:"json",url:g.sUrl,success:function(t){C(e.oLanguage,t),Z(t),P.extend(!0,g,t,h.oInit.oLanguage),R(h,null,"i18n",[h]),Jt(h)},error:function(){Jt(h)}}),i=!0):R(h,null,"i18n",[h]),null===r.asStripeClasses&&(h.asStripeClasses=[p.sStripeOdd,p.sStripeEven]),h.asStripeClasses),b=l.children("tbody").find("tr").eq(0),m=(-1!==P.inArray(!0,P.map(t,function(t,e){return b.hasClass(t)}))&&(P("tbody tr",this).removeClass(t.join(" ")),h.asDestroyStripes=t.slice()),[]),t=this.getElementsByTagName("thead");if(0!==t.length&&(wt(h.aoHeader,t[0]),m=Ct(h)),null===r.aoColumns)for(f=[],o=0,s=m.length;o<s;o++)f.push(null);else f=r.aoColumns;for(o=0,s=f.length;o<s;o++)nt(h,m?m[o]:null);st(h,r.aoColumnDefs,f,function(t,e){at(h,t,e)}),b.length&&(d=function(t,e){return null!==t.getAttribute("data-"+e)?e:null},P(b[0]).children("th, td").each(function(t,e){var n,a=h.aoColumns[t];a||W(h,0,"Incorrect column count",18),a.mData===t&&(n=d(e,"sort")||d(e,"order"),e=d(e,"filter")||d(e,"search"),null===n&&null===e||(a.mData={_:t+".display",sort:null!==n?t+".@data-"+n:H,type:null!==n?t+".@data-"+n:H,filter:null!==e?t+".@data-"+e:H},a._isArrayHost=!0,at(h,t)))}));var S=h.oFeatures,t=function(){if(r.aaSorting===H){var t=h.aaSorting;for(o=0,s=t.length;o<s;o++)t[o][1]=h.aoColumns[o].asSorting[0]}ce(h),S.bSort&&L(h,"aoDrawCallback",function(){var t,n;h.bSorted&&(t=I(h),n={},P.each(t,function(t,e){n[e.src]=e.dir}),R(h,null,"order",[h,t,n]),le(h))}),L(h,"aoDrawCallback",function(){(h.bSorted||"ssp"===E(h)||S.bDeferRender)&&ce(h)},"sc");var e=l.children("caption").each(function(){this._captionSide=P(this).css("caption-side")}),n=l.children("thead"),a=(0===n.length&&(n=P("<thead/>").appendTo(l)),h.nTHead=n[0],l.children("tbody")),n=(0===a.length&&(a=P("<tbody/>").insertAfter(n)),h.nTBody=a[0],l.children("tfoot"));if(0===(n=0===n.length&&0<e.length&&(""!==h.oScroll.sX||""!==h.oScroll.sY)?P("<tfoot/>").appendTo(l):n).length||0===n.children().length?l.addClass(p.sNoFooter):0<n.length&&(h.nTFoot=n[0],wt(h.aoFooter,h.nTFoot)),r.aaData)for(o=0;o<r.aaData.length;o++)x(h,r.aaData[o]);else!h.bDeferLoading&&"dom"!=E(h)||ut(h,P(h.nTBody).children("tr"));h.aiDisplay=h.aiDisplayMaster.slice(),!(h.bInitialised=!0)===i&&Jt(h)};L(h,"aoDrawCallback",de,"state_save"),r.bStateSave?(S.bStateSave=!0,he(h,0,t)):t()}}),y=null,this},c={},U=/[\r\n\u2028]/g,V=/<.*?>/g,X=/^\d{2,4}[\.\/\-]\d{1,2}[\.\/\-]\d{1,2}([T ]{1}\d{1,2}[:\.]\d{2}([\.:]\d{2})?)?$/,J=new RegExp("(\\"+["/",".","*","+","?","|","(",")","[","]","{","}","\\","$","^","-"].join("|\\")+")","g"),q=/['\u00A0,$£€¥%\u2009\u202F\u20BD\u20a9\u20BArfkɃΞ]/gi,h=function(t){return!t||!0===t||"-"===t},$=function(t,e){return c[e]||(c[e]=new RegExp(Ot(e),"g")),"string"==typeof t&&"."!==e?t.replace(/\./g,"").replace(c[e],"."):t},N=function(t,e,n){var a=[],r=0,o=t.length;if(n!==H)for(;r<o;r++)t[r]&&t[r][e]&&a.push(t[r][e][n]);else for(;r<o;r++)t[r]&&a.push(t[r][e]);return a},G=function(t){if(!(t.length<2))for(var e=t.slice().sort(),n=e[0],a=1,r=e.length;a<r;a++){if(e[a]===n)return!1;n=e[a]}return!0},z=function(t){if(G(t))return t.slice();var e,n,a,r=[],o=t.length,i=0;t:for(n=0;n<o;n++){for(e=t[n],a=0;a<i;a++)if(r[a]===e)continue t;r.push(e),i++}return r},Y=function(t,e){if(Array.isArray(e))for(var n=0;n<e.length;n++)Y(t,e[n]);else t.push(e);return t};function i(n){var a,r,o={};P.each(n,function(t,e){(a=t.match(/^([^A-Z]+?)([A-Z])/))&&-1!=="a aa ai ao as b fn i m o s ".indexOf(a[1]+" ")&&(r=t.replace(a[0],a[2].toLowerCase()),o[r]=t,"o"===a[1])&&i(n[t])}),n._hungarianMap=o}function C(n,a,r){var o;n._hungarianMap||i(n),P.each(a,function(t,e){(o=n._hungarianMap[t])===H||!r&&a[o]!==H||("o"===o.charAt(0)?(a[o]||(a[o]={}),P.extend(!0,a[o],a[t]),C(n[o],a[o],r)):a[o]=a[t])})}function Z(t){var e,n=w.defaults.oLanguage,a=n.sDecimal;a&&Me(a),t&&(e=t.sZeroRecords,!t.sEmptyTable&&e&&"No data available in table"===n.sEmptyTable&&F(t,t,"sZeroRecords","sEmptyTable"),!t.sLoadingRecords&&e&&"Loading..."===n.sLoadingRecords&&F(t,t,"sZeroRecords","sLoadingRecords"),t.sInfoThousands&&(t.sThousands=t.sInfoThousands),e=t.sDecimal)&&a!==e&&Me(e)}Array.isArray||(Array.isArray=function(t){return"[object Array]"===Object.prototype.toString.call(t)}),Array.prototype.includes||(Array.prototype.includes=s),String.prototype.trim||(String.prototype.trim=function(){return this.replace(/^[\s\uFEFF\xA0]+|[\s\uFEFF\xA0]+$/g,"")}),String.prototype.includes||(String.prototype.includes=s),w.util={throttle:function(a,t){var r,o,i=t!==H?t:200;return function(){var t=this,e=+new Date,n=arguments;r&&e<r+i?(clearTimeout(o),o=setTimeout(function(){r=H,a.apply(t,n)},i)):(r=e,a.apply(t,n))}},escapeRegex:function(t){return t.replace(J,"\\$1")},set:function(a){var d;return P.isPlainObject(a)?w.util.set(a._):null===a?function(){}:"function"==typeof a?function(t,e,n){a(t,"set",e,n)}:"string"!=typeof a||-1===a.indexOf(".")&&-1===a.indexOf("[")&&-1===a.indexOf("(")?function(t,e){t[a]=e}:(d=function(t,e,n){for(var a,r,o,i,l=dt(n),n=l[l.length-1],s=0,u=l.length-1;s<u;s++){if("__proto__"===l[s]||"constructor"===l[s])throw new Error("Cannot set prototype values");if(a=l[s].match(ft),r=l[s].match(g),a){if(l[s]=l[s].replace(ft,""),t[l[s]]=[],(a=l.slice()).splice(0,s+1),i=a.join("."),Array.isArray(e))for(var c=0,f=e.length;c<f;c++)d(o={},e[c],i),t[l[s]].push(o);else t[l[s]]=e;return}r&&(l[s]=l[s].replace(g,""),t=t[l[s]](e)),null!==t[l[s]]&&t[l[s]]!==H||(t[l[s]]={}),t=t[l[s]]}n.match(g)?t[n.replace(g,"")](e):t[n.replace(ft,"")]=e},function(t,e){return d(t,e,a)})},get:function(r){var o,d;return P.isPlainObject(r)?(o={},P.each(r,function(t,e){e&&(o[t]=w.util.get(e))}),function(t,e,n,a){var r=o[e]||o._;return r!==H?r(t,e,n,a):t}):null===r?function(t){return t}:"function"==typeof r?function(t,e,n,a){return r(t,e,n,a)}:"string"!=typeof r||-1===r.indexOf(".")&&-1===r.indexOf("[")&&-1===r.indexOf("(")?function(t,e){return t[r]}:(d=function(t,e,n){var a,r,o;if(""!==n)for(var i=dt(n),l=0,s=i.length;l<s;l++){if(f=i[l].match(ft),a=i[l].match(g),f){if(i[l]=i[l].replace(ft,""),""!==i[l]&&(t=t[i[l]]),r=[],i.splice(0,l+1),o=i.join("."),Array.isArray(t))for(var u=0,c=t.length;u<c;u++)r.push(d(t[u],e,o));var f=f[0].substring(1,f[0].length-1);t=""===f?r:r.join(f);break}if(a)i[l]=i[l].replace(g,""),t=t[i[l]]();else{if(null===t||null===t[i[l]])return null;if(t===H||t[i[l]]===H)return H;t=t[i[l]]}}return t},function(t,e){return d(t,e,r)})}};var r=function(t,e,n){t[e]!==H&&(t[n]=t[e])};function K(t){r(t,"ordering","bSort"),r(t,"orderMulti","bSortMulti"),r(t,"orderClasses","bSortClasses"),r(t,"orderCellsTop","bSortCellsTop"),r(t,"order","aaSorting"),r(t,"orderFixed","aaSortingFixed"),r(t,"paging","bPaginate"),r(t,"pagingType","sPaginationType"),r(t,"pageLength","iDisplayLength"),r(t,"searching","bFilter"),"boolean"==typeof t.sScrollX&&(t.sScrollX=t.sScrollX?"100%":""),"boolean"==typeof t.scrollX&&(t.scrollX=t.scrollX?"100%":"");var e=t.aoSearchCols;if(e)for(var n=0,a=e.length;n<a;n++)e[n]&&C(w.models.oSearch,e[n])}function Q(t){r(t,"orderable","bSortable"),r(t,"orderData","aDataSort"),r(t,"orderSequence","asSorting"),r(t,"orderDataType","sortDataType");var e=t.aDataSort;"number"!=typeof e||Array.isArray(e)||(t.aDataSort=[e])}function tt(t){var e,n,a,r;w.__browser||(w.__browser=e={},r=(a=(n=P("<div/>").css({position:"fixed",top:0,left:-1*P(j).scrollLeft(),height:1,width:1,overflow:"hidden"}).append(P("<div/>").css({position:"absolute",top:1,left:1,width:100,overflow:"scroll"}).append(P("<div/>").css({width:"100%",height:10}))).appendTo("body")).children()).children(),e.barWidth=a[0].offsetWidth-a[0].clientWidth,e.bScrollOversize=100===r[0].offsetWidth&&100!==a[0].clientWidth,e.bScrollbarLeft=1!==Math.round(r.offset().left),e.bBounding=!!n[0].getBoundingClientRect().width,n.remove()),P.extend(t.oBrowser,w.__browser),t.oScroll.iBarWidth=w.__browser.barWidth}function et(t,e,n,a,r,o){var i,l=a,s=!1;for(n!==H&&(i=n,s=!0);l!==r;)t.hasOwnProperty(l)&&(i=s?e(i,t[l],l,t):t[l],s=!0,l+=o);return i}function nt(t,e){var n=w.defaults.column,a=t.aoColumns.length,n=P.extend({},w.models.oColumn,n,{nTh:e||v.createElement("th"),sTitle:n.sTitle||(e?e.innerHTML:""),aDataSort:n.aDataSort||[a],mData:n.mData||a,idx:a}),n=(t.aoColumns.push(n),t.aoPreSearchCols);n[a]=P.extend({},w.models.oSearch,n[a]),at(t,a,P(e).data())}function at(t,e,n){function a(t){return"string"==typeof t&&-1!==t.indexOf("@")}var e=t.aoColumns[e],r=t.oClasses,o=P(e.nTh),i=(!e.sWidthOrig&&(e.sWidthOrig=o.attr("width")||null,u=(o.attr("style")||"").match(/width:\s*(\d+[pxem%]+)/))&&(e.sWidthOrig=u[1]),n!==H&&null!==n&&(Q(n),C(w.defaults.column,n,!0),n.mDataProp===H||n.mData||(n.mData=n.mDataProp),n.sType&&(e._sManualType=n.sType),n.className&&!n.sClass&&(n.sClass=n.className),n.sClass&&o.addClass(n.sClass),u=e.sClass,P.extend(e,n),F(e,n,"sWidth","sWidthOrig"),u!==e.sClass&&(e.sClass=u+" "+e.sClass),n.iDataSort!==H&&(e.aDataSort=[n.iDataSort]),F(e,n,"aDataSort"),e.ariaTitle||(e.ariaTitle=o.attr("aria-label"))),e.mData),l=A(i),s=e.mRender?A(e.mRender):null,u=(e._bAttrSrc=P.isPlainObject(i)&&(a(i.sort)||a(i.type)||a(i.filter)),e._setter=null,e.fnGetData=function(t,e,n){var a=l(t,e,H,n);return s&&e?s(a,e,t,n):a},e.fnSetData=function(t,e,n){return b(i)(t,e,n)},"number"==typeof i||e._isArrayHost||(t._rowReadObject=!0),t.oFeatures.bSort||(e.bSortable=!1,o.addClass(r.sSortableNone)),-1!==P.inArray("asc",e.asSorting)),n=-1!==P.inArray("desc",e.asSorting);e.bSortable&&(u||n)?u&&!n?(e.sSortingClass=r.sSortableAsc,e.sSortingClassJUI=r.sSortJUIAscAllowed):!u&&n?(e.sSortingClass=r.sSortableDesc,e.sSortingClassJUI=r.sSortJUIDescAllowed):(e.sSortingClass=r.sSortable,e.sSortingClassJUI=r.sSortJUI):(e.sSortingClass=r.sSortableNone,e.sSortingClassJUI="")}function O(t){if(!1!==t.oFeatures.bAutoWidth){var e=t.aoColumns;ee(t);for(var n=0,a=e.length;n<a;n++)e[n].nTh.style.width=e[n].sWidth}var r=t.oScroll;""===r.sY&&""===r.sX||Qt(t),R(t,null,"column-sizing",[t])}function rt(t,e){t=it(t,"bVisible");return"number"==typeof t[e]?t[e]:null}function ot(t,e){t=it(t,"bVisible"),e=P.inArray(e,t);return-1!==e?e:null}function T(t){var n=0;return P.each(t.aoColumns,function(t,e){e.bVisible&&"none"!==P(e.nTh).css("display")&&n++}),n}function it(t,n){var a=[];return P.map(t.aoColumns,function(t,e){t[n]&&a.push(e)}),a}function lt(t){for(var e,n,a,r,o,i,l,s=t.aoColumns,u=t.aoData,c=w.ext.type.detect,f=0,d=s.length;f<d;f++)if(l=[],!(o=s[f]).sType&&o._sManualType)o.sType=o._sManualType;else if(!o.sType){for(e=0,n=c.length;e<n;e++){for(a=0,r=u.length;a<r&&(l[a]===H&&(l[a]=S(t,a,f,"type")),(i=c[e](l[a],t))||e===c.length-1)&&("html"!==i||h(l[a]));a++);if(i){o.sType=i;break}}o.sType||(o.sType="string")}}function st(t,e,n,a){var r,o,i,l,s=t.aoColumns;if(e)for(r=e.length-1;0<=r;r--)for(var u,c=(u=e[r]).target!==H?u.target:u.targets!==H?u.targets:u.aTargets,f=0,d=(c=Array.isArray(c)?c:[c]).length;f<d;f++)if("number"==typeof c[f]&&0<=c[f]){for(;s.length<=c[f];)nt(t);a(c[f],u)}else if("number"==typeof c[f]&&c[f]<0)a(s.length+c[f],u);else if("string"==typeof c[f])for(i=0,l=s.length;i<l;i++)"_all"!=c[f]&&!P(s[i].nTh).hasClass(c[f])||a(i,u);if(n)for(r=0,o=n.length;r<o;r++)a(r,n[r])}function x(t,e,n,a){for(var r=t.aoData.length,o=P.extend(!0,{},w.models.oRow,{src:n?"dom":"data",idx:r}),i=(o._aData=e,t.aoData.push(o),t.aoColumns),l=0,s=i.length;l<s;l++)i[l].sType=null;t.aiDisplayMaster.push(r);e=t.rowIdFn(e);return e!==H&&(t.aIds[e]=o),!n&&t.oFeatures.bDeferRender||St(t,r,n,a),r}function ut(n,t){var a;return(t=t instanceof P?t:P(t)).map(function(t,e){return a=mt(n,e),x(n,a.data,e,a.cells)})}function S(t,e,n,a){"search"===a?a="filter":"order"===a&&(a="sort");var r=t.iDraw,o=t.aoColumns[n],i=t.aoData[e]._aData,l=o.sDefaultContent,s=o.fnGetData(i,a,{settings:t,row:e,col:n});if(s===H)return t.iDrawError!=r&&null===l&&(W(t,0,"Requested unknown parameter "+("function"==typeof o.mData?"{function}":"'"+o.mData+"'")+" for row "+e+", column "+n,4),t.iDrawError=r),l;if(s!==i&&null!==s||null===l||a===H){if("function"==typeof s)return s.call(i)}else s=l;return null===s&&"display"===a?"":"filter"===a&&(e=w.ext.type.search)[o.sType]?e[o.sType](s):s}function ct(t,e,n,a){var r=t.aoColumns[n],o=t.aoData[e]._aData;r.fnSetData(o,a,{settings:t,row:e,col:n})}var ft=/\[.*?\]$/,g=/\(\)$/;function dt(t){return P.map(t.match(/(\\.|[^\.])+/g)||[""],function(t){return t.replace(/\\\./g,".")})}var A=w.util.get,b=w.util.set;function ht(t){return N(t.aoData,"_aData")}function pt(t){t.aoData.length=0,t.aiDisplayMaster.length=0,t.aiDisplay.length=0,t.aIds={}}function gt(t,e,n){for(var a=-1,r=0,o=t.length;r<o;r++)t[r]==e?a=r:t[r]>e&&t[r]--;-1!=a&&n===H&&t.splice(a,1)}function bt(n,a,t,e){function r(t,e){for(;t.childNodes.length;)t.removeChild(t.firstChild);t.innerHTML=S(n,a,e,"display")}var o,i,l=n.aoData[a];if("dom"!==t&&(t&&"auto"!==t||"dom"!==l.src)){var s=l.anCells;if(s)if(e!==H)r(s[e],e);else for(o=0,i=s.length;o<i;o++)r(s[o],o)}else l._aData=mt(n,l,e,e===H?H:l._aData).data;l._aSortData=null,l._aFilterData=null;var u=n.aoColumns;if(e!==H)u[e].sType=null;else{for(o=0,i=u.length;o<i;o++)u[o].sType=null;vt(n,l)}}function mt(t,e,n,a){function r(t,e){var n;"string"==typeof t&&-1!==(n=t.indexOf("@"))&&(n=t.substring(n+1),b(t)(a,e.getAttribute(n)))}function o(t){n!==H&&n!==f||(l=d[f],s=t.innerHTML.trim(),l&&l._bAttrSrc?(b(l.mData._)(a,s),r(l.mData.sort,t),r(l.mData.type,t),r(l.mData.filter,t)):h?(l._setter||(l._setter=b(l.mData)),l._setter(a,s)):a[f]=s),f++}var i,l,s,u=[],c=e.firstChild,f=0,d=t.aoColumns,h=t._rowReadObject;a=a!==H?a:h?{}:[];if(c)for(;c;)"TD"!=(i=c.nodeName.toUpperCase())&&"TH"!=i||(o(c),u.push(c)),c=c.nextSibling;else for(var p=0,g=(u=e.anCells).length;p<g;p++)o(u[p]);var e=e.firstChild?e:e.nTr;return e&&(e=e.getAttribute("id"))&&b(t.rowId)(a,e),{data:a,cells:u}}function St(t,e,n,a){var r,o,i,l,s,u,c=t.aoData[e],f=c._aData,d=[];if(null===c.nTr){for(r=n||v.createElement("tr"),c.nTr=r,c.anCells=d,r._DT_RowIndex=e,vt(t,c),l=0,s=t.aoColumns.length;l<s;l++)i=t.aoColumns[l],(o=(u=!n)?v.createElement(i.sCellType):a[l])||W(t,0,"Incorrect column count",18),o._DT_CellIndex={row:e,column:l},d.push(o),!u&&(!i.mRender&&i.mData===l||P.isPlainObject(i.mData)&&i.mData._===l+".display")||(o.innerHTML=S(t,e,l,"display")),i.sClass&&(o.className+=" "+i.sClass),i.bVisible&&!n?r.appendChild(o):!i.bVisible&&n&&o.parentNode.removeChild(o),i.fnCreatedCell&&i.fnCreatedCell.call(t.oInstance,o,S(t,e,l),f,e,l);R(t,"aoRowCreatedCallback",null,[r,f,e,d])}}function vt(t,e){var n=e.nTr,a=e._aData;n&&((t=t.rowIdFn(a))&&(n.id=t),a.DT_RowClass&&(t=a.DT_RowClass.split(" "),e.__rowc=e.__rowc?z(e.__rowc.concat(t)):t,P(n).removeClass(e.__rowc.join(" ")).addClass(a.DT_RowClass)),a.DT_RowAttr&&P(n).attr(a.DT_RowAttr),a.DT_RowData)&&P(n).data(a.DT_RowData)}function yt(t){var e,n,a,r=t.nTHead,o=t.nTFoot,i=0===P("th, td",r).length,l=t.oClasses,s=t.aoColumns;for(i&&(n=P("<tr/>").appendTo(r)),c=0,f=s.length;c<f;c++)a=s[c],e=P(a.nTh).addClass(a.sClass),i&&e.appendTo(n),t.oFeatures.bSort&&(e.addClass(a.sSortingClass),!1!==a.bSortable)&&(e.attr("tabindex",t.iTabIndex).attr("aria-controls",t.sTableId),ue(t,a.nTh,c)),a.sTitle!=e[0].innerHTML&&e.html(a.sTitle),ve(t,"header")(t,e,a,l);if(i&&wt(t.aoHeader,r),P(r).children("tr").children("th, td").addClass(l.sHeaderTH),P(o).children("tr").children("th, td").addClass(l.sFooterTH),null!==o)for(var u=t.aoFooter[0],c=0,f=u.length;c<f;c++)(a=s[c])?(a.nTf=u[c].cell,a.sClass&&P(a.nTf).addClass(a.sClass)):W(t,0,"Incorrect column count",18)}function Dt(t,e,n){var a,r,o,i,l,s,u,c,f,d=[],h=[],p=t.aoColumns.length;if(e){for(n===H&&(n=!1),a=0,r=e.length;a<r;a++){for(d[a]=e[a].slice(),d[a].nTr=e[a].nTr,o=p-1;0<=o;o--)t.aoColumns[o].bVisible||n||d[a].splice(o,1);h.push([])}for(a=0,r=d.length;a<r;a++){if(u=d[a].nTr)for(;s=u.firstChild;)u.removeChild(s);for(o=0,i=d[a].length;o<i;o++)if(f=c=1,h[a][o]===H){for(u.appendChild(d[a][o].cell),h[a][o]=1;d[a+c]!==H&&d[a][o].cell==d[a+c][o].cell;)h[a+c][o]=1,c++;for(;d[a][o+f]!==H&&d[a][o].cell==d[a][o+f].cell;){for(l=0;l<c;l++)h[a+l][o+f]=1;f++}P(d[a][o].cell).attr("rowspan",c).attr("colspan",f)}}}}function y(t,e){n="ssp"==E(s=t),(l=s.iInitDisplayStart)!==H&&-1!==l&&(s._iDisplayStart=!n&&l>=s.fnRecordsDisplay()?0:l,s.iInitDisplayStart=-1);var n=R(t,"aoPreDrawCallback","preDraw",[t]);if(-1!==P.inArray(!1,n))D(t,!1);else{var a=[],r=0,o=t.asStripeClasses,i=o.length,l=t.oLanguage,s="ssp"==E(t),u=t.aiDisplay,n=t._iDisplayStart,c=t.fnDisplayEnd();if(t.bDrawing=!0,t.bDeferLoading)t.bDeferLoading=!1,t.iDraw++,D(t,!1);else if(s){if(!t.bDestroying&&!e)return void xt(t)}else t.iDraw++;if(0!==u.length)for(var f=s?t.aoData.length:c,d=s?0:n;d<f;d++){var h,p=u[d],g=t.aoData[p],b=(null===g.nTr&&St(t,p),g.nTr);0!==i&&(h=o[r%i],g._sRowStripe!=h)&&(P(b).removeClass(g._sRowStripe).addClass(h),g._sRowStripe=h),R(t,"aoRowCallback",null,[b,g._aData,r,d,p]),a.push(b),r++}else{e=l.sZeroRecords;1==t.iDraw&&"ajax"==E(t)?e=l.sLoadingRecords:l.sEmptyTable&&0===t.fnRecordsTotal()&&(e=l.sEmptyTable),a[0]=P("<tr/>",{class:i?o[0]:""}).append(P("<td />",{valign:"top",colSpan:T(t),class:t.oClasses.sRowEmpty}).html(e))[0]}R(t,"aoHeaderCallback","header",[P(t.nTHead).children("tr")[0],ht(t),n,c,u]),R(t,"aoFooterCallback","footer",[P(t.nTFoot).children("tr")[0],ht(t),n,c,u]);s=P(t.nTBody);s.children().detach(),s.append(P(a)),R(t,"aoDrawCallback","draw",[t]),t.bSorted=!1,t.bFiltered=!1,t.bDrawing=!1}}function u(t,e){var n=t.oFeatures,a=n.bSort,n=n.bFilter;a&&ie(t),n?Rt(t,t.oPreviousSearch):t.aiDisplay=t.aiDisplayMaster.slice(),!0!==e&&(t._iDisplayStart=0),t._drawHold=e,y(t),t._drawHold=!1}function _t(t){for(var e,n,a,r,o,i,l,s=t.oClasses,u=P(t.nTable),u=P("<div/>").insertBefore(u),c=t.oFeatures,f=P("<div/>",{id:t.sTableId+"_wrapper",class:s.sWrapper+(t.nTFoot?"":" "+s.sNoFooter)}),d=(t.nHolding=u[0],t.nTableWrapper=f[0],t.nTableReinsertBefore=t.nTable.nextSibling,t.sDom.split("")),h=0;h<d.length;h++){if(e=null,"<"==(n=d[h])){if(a=P("<div/>")[0],"'"==(r=d[h+1])||'"'==r){for(o="",i=2;d[h+i]!=r;)o+=d[h+i],i++;"H"==o?o=s.sJUIHeader:"F"==o&&(o=s.sJUIFooter),-1!=o.indexOf(".")?(l=o.split("."),a.id=l[0].substr(1,l[0].length-1),a.className=l[1]):"#"==o.charAt(0)?a.id=o.substr(1,o.length-1):a.className=o,h+=i}f.append(a),f=P(a)}else if(">"==n)f=f.parent();else if("l"==n&&c.bPaginate&&c.bLengthChange)e=Gt(t);else if("f"==n&&c.bFilter)e=Lt(t);else if("r"==n&&c.bProcessing)e=Zt(t);else if("t"==n)e=Kt(t);else if("i"==n&&c.bInfo)e=Ut(t);else if("p"==n&&c.bPaginate)e=zt(t);else if(0!==w.ext.feature.length)for(var p=w.ext.feature,g=0,b=p.length;g<b;g++)if(n==p[g].cFeature){e=p[g].fnInit(t);break}e&&((l=t.aanFeatures)[n]||(l[n]=[]),l[n].push(e),f.append(e))}u.replaceWith(f),t.nHolding=null}function wt(t,e){var n,a,r,o,i,l,s,u,c,f,d=P(e).children("tr");for(t.splice(0,t.length),r=0,l=d.length;r<l;r++)t.push([]);for(r=0,l=d.length;r<l;r++)for(a=(n=d[r]).firstChild;a;){if("TD"==a.nodeName.toUpperCase()||"TH"==a.nodeName.toUpperCase())for(u=(u=+a.getAttribute("colspan"))&&0!=u&&1!=u?u:1,c=(c=+a.getAttribute("rowspan"))&&0!=c&&1!=c?c:1,s=function(t,e,n){for(var a=t[e];a[n];)n++;return n}(t,r,0),f=1==u,i=0;i<u;i++)for(o=0;o<c;o++)t[r+o][s+i]={cell:a,unique:f},t[r+o].nTr=n;a=a.nextSibling}}function Ct(t,e,n){var a=[];n||(n=t.aoHeader,e&&wt(n=[],e));for(var r=0,o=n.length;r<o;r++)for(var i=0,l=n[r].length;i<l;i++)!n[r][i].unique||a[i]&&t.bSortCellsTop||(a[i]=n[r][i].cell);return a}function Tt(r,t,n){function e(t){var e=r.jqXHR?r.jqXHR.status:null;(null===t||"number"==typeof e&&204==e)&&Ft(r,t={},[]),(e=t.error||t.sError)&&W(r,0,e),r.json=t,R(r,null,"xhr",[r,t,r.jqXHR]),n(t)}R(r,"aoServerParams","serverParams",[t]),t&&Array.isArray(t)&&(a={},o=/(.*?)\[\]$/,P.each(t,function(t,e){var n=e.name.match(o);n?(n=n[0],a[n]||(a[n]=[]),a[n].push(e.value)):a[e.name]=e.value}),t=a);var a,o,i,l=r.ajax,s=r.oInstance,u=(P.isPlainObject(l)&&l.data&&(u="function"==typeof(i=l.data)?i(t,r):i,t="function"==typeof i&&u?u:P.extend(!0,t,u),delete l.data),{data:t,success:e,dataType:"json",cache:!1,type:r.sServerMethod,error:function(t,e,n){var a=R(r,null,"xhr",[r,null,r.jqXHR]);-1===P.inArray(!0,a)&&("parsererror"==e?W(r,0,"Invalid JSON response",1):4===t.readyState&&W(r,0,"Ajax error",7)),D(r,!1)}});r.oAjaxData=t,R(r,null,"preXhr",[r,t]),r.fnServerData?r.fnServerData.call(s,r.sAjaxSource,P.map(t,function(t,e){return{name:e,value:t}}),e,r):r.sAjaxSource||"string"==typeof l?r.jqXHR=P.ajax(P.extend(u,{url:l||r.sAjaxSource})):"function"==typeof l?r.jqXHR=l.call(s,t,e,r):(r.jqXHR=P.ajax(P.extend(u,l)),l.data=i)}function xt(e){e.iDraw++,D(e,!0);var n=e._drawHold;Tt(e,At(e),function(t){e._drawHold=n,It(e,t),e._drawHold=!1})}function At(t){for(var e,n,a,r=t.aoColumns,o=r.length,i=t.oFeatures,l=t.oPreviousSearch,s=t.aoPreSearchCols,u=[],c=I(t),f=t._iDisplayStart,d=!1!==i.bPaginate?t._iDisplayLength:-1,h=function(t,e){u.push({name:t,value:e})},p=(h("sEcho",t.iDraw),h("iColumns",o),h("sColumns",N(r,"sName").join(",")),h("iDisplayStart",f),h("iDisplayLength",d),{draw:t.iDraw,columns:[],order:[],start:f,length:d,search:{value:l.sSearch,regex:l.bRegex}}),g=0;g<o;g++)n=r[g],a=s[g],e="function"==typeof n.mData?"function":n.mData,p.columns.push({data:e,name:n.sName,searchable:n.bSearchable,orderable:n.bSortable,search:{value:a.sSearch,regex:a.bRegex}}),h("mDataProp_"+g,e),i.bFilter&&(h("sSearch_"+g,a.sSearch),h("bRegex_"+g,a.bRegex),h("bSearchable_"+g,n.bSearchable)),i.bSort&&h("bSortable_"+g,n.bSortable);i.bFilter&&(h("sSearch",l.sSearch),h("bRegex",l.bRegex)),i.bSort&&(P.each(c,function(t,e){p.order.push({column:e.col,dir:e.dir}),h("iSortCol_"+t,e.col),h("sSortDir_"+t,e.dir)}),h("iSortingCols",c.length));f=w.ext.legacy.ajax;return null===f?t.sAjaxSource?u:p:f?u:p}function It(t,n){function e(t,e){return n[t]!==H?n[t]:n[e]}var a=Ft(t,n),r=e("sEcho","draw"),o=e("iTotalRecords","recordsTotal"),i=e("iTotalDisplayRecords","recordsFiltered");if(r!==H){if(+r<t.iDraw)return;t.iDraw=+r}a=a||[],pt(t),t._iRecordsTotal=parseInt(o,10),t._iRecordsDisplay=parseInt(i,10);for(var l=0,s=a.length;l<s;l++)x(t,a[l]);t.aiDisplay=t.aiDisplayMaster.slice(),y(t,!0),t._bInitComplete||qt(t,n),D(t,!1)}function Ft(t,e,n){t=P.isPlainObject(t.ajax)&&t.ajax.dataSrc!==H?t.ajax.dataSrc:t.sAjaxDataProp;if(!n)return"data"===t?e.aaData||e[t]:""!==t?A(t)(e):e;b(t)(e,n)}function Lt(n){function e(t){i.f;var e=this.value||"";o.return&&"Enter"!==t.key||e!=o.sSearch&&(Rt(n,{sSearch:e,bRegex:o.bRegex,bSmart:o.bSmart,bCaseInsensitive:o.bCaseInsensitive,return:o.return}),n._iDisplayStart=0,y(n))}var t=n.oClasses,a=n.sTableId,r=n.oLanguage,o=n.oPreviousSearch,i=n.aanFeatures,l='<input type="search" class="'+t.sFilterInput+'"/>',s=(s=r.sSearch).match(/_INPUT_/)?s.replace("_INPUT_",l):s+l,l=P("<div/>",{id:i.f?null:a+"_filter",class:t.sFilter}).append(P("<label/>").append(s)),t=null!==n.searchDelay?n.searchDelay:"ssp"===E(n)?400:0,u=P("input",l).val(o.sSearch).attr("placeholder",r.sSearchPlaceholder).on("keyup.DT search.DT input.DT paste.DT cut.DT",t?ne(e,t):e).on("mouseup.DT",function(t){setTimeout(function(){e.call(u[0],t)},10)}).on("keypress.DT",function(t){if(13==t.keyCode)return!1}).attr("aria-controls",a);return P(n.nTable).on("search.dt.DT",function(t,e){if(n===e)try{u[0]!==v.activeElement&&u.val(o.sSearch)}catch(t){}}),l[0]}function Rt(t,e,n){function a(t){o.sSearch=t.sSearch,o.bRegex=t.bRegex,o.bSmart=t.bSmart,o.bCaseInsensitive=t.bCaseInsensitive,o.return=t.return}function r(t){return t.bEscapeRegex!==H?!t.bEscapeRegex:t.bRegex}var o=t.oPreviousSearch,i=t.aoPreSearchCols;if(lt(t),"ssp"!=E(t)){Ht(t,e.sSearch,n,r(e),e.bSmart,e.bCaseInsensitive),a(e);for(var l=0;l<i.length;l++)jt(t,i[l].sSearch,l,r(i[l]),i[l].bSmart,i[l].bCaseInsensitive);Pt(t)}else a(e);t.bFiltered=!0,R(t,null,"search",[t])}function Pt(t){for(var e,n,a=w.ext.search,r=t.aiDisplay,o=0,i=a.length;o<i;o++){for(var l=[],s=0,u=r.length;s<u;s++)n=r[s],e=t.aoData[n],a[o](t,e._aFilterData,n,e._aData,s)&&l.push(n);r.length=0,P.merge(r,l)}}function jt(t,e,n,a,r,o){if(""!==e){for(var i,l=[],s=t.aiDisplay,u=Nt(e,a,r,o),c=0;c<s.length;c++)i=t.aoData[s[c]]._aFilterData[n],u.test(i)&&l.push(s[c]);t.aiDisplay=l}}function Ht(t,e,n,a,r,o){var i,l,s,u=Nt(e,a,r,o),r=t.oPreviousSearch.sSearch,o=t.aiDisplayMaster,c=[];if(0!==w.ext.search.length&&(n=!0),l=Wt(t),e.length<=0)t.aiDisplay=o.slice();else{for((l||n||a||r.length>e.length||0!==e.indexOf(r)||t.bSorted)&&(t.aiDisplay=o.slice()),i=t.aiDisplay,s=0;s<i.length;s++)u.test(t.aoData[i[s]]._sFilterRow)&&c.push(i[s]);t.aiDisplay=c}}function Nt(t,e,n,a){return t=e?t:Ot(t),n&&(t="^(?=.*?"+P.map(t.match(/["\u201C][^"\u201D]+["\u201D]|[^ ]+/g)||[""],function(t){var e;return'"'===t.charAt(0)?t=(e=t.match(/^"(.*)"$/))?e[1]:t:"“"===t.charAt(0)&&(t=(e=t.match(/^\u201C(.*)\u201D$/))?e[1]:t),t.replace('"',"")}).join(")(?=.*?")+").*$"),new RegExp(t,a?"i":"")}var Ot=w.util.escapeRegex,kt=P("<div>")[0],Mt=kt.textContent!==H;function Wt(t){for(var e,n,a,r,o,i=t.aoColumns,l=!1,s=0,u=t.aoData.length;s<u;s++)if(!(o=t.aoData[s])._aFilterData){for(a=[],e=0,n=i.length;e<n;e++)i[e].bSearchable?"string"!=typeof(r=null===(r=S(t,s,e,"filter"))?"":r)&&r.toString&&(r=r.toString()):r="",r.indexOf&&-1!==r.indexOf("&")&&(kt.innerHTML=r,r=Mt?kt.textContent:kt.innerText),r.replace&&(r=r.replace(/[\r\n\u2028]/g,"")),a.push(r);o._aFilterData=a,o._sFilterRow=a.join("  "),l=!0}return l}function Et(t){return{search:t.sSearch,smart:t.bSmart,regex:t.bRegex,caseInsensitive:t.bCaseInsensitive}}function Bt(t){return{sSearch:t.search,bSmart:t.smart,bRegex:t.regex,bCaseInsensitive:t.caseInsensitive}}function Ut(t){var e=t.sTableId,n=t.aanFeatures.i,a=P("<div/>",{class:t.oClasses.sInfo,id:n?null:e+"_info"});return n||(t.aoDrawCallback.push({fn:Vt,sName:"information"}),a.attr("role","status").attr("aria-live","polite"),P(t.nTable).attr("aria-describedby",e+"_info")),a[0]}function Vt(t){var e,n,a,r,o,i,l=t.aanFeatures.i;0!==l.length&&(i=t.oLanguage,e=t._iDisplayStart+1,n=t.fnDisplayEnd(),a=t.fnRecordsTotal(),o=(r=t.fnRecordsDisplay())?i.sInfo:i.sInfoEmpty,r!==a&&(o+=" "+i.sInfoFiltered),o=Xt(t,o+=i.sInfoPostFix),null!==(i=i.fnInfoCallback)&&(o=i.call(t.oInstance,t,e,n,a,r,o)),P(l).html(o))}function Xt(t,e){var n=t.fnFormatNumber,a=t._iDisplayStart+1,r=t._iDisplayLength,o=t.fnRecordsDisplay(),i=-1===r;return e.replace(/_START_/g,n.call(t,a)).replace(/_END_/g,n.call(t,t.fnDisplayEnd())).replace(/_MAX_/g,n.call(t,t.fnRecordsTotal())).replace(/_TOTAL_/g,n.call(t,o)).replace(/_PAGE_/g,n.call(t,i?1:Math.ceil(a/r))).replace(/_PAGES_/g,n.call(t,i?1:Math.ceil(o/r)))}function Jt(n){var a,t,e,r=n.iInitDisplayStart,o=n.aoColumns,i=n.oFeatures,l=n.bDeferLoading;if(n.bInitialised){for(_t(n),yt(n),Dt(n,n.aoHeader),Dt(n,n.aoFooter),D(n,!0),i.bAutoWidth&&ee(n),a=0,t=o.length;a<t;a++)(e=o[a]).sWidth&&(e.nTh.style.width=M(e.sWidth));R(n,null,"preInit",[n]),u(n);i=E(n);"ssp"==i&&!l||("ajax"==i?Tt(n,[],function(t){var e=Ft(n,t);for(a=0;a<e.length;a++)x(n,e[a]);n.iInitDisplayStart=r,u(n),D(n,!1),qt(n,t)}):(D(n,!1),qt(n)))}else setTimeout(function(){Jt(n)},200)}function qt(t,e){t._bInitComplete=!0,(e||t.oInit.aaData)&&O(t),R(t,null,"plugin-init",[t,e]),R(t,"aoInitComplete","init",[t,e])}function $t(t,e){e=parseInt(e,10);t._iDisplayLength=e,Se(t),R(t,null,"length",[t,e])}function Gt(a){for(var t=a.oClasses,e=a.sTableId,n=a.aLengthMenu,r=Array.isArray(n[0]),o=r?n[0]:n,i=r?n[1]:n,l=P("<select/>",{name:e+"_length","aria-controls":e,class:t.sLengthSelect}),s=0,u=o.length;s<u;s++)l[0][s]=new Option("number"==typeof i[s]?a.fnFormatNumber(i[s]):i[s],o[s]);var c=P("<div><label/></div>").addClass(t.sLength);return a.aanFeatures.l||(c[0].id=e+"_length"),c.children().append(a.oLanguage.sLengthMenu.replace("_MENU_",l[0].outerHTML)),P("select",c).val(a._iDisplayLength).on("change.DT",function(t){$t(a,P(this).val()),y(a)}),P(a.nTable).on("length.dt.DT",function(t,e,n){a===e&&P("select",c).val(n)}),c[0]}function zt(t){function c(t){y(t)}var e=t.sPaginationType,f=w.ext.pager[e],d="function"==typeof f,e=P("<div/>").addClass(t.oClasses.sPaging+e)[0],h=t.aanFeatures;return d||f.fnInit(t,e,c),h.p||(e.id=t.sTableId+"_paginate",t.aoDrawCallback.push({fn:function(t){if(d)for(var e=t._iDisplayStart,n=t._iDisplayLength,a=t.fnRecordsDisplay(),r=-1===n,o=r?0:Math.ceil(e/n),i=r?1:Math.ceil(a/n),l=f(o,i),s=0,u=h.p.length;s<u;s++)ve(t,"pageButton")(t,h.p[s],s,l,o,i);else f.fnUpdate(t,c)},sName:"pagination"})),e}function Yt(t,e,n){var a=t._iDisplayStart,r=t._iDisplayLength,o=t.fnRecordsDisplay(),o=(0===o||-1===r?a=0:"number"==typeof e?o<(a=e*r)&&(a=0):"first"==e?a=0:"previous"==e?(a=0<=r?a-r:0)<0&&(a=0):"next"==e?a+r<o&&(a+=r):"last"==e?a=Math.floor((o-1)/r)*r:W(t,0,"Unknown paging action: "+e,5),t._iDisplayStart!==a);return t._iDisplayStart=a,o?(R(t,null,"page",[t]),n&&y(t)):R(t,null,"page-nc",[t]),o}function Zt(t){return P("<div/>",{id:t.aanFeatures.r?null:t.sTableId+"_processing",class:t.oClasses.sProcessing,role:"status"}).html(t.oLanguage.sProcessing).append("<div><div></div><div></div><div></div><div></div></div>").insertBefore(t.nTable)[0]}function D(t,e){t.oFeatures.bProcessing&&P(t.aanFeatures.r).css("display",e?"block":"none"),R(t,null,"processing",[t,e])}function Kt(t){var e,n,a,r,o,i,l,s,u,c,f,d,h=P(t.nTable),p=t.oScroll;return""===p.sX&&""===p.sY?t.nTable:(e=p.sX,n=p.sY,a=t.oClasses,o=(r=h.children("caption")).length?r[0]._captionSide:null,s=P(h[0].cloneNode(!1)),i=P(h[0].cloneNode(!1)),u=function(t){return t?M(t):null},(l=h.children("tfoot")).length||(l=null),s=P(f="<div/>",{class:a.sScrollWrapper}).append(P(f,{class:a.sScrollHead}).css({overflow:"hidden",position:"relative",border:0,width:e?u(e):"100%"}).append(P(f,{class:a.sScrollHeadInner}).css({"box-sizing":"content-box",width:p.sXInner||"100%"}).append(s.removeAttr("id").css("margin-left",0).append("top"===o?r:null).append(h.children("thead"))))).append(P(f,{class:a.sScrollBody}).css({position:"relative",overflow:"auto",width:u(e)}).append(h)),l&&s.append(P(f,{class:a.sScrollFoot}).css({overflow:"hidden",border:0,width:e?u(e):"100%"}).append(P(f,{class:a.sScrollFootInner}).append(i.removeAttr("id").css("margin-left",0).append("bottom"===o?r:null).append(h.children("tfoot"))))),u=s.children(),c=u[0],f=u[1],d=l?u[2]:null,e&&P(f).on("scroll.DT",function(t){var e=this.scrollLeft;c.scrollLeft=e,l&&(d.scrollLeft=e)}),P(f).css("max-height",n),p.bCollapse||P(f).css("height",n),t.nScrollHead=c,t.nScrollBody=f,t.nScrollFoot=d,t.aoDrawCallback.push({fn:Qt,sName:"scrolling"}),s[0])}function Qt(n){function t(t){(t=t.style).paddingTop="0",t.paddingBottom="0",t.borderTopWidth="0",t.borderBottomWidth="0",t.height=0}var e,a,r,o,i,l=n.oScroll,s=l.sX,u=l.sXInner,c=l.sY,l=l.iBarWidth,f=P(n.nScrollHead),d=f[0].style,h=f.children("div"),p=h[0].style,h=h.children("table"),g=n.nScrollBody,b=P(g),m=g.style,S=P(n.nScrollFoot).children("div"),v=S.children("table"),y=P(n.nTHead),D=P(n.nTable),_=D[0],w=_.style,C=n.nTFoot?P(n.nTFoot):null,T=n.oBrowser,x=T.bScrollOversize,A=(N(n.aoColumns,"nTh"),[]),I=[],F=[],L=[],R=g.scrollHeight>g.clientHeight;n.scrollBarVis!==R&&n.scrollBarVis!==H?(n.scrollBarVis=R,O(n)):(n.scrollBarVis=R,D.children("thead, tfoot").remove(),C&&(R=C.clone().prependTo(D),i=C.find("tr"),a=R.find("tr"),R.find("[id]").removeAttr("id")),R=y.clone().prependTo(D),y=y.find("tr"),e=R.find("tr"),R.find("th, td").removeAttr("tabindex"),R.find("[id]").removeAttr("id"),s||(m.width="100%",f[0].style.width="100%"),P.each(Ct(n,R),function(t,e){r=rt(n,t),e.style.width=n.aoColumns[r].sWidth}),C&&k(function(t){t.style.width=""},a),f=D.outerWidth(),""===s?(w.width="100%",x&&(D.find("tbody").height()>g.offsetHeight||"scroll"==b.css("overflow-y"))&&(w.width=M(D.outerWidth()-l)),f=D.outerWidth()):""!==u&&(w.width=M(u),f=D.outerWidth()),k(t,e),k(function(t){var e=j.getComputedStyle?j.getComputedStyle(t).width:M(P(t).width());F.push(t.innerHTML),A.push(e)},e),k(function(t,e){t.style.width=A[e]},y),P(e).css("height",0),C&&(k(t,a),k(function(t){L.push(t.innerHTML),I.push(M(P(t).css("width")))},a),k(function(t,e){t.style.width=I[e]},i),P(a).height(0)),k(function(t,e){t.innerHTML='<div class="dataTables_sizing">'+F[e]+"</div>",t.childNodes[0].style.height="0",t.childNodes[0].style.overflow="hidden",t.style.width=A[e]},e),C&&k(function(t,e){t.innerHTML='<div class="dataTables_sizing">'+L[e]+"</div>",t.childNodes[0].style.height="0",t.childNodes[0].style.overflow="hidden",t.style.width=I[e]},a),Math.round(D.outerWidth())<Math.round(f)?(o=g.scrollHeight>g.offsetHeight||"scroll"==b.css("overflow-y")?f+l:f,x&&(g.scrollHeight>g.offsetHeight||"scroll"==b.css("overflow-y"))&&(w.width=M(o-l)),""!==s&&""===u||W(n,1,"Possible column misalignment",6)):o="100%",m.width=M(o),d.width=M(o),C&&(n.nScrollFoot.style.width=M(o)),c||x&&(m.height=M(_.offsetHeight+l)),R=D.outerWidth(),h[0].style.width=M(R),p.width=M(R),y=D.height()>g.clientHeight||"scroll"==b.css("overflow-y"),p[i="padding"+(T.bScrollbarLeft?"Left":"Right")]=y?l+"px":"0px",C&&(v[0].style.width=M(R),S[0].style.width=M(R),S[0].style[i]=y?l+"px":"0px"),D.children("colgroup").insertBefore(D.children("thead")),b.trigger("scroll"),!n.bSorted&&!n.bFiltered||n._drawHold||(g.scrollTop=0))}function k(t,e,n){for(var a,r,o=0,i=0,l=e.length;i<l;){for(a=e[i].firstChild,r=n?n[i].firstChild:null;a;)1===a.nodeType&&(n?t(a,r,o):t(a,o),o++),a=a.nextSibling,r=n?r.nextSibling:null;i++}}var te=/<.*?>/g;function ee(t){var e,n,a=t.nTable,r=t.aoColumns,o=t.oScroll,i=o.sY,l=o.sX,o=o.sXInner,s=r.length,u=it(t,"bVisible"),c=P("th",t.nTHead),f=a.getAttribute("width"),d=a.parentNode,h=!1,p=t.oBrowser,g=p.bScrollOversize,b=a.style.width;for(b&&-1!==b.indexOf("%")&&(f=b),D=0;D<u.length;D++)null!==(e=r[u[D]]).sWidth&&(e.sWidth=ae(e.sWidthOrig,d),h=!0);if(g||!h&&!l&&!i&&s==T(t)&&s==c.length)for(D=0;D<s;D++){var m=rt(t,D);null!==m&&(r[m].sWidth=M(c.eq(D).width()))}else{var b=P(a).clone().css("visibility","hidden").removeAttr("id"),S=(b.find("tbody tr").remove(),P("<tr/>").appendTo(b.find("tbody")));for(b.find("thead, tfoot").remove(),b.append(P(t.nTHead).clone()).append(P(t.nTFoot).clone()),b.find("tfoot th, tfoot td").css("width",""),c=Ct(t,b.find("thead")[0]),D=0;D<u.length;D++)e=r[u[D]],c[D].style.width=null!==e.sWidthOrig&&""!==e.sWidthOrig?M(e.sWidthOrig):"",e.sWidthOrig&&l&&P(c[D]).append(P("<div/>").css({width:e.sWidthOrig,margin:0,padding:0,border:0,height:1}));if(t.aoData.length)for(D=0;D<u.length;D++)e=r[n=u[D]],P(re(t,n)).clone(!1).append(e.sContentPadding).appendTo(S);P("[name]",b).removeAttr("name");for(var v=P("<div/>").css(l||i?{position:"absolute",top:0,left:0,height:1,right:0,overflow:"hidden"}:{}).append(b).appendTo(d),y=(l&&o?b.width(o):l?(b.css("width","auto"),b.removeAttr("width"),b.width()<d.clientWidth&&f&&b.width(d.clientWidth)):i?b.width(d.clientWidth):f&&b.width(f),0),D=0;D<u.length;D++){var _=P(c[D]),w=_.outerWidth()-_.width(),_=p.bBounding?Math.ceil(c[D].getBoundingClientRect().width):_.outerWidth();y+=_,r[u[D]].sWidth=M(_-w)}a.style.width=M(y),v.remove()}f&&(a.style.width=M(f)),!f&&!l||t._reszEvt||(o=function(){P(j).on("resize.DT-"+t.sInstance,ne(function(){O(t)}))},g?setTimeout(o,1e3):o(),t._reszEvt=!0)}var ne=w.util.throttle;function ae(t,e){return t?(e=(t=P("<div/>").css("width",M(t)).appendTo(e||v.body))[0].offsetWidth,t.remove(),e):0}function re(t,e){var n,a=oe(t,e);return a<0?null:(n=t.aoData[a]).nTr?n.anCells[e]:P("<td/>").html(S(t,a,e,"display"))[0]}function oe(t,e){for(var n,a=-1,r=-1,o=0,i=t.aoData.length;o<i;o++)(n=(n=(n=S(t,o,e,"display")+"").replace(te,"")).replace(/&nbsp;/g," ")).length>a&&(a=n.length,r=o);return r}function M(t){return null===t?"0px":"number"==typeof t?t<0?"0px":t+"px":t.match(/\d$/)?t+"px":t}function I(t){function e(t){t.length&&!Array.isArray(t[0])?h.push(t):P.merge(h,t)}var n,a,r,o,i,l,s,u=[],c=t.aoColumns,f=t.aaSortingFixed,d=P.isPlainObject(f),h=[];for(Array.isArray(f)&&e(f),d&&f.pre&&e(f.pre),e(t.aaSorting),d&&f.post&&e(f.post),n=0;n<h.length;n++)for(r=(o=c[s=h[n][a=0]].aDataSort).length;a<r;a++)l=c[i=o[a]].sType||"string",h[n]._idx===H&&(h[n]._idx=P.inArray(h[n][1],c[i].asSorting)),u.push({src:s,col:i,dir:h[n][1],index:h[n]._idx,type:l,formatter:w.ext.type.order[l+"-pre"]});return u}function ie(t){var e,n,a,r,c,f=[],u=w.ext.type.order,d=t.aoData,o=(t.aoColumns,0),i=t.aiDisplayMaster;for(lt(t),e=0,n=(c=I(t)).length;e<n;e++)(r=c[e]).formatter&&o++,fe(t,r.col);if("ssp"!=E(t)&&0!==c.length){for(e=0,a=i.length;e<a;e++)f[i[e]]=e;o===c.length?i.sort(function(t,e){for(var n,a,r,o,i=c.length,l=d[t]._aSortData,s=d[e]._aSortData,u=0;u<i;u++)if(0!=(r=(n=l[(o=c[u]).col])<(a=s[o.col])?-1:a<n?1:0))return"asc"===o.dir?r:-r;return(n=f[t])<(a=f[e])?-1:a<n?1:0}):i.sort(function(t,e){for(var n,a,r,o=c.length,i=d[t]._aSortData,l=d[e]._aSortData,s=0;s<o;s++)if(n=i[(r=c[s]).col],a=l[r.col],0!==(r=(u[r.type+"-"+r.dir]||u["string-"+r.dir])(n,a)))return r;return(n=f[t])<(a=f[e])?-1:a<n?1:0})}t.bSorted=!0}function le(t){for(var e=t.aoColumns,n=I(t),a=t.oLanguage.oAria,r=0,o=e.length;r<o;r++){var i=e[r],l=i.asSorting,s=i.ariaTitle||i.sTitle.replace(/<.*?>/g,""),u=i.nTh;u.removeAttribute("aria-sort"),i=i.bSortable?s+("asc"===(0<n.length&&n[0].col==r&&(u.setAttribute("aria-sort","asc"==n[0].dir?"ascending":"descending"),l[n[0].index+1])||l[0])?a.sSortAscending:a.sSortDescending):s,u.setAttribute("aria-label",i)}}function se(t,e,n,a){function r(t,e){var n=t._idx;return(n=n===H?P.inArray(t[1],s):n)+1<s.length?n+1:e?null:0}var o,i=t.aoColumns[e],l=t.aaSorting,s=i.asSorting;"number"==typeof l[0]&&(l=t.aaSorting=[l]),n&&t.oFeatures.bSortMulti?-1!==(i=P.inArray(e,N(l,"0")))?null===(o=null===(o=r(l[i],!0))&&1===l.length?0:o)?l.splice(i,1):(l[i][1]=s[o],l[i]._idx=o):(l.push([e,s[0],0]),l[l.length-1]._idx=0):l.length&&l[0][0]==e?(o=r(l[0]),l.length=1,l[0][1]=s[o],l[0]._idx=o):(l.length=0,l.push([e,s[0]]),l[0]._idx=0),u(t),"function"==typeof a&&a(t)}function ue(e,t,n,a){var r=e.aoColumns[n];me(t,{},function(t){!1!==r.bSortable&&(e.oFeatures.bProcessing?(D(e,!0),setTimeout(function(){se(e,n,t.shiftKey,a),"ssp"!==E(e)&&D(e,!1)},0)):se(e,n,t.shiftKey,a))})}function ce(t){var e,n,a,r=t.aLastSort,o=t.oClasses.sSortColumn,i=I(t),l=t.oFeatures;if(l.bSort&&l.bSortClasses){for(e=0,n=r.length;e<n;e++)a=r[e].src,P(N(t.aoData,"anCells",a)).removeClass(o+(e<2?e+1:3));for(e=0,n=i.length;e<n;e++)a=i[e].src,P(N(t.aoData,"anCells",a)).addClass(o+(e<2?e+1:3))}t.aLastSort=i}function fe(t,e){for(var n,a,r,o=t.aoColumns[e],i=w.ext.order[o.sSortDataType],l=(i&&(n=i.call(t.oInstance,t,e,ot(t,e))),w.ext.type.order[o.sType+"-pre"]),s=0,u=t.aoData.length;s<u;s++)(a=t.aoData[s])._aSortData||(a._aSortData=[]),a._aSortData[e]&&!i||(r=i?n[s]:S(t,s,e,"sort"),a._aSortData[e]=l?l(r):r)}function de(n){var t;n._bLoadingState||(t={time:+new Date,start:n._iDisplayStart,length:n._iDisplayLength,order:P.extend(!0,[],n.aaSorting),search:Et(n.oPreviousSearch),columns:P.map(n.aoColumns,function(t,e){return{visible:t.bVisible,search:Et(n.aoPreSearchCols[e])}})},n.oSavedState=t,R(n,"aoStateSaveParams","stateSaveParams",[n,t]),n.oFeatures.bStateSave&&!n.bDestroying&&n.fnStateSaveCallback.call(n.oInstance,n,t))}function he(e,t,n){var a;if(e.oFeatures.bStateSave)return(a=e.fnStateLoadCallback.call(e.oInstance,e,function(t){pe(e,t,n)}))!==H&&pe(e,a,n),!0;n()}function pe(n,t,e){var a,r,o=n.aoColumns,i=(n._bLoadingState=!0,n._bInitComplete?new w.Api(n):null);if(t&&t.time){var l=R(n,"aoStateLoadParams","stateLoadParams",[n,t]);if(-1!==P.inArray(!1,l))n._bLoadingState=!1;else{l=n.iStateDuration;if(0<l&&t.time<+new Date-1e3*l)n._bLoadingState=!1;else if(t.columns&&o.length!==t.columns.length)n._bLoadingState=!1;else{if(n.oLoadedState=P.extend(!0,{},t),t.length!==H&&(i?i.page.len(t.length):n._iDisplayLength=t.length),t.start!==H&&(null===i?(n._iDisplayStart=t.start,n.iInitDisplayStart=t.start):Yt(n,t.start/n._iDisplayLength)),t.order!==H&&(n.aaSorting=[],P.each(t.order,function(t,e){n.aaSorting.push(e[0]>=o.length?[0,e[1]]:e)})),t.search!==H&&P.extend(n.oPreviousSearch,Bt(t.search)),t.columns){for(a=0,r=t.columns.length;a<r;a++){var s=t.columns[a];s.visible!==H&&(i?i.column(a).visible(s.visible,!1):o[a].bVisible=s.visible),s.search!==H&&P.extend(n.aoPreSearchCols[a],Bt(s.search))}i&&i.columns.adjust()}n._bLoadingState=!1,R(n,"aoStateLoaded","stateLoaded",[n,t])}}}else n._bLoadingState=!1;e()}function ge(t){var e=w.settings,t=P.inArray(t,N(e,"nTable"));return-1!==t?e[t]:null}function W(t,e,n,a){if(n="DataTables warning: "+(t?"table id="+t.sTableId+" - ":"")+n,a&&(n+=". For more information about this error, please see http://datatables.net/tn/"+a),e)j.console&&console.log&&console.log(n);else{e=w.ext,e=e.sErrMode||e.errMode;if(t&&R(t,null,"error",[t,a,n]),"alert"==e)alert(n);else{if("throw"==e)throw new Error(n);"function"==typeof e&&e(t,a,n)}}}function F(n,a,t,e){Array.isArray(t)?P.each(t,function(t,e){Array.isArray(e)?F(n,a,e[0],e[1]):F(n,a,e)}):(e===H&&(e=t),a[t]!==H&&(n[e]=a[t]))}function be(t,e,n){var a,r;for(r in e)e.hasOwnProperty(r)&&(a=e[r],P.isPlainObject(a)?(P.isPlainObject(t[r])||(t[r]={}),P.extend(!0,t[r],a)):n&&"data"!==r&&"aaData"!==r&&Array.isArray(a)?t[r]=a.slice():t[r]=a);return t}function me(e,t,n){P(e).on("click.DT",t,function(t){P(e).trigger("blur"),n(t)}).on("keypress.DT",t,function(t){13===t.which&&(t.preventDefault(),n(t))}).on("selectstart.DT",function(){return!1})}function L(t,e,n,a){n&&t[e].push({fn:n,sName:a})}function R(n,t,e,a){var r=[];return t&&(r=P.map(n[t].slice().reverse(),function(t,e){return t.fn.apply(n.oInstance,a)})),null!==e&&(t=P.Event(e+".dt"),(e=P(n.nTable)).trigger(t,a),0===e.parents("body").length&&P("body").trigger(t,a),r.push(t.result)),r}function Se(t){var e=t._iDisplayStart,n=t.fnDisplayEnd(),a=t._iDisplayLength;n<=e&&(e=n-a),e-=e%a,t._iDisplayStart=e=-1===a||e<0?0:e}function ve(t,e){var t=t.renderer,n=w.ext.renderer[e];return P.isPlainObject(t)&&t[e]?n[t[e]]||n._:"string"==typeof t&&n[t]||n._}function E(t){return t.oFeatures.bServerSide?"ssp":t.ajax||t.sAjaxSource?"ajax":"dom"}function ye(t,n){var a;return Array.isArray(t)?P.map(t,function(t){return ye(t,n)}):"number"==typeof t?[n[t]]:(a=P.map(n,function(t,e){return t.nTable}),P(a).filter(t).map(function(t){var e=P.inArray(this,a);return n[e]}).toArray())}function De(r,o,t){var e,n;t&&(e=new B(r)).one("draw",function(){t(e.ajax.json())}),"ssp"==E(r)?u(r,o):(D(r,!0),(n=r.jqXHR)&&4!==n.readyState&&n.abort(),Tt(r,[],function(t){pt(r);for(var e=Ft(r,t),n=0,a=e.length;n<a;n++)x(r,e[n]);u(r,o),D(r,!1)}))}function _e(t,e,n,a,r){for(var o,i,l,s,u=[],c=typeof e,f=0,d=(e=e&&"string"!=c&&"function"!=c&&e.length!==H?e:[e]).length;f<d;f++)for(l=0,s=(i=e[f]&&e[f].split&&!e[f].match(/[\[\(:]/)?e[f].split(","):[e[f]]).length;l<s;l++)(o=n("string"==typeof i[l]?i[l].trim():i[l]))&&o.length&&(u=u.concat(o));var h=p.selector[t];if(h.length)for(f=0,d=h.length;f<d;f++)u=h[f](a,r,u);return z(u)}function we(t){return(t=t||{}).filter&&t.search===H&&(t.search=t.filter),P.extend({search:"none",order:"current",page:"all"},t)}function Ce(t){for(var e=0,n=t.length;e<n;e++)if(0<t[e].length)return t[0]=t[e],t[0].length=1,t.length=1,t.context=[t.context[e]],t;return t.length=0,t}function Te(o,t,e,n){function i(t,e){var n;if(Array.isArray(t)||t instanceof P)for(var a=0,r=t.length;a<r;a++)i(t[a],e);else t.nodeName&&"tr"===t.nodeName.toLowerCase()?l.push(t):(n=P("<tr><td></td></tr>").addClass(e),P("td",n).addClass(e).html(t)[0].colSpan=T(o),l.push(n[0]))}var l=[];i(e,n),t._details&&t._details.detach(),t._details=P(l),t._detailsShow&&t._details.insertAfter(t.nTr)}function xe(t,e){var n=t.context;if(n.length&&t.length){var a=n[0].aoData[t[0]];if(a._details){(a._detailsShow=e)?(a._details.insertAfter(a.nTr),P(a.nTr).addClass("dt-hasChild")):(a._details.detach(),P(a.nTr).removeClass("dt-hasChild")),R(n[0],null,"childRow",[e,t.row(t[0])]);var s=n[0],r=new B(s),a=".dt.DT_details",e="draw"+a,t="column-sizing"+a,a="destroy"+a,u=s.aoData;if(r.off(e+" "+t+" "+a),N(u,"_details").length>0){r.on(e,function(t,e){if(s!==e)return;r.rows({page:"current"}).eq(0).each(function(t){var e=u[t];if(e._detailsShow)e._details.insertAfter(e.nTr)})});r.on(t,function(t,e,n,a){if(s!==e)return;var r,o=T(e);for(var i=0,l=u.length;i<l;i++){r=u[i];if(r._details)r._details.children("td[colspan]").attr("colspan",o)}});r.on(a,function(t,e){if(s!==e)return;for(var n=0,a=u.length;n<a;n++)if(u[n]._details)Re(r,n)})}Le(n)}}}function Ae(t,e,n,a,r){for(var o=[],i=0,l=r.length;i<l;i++)o.push(S(t,r[i],e));return o}var Ie=[],o=Array.prototype,B=function(t,e){if(!(this instanceof B))return new B(t,e);function n(t){var e,n,a,r;t=t,a=w.settings,r=P.map(a,function(t,e){return t.nTable}),(t=t?t.nTable&&t.oApi?[t]:t.nodeName&&"table"===t.nodeName.toLowerCase()?-1!==(e=P.inArray(t,r))?[a[e]]:null:t&&"function"==typeof t.settings?t.settings().toArray():("string"==typeof t?n=P(t):t instanceof P&&(n=t),n?n.map(function(t){return-1!==(e=P.inArray(this,r))?a[e]:null}).toArray():void 0):[])&&o.push.apply(o,t)}var o=[];if(Array.isArray(t))for(var a=0,r=t.length;a<r;a++)n(t[a]);else n(t);this.context=z(o),e&&P.merge(this,e),this.selector={rows:null,cols:null,opts:null},B.extend(this,this,Ie)},Fe=(w.Api=B,P.extend(B.prototype,{any:function(){return 0!==this.count()},concat:o.concat,context:[],count:function(){return this.flatten().length},each:function(t){for(var e=0,n=this.length;e<n;e++)t.call(this,this[e],e,this);return this},eq:function(t){var e=this.context;return e.length>t?new B(e[t],this[t]):null},filter:function(t){var e=[];if(o.filter)e=o.filter.call(this,t,this);else for(var n=0,a=this.length;n<a;n++)t.call(this,this[n],n,this)&&e.push(this[n]);return new B(this.context,e)},flatten:function(){var t=[];return new B(this.context,t.concat.apply(t,this.toArray()))},join:o.join,indexOf:o.indexOf||function(t,e){for(var n=e||0,a=this.length;n<a;n++)if(this[n]===t)return n;return-1},iterator:function(t,e,n,a){var r,o,i,l,s,u,c,f,d=[],h=this.context,p=this.selector;for("string"==typeof t&&(a=n,n=e,e=t,t=!1),o=0,i=h.length;o<i;o++){var g=new B(h[o]);if("table"===e)(r=n.call(g,h[o],o))!==H&&d.push(r);else if("columns"===e||"rows"===e)(r=n.call(g,h[o],this[o],o))!==H&&d.push(r);else if("column"===e||"column-rows"===e||"row"===e||"cell"===e)for(c=this[o],"column-rows"===e&&(u=Fe(h[o],p.opts)),l=0,s=c.length;l<s;l++)f=c[l],(r="cell"===e?n.call(g,h[o],f.row,f.column,o,l):n.call(g,h[o],f,o,l,u))!==H&&d.push(r)}return d.length||a?((t=(a=new B(h,t?d.concat.apply([],d):d)).selector).rows=p.rows,t.cols=p.cols,t.opts=p.opts,a):this},lastIndexOf:o.lastIndexOf||function(t,e){return this.indexOf.apply(this.toArray.reverse(),arguments)},length:0,map:function(t){var e=[];if(o.map)e=o.map.call(this,t,this);else for(var n=0,a=this.length;n<a;n++)e.push(t.call(this,this[n],n));return new B(this.context,e)},pluck:function(t){var e=w.util.get(t);return this.map(function(t){return e(t)})},pop:o.pop,push:o.push,reduce:o.reduce||function(t,e){return et(this,t,e,0,this.length,1)},reduceRight:o.reduceRight||function(t,e){return et(this,t,e,this.length-1,-1,-1)},reverse:o.reverse,selector:null,shift:o.shift,slice:function(){return new B(this.context,this)},sort:o.sort,splice:o.splice,toArray:function(){return o.slice.call(this)},to$:function(){return P(this)},toJQuery:function(){return P(this)},unique:function(){return new B(this.context,z(this))},unshift:o.unshift}),B.extend=function(t,e,n){if(n.length&&e&&(e instanceof B||e.__dt_wrapper))for(var a,r=0,o=n.length;r<o;r++)e[(a=n[r]).name]="function"===a.type?function(e,n,a){return function(){var t=n.apply(e,arguments);return B.extend(t,t,a.methodExt),t}}(t,a.val,a):"object"===a.type?{}:a.val,e[a.name].__dt_wrapper=!0,B.extend(t,e[a.name],a.propExt)},B.register=e=function(t,e){if(Array.isArray(t))for(var n=0,a=t.length;n<a;n++)B.register(t[n],e);else for(var r=t.split("."),o=Ie,i=0,l=r.length;i<l;i++){var s,u,c=function(t,e){for(var n=0,a=t.length;n<a;n++)if(t[n].name===e)return t[n];return null}(o,u=(s=-1!==r[i].indexOf("()"))?r[i].replace("()",""):r[i]);c||o.push(c={name:u,val:{},methodExt:[],propExt:[],type:"object"}),i===l-1?(c.val=e,c.type="function"==typeof e?"function":P.isPlainObject(e)?"object":"other"):o=s?c.methodExt:c.propExt}},B.registerPlural=t=function(t,e,n){B.register(t,n),B.register(e,function(){var t=n.apply(this,arguments);return t===this?this:t instanceof B?t.length?Array.isArray(t[0])?new B(t.context,t[0]):t[0]:H:t})},e("tables()",function(t){return t!==H&&null!==t?new B(ye(t,this.context)):this}),e("table()",function(t){var t=this.tables(t),e=t.context;return e.length?new B(e[0]):t}),t("tables().nodes()","table().node()",function(){return this.iterator("table",function(t){return t.nTable},1)}),t("tables().body()","table().body()",function(){return this.iterator("table",function(t){return t.nTBody},1)}),t("tables().header()","table().header()",function(){return this.iterator("table",function(t){return t.nTHead},1)}),t("tables().footer()","table().footer()",function(){return this.iterator("table",function(t){return t.nTFoot},1)}),t("tables().containers()","table().container()",function(){return this.iterator("table",function(t){return t.nTableWrapper},1)}),e("draw()",function(e){return this.iterator("table",function(t){"page"===e?y(t):u(t,!1===(e="string"==typeof e?"full-hold"!==e:e))})}),e("page()",function(e){return e===H?this.page.info().page:this.iterator("table",function(t){Yt(t,e)})}),e("page.info()",function(t){var e,n,a,r,o;return 0===this.context.length?H:(n=(e=this.context[0])._iDisplayStart,a=e.oFeatures.bPaginate?e._iDisplayLength:-1,r=e.fnRecordsDisplay(),{page:(o=-1===a)?0:Math.floor(n/a),pages:o?1:Math.ceil(r/a),start:n,end:e.fnDisplayEnd(),length:a,recordsTotal:e.fnRecordsTotal(),recordsDisplay:r,serverSide:"ssp"===E(e)})}),e("page.len()",function(e){return e===H?0!==this.context.length?this.context[0]._iDisplayLength:H:this.iterator("table",function(t){$t(t,e)})}),e("ajax.json()",function(){var t=this.context;if(0<t.length)return t[0].json}),e("ajax.params()",function(){var t=this.context;if(0<t.length)return t[0].oAjaxData}),e("ajax.reload()",function(e,n){return this.iterator("table",function(t){De(t,!1===n,e)})}),e("ajax.url()",function(e){var t=this.context;return e===H?0===t.length?H:(t=t[0]).ajax?P.isPlainObject(t.ajax)?t.ajax.url:t.ajax:t.sAjaxSource:this.iterator("table",function(t){P.isPlainObject(t.ajax)?t.ajax.url=e:t.ajax=e})}),e("ajax.url().load()",function(e,n){return this.iterator("table",function(t){De(t,!1===n,e)})}),function(t,e){var n,a=[],r=t.aiDisplay,o=t.aiDisplayMaster,i=e.search,l=e.order,e=e.page;if("ssp"==E(t))return"removed"===i?[]:f(0,o.length);if("current"==e)for(u=t._iDisplayStart,c=t.fnDisplayEnd();u<c;u++)a.push(r[u]);else if("current"==l||"applied"==l){if("none"==i)a=o.slice();else if("applied"==i)a=r.slice();else if("removed"==i){for(var s={},u=0,c=r.length;u<c;u++)s[r[u]]=null;a=P.map(o,function(t){return s.hasOwnProperty(t)?null:t})}}else if("index"==l||"original"==l)for(u=0,c=t.aoData.length;u<c;u++)("none"==i||-1===(n=P.inArray(u,r))&&"removed"==i||0<=n&&"applied"==i)&&a.push(u);return a}),Le=(e("rows()",function(e,n){e===H?e="":P.isPlainObject(e)&&(n=e,e=""),n=we(n);var t=this.iterator("table",function(t){return _e("row",e,function(n){var t=d(n),a=r.aoData;if(null!==t&&!o)return[t];if(i=i||Fe(r,o),null!==t&&-1!==P.inArray(t,i))return[t];if(null===n||n===H||""===n)return i;if("function"==typeof n)return P.map(i,function(t){var e=a[t];return n(t,e._aData,e.nTr)?t:null});if(n.nodeName)return t=n._DT_RowIndex,e=n._DT_CellIndex,t!==H?a[t]&&a[t].nTr===n?[t]:[]:e?a[e.row]&&a[e.row].nTr===n.parentNode?[e.row]:[]:(t=P(n).closest("*[data-dt-row]")).length?[t.data("dt-row")]:[];if("string"==typeof n&&"#"===n.charAt(0)){var e=r.aIds[n.replace(/^#/,"")];if(e!==H)return[e.idx]}t=_(m(r.aoData,i,"nTr"));return P(t).filter(n).map(function(){return this._DT_RowIndex}).toArray()},r=t,o=n);var r,o,i},1);return t.selector.rows=e,t.selector.opts=n,t}),e("rows().nodes()",function(){return this.iterator("row",function(t,e){return t.aoData[e].nTr||H},1)}),e("rows().data()",function(){return this.iterator(!0,"rows",function(t,e){return m(t.aoData,e,"_aData")},1)}),t("rows().cache()","row().cache()",function(n){return this.iterator("row",function(t,e){t=t.aoData[e];return"search"===n?t._aFilterData:t._aSortData},1)}),t("rows().invalidate()","row().invalidate()",function(n){return this.iterator("row",function(t,e){bt(t,e,n)})}),t("rows().indexes()","row().index()",function(){return this.iterator("row",function(t,e){return e},1)}),t("rows().ids()","row().id()",function(t){for(var e=[],n=this.context,a=0,r=n.length;a<r;a++)for(var o=0,i=this[a].length;o<i;o++){var l=n[a].rowIdFn(n[a].aoData[this[a][o]]._aData);e.push((!0===t?"#":"")+l)}return new B(n,e)}),t("rows().remove()","row().remove()",function(){var f=this;return this.iterator("row",function(t,e,n){var a,r,o,i,l,s,u=t.aoData,c=u[e];for(u.splice(e,1),a=0,r=u.length;a<r;a++)if(s=(l=u[a]).anCells,null!==l.nTr&&(l.nTr._DT_RowIndex=a),null!==s)for(o=0,i=s.length;o<i;o++)s[o]._DT_CellIndex.row=a;gt(t.aiDisplayMaster,e),gt(t.aiDisplay,e),gt(f[n],e,!1),0<t._iRecordsDisplay&&t._iRecordsDisplay--,Se(t);n=t.rowIdFn(c._aData);n!==H&&delete t.aIds[n]}),this.iterator("table",function(t){for(var e=0,n=t.aoData.length;e<n;e++)t.aoData[e].idx=e}),this}),e("rows.add()",function(o){var t=this.iterator("table",function(t){for(var e,n=[],a=0,r=o.length;a<r;a++)(e=o[a]).nodeName&&"TR"===e.nodeName.toUpperCase()?n.push(ut(t,e)[0]):n.push(x(t,e));return n},1),e=this.rows(-1);return e.pop(),P.merge(e,t),e}),e("row()",function(t,e){return Ce(this.rows(t,e))}),e("row().data()",function(t){var e,n=this.context;return t===H?n.length&&this.length?n[0].aoData[this[0]]._aData:H:((e=n[0].aoData[this[0]])._aData=t,Array.isArray(t)&&e.nTr&&e.nTr.id&&b(n[0].rowId)(t,e.nTr.id),bt(n[0],this[0],"data"),this)}),e("row().node()",function(){var t=this.context;return t.length&&this.length&&t[0].aoData[this[0]].nTr||null}),e("row.add()",function(e){e instanceof P&&e.length&&(e=e[0]);var t=this.iterator("table",function(t){return e.nodeName&&"TR"===e.nodeName.toUpperCase()?ut(t,e)[0]:x(t,e)});return this.row(t[0])}),P(v).on("plugin-init.dt",function(t,e){var n=new B(e),a="on-plugin-init",r="stateSaveParams."+a,o="destroy. "+a,a=(n.on(r,function(t,e,n){for(var a=e.rowIdFn,r=e.aoData,o=[],i=0;i<r.length;i++)r[i]._detailsShow&&o.push("#"+a(r[i]._aData));n.childRows=o}),n.on(o,function(){n.off(r+" "+o)}),n.state.loaded());a&&a.childRows&&n.rows(P.map(a.childRows,function(t){return t.replace(/:/g,"\\:")})).every(function(){R(e,null,"requestChild",[this])})}),w.util.throttle(function(t){de(t[0])},500)),Re=function(t,e){var n=t.context;n.length&&(e=n[0].aoData[e!==H?e:t[0]])&&e._details&&(e._details.remove(),e._detailsShow=H,e._details=H,P(e.nTr).removeClass("dt-hasChild"),Le(n))},Pe="row().child",je=Pe+"()",He=(e(je,function(t,e){var n=this.context;return t===H?n.length&&this.length?n[0].aoData[this[0]]._details:H:(!0===t?this.child.show():!1===t?Re(this):n.length&&this.length&&Te(n[0],n[0].aoData[this[0]],t,e),this)}),e([Pe+".show()",je+".show()"],function(t){return xe(this,!0),this}),e([Pe+".hide()",je+".hide()"],function(){return xe(this,!1),this}),e([Pe+".remove()",je+".remove()"],function(){return Re(this),this}),e(Pe+".isShown()",function(){var t=this.context;return t.length&&this.length&&t[0].aoData[this[0]]._detailsShow||!1}),/^([^:]+):(name|visIdx|visible)$/),Ne=(e("columns()",function(n,a){n===H?n="":P.isPlainObject(n)&&(a=n,n=""),a=we(a);var t=this.iterator("table",function(t){return e=n,l=a,s=(i=t).aoColumns,u=N(s,"sName"),c=N(s,"nTh"),_e("column",e,function(n){var a,t=d(n);if(""===n)return f(s.length);if(null!==t)return[0<=t?t:s.length+t];if("function"==typeof n)return a=Fe(i,l),P.map(s,function(t,e){return n(e,Ae(i,e,0,0,a),c[e])?e:null});var r="string"==typeof n?n.match(He):"";if(r)switch(r[2]){case"visIdx":case"visible":var e,o=parseInt(r[1],10);return o<0?[(e=P.map(s,function(t,e){return t.bVisible?e:null}))[e.length+o]]:[rt(i,o)];case"name":return P.map(u,function(t,e){return t===r[1]?e:null});default:return[]}return n.nodeName&&n._DT_CellIndex?[n._DT_CellIndex.column]:(t=P(c).filter(n).map(function(){return P.inArray(this,c)}).toArray()).length||!n.nodeName?t:(t=P(n).closest("*[data-dt-column]")).length?[t.data("dt-column")]:[]},i,l);var i,e,l,s,u,c},1);return t.selector.cols=n,t.selector.opts=a,t}),t("columns().header()","column().header()",function(t,e){return this.iterator("column",function(t,e){return t.aoColumns[e].nTh},1)}),t("columns().footer()","column().footer()",function(t,e){return this.iterator("column",function(t,e){return t.aoColumns[e].nTf},1)}),t("columns().data()","column().data()",function(){return this.iterator("column-rows",Ae,1)}),t("columns().dataSrc()","column().dataSrc()",function(){return this.iterator("column",function(t,e){return t.aoColumns[e].mData},1)}),t("columns().cache()","column().cache()",function(o){return this.iterator("column-rows",function(t,e,n,a,r){return m(t.aoData,r,"search"===o?"_aFilterData":"_aSortData",e)},1)}),t("columns().nodes()","column().nodes()",function(){return this.iterator("column-rows",function(t,e,n,a,r){return m(t.aoData,r,"anCells",e)},1)}),t("columns().visible()","column().visible()",function(f,n){var e=this,t=this.iterator("column",function(t,e){if(f===H)return t.aoColumns[e].bVisible;var n,a,r=e,e=f,o=t.aoColumns,i=o[r],l=t.aoData;if(e===H)i.bVisible;else if(i.bVisible!==e){if(e)for(var s=P.inArray(!0,N(o,"bVisible"),r+1),u=0,c=l.length;u<c;u++)a=l[u].nTr,n=l[u].anCells,a&&a.insertBefore(n[r],n[s]||null);else P(N(t.aoData,"anCells",r)).detach();i.bVisible=e}});return f!==H&&this.iterator("table",function(t){Dt(t,t.aoHeader),Dt(t,t.aoFooter),t.aiDisplay.length||P(t.nTBody).find("td[colspan]").attr("colspan",T(t)),de(t),e.iterator("column",function(t,e){R(t,null,"column-visibility",[t,e,f,n])}),n!==H&&!n||e.columns.adjust()}),t}),t("columns().indexes()","column().index()",function(n){return this.iterator("column",function(t,e){return"visible"===n?ot(t,e):e},1)}),e("columns.adjust()",function(){return this.iterator("table",function(t){O(t)},1)}),e("column.index()",function(t,e){var n;if(0!==this.context.length)return n=this.context[0],"fromVisible"===t||"toData"===t?rt(n,e):"fromData"===t||"toVisible"===t?ot(n,e):void 0}),e("column()",function(t,e){return Ce(this.columns(t,e))}),e("cells()",function(g,t,b){var a,r,o,i,l,s,e;return P.isPlainObject(g)&&(g.row===H?(b=g,g=null):(b=t,t=null)),P.isPlainObject(t)&&(b=t,t=null),null===t||t===H?this.iterator("table",function(t){return a=t,t=g,e=we(b),f=a.aoData,d=Fe(a,e),n=_(m(f,d,"anCells")),h=P(Y([],n)),p=a.aoColumns.length,_e("cell",t,function(t){var e,n="function"==typeof t;if(null===t||t===H||n){for(o=[],i=0,l=d.length;i<l;i++)for(r=d[i],s=0;s<p;s++)u={row:r,column:s},(!n||(c=f[r],t(u,S(a,r,s),c.anCells?c.anCells[s]:null)))&&o.push(u);return o}return P.isPlainObject(t)?t.column!==H&&t.row!==H&&-1!==P.inArray(t.row,d)?[t]:[]:(e=h.filter(t).map(function(t,e){return{row:e._DT_CellIndex.row,column:e._DT_CellIndex.column}}).toArray()).length||!t.nodeName?e:(c=P(t).closest("*[data-dt-row]")).length?[{row:c.data("dt-row"),column:c.data("dt-column")}]:[]},a,e);var a,e,r,o,i,l,s,u,c,f,d,n,h,p}):(e=b?{page:b.page,order:b.order,search:b.search}:{},a=this.columns(t,e),r=this.rows(g,e),e=this.iterator("table",function(t,e){var n=[];for(o=0,i=r[e].length;o<i;o++)for(l=0,s=a[e].length;l<s;l++)n.push({row:r[e][o],column:a[e][l]});return n},1),e=b&&b.selected?this.cells(e,b):e,P.extend(e.selector,{cols:t,rows:g,opts:b}),e)}),t("cells().nodes()","cell().node()",function(){return this.iterator("cell",function(t,e,n){t=t.aoData[e];return t&&t.anCells?t.anCells[n]:H},1)}),e("cells().data()",function(){return this.iterator("cell",function(t,e,n){return S(t,e,n)},1)}),t("cells().cache()","cell().cache()",function(a){return a="search"===a?"_aFilterData":"_aSortData",this.iterator("cell",function(t,e,n){return t.aoData[e][a][n]},1)}),t("cells().render()","cell().render()",function(a){return this.iterator("cell",function(t,e,n){return S(t,e,n,a)},1)}),t("cells().indexes()","cell().index()",function(){return this.iterator("cell",function(t,e,n){return{row:e,column:n,columnVisible:ot(t,n)}},1)}),t("cells().invalidate()","cell().invalidate()",function(a){return this.iterator("cell",function(t,e,n){bt(t,e,a,n)})}),e("cell()",function(t,e,n){return Ce(this.cells(t,e,n))}),e("cell().data()",function(t){var e=this.context,n=this[0];return t===H?e.length&&n.length?S(e[0],n[0].row,n[0].column):H:(ct(e[0],n[0].row,n[0].column,t),bt(e[0],n[0].row,"data",n[0].column),this)}),e("order()",function(e,t){var n=this.context;return e===H?0!==n.length?n[0].aaSorting:H:("number"==typeof e?e=[[e,t]]:e.length&&!Array.isArray(e[0])&&(e=Array.prototype.slice.call(arguments)),this.iterator("table",function(t){t.aaSorting=e.slice()}))}),e("order.listener()",function(e,n,a){return this.iterator("table",function(t){ue(t,e,n,a)})}),e("order.fixed()",function(e){var t;return e?this.iterator("table",function(t){t.aaSortingFixed=P.extend(!0,{},e)}):(t=(t=this.context).length?t[0].aaSortingFixed:H,Array.isArray(t)?{pre:t}:t)}),e(["columns().order()","column().order()"],function(a){var r=this;return this.iterator("table",function(t,e){var n=[];P.each(r[e],function(t,e){n.push([e,a])}),t.aaSorting=n})}),e("search()",function(e,n,a,r){var t=this.context;return e===H?0!==t.length?t[0].oPreviousSearch.sSearch:H:this.iterator("table",function(t){t.oFeatures.bFilter&&Rt(t,P.extend({},t.oPreviousSearch,{sSearch:e+"",bRegex:null!==n&&n,bSmart:null===a||a,bCaseInsensitive:null===r||r}),1)})}),t("columns().search()","column().search()",function(a,r,o,i){return this.iterator("column",function(t,e){var n=t.aoPreSearchCols;if(a===H)return n[e].sSearch;t.oFeatures.bFilter&&(P.extend(n[e],{sSearch:a+"",bRegex:null!==r&&r,bSmart:null===o||o,bCaseInsensitive:null===i||i}),Rt(t,t.oPreviousSearch,1))})}),e("state()",function(){return this.context.length?this.context[0].oSavedState:null}),e("state.clear()",function(){return this.iterator("table",function(t){t.fnStateSaveCallback.call(t.oInstance,t,{})})}),e("state.loaded()",function(){return this.context.length?this.context[0].oLoadedState:null}),e("state.save()",function(){return this.iterator("table",function(t){de(t)})}),w.use=function(t,e){"lib"===e||t.fn?P=t:"win"==e||t.document?v=(j=t).document:"datetime"!==e&&"DateTime"!==t.type||(w.DateTime=t)},w.factory=function(t,e){var n=!1;return t&&t.document&&(v=(j=t).document),e&&e.fn&&e.fn.jquery&&(P=e,n=!0),n},w.versionCheck=w.fnVersionCheck=function(t){for(var e,n,a=w.version.split("."),r=t.split("."),o=0,i=r.length;o<i;o++)if((e=parseInt(a[o],10)||0)!==(n=parseInt(r[o],10)||0))return n<e;return!0},w.isDataTable=w.fnIsDataTable=function(t){var r=P(t).get(0),o=!1;return t instanceof w.Api||(P.each(w.settings,function(t,e){var n=e.nScrollHead?P("table",e.nScrollHead)[0]:null,a=e.nScrollFoot?P("table",e.nScrollFoot)[0]:null;e.nTable!==r&&n!==r&&a!==r||(o=!0)}),o)},w.tables=w.fnTables=function(e){var t=!1,n=(P.isPlainObject(e)&&(t=e.api,e=e.visible),P.map(w.settings,function(t){if(!e||P(t.nTable).is(":visible"))return t.nTable}));return t?new B(n):n},w.camelToHungarian=C,e("$()",function(t,e){e=this.rows(e).nodes(),e=P(e);return P([].concat(e.filter(t).toArray(),e.find(t).toArray()))}),P.each(["on","one","off"],function(t,n){e(n+"()",function(){var t=Array.prototype.slice.call(arguments),e=(t[0]=P.map(t[0].split(/\s/),function(t){return t.match(/\.dt\b/)?t:t+".dt"}).join(" "),P(this.tables().nodes()));return e[n].apply(e,t),this})}),e("clear()",function(){return this.iterator("table",function(t){pt(t)})}),e("settings()",function(){return new B(this.context,this.context)}),e("init()",function(){var t=this.context;return t.length?t[0].oInit:null}),e("data()",function(){return this.iterator("table",function(t){return N(t.aoData,"_aData")}).flatten()}),e("destroy()",function(c){return c=c||!1,this.iterator("table",function(e){var n,t=e.oClasses,a=e.nTable,r=e.nTBody,o=e.nTHead,i=e.nTFoot,l=P(a),r=P(r),s=P(e.nTableWrapper),u=P.map(e.aoData,function(t){return t.nTr}),i=(e.bDestroying=!0,R(e,"aoDestroyCallback","destroy",[e]),c||new B(e).columns().visible(!0),s.off(".DT").find(":not(tbody *)").off(".DT"),P(j).off(".DT-"+e.sInstance),a!=o.parentNode&&(l.children("thead").detach(),l.append(o)),i&&a!=i.parentNode&&(l.children("tfoot").detach(),l.append(i)),e.aaSorting=[],e.aaSortingFixed=[],ce(e),P(u).removeClass(e.asStripeClasses.join(" ")),P("th, td",o).removeClass(t.sSortable+" "+t.sSortableAsc+" "+t.sSortableDesc+" "+t.sSortableNone),r.children().detach(),r.append(u),e.nTableWrapper.parentNode),o=c?"remove":"detach",u=(l[o](),s[o](),!c&&i&&(i.insertBefore(a,e.nTableReinsertBefore),l.css("width",e.sDestroyWidth).removeClass(t.sTable),n=e.asDestroyStripes.length)&&r.children().each(function(t){P(this).addClass(e.asDestroyStripes[t%n])}),P.inArray(e,w.settings));-1!==u&&w.settings.splice(u,1)})}),P.each(["column","row","cell"],function(t,s){e(s+"s().every()",function(o){var i=this.selector.opts,l=this;return this.iterator(s,function(t,e,n,a,r){o.call(l[s](e,"cell"===s?n:i,"cell"===s?i:H),e,n,a,r)})})}),e("i18n()",function(t,e,n){var a=this.context[0],t=A(t)(a.oLanguage);return t===H&&(t=e),"string"==typeof(t=n!==H&&P.isPlainObject(t)?t[n]!==H?t[n]:t._:t)?t.replace("%d",n):t}),w.version="1.13.6",w.settings=[],w.models={},w.models.oSearch={bCaseInsensitive:!0,sSearch:"",bRegex:!1,bSmart:!0,return:!1},w.models.oRow={nTr:null,anCells:null,_aData:[],_aSortData:null,_aFilterData:null,_sFilterRow:null,_sRowStripe:"",src:null,idx:-1},w.models.oColumn={idx:null,aDataSort:null,asSorting:null,bSearchable:null,bSortable:null,bVisible:null,_sManualType:null,_bAttrSrc:!1,fnCreatedCell:null,fnGetData:null,fnSetData:null,mData:null,mRender:null,nTh:null,nTf:null,sClass:null,sContentPadding:null,sDefaultContent:null,sName:null,sSortDataType:"std",sSortingClass:null,sSortingClassJUI:null,sTitle:null,sType:null,sWidth:null,sWidthOrig:null},w.defaults={aaData:null,aaSorting:[[0,"asc"]],aaSortingFixed:[],ajax:null,aLengthMenu:[10,25,50,100],aoColumns:null,aoColumnDefs:null,aoSearchCols:[],asStripeClasses:null,bAutoWidth:!0,bDeferRender:!1,bDestroy:!1,bFilter:!0,bInfo:!0,bLengthChange:!0,bPaginate:!0,bProcessing:!1,bRetrieve:!1,bScrollCollapse:!1,bServerSide:!1,bSort:!0,bSortMulti:!0,bSortCellsTop:!1,bSortClasses:!0,bStateSave:!1,fnCreatedRow:null,fnDrawCallback:null,fnFooterCallback:null,fnFormatNumber:function(t){return t.toString().replace(/\B(?=(\d{3})+(?!\d))/g,this.oLanguage.sThousands)},fnHeaderCallback:null,fnInfoCallback:null,fnInitComplete:null,fnPreDrawCallback:null,fnRowCallback:null,fnServerData:null,fnServerParams:null,fnStateLoadCallback:function(t){try{return JSON.parse((-1===t.iStateDuration?sessionStorage:localStorage).getItem("DataTables_"+t.sInstance+"_"+location.pathname))}catch(t){return{}}},fnStateLoadParams:null,fnStateLoaded:null,fnStateSaveCallback:function(t,e){try{(-1===t.iStateDuration?sessionStorage:localStorage).setItem("DataTables_"+t.sInstance+"_"+location.pathname,JSON.stringify(e))}catch(t){}},fnStateSaveParams:null,iStateDuration:7200,iDeferLoading:null,iDisplayLength:10,iDisplayStart:0,iTabIndex:0,oClasses:{},oLanguage:{oAria:{sSortAscending:": activate to sort column ascending",sSortDescending:": activate to sort column descending"},oPaginate:{sFirst:"First",sLast:"Last",sNext:"Next",sPrevious:"Previous"},sEmptyTable:"No data available in table",sInfo:"Showing _START_ to _END_ of _TOTAL_ entries",sInfoEmpty:"Showing 0 to 0 of 0 entries",sInfoFiltered:"(filtered from _MAX_ total entries)",sInfoPostFix:"",sDecimal:"",sThousands:",",sLengthMenu:"Show _MENU_ entries",sLoadingRecords:"Loading...",sProcessing:"",sSearch:"Search:",sSearchPlaceholder:"",sUrl:"",sZeroRecords:"No matching records found"},oSearch:P.extend({},w.models.oSearch),sAjaxDataProp:"data",sAjaxSource:null,sDom:"lfrtip",searchDelay:null,sPaginationType:"simple_numbers",sScrollX:"",sScrollXInner:"",sScrollY:"",sServerMethod:"GET",renderer:null,rowId:"DT_RowId"},i(w.defaults),w.defaults.column={aDataSort:null,iDataSort:-1,asSorting:["asc","desc"],bSearchable:!0,bSortable:!0,bVisible:!0,fnCreatedCell:null,mData:null,mRender:null,sCellType:"td",sClass:"",sContentPadding:"",sDefaultContent:null,sName:"",sSortDataType:"std",sTitle:null,sType:null,sWidth:null},i(w.defaults.column),w.models.oSettings={oFeatures:{bAutoWidth:null,bDeferRender:null,bFilter:null,bInfo:null,bLengthChange:null,bPaginate:null,bProcessing:null,bServerSide:null,bSort:null,bSortMulti:null,bSortClasses:null,bStateSave:null},oScroll:{bCollapse:null,iBarWidth:0,sX:null,sXInner:null,sY:null},oLanguage:{fnInfoCallback:null},oBrowser:{bScrollOversize:!1,bScrollbarLeft:!1,bBounding:!1,barWidth:0},ajax:null,aanFeatures:[],aoData:[],aiDisplay:[],aiDisplayMaster:[],aIds:{},aoColumns:[],aoHeader:[],aoFooter:[],oPreviousSearch:{},aoPreSearchCols:[],aaSorting:null,aaSortingFixed:[],asStripeClasses:null,asDestroyStripes:[],sDestroyWidth:0,aoRowCallback:[],aoHeaderCallback:[],aoFooterCallback:[],aoDrawCallback:[],aoRowCreatedCallback:[],aoPreDrawCallback:[],aoInitComplete:[],aoStateSaveParams:[],aoStateLoadParams:[],aoStateLoaded:[],sTableId:"",nTable:null,nTHead:null,nTFoot:null,nTBody:null,nTableWrapper:null,bDeferLoading:!1,bInitialised:!1,aoOpenRows:[],sDom:null,searchDelay:null,sPaginationType:"two_button",iStateDuration:0,aoStateSave:[],aoStateLoad:[],oSavedState:null,oLoadedState:null,sAjaxSource:null,sAjaxDataProp:null,jqXHR:null,json:H,oAjaxData:H,fnServerData:null,aoServerParams:[],sServerMethod:null,fnFormatNumber:null,aLengthMenu:null,iDraw:0,bDrawing:!1,iDrawError:-1,_iDisplayLength:10,_iDisplayStart:0,_iRecordsTotal:0,_iRecordsDisplay:0,oClasses:{},bFiltered:!1,bSorted:!1,bSortCellsTop:null,oInit:null,aoDestroyCallback:[],fnRecordsTotal:function(){return"ssp"==E(this)?+this._iRecordsTotal:this.aiDisplayMaster.length},fnRecordsDisplay:function(){return"ssp"==E(this)?+this._iRecordsDisplay:this.aiDisplay.length},fnDisplayEnd:function(){var t=this._iDisplayLength,e=this._iDisplayStart,n=e+t,a=this.aiDisplay.length,r=this.oFeatures,o=r.bPaginate;return r.bServerSide?!1===o||-1===t?e+a:Math.min(e+t,this._iRecordsDisplay):!o||a<n||-1===t?a:n},oInstance:null,sInstance:null,iTabIndex:0,nScrollHead:null,nScrollFoot:null,aLastSort:[],oPlugins:{},rowIdFn:null,rowId:null},w.ext=p={buttons:{},classes:{},builder:"-source-",errMode:"alert",feature:[],search:[],selector:{cell:[],column:[],row:[]},internal:{},legacy:{ajax:null},pager:{},renderer:{pageButton:{},header:{}},order:{},type:{detect:[],search:{},order:{}},_unique:0,fnVersionCheck:w.fnVersionCheck,iApiIndex:0,oJUIClasses:{},sVersion:w.version},P.extend(p,{afnFiltering:p.search,aTypes:p.type.detect,ofnSearch:p.type.search,oSort:p.type.order,afnSortData:p.order,aoFeatures:p.feature,oApi:p.internal,oStdClasses:p.classes,oPagination:p.pager}),P.extend(w.ext.classes,{sTable:"dataTable",sNoFooter:"no-footer",sPageButton:"paginate_button",sPageButtonActive:"current",sPageButtonDisabled:"disabled",sStripeOdd:"odd",sStripeEven:"even",sRowEmpty:"dataTables_empty",sWrapper:"dataTables_wrapper",sFilter:"dataTables_filter",sInfo:"dataTables_info",sPaging:"dataTables_paginate paging_",sLength:"dataTables_length",sProcessing:"dataTables_processing",sSortAsc:"sorting_asc",sSortDesc:"sorting_desc",sSortable:"sorting",sSortableAsc:"sorting_desc_disabled",sSortableDesc:"sorting_asc_disabled",sSortableNone:"sorting_disabled",sSortColumn:"sorting_",sFilterInput:"",sLengthSelect:"",sScrollWrapper:"dataTables_scroll",sScrollHead:"dataTables_scrollHead",sScrollHeadInner:"dataTables_scrollHeadInner",sScrollBody:"dataTables_scrollBody",sScrollFoot:"dataTables_scrollFoot",sScrollFootInner:"dataTables_scrollFootInner",sHeaderTH:"",sFooterTH:"",sSortJUIAsc:"",sSortJUIDesc:"",sSortJUI:"",sSortJUIAscAllowed:"",sSortJUIDescAllowed:"",sSortJUIWrapper:"",sSortIcon:"",sJUIHeader:"",sJUIFooter:""}),w.ext.pager);function Oe(t,e){var n=[],a=Ne.numbers_length,r=Math.floor(a/2);return e<=a?n=f(0,e):t<=r?((n=f(0,a-2)).push("ellipsis"),n.push(e-1)):((e-1-r<=t?n=f(e-(a-2),e):((n=f(t-r+2,t+r-1)).push("ellipsis"),n.push(e-1),n)).splice(0,0,"ellipsis"),n.splice(0,0,0)),n.DT_el="span",n}P.extend(Ne,{simple:function(t,e){return["previous","next"]},full:function(t,e){return["first","previous","next","last"]},numbers:function(t,e){return[Oe(t,e)]},simple_numbers:function(t,e){return["previous",Oe(t,e),"next"]},full_numbers:function(t,e){return["first","previous",Oe(t,e),"next","last"]},first_last_numbers:function(t,e){return["first",Oe(t,e),"last"]},_numbers:Oe,numbers_length:7}),P.extend(!0,w.ext.renderer,{pageButton:{_:function(u,t,c,e,f,d){function h(t,e){for(var n,a=b.sPageButtonDisabled,r=function(t){Yt(u,t.data.action,!0)},o=0,i=e.length;o<i;o++)if(n=e[o],Array.isArray(n)){var l=P("<"+(n.DT_el||"div")+"/>").appendTo(t);h(l,n)}else{var s=!1;switch(p=null,g=n){case"ellipsis":t.append('<span class="ellipsis">&#x2026;</span>');break;case"first":p=m.sFirst,0===f&&(s=!0);break;case"previous":p=m.sPrevious,0===f&&(s=!0);break;case"next":p=m.sNext,0!==d&&f!==d-1||(s=!0);break;case"last":p=m.sLast,0!==d&&f!==d-1||(s=!0);break;default:p=u.fnFormatNumber(n+1),g=f===n?b.sPageButtonActive:""}null!==p&&(l=u.oInit.pagingTag||"a",s&&(g+=" "+a),me(P("<"+l+">",{class:b.sPageButton+" "+g,"aria-controls":u.sTableId,"aria-disabled":s?"true":null,"aria-label":S[n],role:"link","aria-current":g===b.sPageButtonActive?"page":null,"data-dt-idx":n,tabindex:s?-1:u.iTabIndex,id:0===c&&"string"==typeof n?u.sTableId+"_"+n:null}).html(p).appendTo(t),{action:n},r))}}var p,g,n,b=u.oClasses,m=u.oLanguage.oPaginate,S=u.oLanguage.oAria.paginate||{};try{n=P(t).find(v.activeElement).data("dt-idx")}catch(t){}h(P(t).empty(),e),n!==H&&P(t).find("[data-dt-idx="+n+"]").trigger("focus")}}}),P.extend(w.ext.type.detect,[function(t,e){e=e.oLanguage.sDecimal;return l(t,e)?"num"+e:null},function(t,e){var n;return(!t||t instanceof Date||X.test(t))&&(null!==(n=Date.parse(t))&&!isNaN(n)||h(t))?"date":null},function(t,e){e=e.oLanguage.sDecimal;return l(t,e,!0)?"num-fmt"+e:null},function(t,e){e=e.oLanguage.sDecimal;return a(t,e)?"html-num"+e:null},function(t,e){e=e.oLanguage.sDecimal;return a(t,e,!0)?"html-num-fmt"+e:null},function(t,e){return h(t)||"string"==typeof t&&-1!==t.indexOf("<")?"html":null}]),P.extend(w.ext.type.search,{html:function(t){return h(t)?t:"string"==typeof t?t.replace(U," ").replace(V,""):""},string:function(t){return!h(t)&&"string"==typeof t?t.replace(U," "):t}});function ke(t,e,n,a){var r;return 0===t||t&&"-"!==t?"number"==(r=typeof t)||"bigint"==r?t:+(t=(t=e?$(t,e):t).replace&&(n&&(t=t.replace(n,"")),a)?t.replace(a,""):t):-1/0}function Me(n){P.each({num:function(t){return ke(t,n)},"num-fmt":function(t){return ke(t,n,q)},"html-num":function(t){return ke(t,n,V)},"html-num-fmt":function(t){return ke(t,n,V,q)}},function(t,e){p.type.order[t+n+"-pre"]=e,t.match(/^html\-/)&&(p.type.search[t+n]=p.type.search.html)})}P.extend(p.type.order,{"date-pre":function(t){t=Date.parse(t);return isNaN(t)?-1/0:t},"html-pre":function(t){return h(t)?"":t.replace?t.replace(/<.*?>/g,"").toLowerCase():t+""},"string-pre":function(t){return h(t)?"":"string"==typeof t?t.toLowerCase():t.toString?t.toString():""},"string-asc":function(t,e){return t<e?-1:e<t?1:0},"string-desc":function(t,e){return t<e?1:e<t?-1:0}}),Me(""),P.extend(!0,w.ext.renderer,{header:{_:function(r,o,i,l){P(r.nTable).on("order.dt.DT",function(t,e,n,a){r===e&&(e=i.idx,o.removeClass(l.sSortAsc+" "+l.sSortDesc).addClass("asc"==a[e]?l.sSortAsc:"desc"==a[e]?l.sSortDesc:i.sSortingClass))})},jqueryui:function(r,o,i,l){P("<div/>").addClass(l.sSortJUIWrapper).append(o.contents()).append(P("<span/>").addClass(l.sSortIcon+" "+i.sSortingClassJUI)).appendTo(o),P(r.nTable).on("order.dt.DT",function(t,e,n,a){r===e&&(e=i.idx,o.removeClass(l.sSortAsc+" "+l.sSortDesc).addClass("asc"==a[e]?l.sSortAsc:"desc"==a[e]?l.sSortDesc:i.sSortingClass),o.find("span."+l.sSortIcon).removeClass(l.sSortJUIAsc+" "+l.sSortJUIDesc+" "+l.sSortJUI+" "+l.sSortJUIAscAllowed+" "+l.sSortJUIDescAllowed).addClass("asc"==a[e]?l.sSortJUIAsc:"desc"==a[e]?l.sSortJUIDesc:i.sSortingClassJUI))})}}});function We(t){return"string"==typeof(t=Array.isArray(t)?t.join(","):t)?t.replace(/&/g,"&amp;").replace(/</g,"&lt;").replace(/>/g,"&gt;").replace(/"/g,"&quot;"):t}function Ee(t,e,n,a,r){return j.moment?t[e](r):j.luxon?t[n](r):a?t[a](r):t}var Be=!1;function Ue(t,e,n){var a;if(j.moment){if(!(a=j.moment.utc(t,e,n,!0)).isValid())return null}else if(j.luxon){if(!(a=e&&"string"==typeof t?j.luxon.DateTime.fromFormat(t,e):j.luxon.DateTime.fromISO(t)).isValid)return null;a.setLocale(n)}else e?(Be||alert("DataTables warning: Formatted date without Moment.js or Luxon - https://datatables.net/tn/17"),Be=!0):a=new Date(t);return a}function Ve(s){return function(a,r,o,i){0===arguments.length?(o="en",a=r=null):1===arguments.length?(o="en",r=a,a=null):2===arguments.length&&(o=r,r=a,a=null);var l="datetime-"+r;return w.ext.type.order[l]||(w.ext.type.detect.unshift(function(t){return t===l&&l}),w.ext.type.order[l+"-asc"]=function(t,e){t=t.valueOf(),e=e.valueOf();return t===e?0:t<e?-1:1},w.ext.type.order[l+"-desc"]=function(t,e){t=t.valueOf(),e=e.valueOf();return t===e?0:e<t?-1:1}),function(t,e){var n;return null!==t&&t!==H||(t="--now"===i?(n=new Date,new Date(Date.UTC(n.getFullYear(),n.getMonth(),n.getDate(),n.getHours(),n.getMinutes(),n.getSeconds()))):""),"type"===e?l:""===t?"sort"!==e?"":Ue("0000-01-01 00:00:00",null,o):!(null===r||a!==r||"sort"===e||"type"===e||t instanceof Date)||null===(n=Ue(t,a,o))?t:"sort"===e?n:(t=null===r?Ee(n,"toDate","toJSDate","")[s]():Ee(n,"format","toFormat","toISOString",r),"display"===e?We(t):t)}}}var Xe=",",Je=".";if(j.Intl!==H)try{for(var qe=(new Intl.NumberFormat).formatToParts(100000.1),n=0;n<qe.length;n++)"group"===qe[n].type?Xe=qe[n].value:"decimal"===qe[n].type&&(Je=qe[n].value)}catch(t){}function $e(e){return function(){var t=[ge(this[w.ext.iApiIndex])].concat(Array.prototype.slice.call(arguments));return w.ext.internal[e].apply(this,t)}}return w.datetime=function(n,a){var r="datetime-detect-"+n;a=a||"en",w.ext.type.order[r]||(w.ext.type.detect.unshift(function(t){var e=Ue(t,n,a);return!(""!==t&&!e)&&r}),w.ext.type.order[r+"-pre"]=function(t){return Ue(t,n,a)||0})},w.render={date:Ve("toLocaleDateString"),datetime:Ve("toLocaleString"),time:Ve("toLocaleTimeString"),number:function(a,r,o,i,l){return null!==a&&a!==H||(a=Xe),null!==r&&r!==H||(r=Je),{display:function(t){if("number"!=typeof t&&"string"!=typeof t)return t;if(""===t||null===t)return t;var e=t<0?"-":"",n=parseFloat(t);if(isNaN(n))return We(t);n=n.toFixed(o),t=Math.abs(n);n=parseInt(t,10),t=o?r+(t-n).toFixed(o).substring(2):"";return(e=0===n&&0===parseFloat(t)?"":e)+(i||"")+n.toString().replace(/\B(?=(\d{3})+(?!\d))/g,a)+t+(l||"")}}},text:function(){return{display:We,filter:We}}},P.extend(w.ext.internal,{_fnExternApiFunc:$e,_fnBuildAjax:Tt,_fnAjaxUpdate:xt,_fnAjaxParameters:At,_fnAjaxUpdateDraw:It,_fnAjaxDataSrc:Ft,_fnAddColumn:nt,_fnColumnOptions:at,_fnAdjustColumnSizing:O,_fnVisibleToColumnIndex:rt,_fnColumnIndexToVisible:ot,_fnVisbleColumns:T,_fnGetColumns:it,_fnColumnTypes:lt,_fnApplyColumnDefs:st,_fnHungarianMap:i,_fnCamelToHungarian:C,_fnLanguageCompat:Z,_fnBrowserDetect:tt,_fnAddData:x,_fnAddTr:ut,_fnNodeToDataIndex:function(t,e){return e._DT_RowIndex!==H?e._DT_RowIndex:null},_fnNodeToColumnIndex:function(t,e,n){return P.inArray(n,t.aoData[e].anCells)},_fnGetCellData:S,_fnSetCellData:ct,_fnSplitObjNotation:dt,_fnGetObjectDataFn:A,_fnSetObjectDataFn:b,_fnGetDataMaster:ht,_fnClearTable:pt,_fnDeleteIndex:gt,_fnInvalidate:bt,_fnGetRowElements:mt,_fnCreateTr:St,_fnBuildHead:yt,_fnDrawHead:Dt,_fnDraw:y,_fnReDraw:u,_fnAddOptionsHtml:_t,_fnDetectHeader:wt,_fnGetUniqueThs:Ct,_fnFeatureHtmlFilter:Lt,_fnFilterComplete:Rt,_fnFilterCustom:Pt,_fnFilterColumn:jt,_fnFilter:Ht,_fnFilterCreateSearch:Nt,_fnEscapeRegex:Ot,_fnFilterData:Wt,_fnFeatureHtmlInfo:Ut,_fnUpdateInfo:Vt,_fnInfoMacros:Xt,_fnInitialise:Jt,_fnInitComplete:qt,_fnLengthChange:$t,_fnFeatureHtmlLength:Gt,_fnFeatureHtmlPaginate:zt,_fnPageChange:Yt,_fnFeatureHtmlProcessing:Zt,_fnProcessingDisplay:D,_fnFeatureHtmlTable:Kt,_fnScrollDraw:Qt,_fnApplyToChildren:k,_fnCalculateColumnWidths:ee,_fnThrottle:ne,_fnConvertToWidth:ae,_fnGetWidestNode:re,_fnGetMaxLenString:oe,_fnStringToCss:M,_fnSortFlatten:I,_fnSort:ie,_fnSortAria:le,_fnSortListener:se,_fnSortAttachListener:ue,_fnSortingClasses:ce,_fnSortData:fe,_fnSaveState:de,_fnLoadState:he,_fnImplementState:pe,_fnSettingsFromNode:ge,_fnLog:W,_fnMap:F,_fnBindAction:me,_fnCallbackReg:L,_fnCallbackFire:R,_fnLengthOverflow:Se,_fnRenderer:ve,_fnDataSource:E,_fnRowAttributes:vt,_fnExtend:be,_fnCalculateEnd:function(){}}),((P.fn.dataTable=w).$=P).fn.dataTableSettings=w.settings,P.fn.dataTableExt=w.ext,P.fn.DataTable=function(t){return P(this).dataTable(t).api()},P.each(w,function(t,e){P.fn.DataTable[t]=e}),w});
```
