## supplement_file1 for "GENETH’OFF: a flexible workflow for genome-wide profiling of CRISPR/Cas off-targets": buttons.colVis.min.js

```
/*!
 * Column visibility buttons for Buttons and DataTables.
 * © SpryMedia Ltd - datatables.net/license
 */
!function(t){var o,i;"function"==typeof define&&define.amd?define(["jquery","datatables.net","datatables.net-buttons"],function(n){return t(n,window,document)}):"object"==typeof exports?(o=require("jquery"),i=function(n,e){e.fn.dataTable||require("datatables.net")(n,e),e.fn.dataTable.Buttons||require("datatables.net-buttons")(n,e)},"undefined"==typeof window?module.exports=function(n,e){return n=n||window,e=e||o(n),i(n,e),t(e,0,n.document)}:(i(window,o),module.exports=t(o,window,window.document))):t(jQuery,window,document)}(function(n,e,t,l){"use strict";var o=n.fn.dataTable;return n.extend(o.ext.buttons,{colvis:function(o,i){var l=null,n={extend:"collection",init:function(n,e){l=e},text:function(n){return n.i18n("buttons.colvis","Column visibility")},className:"buttons-colvis",closeButton:!1,buttons:[{extend:"columnsToggle",columns:i.columns,columnText:i.columnText}]};return o.on("column-reorder.dt"+i.namespace,function(n,e,t){o.button(null,o.button(null,l).node()).collectionRebuild([{extend:"columnsToggle",columns:i.columns,columnText:i.columnText}])}),n},columnsToggle:function(n,e){return n.columns(e.columns).indexes().map(function(n){return{extend:"columnToggle",columns:n,columnText:e.columnText}}).toArray()},columnToggle:function(n,e){return{extend:"columnVisibility",columns:e.columns,columnText:e.columnText}},columnsVisibility:function(n,e){return n.columns(e.columns).indexes().map(function(n){return{extend:"columnVisibility",columns:n,visibility:e.visibility,columnText:e.columnText}}).toArray()},columnVisibility:{columns:l,text:function(n,e,t){return t._columnText(n,t)},className:"buttons-columnVisibility",action:function(n,e,t,o){var e=e.columns(o.columns),i=e.visible();e.visible(o.visibility!==l?o.visibility:!(i.length&&i[0]))},init:function(o,n,i){var l=this;n.attr("data-cv-idx",i.columns),o.on("column-visibility.dt"+i.namespace,function(n,e){e.bDestroying||e.nTable!=o.settings()[0].nTable||l.active(o.column(i.columns).visible())}).on("column-reorder.dt"+i.namespace,function(n,e,t){i.destroying||1===o.columns(i.columns).count()&&(l.text(i._columnText(o,i)),l.active(o.column(i.columns).visible()))}),this.active(o.column(i.columns).visible())},destroy:function(n,e,t){n.off("column-visibility.dt"+t.namespace).off("column-reorder.dt"+t.namespace)},_columnText:function(n,e){var t=n.column(e.columns).index(),o=n.settings()[0].aoColumns[t].sTitle;return o=(o=o||n.column(t).header().innerHTML).replace(/\n/g," ").replace(/<br\s*\/?>/gi," ").replace(/<select(.*?)<\/select>/g,"").replace(/<!\-\-.*?\-\->/g,"").replace(/<.*?>/g,"").replace(/^\s+|\s+$/g,""),e.columnText?e.columnText(n,t,o):o}},colvisRestore:{className:"buttons-colvisRestore",text:function(n){return n.i18n("buttons.colvisRestore","Restore visibility")},init:function(e,n,t){t._visOriginal=e.columns().indexes().map(function(n){return e.column(n).visible()}).toArray()},action:function(n,e,t,o){e.columns().every(function(n){n=e.colReorder&&e.colReorder.transpose?e.colReorder.transpose(n,"toOriginal"):n;this.visible(o._visOriginal[n])})}},colvisGroup:{className:"buttons-colvisGroup",action:function(n,e,t,o){e.columns(o.show).visible(!0,!1),e.columns(o.hide).visible(!1,!1),e.columns.adjust()},show:[],hide:[]}}),o});
```
