## supplement_file1 for "GENETH’OFF: a flexible workflow for genome-wide profiling of CRISPR/Cas off-targets": buttons.html5.min.js

```
/*!
 * HTML5 export buttons for Buttons and DataTables.
 * © SpryMedia Ltd - datatables.net/license
 *
 * FileSaver.js (1.3.3) - MIT license
 * Copyright © 2016 Eli Grey - http://eligrey.com
 */
!function(n){var r,a;"function"==typeof define&&define.amd?define(["jquery","datatables.net","datatables.net-buttons"],function(t){return n(t,window,document)}):"object"==typeof exports?(r=require("jquery"),a=function(t,e){e.fn.dataTable||require("datatables.net")(t,e),e.fn.dataTable.Buttons||require("datatables.net-buttons")(t,e)},"undefined"==typeof window?module.exports=function(t,e,o,l){return t=t||window,e=e||r(t),a(t,e),n(e,t,t.document)}:(a(window,r),module.exports=n(r,window,window.document))):n(jQuery,window,document)}(function(C,T,y,t,e,k){"use strict";var o,l,n=C.fn.dataTable;function S(){return o||T.JSZip}function s(){return l||T.pdfMake}n.Buttons.pdfMake=function(t){if(!t)return s();l=t},n.Buttons.jszip=function(t){if(!t)return S();o=t};function N(t){var e="Sheet1";return e=t.sheetName?t.sheetName.replace(/[\[\]\*\/\\\?\:]/g,""):e}function u(t,e){for(var o=c(e),l=t.buttons.exportData(e.exportOptions),n=e.fieldBoundary,r=e.fieldSeparator,a=new RegExp(n,"g"),d=e.escapeChar!==k?e.escapeChar:"\\",p=function(t){for(var e="",o=0,l=t.length;o<l;o++)0<o&&(e+=r),e+=n?n+(""+t[o]).replace(a,d+n)+n:t[o];return e},t=e.header?p(l.header)+o:"",e=e.footer&&l.footer?o+p(l.footer):"",i=[],s=0,f=l.body.length;s<f;s++)i.push(p(l.body[s]));return{str:t+i.join(o)+e,rows:i.length}}function f(){var t;return-1!==navigator.userAgent.indexOf("Safari")&&-1===navigator.userAgent.indexOf("Chrome")&&-1===navigator.userAgent.indexOf("Opera")&&!!((t=navigator.userAgent.match(/AppleWebKit\/(\d+\.\d+)/))&&1<t.length&&+t[1]<603.1)}var O=function(d){var p,i,s,f,m,y,e,u,c,l,t;if(!(void 0===d||"undefined"!=typeof navigator&&/MSIE [1-9]\./.test(navigator.userAgent)))return t=d.document,p=function(){return d.URL||d.webkitURL||d},i=t.createElementNS("http://www.w3.org/1999/xhtml","a"),s="download"in i,f=/constructor/i.test(d.HTMLElement)||d.safari,m=/CriOS\/[\d]+/.test(navigator.userAgent),y=function(t){(d.setImmediate||d.setTimeout)(function(){throw t},0)},e=4e4,u=function(t){setTimeout(function(){"string"==typeof t?p().revokeObjectURL(t):t.remove()},e)},c=function(t){return/^\s*(?:text\/\S*|application\/xml|\S*\/\S*\+xml)\s*;.*charset\s*=\s*utf-8/i.test(t.type)?new Blob([String.fromCharCode(65279),t],{type:t.type}):t},t=(l=function(t,o,e){e||(t=c(t));var l,n,r=this,e="application/octet-stream"===t.type,a=function(){for(var t=r,e="writestart progress write writeend".split(" "),o=void 0,l=(e=[].concat(e)).length;l--;){var n=t["on"+e[l]];if("function"==typeof n)try{n.call(t,o||t)}catch(t){y(t)}}};r.readyState=r.INIT,s?(l=p().createObjectURL(t),setTimeout(function(){var t,e;i.href=l,i.download=o,t=i,e=new MouseEvent("click"),t.dispatchEvent(e),a(),u(l),r.readyState=r.DONE})):(m||e&&f)&&d.FileReader?((n=new FileReader).onloadend=function(){var t=m?n.result:n.result.replace(/^data:[^;]*;/,"data:attachment/file;");d.open(t,"_blank")||(d.location.href=t),r.readyState=r.DONE,a()},n.readAsDataURL(t),r.readyState=r.INIT):(l=l||p().createObjectURL(t),!e&&d.open(l,"_blank")||(d.location.href=l),r.readyState=r.DONE,a(),u(l))}).prototype,"undefined"!=typeof navigator&&navigator.msSaveOrOpenBlob?function(t,e,o){return e=e||t.name||"download",o||(t=c(t)),navigator.msSaveOrOpenBlob(t,e)}:(t.abort=function(){},t.readyState=t.INIT=0,t.WRITING=1,t.DONE=2,t.error=t.onwritestart=t.onprogress=t.onwrite=t.onabort=t.onerror=t.onwriteend=null,function(t,e,o){return new l(t,e||t.name||"download",o)})}("undefined"!=typeof self&&self||void 0!==T&&T||this.content),c=(n.fileSave=O,function(t){return t.newline||(navigator.userAgent.match(/Windows/)?"\r\n":"\n")});function z(t){for(var e="A".charCodeAt(0),o="Z".charCodeAt(0)-e+1,l="";0<=t;)l=String.fromCharCode(t%o+e)+l,t=Math.floor(t/o)-1;return l}try{var D,A=new XMLSerializer}catch(t){}function E(t,e,o){var l=t.createElement(e);return o&&(o.attr&&C(l).attr(o.attr),o.children&&C.each(o.children,function(t,e){l.appendChild(e)}),null!==o.text)&&o.text!==k&&l.appendChild(t.createTextNode(o.text)),l}var _={"_rels/.rels":'<?xml version="1.0" encoding="UTF-8" standalone="yes"?><Relationships xmlns="http://schemas.openxmlformats.org/package/2006/relationships"><Relationship Id="rId1" Type="http://schemas.openxmlformats.org/officeDocument/2006/relationships/officeDocument" Target="xl/workbook.xml"/></Relationships>',"xl/_rels/workbook.xml.rels":'<?xml version="1.0" encoding="UTF-8" standalone="yes"?><Relationships xmlns="http://schemas.openxmlformats.org/package/2006/relationships"><Relationship Id="rId1" Type="http://schemas.openxmlformats.org/officeDocument/2006/relationships/worksheet" Target="worksheets/sheet1.xml"/><Relationship Id="rId2" Type="http://schemas.openxmlformats.org/officeDocument/2006/relationships/styles" Target="styles.xml"/></Relationships>',"[Content_Types].xml":'<?xml version="1.0" encoding="UTF-8" standalone="yes"?><Types xmlns="http://schemas.openxmlformats.org/package/2006/content-types"><Default Extension="xml" ContentType="application/xml" /><Default Extension="rels" ContentType="application/vnd.openxmlformats-package.relationships+xml" /><Default Extension="jpeg" ContentType="image/jpeg" /><Override PartName="/xl/workbook.xml" ContentType="application/vnd.openxmlformats-officedocument.spreadsheetml.sheet.main+xml" /><Override PartName="/xl/worksheets/sheet1.xml" ContentType="application/vnd.openxmlformats-officedocument.spreadsheetml.worksheet+xml" /><Override PartName="/xl/styles.xml" ContentType="application/vnd.openxmlformats-officedocument.spreadsheetml.styles+xml" /></Types>',"xl/workbook.xml":'<?xml version="1.0" encoding="UTF-8" standalone="yes"?><workbook xmlns="http://schemas.openxmlformats.org/spreadsheetml/2006/main" xmlns:r="http://schemas.openxmlformats.org/officeDocument/2006/relationships"><fileVersion appName="xl" lastEdited="5" lowestEdited="5" rupBuild="24816"/><workbookPr showInkAnnotation="0" autoCompressPictures="0"/><bookViews><workbookView xWindow="0" yWindow="0" windowWidth="25600" windowHeight="19020" tabRatio="500"/></bookViews><sheets><sheet name="Sheet1" sheetId="1" r:id="rId1"/></sheets><definedNames/></workbook>',"xl/worksheets/sheet1.xml":'<?xml version="1.0" encoding="UTF-8" standalone="yes"?><worksheet xmlns="http://schemas.openxmlformats.org/spreadsheetml/2006/main" xmlns:r="http://schemas.openxmlformats.org/officeDocument/2006/relationships" xmlns:mc="http://schemas.openxmlformats.org/markup-compatibility/2006" mc:Ignorable="x14ac" xmlns:x14ac="http://schemas.microsoft.com/office/spreadsheetml/2009/9/ac"><sheetData/><mergeCells count="0"/></worksheet>',"xl/styles.xml":'<?xml version="1.0" encoding="UTF-8"?><styleSheet xmlns="http://schemas.openxmlformats.org/spreadsheetml/2006/main" xmlns:mc="http://schemas.openxmlformats.org/markup-compatibility/2006" mc:Ignorable="x14ac" xmlns:x14ac="http://schemas.microsoft.com/office/spreadsheetml/2009/9/ac"><numFmts count="6"><numFmt numFmtId="164" formatCode="#,##0.00_- [$$-45C]"/><numFmt numFmtId="165" formatCode="&quot;£&quot;#,##0.00"/><numFmt numFmtId="166" formatCode="[$€-2] #,##0.00"/><numFmt numFmtId="167" formatCode="0.0%"/><numFmt numFmtId="168" formatCode="#,##0;(#,##0)"/><numFmt numFmtId="169" formatCode="#,##0.00;(#,##0.00)"/></numFmts><fonts count="5" x14ac:knownFonts="1"><font><sz val="11" /><name val="Calibri" /></font><font><sz val="11" /><name val="Calibri" /><color rgb="FFFFFFFF" /></font><font><sz val="11" /><name val="Calibri" /><b /></font><font><sz val="11" /><name val="Calibri" /><i /></font><font><sz val="11" /><name val="Calibri" /><u /></font></fonts><fills count="6"><fill><patternFill patternType="none" /></fill><fill><patternFill patternType="none" /></fill><fill><patternFill patternType="solid"><fgColor rgb="FFD9D9D9" /><bgColor indexed="64" /></patternFill></fill><fill><patternFill patternType="solid"><fgColor rgb="FFD99795" /><bgColor indexed="64" /></patternFill></fill><fill><patternFill patternType="solid"><fgColor rgb="ffc6efce" /><bgColor indexed="64" /></patternFill></fill><fill><patternFill patternType="solid"><fgColor rgb="ffc6cfef" /><bgColor indexed="64" /></patternFill></fill></fills><borders count="2"><border><left /><right /><top /><bottom /><diagonal /></border><border diagonalUp="false" diagonalDown="false"><left style="thin"><color auto="1" /></left><right style="thin"><color auto="1" /></right><top style="thin"><color auto="1" /></top><bottom style="thin"><color auto="1" /></bottom><diagonal /></border></borders><cellStyleXfs count="1"><xf numFmtId="0" fontId="0" fillId="0" borderId="0" /></cellStyleXfs><cellXfs count="68"><xf numFmtId="0" fontId="0" fillId="0" borderId="0" applyFont="1" applyFill="1" applyBorder="1"/><xf numFmtId="0" fontId="1" fillId="0" borderId="0" applyFont="1" applyFill="1" applyBorder="1"/><xf numFmtId="0" fontId="2" fillId="0" borderId="0" applyFont="1" applyFill="1" applyBorder="1"/><xf numFmtId="0" fontId="3" fillId="0" borderId="0" applyFont="1" applyFill="1" applyBorder="1"/><xf numFmtId="0" fontId="4" fillId="0" borderId="0" applyFont="1" applyFill="1" applyBorder="1"/><xf numFmtId="0" fontId="0" fillId="2" borderId="0" applyFont="1" applyFill="1" applyBorder="1"/><xf numFmtId="0" fontId="1" fillId="2" borderId="0" applyFont="1" applyFill="1" applyBorder="1"/><xf numFmtId="0" fontId="2" fillId="2" borderId="0" applyFont="1" applyFill="1" applyBorder="1"/><xf numFmtId="0" fontId="3" fillId="2" borderId="0" applyFont="1" applyFill="1" applyBorder="1"/><xf numFmtId="0" fontId="4" fillId="2" borderId="0" applyFont="1" applyFill="1" applyBorder="1"/><xf numFmtId="0" fontId="0" fillId="3" borderId="0" applyFont="1" applyFill="1" applyBorder="1"/><xf numFmtId="0" fontId="1" fillId="3" borderId="0" applyFont="1" applyFill="1" applyBorder="1"/><xf numFmtId="0" fontId="2" fillId="3" borderId="0" applyFont="1" applyFill="1" applyBorder="1"/><xf numFmtId="0" fontId="3" fillId="3" borderId="0" applyFont="1" applyFill="1" applyBorder="1"/><xf numFmtId="0" fontId="4" fillId="3" borderId="0" applyFont="1" applyFill="1" applyBorder="1"/><xf numFmtId="0" fontId="0" fillId="4" borderId="0" applyFont="1" applyFill="1" applyBorder="1"/><xf numFmtId="0" fontId="1" fillId="4" borderId="0" applyFont="1" applyFill="1" applyBorder="1"/><xf numFmtId="0" fontId="2" fillId="4" borderId="0" applyFont="1" applyFill="1" applyBorder="1"/><xf numFmtId="0" fontId="3" fillId="4" borderId="0" applyFont="1" applyFill="1" applyBorder="1"/><xf numFmtId="0" fontId="4" fillId="4" borderId="0" applyFont="1" applyFill="1" applyBorder="1"/><xf numFmtId="0" fontId="0" fillId="5" borderId="0" applyFont="1" applyFill="1" applyBorder="1"/><xf numFmtId="0" fontId="1" fillId="5" borderId="0" applyFont="1" applyFill="1" applyBorder="1"/><xf numFmtId="0" fontId="2" fillId="5" borderId="0" applyFont="1" applyFill="1" applyBorder="1"/><xf numFmtId="0" fontId="3" fillId="5" borderId="0" applyFont="1" applyFill="1" applyBorder="1"/><xf numFmtId="0" fontId="4" fillId="5" borderId="0" applyFont="1" applyFill="1" applyBorder="1"/><xf numFmtId="0" fontId="0" fillId="0" borderId="1" applyFont="1" applyFill="1" applyBorder="1"/><xf numFmtId="0" fontId="1" fillId="0" borderId="1" applyFont="1" applyFill="1" applyBorder="1"/><xf numFmtId="0" fontId="2" fillId="0" borderId="1" applyFont="1" applyFill="1" applyBorder="1"/><xf numFmtId="0" fontId="3" fillId="0" borderId="1" applyFont="1" applyFill="1" applyBorder="1"/><xf numFmtId="0" fontId="4" fillId="0" borderId="1" applyFont="1" applyFill="1" applyBorder="1"/><xf numFmtId="0" fontId="0" fillId="2" borderId="1" applyFont="1" applyFill="1" applyBorder="1"/><xf numFmtId="0" fontId="1" fillId="2" borderId="1" applyFont="1" applyFill="1" applyBorder="1"/><xf numFmtId="0" fontId="2" fillId="2" borderId="1" applyFont="1" applyFill="1" applyBorder="1"/><xf numFmtId="0" fontId="3" fillId="2" borderId="1" applyFont="1" applyFill="1" applyBorder="1"/><xf numFmtId="0" fontId="4" fillId="2" borderId="1" applyFont="1" applyFill="1" applyBorder="1"/><xf numFmtId="0" fontId="0" fillId="3" borderId="1" applyFont="1" applyFill="1" applyBorder="1"/><xf numFmtId="0" fontId="1" fillId="3" borderId="1" applyFont="1" applyFill="1" applyBorder="1"/><xf numFmtId="0" fontId="2" fillId="3" borderId="1" applyFont="1" applyFill="1" applyBorder="1"/><xf numFmtId="0" fontId="3" fillId="3" borderId="1" applyFont="1" applyFill="1" applyBorder="1"/><xf numFmtId="0" fontId="4" fillId="3" borderId="1" applyFont="1" applyFill="1" applyBorder="1"/><xf numFmtId="0" fontId="0" fillId="4" borderId="1" applyFont="1" applyFill="1" applyBorder="1"/><xf numFmtId="0" fontId="1" fillId="4" borderId="1" applyFont="1" applyFill="1" applyBorder="1"/><xf numFmtId="0" fontId="2" fillId="4" borderId="1" applyFont="1" applyFill="1" applyBorder="1"/><xf numFmtId="0" fontId="3" fillId="4" borderId="1" applyFont="1" applyFill="1" applyBorder="1"/><xf numFmtId="0" fontId="4" fillId="4" borderId="1" applyFont="1" applyFill="1" applyBorder="1"/><xf numFmtId="0" fontId="0" fillId="5" borderId="1" applyFont="1" applyFill="1" applyBorder="1"/><xf numFmtId="0" fontId="1" fillId="5" borderId="1" applyFont="1" applyFill="1" applyBorder="1"/><xf numFmtId="0" fontId="2" fillId="5" borderId="1" applyFont="1" applyFill="1" applyBorder="1"/><xf numFmtId="0" fontId="3" fillId="5" borderId="1" applyFont="1" applyFill="1" applyBorder="1"/><xf numFmtId="0" fontId="4" fillId="5" borderId="1" applyFont="1" applyFill="1" applyBorder="1"/><xf numFmtId="0" fontId="0" fillId="0" borderId="0" applyFont="1" applyFill="1" applyBorder="1" xfId="0" applyAlignment="1"><alignment horizontal="left"/></xf><xf numFmtId="0" fontId="0" fillId="0" borderId="0" applyFont="1" applyFill="1" applyBorder="1" xfId="0" applyAlignment="1"><alignment horizontal="center"/></xf><xf numFmtId="0" fontId="0" fillId="0" borderId="0" applyFont="1" applyFill="1" applyBorder="1" xfId="0" applyAlignment="1"><alignment horizontal="right"/></xf><xf numFmtId="0" fontId="0" fillId="0" borderId="0" applyFont="1" applyFill="1" applyBorder="1" xfId="0" applyAlignment="1"><alignment horizontal="fill"/></xf><xf numFmtId="0" fontId="0" fillId="0" borderId="0" applyFont="1" applyFill="1" applyBorder="1" xfId="0" applyAlignment="1"><alignment textRotation="90"/></xf><xf numFmtId="0" fontId="0" fillId="0" borderId="0" applyFont="1" applyFill="1" applyBorder="1" xfId="0" applyAlignment="1"><alignment wrapText="1"/></xf><xf numFmtId="9"   fontId="0" fillId="0" borderId="0" applyFont="1" applyFill="1" applyBorder="1" xfId="0" applyNumberFormat="1"/><xf numFmtId="164" fontId="0" fillId="0" borderId="0" applyFont="1" applyFill="1" applyBorder="1" xfId="0" applyNumberFormat="1"/><xf numFmtId="165" fontId="0" fillId="0" borderId="0" applyFont="1" applyFill="1" applyBorder="1" xfId="0" applyNumberFormat="1"/><xf numFmtId="166" fontId="0" fillId="0" borderId="0" applyFont="1" applyFill="1" applyBorder="1" xfId="0" applyNumberFormat="1"/><xf numFmtId="167" fontId="0" fillId="0" borderId="0" applyFont="1" applyFill="1" applyBorder="1" xfId="0" applyNumberFormat="1"/><xf numFmtId="168" fontId="0" fillId="0" borderId="0" applyFont="1" applyFill="1" applyBorder="1" xfId="0" applyNumberFormat="1"/><xf numFmtId="169" fontId="0" fillId="0" borderId="0" applyFont="1" applyFill="1" applyBorder="1" xfId="0" applyNumberFormat="1"/><xf numFmtId="3" fontId="0" fillId="0" borderId="0" applyFont="1" applyFill="1" applyBorder="1" xfId="0" applyNumberFormat="1"/><xf numFmtId="4" fontId="0" fillId="0" borderId="0" applyFont="1" applyFill="1" applyBorder="1" xfId="0" applyNumberFormat="1"/><xf numFmtId="1" fontId="0" fillId="0" borderId="0" applyFont="1" applyFill="1" applyBorder="1" xfId="0" applyNumberFormat="1"/><xf numFmtId="2" fontId="0" fillId="0" borderId="0" applyFont="1" applyFill="1" applyBorder="1" xfId="0" applyNumberFormat="1"/><xf numFmtId="14" fontId="0" fillId="0" borderId="0" applyFont="1" applyFill="1" applyBorder="1" xfId="0" applyNumberFormat="1"/></cellXfs><cellStyles count="1"><cellStyle name="Normal" xfId="0" builtinId="0" /></cellStyles><dxfs count="0" /><tableStyles count="0" defaultTableStyle="TableStyleMedium9" defaultPivotStyle="PivotStyleMedium4" /></styleSheet>'},R=[{match:/^\-?\d+\.\d%$/,style:60,fmt:function(t){return t/100}},{match:/^\-?\d+\.?\d*%$/,style:56,fmt:function(t){return t/100}},{match:/^\-?\$[\d,]+.?\d*$/,style:57},{match:/^\-?£[\d,]+.?\d*$/,style:58},{match:/^\-?€[\d,]+.?\d*$/,style:59},{match:/^\-?\d+$/,style:65},{match:/^\-?\d+\.\d{2}$/,style:66},{match:/^\([\d,]+\)$/,style:61,fmt:function(t){return-1*t.replace(/[\(\)]/g,"")}},{match:/^\([\d,]+\.\d{2}\)$/,style:62,fmt:function(t){return-1*t.replace(/[\(\)]/g,"")}},{match:/^\-?[\d,]+$/,style:63},{match:/^\-?[\d,]+\.\d{2}$/,style:64},{match:/^[\d]{4}\-[01][\d]\-[0123][\d]$/,style:67,fmt:function(t){return Math.round(25569+Date.parse(t)/864e5)}}];return n.ext.buttons.copyHtml5={className:"buttons-copy buttons-html5",text:function(t){return t.i18n("buttons.copy","Copy")},action:function(t,e,o,l){this.processing(!0);var n=this,r=u(e,l),a=e.buttons.exportInfo(l),d=c(l),p=r.str,i=C("<div/>").css({height:1,width:1,overflow:"hidden",position:"fixed",top:0,left:0}),d=(a.title&&(p=a.title+d+d+p),a.messageTop&&(p=a.messageTop+d+d+p),a.messageBottom&&(p=p+d+d+a.messageBottom),l.customize&&(p=l.customize(p,l,e)),C("<textarea readonly/>").val(p).appendTo(i));if(y.queryCommandSupported("copy")){i.appendTo(e.table().container()),d[0].focus(),d[0].select();try{var s=y.execCommand("copy");if(i.remove(),s)return e.buttons.info(e.i18n("buttons.copyTitle","Copy to clipboard"),e.i18n("buttons.copySuccess",{1:"Copied one row to clipboard",_:"Copied %d rows to clipboard"},r.rows),2e3),void this.processing(!1)}catch(t){}}function f(){m.off("click.buttons-copy"),C(y).off(".buttons-copy"),e.buttons.info(!1)}var a=C("<span>"+e.i18n("buttons.copyKeys","Press <i>ctrl</i> or <i>⌘</i> + <i>C</i> to copy the table data<br>to your system clipboard.<br><br>To cancel, click this message or press escape.")+"</span>").append(i),m=(e.buttons.info(e.i18n("buttons.copyTitle","Copy to clipboard"),a,0),d[0].focus(),d[0].select(),C(a).closest(".dt-button-info"));m.on("click.buttons-copy",f),C(y).on("keydown.buttons-copy",function(t){27===t.keyCode&&(f(),n.processing(!1))}).on("copy.buttons-copy cut.buttons-copy",function(){f(),n.processing(!1)})},exportOptions:{},fieldSeparator:"\t",fieldBoundary:"",header:!0,footer:!1,title:"*",messageTop:"*",messageBottom:"*"},n.ext.buttons.csvHtml5={bom:!1,className:"buttons-csv buttons-html5",available:function(){return T.FileReader!==k&&T.Blob},text:function(t){return t.i18n("buttons.csv","CSV")},action:function(t,e,o,l){this.processing(!0);var n=u(e,l).str,r=e.buttons.exportInfo(l),a=l.charset;l.customize&&(n=l.customize(n,l,e)),a=!1!==a?(a=a||y.characterSet||y.charset)&&";charset="+a:"",l.bom&&(n=String.fromCharCode(65279)+n),O(new Blob([n],{type:"text/csv"+a}),r.filename,!0),this.processing(!1)},filename:"*",extension:".csv",exportOptions:{},fieldSeparator:",",fieldBoundary:'"',escapeChar:'"',charset:null,header:!0,footer:!1},n.ext.buttons.excelHtml5={className:"buttons-excel buttons-html5",available:function(){return T.FileReader!==k&&S()!==k&&!f()&&A},text:function(t){return t.i18n("buttons.excel","Excel")},action:function(t,e,o,s){this.processing(!0);function l(t){return t=_[t],C.parseXML(t)}function n(t){m=E(u,"row",{attr:{r:f=y+1}});for(var e=0,o=t.length;e<o;e++){var l=z(e)+""+f,n=null;if(null===t[e]||t[e]===k||""===t[e]){if(!0!==s.createEmptyCells)continue;t[e]=""}var r=t[e];t[e]="function"==typeof t[e].trim?t[e].trim():t[e];for(var a=0,d=R.length;a<d;a++){var p=R[a];if(t[e].match&&!t[e].match(/^0\d+/)&&t[e].match(p.match)){var i=t[e].replace(/[^\d\.\-]/g,"");p.fmt&&(i=p.fmt(i)),n=E(u,"c",{attr:{r:l,s:p.style},children:[E(u,"v",{text:i})]});break}}n=n||("number"==typeof t[e]||t[e].match&&t[e].match(/^-?\d+(\.\d+)?([eE]\-?\d+)?$/)&&!t[e].match(/^0\d+/)?E(u,"c",{attr:{t:"n",r:l},children:[E(u,"v",{text:t[e]})]}):(r=r.replace?r.replace(/[\x00-\x09\x0B\x0C\x0E-\x1F\x7F-\x9F]/g,""):r,E(u,"c",{attr:{t:"inlineStr",r:l},children:{row:E(u,"is",{children:{row:E(u,"t",{text:r,attr:{"xml:space":"preserve"}})}})}}))),m.appendChild(n)}c.appendChild(m),y++}function r(t,e){var o=C("mergeCells",u);o[0].appendChild(E(u,"mergeCell",{attr:{ref:"A"+t+":"+z(e)+t}})),o.attr("count",parseFloat(o.attr("count"))+1),C("row:eq("+(t-1)+") c",u).attr("s","51")}var a,f,m,d=this,y=0,u=l("xl/worksheets/sheet1.xml"),c=u.getElementsByTagName("sheetData")[0],p={_rels:{".rels":l("_rels/.rels")},xl:{_rels:{"workbook.xml.rels":l("xl/_rels/workbook.xml.rels")},"workbook.xml":l("xl/workbook.xml"),"styles.xml":l("xl/styles.xml"),worksheets:{"sheet1.xml":u}},"[Content_Types].xml":l("[Content_Types].xml")},i=e.buttons.exportData(s.exportOptions),I=(s.customizeData&&s.customizeData(i),e.buttons.exportInfo(s));I.title&&(n([I.title]),r(y,i.header.length-1)),I.messageTop&&(n([I.messageTop]),r(y,i.header.length-1)),s.header&&(n(i.header),C("row:last c",u).attr("s","2"));for(var F=y,x=0,h=i.body.length;x<h;x++)n(i.body[x]);a=y,s.footer&&i.footer&&(n(i.footer),C("row:last c",u).attr("s","2")),I.messageBottom&&(n([I.messageBottom]),r(y,i.header.length-1));var b=E(u,"cols");C("worksheet",u).prepend(b);for(var g=0,v=i.header.length;g<v;g++)b.appendChild(E(u,"col",{attr:{min:g+1,max:g+1,width:function(t,e){var o=t.header[e].length;t.footer&&t.footer[e].length>o&&(o=t.footer[e].length);for(var l=0,n=t.body.length;l<n;l++){var r,a=t.body[l][e];if(40<(o=o<(r=(-1!==(a=null!==a&&a!==k?a.toString():"").indexOf("\n")?((r=a.split("\n")).sort(function(t,e){return e.length-t.length}),r[0]):a).length)?r:o))return 54}return 6<(o*=1.35)?o:6}(i,g),customWidth:1}}));var w=p.xl["workbook.xml"];C("sheets sheet",w).attr("name",N(s)),s.autoFilter&&(C("mergeCells",u).before(E(u,"autoFilter",{attr:{ref:"A"+F+":"+z(i.header.length-1)+a}})),C("definedNames",w).append(E(w,"definedName",{attr:{name:"_xlnm._FilterDatabase",localSheetId:"0",hidden:1},text:N(s)+"!$A$"+F+":"+z(i.header.length-1)+a}))),s.customize&&s.customize(p,s,e),0===C("mergeCells",u).children().length&&C("mergeCells",u).remove();var w=new(S()),F={compression:"DEFLATE",type:"blob",mimeType:"application/vnd.openxmlformats-officedocument.spreadsheetml.sheet"},B=(!function s(f,t){D===k&&(D=-1===A.serializeToString((new T.DOMParser).parseFromString(_["xl/worksheets/sheet1.xml"],"text/xml")).indexOf("xmlns:r")),C.each(t,function(t,e){if(C.isPlainObject(e))s(f.folder(t),e);else{if(D){for(var o,l=e.childNodes[0],n=[],r=l.attributes.length-1;0<=r;r--){var a=l.attributes[r].nodeName,d=l.attributes[r].nodeValue;-1!==a.indexOf(":")&&(n.push({name:a,value:d}),l.removeAttribute(a))}for(r=0,o=n.length;r<o;r++){var p=e.createAttribute(n[r].name.replace(":","_dt_b_namespace_token_"));p.value=n[r].value,l.setAttributeNode(p)}}var i=A.serializeToString(e),i=(i=D?(i=(i=-1===i.indexOf("<?xml")?'<?xml version="1.0" encoding="UTF-8" standalone="yes"?>'+i:i).replace(/_dt_b_namespace_token_/g,":")).replace(/xmlns:NS[\d]+="" NS[\d]+:/g,""):i).replace(/<([^<>]*?) xmlns=""([^<>]*?)>/g,"<$1 $2>");f.file(t,i)}})}(w,p),I.filename);175<B&&(B=B.substr(0,175)),w.generateAsync?w.generateAsync(F).then(function(t){O(t,B),d.processing(!1)}):(O(w.generate(F),B),this.processing(!1))},filename:"*",extension:".xlsx",exportOptions:{},header:!0,footer:!1,title:"*",messageTop:"*",messageBottom:"*",createEmptyCells:!1,autoFilter:!1,sheetName:""},n.ext.buttons.pdfHtml5={className:"buttons-pdf buttons-html5",available:function(){return T.FileReader!==k&&s()},text:function(t){return t.i18n("buttons.pdf","PDF")},action:function(t,e,o,l){this.processing(!0);var n=e.buttons.exportData(l.exportOptions),r=e.buttons.exportInfo(l),a=[];l.header&&a.push(C.map(n.header,function(t){return{text:"string"==typeof t?t:t+"",style:"tableHeader"}}));for(var d=0,p=n.body.length;d<p;d++)a.push(C.map(n.body[d],function(t){return{text:"string"==typeof(t=null!==t&&t!==k?t:"")?t:t+"",style:d%2?"tableBodyEven":"tableBodyOdd"}}));l.footer&&n.footer&&a.push(C.map(n.footer,function(t){return{text:"string"==typeof t?t:t+"",style:"tableFooter"}}));var i={pageSize:l.pageSize,pageOrientation:l.orientation,content:[{table:{headerRows:1,body:a},layout:"noBorders"}],styles:{tableHeader:{bold:!0,fontSize:11,color:"white",fillColor:"#2d4154",alignment:"center"},tableBodyEven:{},tableBodyOdd:{fillColor:"#f3f3f3"},tableFooter:{bold:!0,fontSize:11,color:"white",fillColor:"#2d4154"},title:{alignment:"center",fontSize:15},message:{}},defaultStyle:{fontSize:10}},e=(r.messageTop&&i.content.unshift({text:r.messageTop,style:"message",margin:[0,0,0,12]}),r.messageBottom&&i.content.push({text:r.messageBottom,style:"message",margin:[0,0,0,12]}),r.title&&i.content.unshift({text:r.title,style:"title",margin:[0,0,0,12]}),l.customize&&l.customize(i,l,e),s().createPdf(i));"open"!==l.download||f()?e.download(r.filename):e.open(),this.processing(!1)},title:"*",filename:"*",extension:".pdf",exportOptions:{},orientation:"portrait",pageSize:"A4",header:!0,footer:!1,messageTop:"*",messageBottom:"*",customize:null,download:"download"},n});
```
