## supplement_file1 for "GENETH’OFF: a flexible workflow for genome-wide profiling of CRISPR/Cas off-targets": buttons.print.min.js

```
/*!
 * Print button for Buttons and DataTables.
 * © SpryMedia Ltd - datatables.net/license
 */
!function(n){var o,r;"function"==typeof define&&define.amd?define(["jquery","datatables.net","datatables.net-buttons"],function(t){return n(t,window,document)}):"object"==typeof exports?(o=require("jquery"),r=function(t,e){e.fn.dataTable||require("datatables.net")(t,e),e.fn.dataTable.Buttons||require("datatables.net-buttons")(t,e)},"undefined"==typeof window?module.exports=function(t,e){return t=t||window,e=e||o(t),r(t,e),n(e,t,t.document)}:(r(window,o),module.exports=n(o,window,window.document))):n(jQuery,window,document)}(function(m,b,t,p){"use strict";function h(t){return n.href=t,-1===(t=n.host).indexOf("/")&&0!==n.pathname.indexOf("/")&&(t+="/"),n.protocol+"//"+t+n.pathname+n.search}var e=m.fn.dataTable,n=t.createElement("a");return e.ext.buttons.print={className:"buttons-print",text:function(t){return t.i18n("buttons.print","Print")},action:function(t,e,n,o){function r(t,e){for(var n="<tr>",o=0,r=t.length;o<r;o++){var i=null===t[o]||t[o]===p?"":t[o];n+="<"+e+" "+(s[o]?'class="'+s[o]+'"':"")+">"+i+"</"+e+">"}return n+"</tr>"}var i=e.buttons.exportData(m.extend({decodeEntities:!1},o.exportOptions)),a=e.buttons.exportInfo(o),s=e.columns(o.exportOptions.columns).flatten().map(function(t){return e.settings()[0].aoColumns[e.column(t).index()].sClass}).toArray(),u='<table class="'+e.table().node().className+'">';o.header&&(u+="<thead>"+r(i.header,"th")+"</thead>"),u+="<tbody>";for(var d=0,c=i.body.length;d<c;d++)u+=r(i.body[d],"td");u+="</tbody>",o.footer&&i.footer&&(u+="<tfoot>"+r(i.footer,"th")+"</tfoot>"),u+="</table>";var l=b.open("","");if(l){l.document.close();var f="<title>"+a.title+"</title>";m("style, link").each(function(){f+=function(t){t=m(t).clone()[0];return"link"===t.nodeName.toLowerCase()&&(t.href=h(t.href)),t.outerHTML}(this)});try{l.document.head.innerHTML=f}catch(t){m(l.document.head).html(f)}l.document.body.innerHTML="<h1>"+a.title+"</h1><div>"+(a.messageTop||"")+"</div>"+u+"<div>"+(a.messageBottom||"")+"</div>",m(l.document.body).addClass("dt-print-view"),m("img",l.document.body).each(function(t,e){e.setAttribute("src",h(e.getAttribute("src")))}),o.customize&&o.customize(l,o,e);a=function(){o.autoPrint&&(l.print(),l.close())};navigator.userAgent.match(/Trident\/\d.\d/)?a():l.setTimeout(a,1e3)}else e.buttons.info(e.i18n("buttons.printErrorTitle","Unable to open print view"),e.i18n("buttons.printErrorMsg","Please allow popups in your browser for this site to be able to view the print view."),5e3)},title:"*",messageTop:"*",messageBottom:"*",exportOptions:{},header:!0,footer:!1,autoPrint:!0,customize:null},e});
```
