## supplement_file1 for "GENETH’OFF: a flexible workflow for genome-wide profiling of CRISPR/Cas off-targets": dataTables.buttons.min.js

```
/*! Buttons for DataTables 2.4.2
 * © SpryMedia Ltd - datatables.net/license
 */
!function(e){var o,i;"function"==typeof define&&define.amd?define(["jquery","datatables.net"],function(t){return e(t,window,document)}):"object"==typeof exports?(o=require("jquery"),i=function(t,n){n.fn.dataTable||require("datatables.net")(t,n)},"undefined"==typeof window?module.exports=function(t,n){return t=t||window,n=n||o(t),i(t,n),e(n,t,t.document)}:(i(window,o),module.exports=e(o,window,window.document))):e(jQuery,window,document)}(function(y,v,x,w){"use strict";var e=y.fn.dataTable,o=0,C=0,_=e.ext.buttons,i=null;function A(t,n,e){y.fn.animate?t.stop().fadeIn(n,e):(t.css("display","block"),e&&e.call(t))}function k(t,n,e){y.fn.animate?t.stop().fadeOut(n,e):(t.css("display","none"),e&&e.call(t))}function B(n,t){if(!(this instanceof B))return function(t){return new B(t,n).container()};!0===(t=void 0===t?{}:t)&&(t={}),Array.isArray(t)&&(t={buttons:t}),this.c=y.extend(!0,{},B.defaults,t),t.buttons&&(this.c.buttons=t.buttons),this.s={dt:new e.Api(n),buttons:[],listenKeys:"",namespace:"dtb"+o++},this.dom={container:y("<"+this.c.dom.container.tag+"/>").addClass(this.c.dom.container.className)},this._constructor()}y.extend(B.prototype,{action:function(t,n){t=this._nodeToButton(t);return n===w?t.conf.action:(t.conf.action=n,this)},active:function(t,n){var t=this._nodeToButton(t),e=this.c.dom.button.active,o=y(t.node);return t.inCollection&&this.c.dom.collection.button&&this.c.dom.collection.button.active!==w&&(e=this.c.dom.collection.button.active),n===w?o.hasClass(e):(o.toggleClass(e,n===w||n),this)},add:function(t,n,e){var o=this.s.buttons;if("string"==typeof n){for(var i=n.split("-"),s=this.s,a=0,r=i.length-1;a<r;a++)s=s.buttons[+i[a]];o=s.buttons,n=+i[i.length-1]}return this._expandButton(o,t,t!==w?t.split:w,(t===w||t.split===w||0===t.split.length)&&s!==w,!1,n),e!==w&&!0!==e||this._draw(),this},collectionRebuild:function(t,n){var e=this._nodeToButton(t);if(n!==w){for(var o=e.buttons.length-1;0<=o;o--)this.remove(e.buttons[o].node);for(e.conf.prefixButtons&&n.unshift.apply(n,e.conf.prefixButtons),e.conf.postfixButtons&&n.push.apply(n,e.conf.postfixButtons),o=0;o<n.length;o++){var i=n[o];this._expandButton(e.buttons,i,i!==w&&i.config!==w&&i.config.split!==w,!0,i.parentConf!==w&&i.parentConf.split!==w,null,i.parentConf)}}this._draw(e.collection,e.buttons)},container:function(){return this.dom.container},disable:function(t){t=this._nodeToButton(t);return y(t.node).addClass(this.c.dom.button.disabled).prop("disabled",!0),this},destroy:function(){y("body").off("keyup."+this.s.namespace);for(var t=this.s.buttons.slice(),n=0,e=t.length;n<e;n++)this.remove(t[n].node);this.dom.container.remove();var o=this.s.dt.settings()[0];for(n=0,e=o.length;n<e;n++)if(o.inst===this){o.splice(n,1);break}return this},enable:function(t,n){return!1===n?this.disable(t):(n=this._nodeToButton(t),y(n.node).removeClass(this.c.dom.button.disabled).prop("disabled",!1),this)},index:function(t,n,e){n||(n="",e=this.s.buttons);for(var o=0,i=e.length;o<i;o++){var s=e[o].buttons;if(e[o].node===t)return n+o;if(s&&s.length){s=this.index(t,o+"-",s);if(null!==s)return s}}return null},name:function(){return this.c.name},node:function(t){return t?(t=this._nodeToButton(t),y(t.node)):this.dom.container},processing:function(t,n){var e=this.s.dt,o=this._nodeToButton(t);return n===w?y(o.node).hasClass("processing"):(y(o.node).toggleClass("processing",n),y(e.table().node()).triggerHandler("buttons-processing.dt",[n,e.button(t),e,y(t),o.conf]),this)},remove:function(t){var n=this._nodeToButton(t),e=this._nodeToHost(t),o=this.s.dt;if(n.buttons.length)for(var i=n.buttons.length-1;0<=i;i--)this.remove(n.buttons[i].node);n.conf.destroying=!0,n.conf.destroy&&n.conf.destroy.call(o.button(t),o,y(t),n.conf),this._removeKey(n.conf),y(n.node).remove();o=y.inArray(n,e);return e.splice(o,1),this},text:function(t,n){function e(t){return"function"==typeof t?t(i,s,o.conf):t}var o=this._nodeToButton(t),t=o.textNode,i=this.s.dt,s=y(o.node);return n===w?e(o.conf.text):(o.conf.text=n,t.html(e(n)),this)},_constructor:function(){var e=this,t=this.s.dt,o=t.settings()[0],n=this.c.buttons;o._buttons||(o._buttons=[]),o._buttons.push({inst:this,name:this.c.name});for(var i=0,s=n.length;i<s;i++)this.add(n[i]);t.on("destroy",function(t,n){n===o&&e.destroy()}),y("body").on("keyup."+this.s.namespace,function(t){var n;x.activeElement&&x.activeElement!==x.body||(n=String.fromCharCode(t.keyCode).toLowerCase(),-1!==e.s.listenKeys.toLowerCase().indexOf(n)&&e._keypress(n,t))})},_addKey:function(t){t.key&&(this.s.listenKeys+=(y.isPlainObject(t.key)?t.key:t).key)},_draw:function(t,n){t||(t=this.dom.container,n=this.s.buttons),t.children().detach();for(var e=0,o=n.length;e<o;e++)t.append(n[e].inserter),t.append(" "),n[e].buttons&&n[e].buttons.length&&this._draw(n[e].collection,n[e].buttons)},_expandButton:function(t,n,e,o,i,s,a){for(var r,l=this.s.dt,c=this.c.dom.collection,u=Array.isArray(n)?n:[n],d=0,f=(u=n===w?Array.isArray(e)?e:[e]:u).length;d<f;d++){var p=this._resolveExtends(u[d]);if(p)if(r=!(!p.config||!p.config.split),Array.isArray(p))this._expandButton(t,p,h!==w&&h.conf!==w?h.conf.split:w,o,a!==w&&a.split!==w,s,a);else{var h=this._buildButton(p,o,p.split!==w||p.config!==w&&p.config.split!==w,i);if(h){if(s!==w&&null!==s?(t.splice(s,0,h),s++):t.push(h),h.conf.buttons&&(h.collection=y("<"+c.container.content.tag+"/>"),h.conf._collection=h.collection,y(h.node).append(c.action.dropHtml),this._expandButton(h.buttons,h.conf.buttons,h.conf.split,!r,r,s,h.conf)),h.conf.split){h.collection=y("<"+c.container.tag+"/>"),h.conf._collection=h.collection;for(var b=0;b<h.conf.split.length;b++){var g=h.conf.split[b];"object"==typeof g&&(g.parent=a,g.collectionLayout===w&&(g.collectionLayout=h.conf.collectionLayout),g.dropup===w&&(g.dropup=h.conf.dropup),g.fade===w)&&(g.fade=h.conf.fade)}this._expandButton(h.buttons,h.conf.buttons,h.conf.split,!r,r,s,h.conf)}h.conf.parent=a,p.init&&p.init.call(l.button(h.node),l,y(h.node),p)}}}},_buildButton:function(n,t,e,o){function i(t){return"function"==typeof t?t(u,l,n):t}var s,a,r,l,c=this.c.dom,u=this.s.dt,d=y.extend(!0,{},c.button);if(t&&e&&c.collection.split?y.extend(!0,d,c.collection.split.action):o||t?y.extend(!0,d,c.collection.button):e&&y.extend(!0,d,c.split.button),n.spacer)return c=y("<"+d.spacer.tag+"/>").addClass("dt-button-spacer "+n.style+" "+d.spacer.className).html(i(n.text)),{conf:n,node:c,inserter:c,buttons:[],inCollection:t,isSplit:e,collection:null,textNode:c};if(n.available&&!n.available(u,n)&&!n.hasOwnProperty("html"))return!1;n.hasOwnProperty("html")?l=y(n.html):(a=function(t,n,e,o){o.action.call(n.button(e),t,n,e,o),y(n.table().node()).triggerHandler("buttons-action.dt",[n.button(e),n,e,o])},c=n.tag||d.tag,r=n.clickBlurs===w||n.clickBlurs,l=y("<"+c+"/>").addClass(d.className).attr("tabindex",this.s.dt.settings()[0].iTabIndex).attr("aria-controls",this.s.dt.table().node().id).on("click.dtb",function(t){t.preventDefault(),!l.hasClass(d.disabled)&&n.action&&a(t,u,l,n),r&&l.trigger("blur")}).on("keypress.dtb",function(t){13===t.keyCode&&(t.preventDefault(),!l.hasClass(d.disabled))&&n.action&&a(t,u,l,n)}),"a"===c.toLowerCase()&&l.attr("href","#"),"button"===c.toLowerCase()&&l.attr("type","button"),s=d.liner.tag?(c=y("<"+d.liner.tag+"/>").html(i(n.text)).addClass(d.liner.className),"a"===d.liner.tag.toLowerCase()&&c.attr("href","#"),l.append(c),c):(l.html(i(n.text)),l),!1===n.enabled&&l.addClass(d.disabled),n.className&&l.addClass(n.className),n.titleAttr&&l.attr("title",i(n.titleAttr)),n.attr&&l.attr(n.attr),n.namespace||(n.namespace=".dt-button-"+C++),n.config!==w&&n.config.split&&(n.split=n.config.split));var f,p,h,b,g,m,c=this.c.dom.buttonContainer,c=c&&c.tag?y("<"+c.tag+"/>").addClass(c.className).append(l):l;return this._addKey(n),this.c.buttonCreated&&(c=this.c.buttonCreated(n,c)),e&&(p=(f=t?y.extend(!0,this.c.dom.split,this.c.dom.collection.split):this.c.dom.split).wrapper,h=y("<"+p.tag+"/>").addClass(p.className).append(l),b=y.extend(n,{align:f.dropdown.align,attr:{"aria-haspopup":"dialog","aria-expanded":!1},className:f.dropdown.className,closeButton:!1,splitAlignClass:f.dropdown.splitAlignClass,text:f.dropdown.text}),this._addKey(b),g=function(t,n,e,o){_.split.action.call(n.button(h),t,n,e,o),y(n.table().node()).triggerHandler("buttons-action.dt",[n.button(e),n,e,o]),e.attr("aria-expanded",!0)},m=y('<button class="'+f.dropdown.className+' dt-button"></button>').html(f.dropdown.dropHtml).on("click.dtb",function(t){t.preventDefault(),t.stopPropagation(),m.hasClass(d.disabled)||g(t,u,m,b),r&&m.trigger("blur")}).on("keypress.dtb",function(t){13===t.keyCode&&(t.preventDefault(),m.hasClass(d.disabled)||g(t,u,m,b))}),0===n.split.length&&m.addClass("dtb-hide-drop"),h.append(m).attr(b.attr)),{conf:n,node:(e?h:l).get(0),inserter:e?h:c,buttons:[],inCollection:t,isSplit:e,inSplit:o,collection:null,textNode:s}},_nodeToButton:function(t,n){for(var e=0,o=(n=n||this.s.buttons).length;e<o;e++){if(n[e].node===t)return n[e];if(n[e].buttons.length){var i=this._nodeToButton(t,n[e].buttons);if(i)return i}}},_nodeToHost:function(t,n){for(var e=0,o=(n=n||this.s.buttons).length;e<o;e++){if(n[e].node===t)return n;if(n[e].buttons.length){var i=this._nodeToHost(t,n[e].buttons);if(i)return i}}},_keypress:function(s,a){var r;a._buttonsHandled||(r=function(t){for(var n,e,o=0,i=t.length;o<i;o++)n=t[o].conf,e=t[o].node,!n.key||n.key!==s&&(!y.isPlainObject(n.key)||n.key.key!==s||n.key.shiftKey&&!a.shiftKey||n.key.altKey&&!a.altKey||n.key.ctrlKey&&!a.ctrlKey||n.key.metaKey&&!a.metaKey)||(a._buttonsHandled=!0,y(e).click()),t[o].buttons.length&&r(t[o].buttons)})(this.s.buttons)},_removeKey:function(t){var n;t.key&&(t=(y.isPlainObject(t.key)?t.key:t).key,n=this.s.listenKeys.split(""),t=y.inArray(t,n),n.splice(t,1),this.s.listenKeys=n.join(""))},_resolveExtends:function(e){function t(t){for(var n=0;!y.isPlainObject(t)&&!Array.isArray(t);){if(t===w)return;if("function"==typeof t){if(!(t=t.call(i,s,e)))return!1}else if("string"==typeof t){if(!_[t])return{html:t};t=_[t]}if(30<++n)throw"Buttons: Too many iterations"}return Array.isArray(t)?t:y.extend({},t)}var n,o,i=this,s=this.s.dt;for(e=t(e);e&&e.extend;){if(!_[e.extend])throw"Cannot extend unknown button type: "+e.extend;var a=t(_[e.extend]);if(Array.isArray(a))return a;if(!a)return!1;var r=a.className;e.config!==w&&a.config!==w&&(e.config=y.extend({},a.config,e.config)),e=y.extend({},a,e),r&&e.className!==r&&(e.className=r+" "+e.className),e.extend=a.extend}var l=e.postfixButtons;if(l)for(e.buttons||(e.buttons=[]),n=0,o=l.length;n<o;n++)e.buttons.push(l[n]);var c=e.prefixButtons;if(c)for(e.buttons||(e.buttons=[]),n=0,o=c.length;n<o;n++)e.buttons.splice(n,0,c[n]);return e},_popover:function(o,t,n,e){function i(){h=!0,k(y(g),b.fade,function(){y(this).detach()}),y(f.buttons('[aria-haspopup="dialog"][aria-expanded="true"]').nodes()).attr("aria-expanded","false"),y("div.dt-button-background").off("click.dtb-collection"),B.background(!1,b.backgroundClassName,b.fade,m),y(v).off("resize.resize.dtb-collection"),y("body").off(".dtb-collection"),f.off("buttons-action.b-internal"),f.off("destroy")}var s,a,r,l,c,u,d,f=t,p=this.c,h=!1,b=y.extend({align:"button-left",autoClose:!1,background:!0,backgroundClassName:"dt-button-background",closeButton:!0,containerClassName:p.dom.collection.container.className,contentClassName:p.dom.collection.container.content.className,collectionLayout:"",collectionTitle:"",dropup:!1,fade:400,popoverTitle:"",rightAlignClassName:"dt-button-right",tag:p.dom.collection.container.tag},n),g=b.tag+"."+b.containerClassName.replace(/ /g,"."),m=t.node();!1===o?i():((p=y(f.buttons('[aria-haspopup="dialog"][aria-expanded="true"]').nodes())).length&&(m.closest(g).length&&(m=p.eq(0)),i()),n=y(".dt-button",o).length,p="",3===n?p="dtb-b3":2===n?p="dtb-b2":1===n&&(p="dtb-b1"),s=y("<"+b.tag+"/>").addClass(b.containerClassName).addClass(b.collectionLayout).addClass(b.splitAlignClass).addClass(p).css("display","none").attr({"aria-modal":!0,role:"dialog"}),o=y(o).addClass(b.contentClassName).attr("role","menu").appendTo(s),m.attr("aria-expanded","true"),m.parents("body")[0]!==x.body&&(m=x.body.lastChild),b.popoverTitle?s.prepend('<div class="dt-button-collection-title">'+b.popoverTitle+"</div>"):b.collectionTitle&&s.prepend('<div class="dt-button-collection-title">'+b.collectionTitle+"</div>"),b.closeButton&&s.prepend('<div class="dtb-popover-close">&times;</div>').addClass("dtb-collection-closeable"),A(s.insertAfter(m),b.fade),n=y(t.table().container()),d=s.css("position"),"container"!==b.span&&"dt-container"!==b.align||(m=m.parent(),s.css("width",n.width())),"absolute"===d?(p=y(m[0].offsetParent),t=m.position(),n=m.offset(),a=p.offset(),r=p.position(),l=v.getComputedStyle(p[0]),a.height=p.outerHeight(),a.width=p.width()+parseFloat(l.paddingLeft),a.right=a.left+a.width,a.bottom=a.top+a.height,p=t.top+m.outerHeight(),c=t.left,s.css({top:p,left:c}),l=v.getComputedStyle(s[0]),(u=s.offset()).height=s.outerHeight(),u.width=s.outerWidth(),u.right=u.left+u.width,u.bottom=u.top+u.height,u.marginTop=parseFloat(l.marginTop),u.marginBottom=parseFloat(l.marginBottom),b.dropup&&(p=t.top-u.height-u.marginTop-u.marginBottom),"button-right"!==b.align&&!s.hasClass(b.rightAlignClassName)||(c=t.left-u.width+m.outerWidth()),"dt-container"!==b.align&&"container"!==b.align||(c=c<t.left?-t.left:c)+u.width>a.width&&(c=a.width-u.width),r.left+c+u.width>y(v).width()&&(c=y(v).width()-u.width-r.left),n.left+c<0&&(c=-n.left),r.top+p+u.height>y(v).height()+y(v).scrollTop()&&(p=t.top-u.height-u.marginTop-u.marginBottom),r.top+p<y(v).scrollTop()&&(p=t.top+m.outerHeight()),s.css({top:p,left:c})):((d=function(){var t=y(v).height()/2,n=s.height()/2;s.css("marginTop",-1*(n=t<n?t:n))})(),y(v).on("resize.dtb-collection",function(){d()})),b.background&&B.background(!0,b.backgroundClassName,b.fade,b.backgroundHost||m),y("div.dt-button-background").on("click.dtb-collection",function(){}),b.autoClose&&setTimeout(function(){f.on("buttons-action.b-internal",function(t,n,e,o){o[0]!==m[0]&&i()})},0),y(s).trigger("buttons-popover.dt"),f.on("destroy",i),setTimeout(function(){h=!1,y("body").on("click.dtb-collection",function(t){var n,e;!h&&(n=y.fn.addBack?"addBack":"andSelf",e=y(t.target).parent()[0],!y(t.target).parents()[n]().filter(o).length&&!y(e).hasClass("dt-buttons")||y(t.target).hasClass("dt-button-background"))&&i()}).on("keyup.dtb-collection",function(t){27===t.keyCode&&i()}).on("keydown.dtb-collection",function(t){var n=y("a, button",o),e=x.activeElement;9===t.keyCode&&(-1===n.index(e)?(n.first().focus(),t.preventDefault()):t.shiftKey?e===n[0]&&(n.last().focus(),t.preventDefault()):e===n.last()[0]&&(n.first().focus(),t.preventDefault()))})},0))}}),B.background=function(t,n,e,o){e===w&&(e=400),o=o||x.body,t?A(y("<div/>").addClass(n).css("display","none").insertAfter(o),e):k(y("div."+n),e,function(){y(this).removeClass(n).remove()})},B.instanceSelector=function(t,i){var s,a,r;return t===w||null===t?y.map(i,function(t){return t.inst}):(s=[],a=y.map(i,function(t){return t.name}),(r=function(t){var n;if(Array.isArray(t))for(var e=0,o=t.length;e<o;e++)r(t[e]);else"string"==typeof t?-1!==t.indexOf(",")?r(t.split(",")):-1!==(n=y.inArray(t.trim(),a))&&s.push(i[n].inst):"number"==typeof t?s.push(i[t].inst):"object"==typeof t&&s.push(t)})(t),s)},B.buttonSelector=function(t,n){for(var c=[],u=function(t,n,e){for(var o,i,s=0,a=n.length;s<a;s++)(o=n[s])&&(t.push({node:o.node,name:o.conf.name,idx:i=e!==w?e+s:s+""}),o.buttons)&&u(t,o.buttons,i+"-")},d=function(t,n){var e=[],o=(u(e,n.s.buttons),y.map(e,function(t){return t.node}));if(Array.isArray(t)||t instanceof y)for(s=0,a=t.length;s<a;s++)d(t[s],n);else if(null===t||t===w||"*"===t)for(s=0,a=e.length;s<a;s++)c.push({inst:n,node:e[s].node});else if("number"==typeof t)n.s.buttons[t]&&c.push({inst:n,node:n.s.buttons[t].node});else if("string"==typeof t)if(-1!==t.indexOf(","))for(var i=t.split(","),s=0,a=i.length;s<a;s++)d(i[s].trim(),n);else if(t.match(/^\d+(\-\d+)*$/)){var r=y.map(e,function(t){return t.idx});c.push({inst:n,node:e[y.inArray(t,r)].node})}else if(-1!==t.indexOf(":name")){var l=t.replace(":name","");for(s=0,a=e.length;s<a;s++)e[s].name===l&&c.push({inst:n,node:e[s].node})}else y(o).filter(t).each(function(){c.push({inst:n,node:this})});else"object"==typeof t&&t.nodeName&&-1!==(r=y.inArray(t,o))&&c.push({inst:n,node:o[r]})},e=0,o=t.length;e<o;e++){var i=t[e];d(n,i)}return c},B.stripData=function(t,n){return t="string"==typeof t&&(t=(t=t.replace(/<script\b[^<]*(?:(?!<\/script>)<[^<]*)*<\/script>/gi,"")).replace(/<!\-\-.*?\-\->/g,""),n&&!n.stripHtml||(t=t.replace(/<[^>]*>/g,"")),n&&!n.trim||(t=t.replace(/^\s+|\s+$/g,"")),n&&!n.stripNewlines||(t=t.replace(/\n/g," ")),!n||n.decodeEntities)?i?i(t):(c.innerHTML=t,c.value):t},B.entityDecoder=function(t){i=t},B.defaults={buttons:["copy","excel","csv","pdf","print"],name:"main",tabIndex:0,dom:{container:{tag:"div",className:"dt-buttons"},collection:{action:{dropHtml:'<span class="dt-button-down-arrow">&#x25BC;</span>'},container:{className:"dt-button-collection",content:{className:"",tag:"div"},tag:"div"}},button:{tag:"button",className:"dt-button",active:"dt-button-active",disabled:"disabled",spacer:{className:"dt-button-spacer",tag:"span"},liner:{tag:"span",className:""}},split:{action:{className:"dt-button-split-drop-button dt-button",tag:"button"},dropdown:{align:"split-right",className:"dt-button-split-drop",dropHtml:'<span class="dt-button-down-arrow">&#x25BC;</span>',splitAlignClass:"dt-button-split-left",tag:"button"},wrapper:{className:"dt-button-split",tag:"div"}}}},y.extend(_,{collection:{text:function(t){return t.i18n("buttons.collection","Collection")},className:"buttons-collection",closeButton:!(B.version="2.4.2"),init:function(t,n,e){n.attr("aria-expanded",!1)},action:function(t,n,e,o){o._collection.parents("body").length?this.popover(!1,o):this.popover(o._collection,o),"keypress"===t.type&&y("a, button",o._collection).eq(0).focus()},attr:{"aria-haspopup":"dialog"}},split:{text:function(t){return t.i18n("buttons.split","Split")},className:"buttons-split",closeButton:!1,init:function(t,n,e){return n.attr("aria-expanded",!1)},action:function(t,n,e,o){this.popover(o._collection,o)},attr:{"aria-haspopup":"dialog"}},copy:function(t,n){if(_.copyHtml5)return"copyHtml5"},csv:function(t,n){if(_.csvHtml5&&_.csvHtml5.available(t,n))return"csvHtml5"},excel:function(t,n){if(_.excelHtml5&&_.excelHtml5.available(t,n))return"excelHtml5"},pdf:function(t,n){if(_.pdfHtml5&&_.pdfHtml5.available(t,n))return"pdfHtml5"},pageLength:function(t){var n=t.settings()[0].aLengthMenu,e=[],o=[];if(Array.isArray(n[0]))e=n[0],o=n[1];else for(var i=0;i<n.length;i++){var s=n[i];y.isPlainObject(s)?(e.push(s.value),o.push(s.label)):(e.push(s),o.push(s))}return{extend:"collection",text:function(t){return t.i18n("buttons.pageLength",{"-1":"Show all rows",_:"Show %d rows"},t.page.len())},className:"buttons-page-length",autoClose:!0,buttons:y.map(e,function(s,t){return{text:o[t],className:"button-page-length",action:function(t,n){n.page.len(s).draw()},init:function(t,n,e){function o(){i.active(t.page.len()===s)}var i=this;t.on("length.dt"+e.namespace,o),o()},destroy:function(t,n,e){t.off("length.dt"+e.namespace)}}}),init:function(t,n,e){var o=this;t.on("length.dt"+e.namespace,function(){o.text(e.text)})},destroy:function(t,n,e){t.off("length.dt"+e.namespace)}}},spacer:{style:"empty",spacer:!0,text:function(t){return t.i18n("buttons.spacer","")}}}),e.Api.register("buttons()",function(n,e){e===w&&(e=n,n=w),this.selector.buttonGroup=n;var t=this.iterator(!0,"table",function(t){if(t._buttons)return B.buttonSelector(B.instanceSelector(n,t._buttons),e)},!0);return t._groupSelector=n,t}),e.Api.register("button()",function(t,n){t=this.buttons(t,n);return 1<t.length&&t.splice(1,t.length),t}),e.Api.registerPlural("buttons().active()","button().active()",function(n){return n===w?this.map(function(t){return t.inst.active(t.node)}):this.each(function(t){t.inst.active(t.node,n)})}),e.Api.registerPlural("buttons().action()","button().action()",function(n){return n===w?this.map(function(t){return t.inst.action(t.node)}):this.each(function(t){t.inst.action(t.node,n)})}),e.Api.registerPlural("buttons().collectionRebuild()","button().collectionRebuild()",function(e){return this.each(function(t){for(var n=0;n<e.length;n++)"object"==typeof e[n]&&(e[n].parentConf=t);t.inst.collectionRebuild(t.node,e)})}),e.Api.register(["buttons().enable()","button().enable()"],function(n){return this.each(function(t){t.inst.enable(t.node,n)})}),e.Api.register(["buttons().disable()","button().disable()"],function(){return this.each(function(t){t.inst.disable(t.node)})}),e.Api.register("button().index()",function(){var n=null;return this.each(function(t){t=t.inst.index(t.node);null!==t&&(n=t)}),n}),e.Api.registerPlural("buttons().nodes()","button().node()",function(){var n=y();return y(this.each(function(t){n=n.add(t.inst.node(t.node))})),n}),e.Api.registerPlural("buttons().processing()","button().processing()",function(n){return n===w?this.map(function(t){return t.inst.processing(t.node)}):this.each(function(t){t.inst.processing(t.node,n)})}),e.Api.registerPlural("buttons().text()","button().text()",function(n){return n===w?this.map(function(t){return t.inst.text(t.node)}):this.each(function(t){t.inst.text(t.node,n)})}),e.Api.registerPlural("buttons().trigger()","button().trigger()",function(){return this.each(function(t){t.inst.node(t.node).trigger("click")})}),e.Api.register("button().popover()",function(n,e){return this.map(function(t){return t.inst._popover(n,this.button(this[0].node),e)})}),e.Api.register("buttons().containers()",function(){var i=y(),s=this._groupSelector;return this.iterator(!0,"table",function(t){if(t._buttons)for(var n=B.instanceSelector(s,t._buttons),e=0,o=n.length;e<o;e++)i=i.add(n[e].container())}),i}),e.Api.register("buttons().container()",function(){return this.containers().eq(0)}),e.Api.register("button().add()",function(t,n,e){var o=this.context;return o.length&&(o=B.instanceSelector(this._groupSelector,o[0]._buttons)).length&&o[0].add(n,t,e),this.button(this._groupSelector,t)}),e.Api.register("buttons().destroy()",function(){return this.pluck("inst").unique().each(function(t){t.destroy()}),this}),e.Api.registerPlural("buttons().remove()","buttons().remove()",function(){return this.each(function(t){t.inst.remove(t.node)}),this}),e.Api.register("buttons.info()",function(t,n,e){var o=this;return!1===t?(this.off("destroy.btn-info"),k(y("#datatables_buttons_info"),400,function(){y(this).remove()}),clearTimeout(s),s=null):(s&&clearTimeout(s),y("#datatables_buttons_info").length&&y("#datatables_buttons_info").remove(),t=t?"<h2>"+t+"</h2>":"",A(y('<div id="datatables_buttons_info" class="dt-button-info"/>').html(t).append(y("<div/>")["string"==typeof n?"html":"append"](n)).css("display","none").appendTo("body")),e!==w&&0!==e&&(s=setTimeout(function(){o.buttons.info(!1)},e)),this.on("destroy.btn-info",function(){o.buttons.info(!1)})),this}),e.Api.register("buttons.exportData()",function(t){if(this.context.length)return u(new e.Api(this.context[0]),t)}),e.Api.register("buttons.exportInfo()",function(t){return{filename:n(t=t||{}),title:r(t),messageTop:l(this,t.message||t.messageTop,"top"),messageBottom:l(this,t.messageBottom,"bottom")}});var s,n=function(t){var n;return(n="function"==typeof(n="*"===t.filename&&"*"!==t.title&&t.title!==w&&null!==t.title&&""!==t.title?t.title:t.filename)?n():n)===w||null===n?null:(n=(n=-1!==n.indexOf("*")?n.replace("*",y("head > title").text()).trim():n).replace(/[^a-zA-Z0-9_\u00A1-\uFFFF\.,\-_ !\(\)]/g,""))+(a(t.extension)||"")},a=function(t){return null===t||t===w?null:"function"==typeof t?t():t},r=function(t){t=a(t.title);return null===t?null:-1!==t.indexOf("*")?t.replace("*",y("head > title").text()||"Exported data"):t},l=function(t,n,e){n=a(n);return null===n?null:(t=y("caption",t.table().container()).eq(0),"*"===n?t.css("caption-side")!==e?null:t.length?t.text():"":n)},c=y("<textarea/>")[0],u=function(e,t){for(var o=y.extend(!0,{},{rows:null,columns:"",modifier:{search:"applied",order:"applied"},orthogonal:"display",stripHtml:!0,stripNewlines:!0,decodeEntities:!0,trim:!0,format:{header:function(t){return B.stripData(t,o)},footer:function(t){return B.stripData(t,o)},body:function(t){return B.stripData(t,o)}},customizeData:null},t),t=e.columns(o.columns).indexes().map(function(t){var n=e.column(t).header();return o.format.header(n.innerHTML,t,n)}).toArray(),n=e.table().footer()?e.columns(o.columns).indexes().map(function(t){var n=e.column(t).footer();return o.format.footer(n?n.innerHTML:"",t,n)}).toArray():null,i=y.extend({},o.modifier),i=(e.select&&"function"==typeof e.select.info&&i.selected===w&&e.rows(o.rows,y.extend({selected:!0},i)).any()&&y.extend(i,{selected:!0}),e.rows(o.rows,i).indexes().toArray()),i=e.cells(i,o.columns),s=i.render(o.orthogonal).toArray(),a=i.nodes().toArray(),r=t.length,l=[],c=0,u=0,d=0<r?s.length/r:0;u<d;u++){for(var f=[r],p=0;p<r;p++)f[p]=o.format.body(s[c],u,p,a[c]),c++;l[u]=f}i={header:t,footer:n,body:l};return o.customizeData&&o.customizeData(i),i};function t(t,n){t=new e.Api(t),n=n||t.init().buttons||e.defaults.buttons;return new B(t,n).container()}return y.fn.dataTable.Buttons=B,y.fn.DataTable.Buttons=B,y(x).on("init.dt plugin-init.dt",function(t,n){"dt"===t.namespace&&(t=n.oInit.buttons||e.defaults.buttons)&&!n._buttons&&new B(n,t).container()}),e.ext.feature.push({fnInit:t,cFeature:"B"}),e.ext.features&&e.ext.features.register("buttons",t),e});
```
