## supplement_file1 for "GENETH’OFF: a flexible workflow for genome-wide profiling of CRISPR/Cas off-targets": jszip.min.js

```
/*!

JSZip v3.10.1 - A JavaScript class for generating and reading zip files
<http://stuartk.com/jszip>

(c) 2009-2016 Stuart Knightley <stuart [at] stuartk.com>
Dual licenced under the MIT license or GPLv3. See https://raw.github.com/Stuk/jszip/main/LICENSE.markdown.

JSZip uses the library pako released under the MIT license :
https://github.com/nodeca/pako/blob/main/LICENSE
*/

!function(e){if("object"==typeof exports&&"undefined"!=typeof module)module.exports=e();else if("function"==typeof define&&define.amd)define([],e);else{("undefined"!=typeof window?window:"undefined"!=typeof global?global:"undefined"!=typeof self?self:this).JSZip=e()}}(function(){return function s(a,o,h){function u(r,e){if(!o[r]){if(!a[r]){var t="function"==typeof require&&require;if(!e&&t)return t(r,!0);if(l)return l(r,!0);var n=new Error("Cannot find module '"+r+"'");throw n.code="MODULE_NOT_FOUND",n}var i=o[r]={exports:{}};a[r][0].call(i.exports,function(e){var t=a[r][1][e];return u(t||e)},i,i.exports,s,a,o,h)}return o[r].exports}for(var l="function"==typeof require&&require,e=0;e<h.length;e++)u(h[e]);return u}({1:[function(e,t,r){"use strict";var d=e("./utils"),c=e("./support"),p="ABCDEFGHIJKLMNOPQRSTUVWXYZabcdefghijklmnopqrstuvwxyz0123456789+/=";r.encode=function(e){for(var t,r,n,i,s,a,o,h=[],u=0,l=e.length,f=l,c="string"!==d.getTypeOf(e);u<e.length;)f=l-u,n=c?(t=e[u++],r=u<l?e[u++]:0,u<l?e[u++]:0):(t=e.charCodeAt(u++),r=u<l?e.charCodeAt(u++):0,u<l?e.charCodeAt(u++):0),i=t>>2,s=(3&t)<<4|r>>4,a=1<f?(15&r)<<2|n>>6:64,o=2<f?63&n:64,h.push(p.charAt(i)+p.charAt(s)+p.charAt(a)+p.charAt(o));return h.join("")},r.decode=function(e){var t,r,n,i,s,a,o=0,h=0,u="data:";if(e.substr(0,u.length)===u)throw new Error("Invalid base64 input, it looks like a data url.");var l,f=3*(e=e.replace(/[^A-Za-z0-9+/=]/g,"")).length/4;if(e.charAt(e.length-1)===p.charAt(64)&&f--,e.charAt(e.length-2)===p.charAt(64)&&f--,f%1!=0)throw new Error("Invalid base64 input, bad content length.");for(l=c.uint8array?new Uint8Array(0|f):new Array(0|f);o<e.length;)t=p.indexOf(e.charAt(o++))<<2|(i=p.indexOf(e.charAt(o++)))>>4,r=(15&i)<<4|(s=p.indexOf(e.charAt(o++)))>>2,n=(3&s)<<6|(a=p.indexOf(e.charAt(o++))),l[h++]=t,64!==s&&(l[h++]=r),64!==a&&(l[h++]=n);return l}},{"./support":30,"./utils":32}],2:[function(e,t,r){"use strict";var n=e("./external"),i=e("./stream/DataWorker"),s=e("./stream/Crc32Probe"),a=e("./stream/DataLengthProbe");function o(e,t,r,n,i){this.compressedSize=e,this.uncompressedSize=t,this.crc32=r,this.compression=n,this.compressedContent=i}o.prototype={getContentWorker:function(){var e=new i(n.Promise.resolve(this.compressedContent)).pipe(this.compression.uncompressWorker()).pipe(new a("data_length")),t=this;return e.on("end",function(){if(this.streamInfo.data_length!==t.uncompressedSize)throw new Error("Bug : uncompressed data size mismatch")}),e},getCompressedWorker:function(){return new i(n.Promise.resolve(this.compressedContent)).withStreamInfo("compressedSize",this.compressedSize).withStreamInfo("uncompressedSize",this.uncompressedSize).withStreamInfo("crc32",this.crc32).withStreamInfo("compression",this.compression)}},o.createWorkerFrom=function(e,t,r){return e.pipe(new s).pipe(new a("uncompressedSize")).pipe(t.compressWorker(r)).pipe(new a("compressedSize")).withStreamInfo("compression",t)},t.exports=o},{"./external":6,"./stream/Crc32Probe":25,"./stream/DataLengthProbe":26,"./stream/DataWorker":27}],3:[function(e,t,r){"use strict";var n=e("./stream/GenericWorker");r.STORE={magic:"\0\0",compressWorker:function(){return new n("STORE compression")},uncompressWorker:function(){return new n("STORE decompression")}},r.DEFLATE=e("./flate")},{"./flate":7,"./stream/GenericWorker":28}],4:[function(e,t,r){"use strict";var n=e("./utils");var o=function(){for(var e,t=[],r=0;r<256;r++){e=r;for(var n=0;n<8;n++)e=1&e?3988292384^e>>>1:e>>>1;t[r]=e}return t}();t.exports=function(e,t){return void 0!==e&&e.length?"string"!==n.getTypeOf(e)?function(e,t,r,n){var i=o,s=n+r;e^=-1;for(var a=n;a<s;a++)e=e>>>8^i[255&(e^t[a])];return-1^e}(0|t,e,e.length,0):function(e,t,r,n){var i=o,s=n+r;e^=-1;for(var a=n;a<s;a++)e=e>>>8^i[255&(e^t.charCodeAt(a))];return-1^e}(0|t,e,e.length,0):0}},{"./utils":32}],5:[function(e,t,r){"use strict";r.base64=!1,r.binary=!1,r.dir=!1,r.createFolders=!0,r.date=null,r.compression=null,r.compressionOptions=null,r.comment=null,r.unixPermissions=null,r.dosPermissions=null},{}],6:[function(e,t,r){"use strict";var n=null;n="undefined"!=typeof Promise?Promise:e("lie"),t.exports={Promise:n}},{lie:37}],7:[function(e,t,r){"use strict";var n="undefined"!=typeof Uint8Array&&"undefined"!=typeof Uint16Array&&"undefined"!=typeof Uint32Array,i=e("pako"),s=e("./utils"),a=e("./stream/GenericWorker"),o=n?"uint8array":"array";function h(e,t){a.call(this,"FlateWorker/"+e),this._pako=null,this._pakoAction=e,this._pakoOptions=t,this.meta={}}r.magic="\b\0",s.inherits(h,a),h.prototype.processChunk=function(e){this.meta=e.meta,null===this._pako&&this._createPako(),this._pako.push(s.transformTo(o,e.data),!1)},h.prototype.flush=function(){a.prototype.flush.call(this),null===this._pako&&this._createPako(),this._pako.push([],!0)},h.prototype.cleanUp=function(){a.prototype.cleanUp.call(this),this._pako=null},h.prototype._createPako=function(){this._pako=new i[this._pakoAction]({raw:!0,level:this._pakoOptions.level||-1});var t=this;this._pako.onData=function(e){t.push({data:e,meta:t.meta})}},r.compressWorker=function(e){return new h("Deflate",e)},r.uncompressWorker=function(){return new h("Inflate",{})}},{"./stream/GenericWorker":28,"./utils":32,pako:38}],8:[function(e,t,r){"use strict";function A(e,t){var r,n="";for(r=0;r<t;r++)n+=String.fromCharCode(255&e),e>>>=8;return n}function n(e,t,r,n,i,s){var a,o,h=e.file,u=e.compression,l=s!==O.utf8encode,f=I.transformTo("string",s(h.name)),c=I.transformTo("string",O.utf8encode(h.name)),d=h.comment,p=I.transformTo("string",s(d)),m=I.transformTo("string",O.utf8encode(d)),_=c.length!==h.name.length,g=m.length!==d.length,b="",v="",y="",w=h.dir,k=h.date,x={crc32:0,compressedSize:0,uncompressedSize:0};t&&!r||(x.crc32=e.crc32,x.compressedSize=e.compressedSize,x.uncompressedSize=e.uncompressedSize);var S=0;t&&(S|=8),l||!_&&!g||(S|=2048);var z=0,C=0;w&&(z|=16),"UNIX"===i?(C=798,z|=function(e,t){var r=e;return e||(r=t?16893:33204),(65535&r)<<16}(h.unixPermissions,w)):(C=20,z|=function(e){return 63&(e||0)}(h.dosPermissions)),a=k.getUTCHours(),a<<=6,a|=k.getUTCMinutes(),a<<=5,a|=k.getUTCSeconds()/2,o=k.getUTCFullYear()-1980,o<<=4,o|=k.getUTCMonth()+1,o<<=5,o|=k.getUTCDate(),_&&(v=A(1,1)+A(B(f),4)+c,b+="up"+A(v.length,2)+v),g&&(y=A(1,1)+A(B(p),4)+m,b+="uc"+A(y.length,2)+y);var E="";return E+="\n\0",E+=A(S,2),E+=u.magic,E+=A(a,2),E+=A(o,2),E+=A(x.crc32,4),E+=A(x.compressedSize,4),E+=A(x.uncompressedSize,4),E+=A(f.length,2),E+=A(b.length,2),{fileRecord:R.LOCAL_FILE_HEADER+E+f+b,dirRecord:R.CENTRAL_FILE_HEADER+A(C,2)+E+A(p.length,2)+"\0\0\0\0"+A(z,4)+A(n,4)+f+b+p}}var I=e("../utils"),i=e("../stream/GenericWorker"),O=e("../utf8"),B=e("../crc32"),R=e("../signature");function s(e,t,r,n){i.call(this,"ZipFileWorker"),this.bytesWritten=0,this.zipComment=t,this.zipPlatform=r,this.encodeFileName=n,this.streamFiles=e,this.accumulate=!1,this.contentBuffer=[],this.dirRecords=[],this.currentSourceOffset=0,this.entriesCount=0,this.currentFile=null,this._sources=[]}I.inherits(s,i),s.prototype.push=function(e){var t=e.meta.percent||0,r=this.entriesCount,n=this._sources.length;this.accumulate?this.contentBuffer.push(e):(this.bytesWritten+=e.data.length,i.prototype.push.call(this,{data:e.data,meta:{currentFile:this.currentFile,percent:r?(t+100*(r-n-1))/r:100}}))},s.prototype.openedSource=function(e){this.currentSourceOffset=this.bytesWritten,this.currentFile=e.file.name;var t=this.streamFiles&&!e.file.dir;if(t){var r=n(e,t,!1,this.currentSourceOffset,this.zipPlatform,this.encodeFileName);this.push({data:r.fileRecord,meta:{percent:0}})}else this.accumulate=!0},s.prototype.closedSource=function(e){this.accumulate=!1;var t=this.streamFiles&&!e.file.dir,r=n(e,t,!0,this.currentSourceOffset,this.zipPlatform,this.encodeFileName);if(this.dirRecords.push(r.dirRecord),t)this.push({data:function(e){return R.DATA_DESCRIPTOR+A(e.crc32,4)+A(e.compressedSize,4)+A(e.uncompressedSize,4)}(e),meta:{percent:100}});else for(this.push({data:r.fileRecord,meta:{percent:0}});this.contentBuffer.length;)this.push(this.contentBuffer.shift());this.currentFile=null},s.prototype.flush=function(){for(var e=this.bytesWritten,t=0;t<this.dirRecords.length;t++)this.push({data:this.dirRecords[t],meta:{percent:100}});var r=this.bytesWritten-e,n=function(e,t,r,n,i){var s=I.transformTo("string",i(n));return R.CENTRAL_DIRECTORY_END+"\0\0\0\0"+A(e,2)+A(e,2)+A(t,4)+A(r,4)+A(s.length,2)+s}(this.dirRecords.length,r,e,this.zipComment,this.encodeFileName);this.push({data:n,meta:{percent:100}})},s.prototype.prepareNextSource=function(){this.previous=this._sources.shift(),this.openedSource(this.previous.streamInfo),this.isPaused?this.previous.pause():this.previous.resume()},s.prototype.registerPrevious=function(e){this._sources.push(e);var t=this;return e.on("data",function(e){t.processChunk(e)}),e.on("end",function(){t.closedSource(t.previous.streamInfo),t._sources.length?t.prepareNextSource():t.end()}),e.on("error",function(e){t.error(e)}),this},s.prototype.resume=function(){return!!i.prototype.resume.call(this)&&(!this.previous&&this._sources.length?(this.prepareNextSource(),!0):this.previous||this._sources.length||this.generatedError?void 0:(this.end(),!0))},s.prototype.error=function(e){var t=this._sources;if(!i.prototype.error.call(this,e))return!1;for(var r=0;r<t.length;r++)try{t[r].error(e)}catch(e){}return!0},s.prototype.lock=function(){i.prototype.lock.call(this);for(var e=this._sources,t=0;t<e.length;t++)e[t].lock()},t.exports=s},{"../crc32":4,"../signature":23,"../stream/GenericWorker":28,"../utf8":31,"../utils":32}],9:[function(e,t,r){"use strict";var u=e("../compressions"),n=e("./ZipFileWorker");r.generateWorker=function(e,a,t){var o=new n(a.streamFiles,t,a.platform,a.encodeFileName),h=0;try{e.forEach(function(e,t){h++;var r=function(e,t){var r=e||t,n=u[r];if(!n)throw new Error(r+" is not a valid compression method !");return n}(t.options.compression,a.compression),n=t.options.compressionOptions||a.compressionOptions||{},i=t.dir,s=t.date;t._compressWorker(r,n).withStreamInfo("file",{name:e,dir:i,date:s,comment:t.comment||"",unixPermissions:t.unixPermissions,dosPermissions:t.dosPermissions}).pipe(o)}),o.entriesCount=h}catch(e){o.error(e)}return o}},{"../compressions":3,"./ZipFileWorker":8}],10:[function(e,t,r){"use strict";function n(){if(!(this instanceof n))return new n;if(arguments.length)throw new Error("The constructor with parameters has been removed in JSZip 3.0, please check the upgrade guide.");this.files=Object.create(null),this.comment=null,this.root="",this.clone=function(){var e=new n;for(var t in this)"function"!=typeof this[t]&&(e[t]=this[t]);return e}}(n.prototype=e("./object")).loadAsync=e("./load"),n.support=e("./support"),n.defaults=e("./defaults"),n.version="3.10.1",n.loadAsync=function(e,t){return(new n).loadAsync(e,t)},n.external=e("./external"),t.exports=n},{"./defaults":5,"./external":6,"./load":11,"./object":15,"./support":30}],11:[function(e,t,r){"use strict";var u=e("./utils"),i=e("./external"),n=e("./utf8"),s=e("./zipEntries"),a=e("./stream/Crc32Probe"),l=e("./nodejsUtils");function f(n){return new i.Promise(function(e,t){var r=n.decompressed.getContentWorker().pipe(new a);r.on("error",function(e){t(e)}).on("end",function(){r.streamInfo.crc32!==n.decompressed.crc32?t(new Error("Corrupted zip : CRC32 mismatch")):e()}).resume()})}t.exports=function(e,o){var h=this;return o=u.extend(o||{},{base64:!1,checkCRC32:!1,optimizedBinaryString:!1,createFolders:!1,decodeFileName:n.utf8decode}),l.isNode&&l.isStream(e)?i.Promise.reject(new Error("JSZip can't accept a stream when loading a zip file.")):u.prepareContent("the loaded zip file",e,!0,o.optimizedBinaryString,o.base64).then(function(e){var t=new s(o);return t.load(e),t}).then(function(e){var t=[i.Promise.resolve(e)],r=e.files;if(o.checkCRC32)for(var n=0;n<r.length;n++)t.push(f(r[n]));return i.Promise.all(t)}).then(function(e){for(var t=e.shift(),r=t.files,n=0;n<r.length;n++){var i=r[n],s=i.fileNameStr,a=u.resolve(i.fileNameStr);h.file(a,i.decompressed,{binary:!0,optimizedBinaryString:!0,date:i.date,dir:i.dir,comment:i.fileCommentStr.length?i.fileCommentStr:null,unixPermissions:i.unixPermissions,dosPermissions:i.dosPermissions,createFolders:o.createFolders}),i.dir||(h.file(a).unsafeOriginalName=s)}return t.zipComment.length&&(h.comment=t.zipComment),h})}},{"./external":6,"./nodejsUtils":14,"./stream/Crc32Probe":25,"./utf8":31,"./utils":32,"./zipEntries":33}],12:[function(e,t,r){"use strict";var n=e("../utils"),i=e("../stream/GenericWorker");function s(e,t){i.call(this,"Nodejs stream input adapter for "+e),this._upstreamEnded=!1,this._bindStream(t)}n.inherits(s,i),s.prototype._bindStream=function(e){var t=this;(this._stream=e).pause(),e.on("data",function(e){t.push({data:e,meta:{percent:0}})}).on("error",function(e){t.isPaused?this.generatedError=e:t.error(e)}).on("end",function(){t.isPaused?t._upstreamEnded=!0:t.end()})},s.prototype.pause=function(){return!!i.prototype.pause.call(this)&&(this._stream.pause(),!0)},s.prototype.resume=function(){return!!i.prototype.resume.call(this)&&(this._upstreamEnded?this.end():this._stream.resume(),!0)},t.exports=s},{"../stream/GenericWorker":28,"../utils":32}],13:[function(e,t,r){"use strict";var i=e("readable-stream").Readable;function n(e,t,r){i.call(this,t),this._helper=e;var n=this;e.on("data",function(e,t){n.push(e)||n._helper.pause(),r&&r(t)}).on("error",function(e){n.emit("error",e)}).on("end",function(){n.push(null)})}e("../utils").inherits(n,i),n.prototype._read=function(){this._helper.resume()},t.exports=n},{"../utils":32,"readable-stream":16}],14:[function(e,t,r){"use strict";t.exports={isNode:"undefined"!=typeof Buffer,newBufferFrom:function(e,t){if(Buffer.from&&Buffer.from!==Uint8Array.from)return Buffer.from(e,t);if("number"==typeof e)throw new Error('The "data" argument must not be a number');return new Buffer(e,t)},allocBuffer:function(e){if(Buffer.alloc)return Buffer.alloc(e);var t=new Buffer(e);return t.fill(0),t},isBuffer:function(e){return Buffer.isBuffer(e)},isStream:function(e){return e&&"function"==typeof e.on&&"function"==typeof e.pause&&"function"==typeof e.resume}}},{}],15:[function(e,t,r){"use strict";function s(e,t,r){var n,i=u.getTypeOf(t),s=u.extend(r||{},f);s.date=s.date||new Date,null!==s.compression&&(s.compression=s.compression.toUpperCase()),"string"==typeof s.unixPermissions&&(s.unixPermissions=parseInt(s.unixPermissions,8)),s.unixPermissions&&16384&s.unixPermissions&&(s.dir=!0),s.dosPermissions&&16&s.dosPermissions&&(s.dir=!0),s.dir&&(e=g(e)),s.createFolders&&(n=_(e))&&b.call(this,n,!0);var a="string"===i&&!1===s.binary&&!1===s.base64;r&&void 0!==r.binary||(s.binary=!a),(t instanceof c&&0===t.uncompressedSize||s.dir||!t||0===t.length)&&(s.base64=!1,s.binary=!0,t="",s.compression="STORE",i="string");var o=null;o=t instanceof c||t instanceof l?t:p.isNode&&p.isStream(t)?new m(e,t):u.prepareContent(e,t,s.binary,s.optimizedBinaryString,s.base64);var h=new d(e,o,s);this.files[e]=h}var i=e("./utf8"),u=e("./utils"),l=e("./stream/GenericWorker"),a=e("./stream/StreamHelper"),f=e("./defaults"),c=e("./compressedObject"),d=e("./zipObject"),o=e("./generate"),p=e("./nodejsUtils"),m=e("./nodejs/NodejsStreamInputAdapter"),_=function(e){"/"===e.slice(-1)&&(e=e.substring(0,e.length-1));var t=e.lastIndexOf("/");return 0<t?e.substring(0,t):""},g=function(e){return"/"!==e.slice(-1)&&(e+="/"),e},b=function(e,t){return t=void 0!==t?t:f.createFolders,e=g(e),this.files[e]||s.call(this,e,null,{dir:!0,createFolders:t}),this.files[e]};function h(e){return"[object RegExp]"===Object.prototype.toString.call(e)}var n={load:function(){throw new Error("This method has been removed in JSZip 3.0, please check the upgrade guide.")},forEach:function(e){var t,r,n;for(t in this.files)n=this.files[t],(r=t.slice(this.root.length,t.length))&&t.slice(0,this.root.length)===this.root&&e(r,n)},filter:function(r){var n=[];return this.forEach(function(e,t){r(e,t)&&n.push(t)}),n},file:function(e,t,r){if(1!==arguments.length)return e=this.root+e,s.call(this,e,t,r),this;if(h(e)){var n=e;return this.filter(function(e,t){return!t.dir&&n.test(e)})}var i=this.files[this.root+e];return i&&!i.dir?i:null},folder:function(r){if(!r)return this;if(h(r))return this.filter(function(e,t){return t.dir&&r.test(e)});var e=this.root+r,t=b.call(this,e),n=this.clone();return n.root=t.name,n},remove:function(r){r=this.root+r;var e=this.files[r];if(e||("/"!==r.slice(-1)&&(r+="/"),e=this.files[r]),e&&!e.dir)delete this.files[r];else for(var t=this.filter(function(e,t){return t.name.slice(0,r.length)===r}),n=0;n<t.length;n++)delete this.files[t[n].name];return this},generate:function(){throw new Error("This method has been removed in JSZip 3.0, please check the upgrade guide.")},generateInternalStream:function(e){var t,r={};try{if((r=u.extend(e||{},{streamFiles:!1,compression:"STORE",compressionOptions:null,type:"",platform:"DOS",comment:null,mimeType:"application/zip",encodeFileName:i.utf8encode})).type=r.type.toLowerCase(),r.compression=r.compression.toUpperCase(),"binarystring"===r.type&&(r.type="string"),!r.type)throw new Error("No output type specified.");u.checkSupport(r.type),"darwin"!==r.platform&&"freebsd"!==r.platform&&"linux"!==r.platform&&"sunos"!==r.platform||(r.platform="UNIX"),"win32"===r.platform&&(r.platform="DOS");var n=r.comment||this.comment||"";t=o.generateWorker(this,r,n)}catch(e){(t=new l("error")).error(e)}return new a(t,r.type||"string",r.mimeType)},generateAsync:function(e,t){return this.generateInternalStream(e).accumulate(t)},generateNodeStream:function(e,t){return(e=e||{}).type||(e.type="nodebuffer"),this.generateInternalStream(e).toNodejsStream(t)}};t.exports=n},{"./compressedObject":2,"./defaults":5,"./generate":9,"./nodejs/NodejsStreamInputAdapter":12,"./nodejsUtils":14,"./stream/GenericWorker":28,"./stream/StreamHelper":29,"./utf8":31,"./utils":32,"./zipObject":35}],16:[function(e,t,r){"use strict";t.exports=e("stream")},{stream:void 0}],17:[function(e,t,r){"use strict";var n=e("./DataReader");function i(e){n.call(this,e);for(var t=0;t<this.data.length;t++)e[t]=255&e[t]}e("../utils").inherits(i,n),i.prototype.byteAt=function(e){return this.data[this.zero+e]},i.prototype.lastIndexOfSignature=function(e){for(var t=e.charCodeAt(0),r=e.charCodeAt(1),n=e.charCodeAt(2),i=e.charCodeAt(3),s=this.length-4;0<=s;--s)if(this.data[s]===t&&this.data[s+1]===r&&this.data[s+2]===n&&this.data[s+3]===i)return s-this.zero;return-1},i.prototype.readAndCheckSignature=function(e){var t=e.charCodeAt(0),r=e.charCodeAt(1),n=e.charCodeAt(2),i=e.charCodeAt(3),s=this.readData(4);return t===s[0]&&r===s[1]&&n===s[2]&&i===s[3]},i.prototype.readData=function(e){if(this.checkOffset(e),0===e)return[];var t=this.data.slice(this.zero+this.index,this.zero+this.index+e);return this.index+=e,t},t.exports=i},{"../utils":32,"./DataReader":18}],18:[function(e,t,r){"use strict";var n=e("../utils");function i(e){this.data=e,this.length=e.length,this.index=0,this.zero=0}i.prototype={checkOffset:function(e){this.checkIndex(this.index+e)},checkIndex:function(e){if(this.length<this.zero+e||e<0)throw new Error("End of data reached (data length = "+this.length+", asked index = "+e+"). Corrupted zip ?")},setIndex:function(e){this.checkIndex(e),this.index=e},skip:function(e){this.setIndex(this.index+e)},byteAt:function(){},readInt:function(e){var t,r=0;for(this.checkOffset(e),t=this.index+e-1;t>=this.index;t--)r=(r<<8)+this.byteAt(t);return this.index+=e,r},readString:function(e){return n.transformTo("string",this.readData(e))},readData:function(){},lastIndexOfSignature:function(){},readAndCheckSignature:function(){},readDate:function(){var e=this.readInt(4);return new Date(Date.UTC(1980+(e>>25&127),(e>>21&15)-1,e>>16&31,e>>11&31,e>>5&63,(31&e)<<1))}},t.exports=i},{"../utils":32}],19:[function(e,t,r){"use strict";var n=e("./Uint8ArrayReader");function i(e){n.call(this,e)}e("../utils").inherits(i,n),i.prototype.readData=function(e){this.checkOffset(e);var t=this.data.slice(this.zero+this.index,this.zero+this.index+e);return this.index+=e,t},t.exports=i},{"../utils":32,"./Uint8ArrayReader":21}],20:[function(e,t,r){"use strict";var n=e("./DataReader");function i(e){n.call(this,e)}e("../utils").inherits(i,n),i.prototype.byteAt=function(e){return this.data.charCodeAt(this.zero+e)},i.prototype.lastIndexOfSignature=function(e){return this.data.lastIndexOf(e)-this.zero},i.prototype.readAndCheckSignature=function(e){return e===this.readData(4)},i.prototype.readData=function(e){this.checkOffset(e);var t=this.data.slice(this.zero+this.index,this.zero+this.index+e);return this.index+=e,t},t.exports=i},{"../utils":32,"./DataReader":18}],21:[function(e,t,r){"use strict";var n=e("./ArrayReader");function i(e){n.call(this,e)}e("../utils").inherits(i,n),i.prototype.readData=function(e){if(this.checkOffset(e),0===e)return new Uint8Array(0);var t=this.data.subarray(this.zero+this.index,this.zero+this.index+e);return this.index+=e,t},t.exports=i},{"../utils":32,"./ArrayReader":17}],22:[function(e,t,r){"use strict";var n=e("../utils"),i=e("../support"),s=e("./ArrayReader"),a=e("./StringReader"),o=e("./NodeBufferReader"),h=e("./Uint8ArrayReader");t.exports=function(e){var t=n.getTypeOf(e);return n.checkSupport(t),"string"!==t||i.uint8array?"nodebuffer"===t?new o(e):i.uint8array?new h(n.transformTo("uint8array",e)):new s(n.transformTo("array",e)):new a(e)}},{"../support":30,"../utils":32,"./ArrayReader":17,"./NodeBufferReader":19,"./StringReader":20,"./Uint8ArrayReader":21}],23:[function(e,t,r){"use strict";r.LOCAL_FILE_HEADER="PK",r.CENTRAL_FILE_HEADER="PK",r.CENTRAL_DIRECTORY_END="PK",r.ZIP64_CENTRAL_DIRECTORY_LOCATOR="PK",r.ZIP64_CENTRAL_DIRECTORY_END="PK",r.DATA_DESCRIPTOR="PK\b"},{}],24:[function(e,t,r){"use strict";var n=e("./GenericWorker"),i=e("../utils");function s(e){n.call(this,"ConvertWorker to "+e),this.destType=e}i.inherits(s,n),s.prototype.processChunk=function(e){this.push({data:i.transformTo(this.destType,e.data),meta:e.meta})},t.exports=s},{"../utils":32,"./GenericWorker":28}],25:[function(e,t,r){"use strict";var n=e("./GenericWorker"),i=e("../crc32");function s(){n.call(this,"Crc32Probe"),this.withStreamInfo("crc32",0)}e("../utils").inherits(s,n),s.prototype.processChunk=function(e){this.streamInfo.crc32=i(e.data,this.streamInfo.crc32||0),this.push(e)},t.exports=s},{"../crc32":4,"../utils":32,"./GenericWorker":28}],26:[function(e,t,r){"use strict";var n=e("../utils"),i=e("./GenericWorker");function s(e){i.call(this,"DataLengthProbe for "+e),this.propName=e,this.withStreamInfo(e,0)}n.inherits(s,i),s.prototype.processChunk=function(e){if(e){var t=this.streamInfo[this.propName]||0;this.streamInfo[this.propName]=t+e.data.length}i.prototype.processChunk.call(this,e)},t.exports=s},{"../utils":32,"./GenericWorker":28}],27:[function(e,t,r){"use strict";var n=e("../utils"),i=e("./GenericWorker");function s(e){i.call(this,"DataWorker");var t=this;this.dataIsReady=!1,this.index=0,this.max=0,this.data=null,this.type="",this._tickScheduled=!1,e.then(function(e){t.dataIsReady=!0,t.data=e,t.max=e&&e.length||0,t.type=n.getTypeOf(e),t.isPaused||t._tickAndRepeat()},function(e){t.error(e)})}n.inherits(s,i),s.prototype.cleanUp=function(){i.prototype.cleanUp.call(this),this.data=null},s.prototype.resume=function(){return!!i.prototype.resume.call(this)&&(!this._tickScheduled&&this.dataIsReady&&(this._tickScheduled=!0,n.delay(this._tickAndRepeat,[],this)),!0)},s.prototype._tickAndRepeat=function(){this._tickScheduled=!1,this.isPaused||this.isFinished||(this._tick(),this.isFinished||(n.delay(this._tickAndRepeat,[],this),this._tickScheduled=!0))},s.prototype._tick=function(){if(this.isPaused||this.isFinished)return!1;var e=null,t=Math.min(this.max,this.index+16384);if(this.index>=this.max)return this.end();switch(this.type){case"string":e=this.data.substring(this.index,t);break;case"uint8array":e=this.data.subarray(this.index,t);break;case"array":case"nodebuffer":e=this.data.slice(this.index,t)}return this.index=t,this.push({data:e,meta:{percent:this.max?this.index/this.max*100:0}})},t.exports=s},{"../utils":32,"./GenericWorker":28}],28:[function(e,t,r){"use strict";function n(e){this.name=e||"default",this.streamInfo={},this.generatedError=null,this.extraStreamInfo={},this.isPaused=!0,this.isFinished=!1,this.isLocked=!1,this._listeners={data:[],end:[],error:[]},this.previous=null}n.prototype={push:function(e){this.emit("data",e)},end:function(){if(this.isFinished)return!1;this.flush();try{this.emit("end"),this.cleanUp(),this.isFinished=!0}catch(e){this.emit("error",e)}return!0},error:function(e){return!this.isFinished&&(this.isPaused?this.generatedError=e:(this.isFinished=!0,this.emit("error",e),this.previous&&this.previous.error(e),this.cleanUp()),!0)},on:function(e,t){return this._listeners[e].push(t),this},cleanUp:function(){this.streamInfo=this.generatedError=this.extraStreamInfo=null,this._listeners=[]},emit:function(e,t){if(this._listeners[e])for(var r=0;r<this._listeners[e].length;r++)this._listeners[e][r].call(this,t)},pipe:function(e){return e.registerPrevious(this)},registerPrevious:function(e){if(this.isLocked)throw new Error("The stream '"+this+"' has already been used.");this.streamInfo=e.streamInfo,this.mergeStreamInfo(),this.previous=e;var t=this;return e.on("data",function(e){t.processChunk(e)}),e.on("end",function(){t.end()}),e.on("error",function(e){t.error(e)}),this},pause:function(){return!this.isPaused&&!this.isFinished&&(this.isPaused=!0,this.previous&&this.previous.pause(),!0)},resume:function(){if(!this.isPaused||this.isFinished)return!1;var e=this.isPaused=!1;return this.generatedError&&(this.error(this.generatedError),e=!0),this.previous&&this.previous.resume(),!e},flush:function(){},processChunk:function(e){this.push(e)},withStreamInfo:function(e,t){return this.extraStreamInfo[e]=t,this.mergeStreamInfo(),this},mergeStreamInfo:function(){for(var e in this.extraStreamInfo)Object.prototype.hasOwnProperty.call(this.extraStreamInfo,e)&&(this.streamInfo[e]=this.extraStreamInfo[e])},lock:function(){if(this.isLocked)throw new Error("The stream '"+this+"' has already been used.");this.isLocked=!0,this.previous&&this.previous.lock()},toString:function(){var e="Worker "+this.name;return this.previous?this.previous+" -> "+e:e}},t.exports=n},{}],29:[function(e,t,r){"use strict";var h=e("../utils"),i=e("./ConvertWorker"),s=e("./GenericWorker"),u=e("../base64"),n=e("../support"),a=e("../external"),o=null;if(n.nodestream)try{o=e("../nodejs/NodejsStreamOutputAdapter")}catch(e){}function l(e,o){return new a.Promise(function(t,r){var n=[],i=e._internalType,s=e._outputType,a=e._mimeType;e.on("data",function(e,t){n.push(e),o&&o(t)}).on("error",function(e){n=[],r(e)}).on("end",function(){try{var e=function(e,t,r){switch(e){case"blob":return h.newBlob(h.transformTo("arraybuffer",t),r);case"base64":return u.encode(t);default:return h.transformTo(e,t)}}(s,function(e,t){var r,n=0,i=null,s=0;for(r=0;r<t.length;r++)s+=t[r].length;switch(e){case"string":return t.join("");case"array":return Array.prototype.concat.apply([],t);case"uint8array":for(i=new Uint8Array(s),r=0;r<t.length;r++)i.set(t[r],n),n+=t[r].length;return i;case"nodebuffer":return Buffer.concat(t);default:throw new Error("concat : unsupported type '"+e+"'")}}(i,n),a);t(e)}catch(e){r(e)}n=[]}).resume()})}function f(e,t,r){var n=t;switch(t){case"blob":case"arraybuffer":n="uint8array";break;case"base64":n="string"}try{this._internalType=n,this._outputType=t,this._mimeType=r,h.checkSupport(n),this._worker=e.pipe(new i(n)),e.lock()}catch(e){this._worker=new s("error"),this._worker.error(e)}}f.prototype={accumulate:function(e){return l(this,e)},on:function(e,t){var r=this;return"data"===e?this._worker.on(e,function(e){t.call(r,e.data,e.meta)}):this._worker.on(e,function(){h.delay(t,arguments,r)}),this},resume:function(){return h.delay(this._worker.resume,[],this._worker),this},pause:function(){return this._worker.pause(),this},toNodejsStream:function(e){if(h.checkSupport("nodestream"),"nodebuffer"!==this._outputType)throw new Error(this._outputType+" is not supported by this method");return new o(this,{objectMode:"nodebuffer"!==this._outputType},e)}},t.exports=f},{"../base64":1,"../external":6,"../nodejs/NodejsStreamOutputAdapter":13,"../support":30,"../utils":32,"./ConvertWorker":24,"./GenericWorker":28}],30:[function(e,t,r){"use strict";if(r.base64=!0,r.array=!0,r.string=!0,r.arraybuffer="undefined"!=typeof ArrayBuffer&&"undefined"!=typeof Uint8Array,r.nodebuffer="undefined"!=typeof Buffer,r.uint8array="undefined"!=typeof Uint8Array,"undefined"==typeof ArrayBuffer)r.blob=!1;else{var n=new ArrayBuffer(0);try{r.blob=0===new Blob([n],{type:"application/zip"}).size}catch(e){try{var i=new(self.BlobBuilder||self.WebKitBlobBuilder||self.MozBlobBuilder||self.MSBlobBuilder);i.append(n),r.blob=0===i.getBlob("application/zip").size}catch(e){r.blob=!1}}}try{r.nodestream=!!e("readable-stream").Readable}catch(e){r.nodestream=!1}},{"readable-stream":16}],31:[function(e,t,s){"use strict";for(var o=e("./utils"),h=e("./support"),r=e("./nodejsUtils"),n=e("./stream/GenericWorker"),u=new Array(256),i=0;i<256;i++)u[i]=252<=i?6:248<=i?5:240<=i?4:224<=i?3:192<=i?2:1;u[254]=u[254]=1;function a(){n.call(this,"utf-8 decode"),this.leftOver=null}function l(){n.call(this,"utf-8 encode")}s.utf8encode=function(e){return h.nodebuffer?r.newBufferFrom(e,"utf-8"):function(e){var t,r,n,i,s,a=e.length,o=0;for(i=0;i<a;i++)55296==(64512&(r=e.charCodeAt(i)))&&i+1<a&&56320==(64512&(n=e.charCodeAt(i+1)))&&(r=65536+(r-55296<<10)+(n-56320),i++),o+=r<128?1:r<2048?2:r<65536?3:4;for(t=h.uint8array?new Uint8Array(o):new Array(o),i=s=0;s<o;i++)55296==(64512&(r=e.charCodeAt(i)))&&i+1<a&&56320==(64512&(n=e.charCodeAt(i+1)))&&(r=65536+(r-55296<<10)+(n-56320),i++),r<128?t[s++]=r:(r<2048?t[s++]=192|r>>>6:(r<65536?t[s++]=224|r>>>12:(t[s++]=240|r>>>18,t[s++]=128|r>>>12&63),t[s++]=128|r>>>6&63),t[s++]=128|63&r);return t}(e)},s.utf8decode=function(e){return h.nodebuffer?o.transformTo("nodebuffer",e).toString("utf-8"):function(e){var t,r,n,i,s=e.length,a=new Array(2*s);for(t=r=0;t<s;)if((n=e[t++])<128)a[r++]=n;else if(4<(i=u[n]))a[r++]=65533,t+=i-1;else{for(n&=2===i?31:3===i?15:7;1<i&&t<s;)n=n<<6|63&e[t++],i--;1<i?a[r++]=65533:n<65536?a[r++]=n:(n-=65536,a[r++]=55296|n>>10&1023,a[r++]=56320|1023&n)}return a.length!==r&&(a.subarray?a=a.subarray(0,r):a.length=r),o.applyFromCharCode(a)}(e=o.transformTo(h.uint8array?"uint8array":"array",e))},o.inherits(a,n),a.prototype.processChunk=function(e){var t=o.transformTo(h.uint8array?"uint8array":"array",e.data);if(this.leftOver&&this.leftOver.length){if(h.uint8array){var r=t;(t=new Uint8Array(r.length+this.leftOver.length)).set(this.leftOver,0),t.set(r,this.leftOver.length)}else t=this.leftOver.concat(t);this.leftOver=null}var n=function(e,t){var r;for((t=t||e.length)>e.length&&(t=e.length),r=t-1;0<=r&&128==(192&e[r]);)r--;return r<0?t:0===r?t:r+u[e[r]]>t?r:t}(t),i=t;n!==t.length&&(h.uint8array?(i=t.subarray(0,n),this.leftOver=t.subarray(n,t.length)):(i=t.slice(0,n),this.leftOver=t.slice(n,t.length))),this.push({data:s.utf8decode(i),meta:e.meta})},a.prototype.flush=function(){this.leftOver&&this.leftOver.length&&(this.push({data:s.utf8decode(this.leftOver),meta:{}}),this.leftOver=null)},s.Utf8DecodeWorker=a,o.inherits(l,n),l.prototype.processChunk=function(e){this.push({data:s.utf8encode(e.data),meta:e.meta})},s.Utf8EncodeWorker=l},{"./nodejsUtils":14,"./stream/GenericWorker":28,"./support":30,"./utils":32}],32:[function(e,t,a){"use strict";var o=e("./support"),h=e("./base64"),r=e("./nodejsUtils"),u=e("./external");function n(e){return e}function l(e,t){for(var r=0;r<e.length;++r)t[r]=255&e.charCodeAt(r);return t}e("setimmediate"),a.newBlob=function(t,r){a.checkSupport("blob");try{return new Blob([t],{type:r})}catch(e){try{var n=new(self.BlobBuilder||self.WebKitBlobBuilder||self.MozBlobBuilder||self.MSBlobBuilder);return n.append(t),n.getBlob(r)}catch(e){throw new Error("Bug : can't construct the Blob.")}}};var i={stringifyByChunk:function(e,t,r){var n=[],i=0,s=e.length;if(s<=r)return String.fromCharCode.apply(null,e);for(;i<s;)"array"===t||"nodebuffer"===t?n.push(String.fromCharCode.apply(null,e.slice(i,Math.min(i+r,s)))):n.push(String.fromCharCode.apply(null,e.subarray(i,Math.min(i+r,s)))),i+=r;return n.join("")},stringifyByChar:function(e){for(var t="",r=0;r<e.length;r++)t+=String.fromCharCode(e[r]);return t},applyCanBeUsed:{uint8array:function(){try{return o.uint8array&&1===String.fromCharCode.apply(null,new Uint8Array(1)).length}catch(e){return!1}}(),nodebuffer:function(){try{return o.nodebuffer&&1===String.fromCharCode.apply(null,r.allocBuffer(1)).length}catch(e){return!1}}()}};function s(e){var t=65536,r=a.getTypeOf(e),n=!0;if("uint8array"===r?n=i.applyCanBeUsed.uint8array:"nodebuffer"===r&&(n=i.applyCanBeUsed.nodebuffer),n)for(;1<t;)try{return i.stringifyByChunk(e,r,t)}catch(e){t=Math.floor(t/2)}return i.stringifyByChar(e)}function f(e,t){for(var r=0;r<e.length;r++)t[r]=e[r];return t}a.applyFromCharCode=s;var c={};c.string={string:n,array:function(e){return l(e,new Array(e.length))},arraybuffer:function(e){return c.string.uint8array(e).buffer},uint8array:function(e){return l(e,new Uint8Array(e.length))},nodebuffer:function(e){return l(e,r.allocBuffer(e.length))}},c.array={string:s,array:n,arraybuffer:function(e){return new Uint8Array(e).buffer},uint8array:function(e){return new Uint8Array(e)},nodebuffer:function(e){return r.newBufferFrom(e)}},c.arraybuffer={string:function(e){return s(new Uint8Array(e))},array:function(e){return f(new Uint8Array(e),new Array(e.byteLength))},arraybuffer:n,uint8array:function(e){return new Uint8Array(e)},nodebuffer:function(e){return r.newBufferFrom(new Uint8Array(e))}},c.uint8array={string:s,array:function(e){return f(e,new Array(e.length))},arraybuffer:function(e){return e.buffer},uint8array:n,nodebuffer:function(e){return r.newBufferFrom(e)}},c.nodebuffer={string:s,array:function(e){return f(e,new Array(e.length))},arraybuffer:function(e){return c.nodebuffer.uint8array(e).buffer},uint8array:function(e){return f(e,new Uint8Array(e.length))},nodebuffer:n},a.transformTo=function(e,t){if(t=t||"",!e)return t;a.checkSupport(e);var r=a.getTypeOf(t);return c[r][e](t)},a.resolve=function(e){for(var t=e.split("/"),r=[],n=0;n<t.length;n++){var i=t[n];"."===i||""===i&&0!==n&&n!==t.length-1||(".."===i?r.pop():r.push(i))}return r.join("/")},a.getTypeOf=function(e){return"string"==typeof e?"string":"[object Array]"===Object.prototype.toString.call(e)?"array":o.nodebuffer&&r.isBuffer(e)?"nodebuffer":o.uint8array&&e instanceof Uint8Array?"uint8array":o.arraybuffer&&e instanceof ArrayBuffer?"arraybuffer":void 0},a.checkSupport=function(e){if(!o[e.toLowerCase()])throw new Error(e+" is not supported by this platform")},a.MAX_VALUE_16BITS=65535,a.MAX_VALUE_32BITS=-1,a.pretty=function(e){var t,r,n="";for(r=0;r<(e||"").length;r++)n+="\\x"+((t=e.charCodeAt(r))<16?"0":"")+t.toString(16).toUpperCase();return n},a.delay=function(e,t,r){setImmediate(function(){e.apply(r||null,t||[])})},a.inherits=function(e,t){function r(){}r.prototype=t.prototype,e.prototype=new r},a.extend=function(){var e,t,r={};for(e=0;e<arguments.length;e++)for(t in arguments[e])Object.prototype.hasOwnProperty.call(arguments[e],t)&&void 0===r[t]&&(r[t]=arguments[e][t]);return r},a.prepareContent=function(r,e,n,i,s){return u.Promise.resolve(e).then(function(n){return o.blob&&(n instanceof Blob||-1!==["[object File]","[object Blob]"].indexOf(Object.prototype.toString.call(n)))&&"undefined"!=typeof FileReader?new u.Promise(function(t,r){var e=new FileReader;e.onload=function(e){t(e.target.result)},e.onerror=function(e){r(e.target.error)},e.readAsArrayBuffer(n)}):n}).then(function(e){var t=a.getTypeOf(e);return t?("arraybuffer"===t?e=a.transformTo("uint8array",e):"string"===t&&(s?e=h.decode(e):n&&!0!==i&&(e=function(e){return l(e,o.uint8array?new Uint8Array(e.length):new Array(e.length))}(e))),e):u.Promise.reject(new Error("Can't read the data of '"+r+"'. Is it in a supported JavaScript type (String, Blob, ArrayBuffer, etc) ?"))})}},{"./base64":1,"./external":6,"./nodejsUtils":14,"./support":30,setimmediate:54}],33:[function(e,t,r){"use strict";var n=e("./reader/readerFor"),i=e("./utils"),s=e("./signature"),a=e("./zipEntry"),o=e("./support");function h(e){this.files=[],this.loadOptions=e}h.prototype={checkSignature:function(e){if(!this.reader.readAndCheckSignature(e)){this.reader.index-=4;var t=this.reader.readString(4);throw new Error("Corrupted zip or bug: unexpected signature ("+i.pretty(t)+", expected "+i.pretty(e)+")")}},isSignature:function(e,t){var r=this.reader.index;this.reader.setIndex(e);var n=this.reader.readString(4)===t;return this.reader.setIndex(r),n},readBlockEndOfCentral:function(){this.diskNumber=this.reader.readInt(2),this.diskWithCentralDirStart=this.reader.readInt(2),this.centralDirRecordsOnThisDisk=this.reader.readInt(2),this.centralDirRecords=this.reader.readInt(2),this.centralDirSize=this.reader.readInt(4),this.centralDirOffset=this.reader.readInt(4),this.zipCommentLength=this.reader.readInt(2);var e=this.reader.readData(this.zipCommentLength),t=o.uint8array?"uint8array":"array",r=i.transformTo(t,e);this.zipComment=this.loadOptions.decodeFileName(r)},readBlockZip64EndOfCentral:function(){this.zip64EndOfCentralSize=this.reader.readInt(8),this.reader.skip(4),this.diskNumber=this.reader.readInt(4),this.diskWithCentralDirStart=this.reader.readInt(4),this.centralDirRecordsOnThisDisk=this.reader.readInt(8),this.centralDirRecords=this.reader.readInt(8),this.centralDirSize=this.reader.readInt(8),this.centralDirOffset=this.reader.readInt(8),this.zip64ExtensibleData={};for(var e,t,r,n=this.zip64EndOfCentralSize-44;0<n;)e=this.reader.readInt(2),t=this.reader.readInt(4),r=this.reader.readData(t),this.zip64ExtensibleData[e]={id:e,length:t,value:r}},readBlockZip64EndOfCentralLocator:function(){if(this.diskWithZip64CentralDirStart=this.reader.readInt(4),this.relativeOffsetEndOfZip64CentralDir=this.reader.readInt(8),this.disksCount=this.reader.readInt(4),1<this.disksCount)throw new Error("Multi-volumes zip are not supported")},readLocalFiles:function(){var e,t;for(e=0;e<this.files.length;e++)t=this.files[e],this.reader.setIndex(t.localHeaderOffset),this.checkSignature(s.LOCAL_FILE_HEADER),t.readLocalPart(this.reader),t.handleUTF8(),t.processAttributes()},readCentralDir:function(){var e;for(this.reader.setIndex(this.centralDirOffset);this.reader.readAndCheckSignature(s.CENTRAL_FILE_HEADER);)(e=new a({zip64:this.zip64},this.loadOptions)).readCentralPart(this.reader),this.files.push(e);if(this.centralDirRecords!==this.files.length&&0!==this.centralDirRecords&&0===this.files.length)throw new Error("Corrupted zip or bug: expected "+this.centralDirRecords+" records in central dir, got "+this.files.length)},readEndOfCentral:function(){var e=this.reader.lastIndexOfSignature(s.CENTRAL_DIRECTORY_END);if(e<0)throw!this.isSignature(0,s.LOCAL_FILE_HEADER)?new Error("Can't find end of central directory : is this a zip file ? If it is, see https://stuk.github.io/jszip/documentation/howto/read_zip.html"):new Error("Corrupted zip: can't find end of central directory");this.reader.setIndex(e);var t=e;if(this.checkSignature(s.CENTRAL_DIRECTORY_END),this.readBlockEndOfCentral(),this.diskNumber===i.MAX_VALUE_16BITS||this.diskWithCentralDirStart===i.MAX_VALUE_16BITS||this.centralDirRecordsOnThisDisk===i.MAX_VALUE_16BITS||this.centralDirRecords===i.MAX_VALUE_16BITS||this.centralDirSize===i.MAX_VALUE_32BITS||this.centralDirOffset===i.MAX_VALUE_32BITS){if(this.zip64=!0,(e=this.reader.lastIndexOfSignature(s.ZIP64_CENTRAL_DIRECTORY_LOCATOR))<0)throw new Error("Corrupted zip: can't find the ZIP64 end of central directory locator");if(this.reader.setIndex(e),this.checkSignature(s.ZIP64_CENTRAL_DIRECTORY_LOCATOR),this.readBlockZip64EndOfCentralLocator(),!this.isSignature(this.relativeOffsetEndOfZip64CentralDir,s.ZIP64_CENTRAL_DIRECTORY_END)&&(this.relativeOffsetEndOfZip64CentralDir=this.reader.lastIndexOfSignature(s.ZIP64_CENTRAL_DIRECTORY_END),this.relativeOffsetEndOfZip64CentralDir<0))throw new Error("Corrupted zip: can't find the ZIP64 end of central directory");this.reader.setIndex(this.relativeOffsetEndOfZip64CentralDir),this.checkSignature(s.ZIP64_CENTRAL_DIRECTORY_END),this.readBlockZip64EndOfCentral()}var r=this.centralDirOffset+this.centralDirSize;this.zip64&&(r+=20,r+=12+this.zip64EndOfCentralSize);var n=t-r;if(0<n)this.isSignature(t,s.CENTRAL_FILE_HEADER)||(this.reader.zero=n);else if(n<0)throw new Error("Corrupted zip: missing "+Math.abs(n)+" bytes.")},prepareReader:function(e){this.reader=n(e)},load:function(e){this.prepareReader(e),this.readEndOfCentral(),this.readCentralDir(),this.readLocalFiles()}},t.exports=h},{"./reader/readerFor":22,"./signature":23,"./support":30,"./utils":32,"./zipEntry":34}],34:[function(e,t,r){"use strict";var n=e("./reader/readerFor"),s=e("./utils"),i=e("./compressedObject"),a=e("./crc32"),o=e("./utf8"),h=e("./compressions"),u=e("./support");function l(e,t){this.options=e,this.loadOptions=t}l.prototype={isEncrypted:function(){return 1==(1&this.bitFlag)},useUTF8:function(){return 2048==(2048&this.bitFlag)},readLocalPart:function(e){var t,r;if(e.skip(22),this.fileNameLength=e.readInt(2),r=e.readInt(2),this.fileName=e.readData(this.fileNameLength),e.skip(r),-1===this.compressedSize||-1===this.uncompressedSize)throw new Error("Bug or corrupted zip : didn't get enough information from the central directory (compressedSize === -1 || uncompressedSize === -1)");if(null===(t=function(e){for(var t in h)if(Object.prototype.hasOwnProperty.call(h,t)&&h[t].magic===e)return h[t];return null}(this.compressionMethod)))throw new Error("Corrupted zip : compression "+s.pretty(this.compressionMethod)+" unknown (inner file : "+s.transformTo("string",this.fileName)+")");this.decompressed=new i(this.compressedSize,this.uncompressedSize,this.crc32,t,e.readData(this.compressedSize))},readCentralPart:function(e){this.versionMadeBy=e.readInt(2),e.skip(2),this.bitFlag=e.readInt(2),this.compressionMethod=e.readString(2),this.date=e.readDate(),this.crc32=e.readInt(4),this.compressedSize=e.readInt(4),this.uncompressedSize=e.readInt(4);var t=e.readInt(2);if(this.extraFieldsLength=e.readInt(2),this.fileCommentLength=e.readInt(2),this.diskNumberStart=e.readInt(2),this.internalFileAttributes=e.readInt(2),this.externalFileAttributes=e.readInt(4),this.localHeaderOffset=e.readInt(4),this.isEncrypted())throw new Error("Encrypted zip are not supported");e.skip(t),this.readExtraFields(e),this.parseZIP64ExtraField(e),this.fileComment=e.readData(this.fileCommentLength)},processAttributes:function(){this.unixPermissions=null,this.dosPermissions=null;var e=this.versionMadeBy>>8;this.dir=!!(16&this.externalFileAttributes),0==e&&(this.dosPermissions=63&this.externalFileAttributes),3==e&&(this.unixPermissions=this.externalFileAttributes>>16&65535),this.dir||"/"!==this.fileNameStr.slice(-1)||(this.dir=!0)},parseZIP64ExtraField:function(){if(this.extraFields[1]){var e=n(this.extraFields[1].value);this.uncompressedSize===s.MAX_VALUE_32BITS&&(this.uncompressedSize=e.readInt(8)),this.compressedSize===s.MAX_VALUE_32BITS&&(this.compressedSize=e.readInt(8)),this.localHeaderOffset===s.MAX_VALUE_32BITS&&(this.localHeaderOffset=e.readInt(8)),this.diskNumberStart===s.MAX_VALUE_32BITS&&(this.diskNumberStart=e.readInt(4))}},readExtraFields:function(e){var t,r,n,i=e.index+this.extraFieldsLength;for(this.extraFields||(this.extraFields={});e.index+4<i;)t=e.readInt(2),r=e.readInt(2),n=e.readData(r),this.extraFields[t]={id:t,length:r,value:n};e.setIndex(i)},handleUTF8:function(){var e=u.uint8array?"uint8array":"array";if(this.useUTF8())this.fileNameStr=o.utf8decode(this.fileName),this.fileCommentStr=o.utf8decode(this.fileComment);else{var t=this.findExtraFieldUnicodePath();if(null!==t)this.fileNameStr=t;else{var r=s.transformTo(e,this.fileName);this.fileNameStr=this.loadOptions.decodeFileName(r)}var n=this.findExtraFieldUnicodeComment();if(null!==n)this.fileCommentStr=n;else{var i=s.transformTo(e,this.fileComment);this.fileCommentStr=this.loadOptions.decodeFileName(i)}}},findExtraFieldUnicodePath:function(){var e=this.extraFields[28789];if(e){var t=n(e.value);return 1!==t.readInt(1)?null:a(this.fileName)!==t.readInt(4)?null:o.utf8decode(t.readData(e.length-5))}return null},findExtraFieldUnicodeComment:function(){var e=this.extraFields[25461];if(e){var t=n(e.value);return 1!==t.readInt(1)?null:a(this.fileComment)!==t.readInt(4)?null:o.utf8decode(t.readData(e.length-5))}return null}},t.exports=l},{"./compressedObject":2,"./compressions":3,"./crc32":4,"./reader/readerFor":22,"./support":30,"./utf8":31,"./utils":32}],35:[function(e,t,r){"use strict";function n(e,t,r){this.name=e,this.dir=r.dir,this.date=r.date,this.comment=r.comment,this.unixPermissions=r.unixPermissions,this.dosPermissions=r.dosPermissions,this._data=t,this._dataBinary=r.binary,this.options={compression:r.compression,compressionOptions:r.compressionOptions}}var s=e("./stream/StreamHelper"),i=e("./stream/DataWorker"),a=e("./utf8"),o=e("./compressedObject"),h=e("./stream/GenericWorker");n.prototype={internalStream:function(e){var t=null,r="string";try{if(!e)throw new Error("No output type specified.");var n="string"===(r=e.toLowerCase())||"text"===r;"binarystring"!==r&&"text"!==r||(r="string"),t=this._decompressWorker();var i=!this._dataBinary;i&&!n&&(t=t.pipe(new a.Utf8EncodeWorker)),!i&&n&&(t=t.pipe(new a.Utf8DecodeWorker))}catch(e){(t=new h("error")).error(e)}return new s(t,r,"")},async:function(e,t){return this.internalStream(e).accumulate(t)},nodeStream:function(e,t){return this.internalStream(e||"nodebuffer").toNodejsStream(t)},_compressWorker:function(e,t){if(this._data instanceof o&&this._data.compression.magic===e.magic)return this._data.getCompressedWorker();var r=this._decompressWorker();return this._dataBinary||(r=r.pipe(new a.Utf8EncodeWorker)),o.createWorkerFrom(r,e,t)},_decompressWorker:function(){return this._data instanceof o?this._data.getContentWorker():this._data instanceof h?this._data:new i(this._data)}};for(var u=["asText","asBinary","asNodeBuffer","asUint8Array","asArrayBuffer"],l=function(){throw new Error("This method has been removed in JSZip 3.0, please check the upgrade guide.")},f=0;f<u.length;f++)n.prototype[u[f]]=l;t.exports=n},{"./compressedObject":2,"./stream/DataWorker":27,"./stream/GenericWorker":28,"./stream/StreamHelper":29,"./utf8":31}],36:[function(e,l,t){(function(t){"use strict";var r,n,e=t.MutationObserver||t.WebKitMutationObserver;if(e){var i=0,s=new e(u),a=t.document.createTextNode("");s.observe(a,{characterData:!0}),r=function(){a.data=i=++i%2}}else if(t.setImmediate||void 0===t.MessageChannel)r="document"in t&&"onreadystatechange"in t.document.createElement("script")?function(){var e=t.document.createElement("script");e.onreadystatechange=function(){u(),e.onreadystatechange=null,e.parentNode.removeChild(e),e=null},t.document.documentElement.appendChild(e)}:function(){setTimeout(u,0)};else{var o=new t.MessageChannel;o.port1.onmessage=u,r=function(){o.port2.postMessage(0)}}var h=[];function u(){var e,t;n=!0;for(var r=h.length;r;){for(t=h,h=[],e=-1;++e<r;)t[e]();r=h.length}n=!1}l.exports=function(e){1!==h.push(e)||n||r()}}).call(this,"undefined"!=typeof global?global:"undefined"!=typeof self?self:"undefined"!=typeof window?window:{})},{}],37:[function(e,t,r){"use strict";var i=e("immediate");function u(){}var l={},s=["REJECTED"],a=["FULFILLED"],n=["PENDING"];function o(e){if("function"!=typeof e)throw new TypeError("resolver must be a function");this.state=n,this.queue=[],this.outcome=void 0,e!==u&&d(this,e)}function h(e,t,r){this.promise=e,"function"==typeof t&&(this.onFulfilled=t,this.callFulfilled=this.otherCallFulfilled),"function"==typeof r&&(this.onRejected=r,this.callRejected=this.otherCallRejected)}function f(t,r,n){i(function(){var e;try{e=r(n)}catch(e){return l.reject(t,e)}e===t?l.reject(t,new TypeError("Cannot resolve promise with itself")):l.resolve(t,e)})}function c(e){var t=e&&e.then;if(e&&("object"==typeof e||"function"==typeof e)&&"function"==typeof t)return function(){t.apply(e,arguments)}}function d(t,e){var r=!1;function n(e){r||(r=!0,l.reject(t,e))}function i(e){r||(r=!0,l.resolve(t,e))}var s=p(function(){e(i,n)});"error"===s.status&&n(s.value)}function p(e,t){var r={};try{r.value=e(t),r.status="success"}catch(e){r.status="error",r.value=e}return r}(t.exports=o).prototype.finally=function(t){if("function"!=typeof t)return this;var r=this.constructor;return this.then(function(e){return r.resolve(t()).then(function(){return e})},function(e){return r.resolve(t()).then(function(){throw e})})},o.prototype.catch=function(e){return this.then(null,e)},o.prototype.then=function(e,t){if("function"!=typeof e&&this.state===a||"function"!=typeof t&&this.state===s)return this;var r=new this.constructor(u);this.state!==n?f(r,this.state===a?e:t,this.outcome):this.queue.push(new h(r,e,t));return r},h.prototype.callFulfilled=function(e){l.resolve(this.promise,e)},h.prototype.otherCallFulfilled=function(e){f(this.promise,this.onFulfilled,e)},h.prototype.callRejected=function(e){l.reject(this.promise,e)},h.prototype.otherCallRejected=function(e){f(this.promise,this.onRejected,e)},l.resolve=function(e,t){var r=p(c,t);if("error"===r.status)return l.reject(e,r.value);var n=r.value;if(n)d(e,n);else{e.state=a,e.outcome=t;for(var i=-1,s=e.queue.length;++i<s;)e.queue[i].callFulfilled(t)}return e},l.reject=function(e,t){e.state=s,e.outcome=t;for(var r=-1,n=e.queue.length;++r<n;)e.queue[r].callRejected(t);return e},o.resolve=function(e){if(e instanceof this)return e;return l.resolve(new this(u),e)},o.reject=function(e){var t=new this(u);return l.reject(t,e)},o.all=function(e){var r=this;if("[object Array]"!==Object.prototype.toString.call(e))return this.reject(new TypeError("must be an array"));var n=e.length,i=!1;if(!n)return this.resolve([]);var s=new Array(n),a=0,t=-1,o=new this(u);for(;++t<n;)h(e[t],t);return o;function h(e,t){r.resolve(e).then(function(e){s[t]=e,++a!==n||i||(i=!0,l.resolve(o,s))},function(e){i||(i=!0,l.reject(o,e))})}},o.race=function(e){var t=this;if("[object Array]"!==Object.prototype.toString.call(e))return this.reject(new TypeError("must be an array"));var r=e.length,n=!1;if(!r)return this.resolve([]);var i=-1,s=new this(u);for(;++i<r;)a=e[i],t.resolve(a).then(function(e){n||(n=!0,l.resolve(s,e))},function(e){n||(n=!0,l.reject(s,e))});var a;return s}},{immediate:36}],38:[function(e,t,r){"use strict";var n={};(0,e("./lib/utils/common").assign)(n,e("./lib/deflate"),e("./lib/inflate"),e("./lib/zlib/constants")),t.exports=n},{"./lib/deflate":39,"./lib/inflate":40,"./lib/utils/common":41,"./lib/zlib/constants":44}],39:[function(e,t,r){"use strict";var a=e("./zlib/deflate"),o=e("./utils/common"),h=e("./utils/strings"),i=e("./zlib/messages"),s=e("./zlib/zstream"),u=Object.prototype.toString,l=0,f=-1,c=0,d=8;function p(e){if(!(this instanceof p))return new p(e);this.options=o.assign({level:f,method:d,chunkSize:16384,windowBits:15,memLevel:8,strategy:c,to:""},e||{});var t=this.options;t.raw&&0<t.windowBits?t.windowBits=-t.windowBits:t.gzip&&0<t.windowBits&&t.windowBits<16&&(t.windowBits+=16),this.err=0,this.msg="",this.ended=!1,this.chunks=[],this.strm=new s,this.strm.avail_out=0;var r=a.deflateInit2(this.strm,t.level,t.method,t.windowBits,t.memLevel,t.strategy);if(r!==l)throw new Error(i[r]);if(t.header&&a.deflateSetHeader(this.strm,t.header),t.dictionary){var n;if(n="string"==typeof t.dictionary?h.string2buf(t.dictionary):"[object ArrayBuffer]"===u.call(t.dictionary)?new Uint8Array(t.dictionary):t.dictionary,(r=a.deflateSetDictionary(this.strm,n))!==l)throw new Error(i[r]);this._dict_set=!0}}function n(e,t){var r=new p(t);if(r.push(e,!0),r.err)throw r.msg||i[r.err];return r.result}p.prototype.push=function(e,t){var r,n,i=this.strm,s=this.options.chunkSize;if(this.ended)return!1;n=t===~~t?t:!0===t?4:0,"string"==typeof e?i.input=h.string2buf(e):"[object ArrayBuffer]"===u.call(e)?i.input=new Uint8Array(e):i.input=e,i.next_in=0,i.avail_in=i.input.length;do{if(0===i.avail_out&&(i.output=new o.Buf8(s),i.next_out=0,i.avail_out=s),1!==(r=a.deflate(i,n))&&r!==l)return this.onEnd(r),!(this.ended=!0);0!==i.avail_out&&(0!==i.avail_in||4!==n&&2!==n)||("string"===this.options.to?this.onData(h.buf2binstring(o.shrinkBuf(i.output,i.next_out))):this.onData(o.shrinkBuf(i.output,i.next_out)))}while((0<i.avail_in||0===i.avail_out)&&1!==r);return 4===n?(r=a.deflateEnd(this.strm),this.onEnd(r),this.ended=!0,r===l):2!==n||(this.onEnd(l),!(i.avail_out=0))},p.prototype.onData=function(e){this.chunks.push(e)},p.prototype.onEnd=function(e){e===l&&("string"===this.options.to?this.result=this.chunks.join(""):this.result=o.flattenChunks(this.chunks)),this.chunks=[],this.err=e,this.msg=this.strm.msg},r.Deflate=p,r.deflate=n,r.deflateRaw=function(e,t){return(t=t||{}).raw=!0,n(e,t)},r.gzip=function(e,t){return(t=t||{}).gzip=!0,n(e,t)}},{"./utils/common":41,"./utils/strings":42,"./zlib/deflate":46,"./zlib/messages":51,"./zlib/zstream":53}],40:[function(e,t,r){"use strict";var c=e("./zlib/inflate"),d=e("./utils/common"),p=e("./utils/strings"),m=e("./zlib/constants"),n=e("./zlib/messages"),i=e("./zlib/zstream"),s=e("./zlib/gzheader"),_=Object.prototype.toString;function a(e){if(!(this instanceof a))return new a(e);this.options=d.assign({chunkSize:16384,windowBits:0,to:""},e||{});var t=this.options;t.raw&&0<=t.windowBits&&t.windowBits<16&&(t.windowBits=-t.windowBits,0===t.windowBits&&(t.windowBits=-15)),!(0<=t.windowBits&&t.windowBits<16)||e&&e.windowBits||(t.windowBits+=32),15<t.windowBits&&t.windowBits<48&&0==(15&t.windowBits)&&(t.windowBits|=15),this.err=0,this.msg="",this.ended=!1,this.chunks=[],this.strm=new i,this.strm.avail_out=0;var r=c.inflateInit2(this.strm,t.windowBits);if(r!==m.Z_OK)throw new Error(n[r]);this.header=new s,c.inflateGetHeader(this.strm,this.header)}function o(e,t){var r=new a(t);if(r.push(e,!0),r.err)throw r.msg||n[r.err];return r.result}a.prototype.push=function(e,t){var r,n,i,s,a,o,h=this.strm,u=this.options.chunkSize,l=this.options.dictionary,f=!1;if(this.ended)return!1;n=t===~~t?t:!0===t?m.Z_FINISH:m.Z_NO_FLUSH,"string"==typeof e?h.input=p.binstring2buf(e):"[object ArrayBuffer]"===_.call(e)?h.input=new Uint8Array(e):h.input=e,h.next_in=0,h.avail_in=h.input.length;do{if(0===h.avail_out&&(h.output=new d.Buf8(u),h.next_out=0,h.avail_out=u),(r=c.inflate(h,m.Z_NO_FLUSH))===m.Z_NEED_DICT&&l&&(o="string"==typeof l?p.string2buf(l):"[object ArrayBuffer]"===_.call(l)?new Uint8Array(l):l,r=c.inflateSetDictionary(this.strm,o)),r===m.Z_BUF_ERROR&&!0===f&&(r=m.Z_OK,f=!1),r!==m.Z_STREAM_END&&r!==m.Z_OK)return this.onEnd(r),!(this.ended=!0);h.next_out&&(0!==h.avail_out&&r!==m.Z_STREAM_END&&(0!==h.avail_in||n!==m.Z_FINISH&&n!==m.Z_SYNC_FLUSH)||("string"===this.options.to?(i=p.utf8border(h.output,h.next_out),s=h.next_out-i,a=p.buf2string(h.output,i),h.next_out=s,h.avail_out=u-s,s&&d.arraySet(h.output,h.output,i,s,0),this.onData(a)):this.onData(d.shrinkBuf(h.output,h.next_out)))),0===h.avail_in&&0===h.avail_out&&(f=!0)}while((0<h.avail_in||0===h.avail_out)&&r!==m.Z_STREAM_END);return r===m.Z_STREAM_END&&(n=m.Z_FINISH),n===m.Z_FINISH?(r=c.inflateEnd(this.strm),this.onEnd(r),this.ended=!0,r===m.Z_OK):n!==m.Z_SYNC_FLUSH||(this.onEnd(m.Z_OK),!(h.avail_out=0))},a.prototype.onData=function(e){this.chunks.push(e)},a.prototype.onEnd=function(e){e===m.Z_OK&&("string"===this.options.to?this.result=this.chunks.join(""):this.result=d.flattenChunks(this.chunks)),this.chunks=[],this.err=e,this.msg=this.strm.msg},r.Inflate=a,r.inflate=o,r.inflateRaw=function(e,t){return(t=t||{}).raw=!0,o(e,t)},r.ungzip=o},{"./utils/common":41,"./utils/strings":42,"./zlib/constants":44,"./zlib/gzheader":47,"./zlib/inflate":49,"./zlib/messages":51,"./zlib/zstream":53}],41:[function(e,t,r){"use strict";var n="undefined"!=typeof Uint8Array&&"undefined"!=typeof Uint16Array&&"undefined"!=typeof Int32Array;r.assign=function(e){for(var t=Array.prototype.slice.call(arguments,1);t.length;){var r=t.shift();if(r){if("object"!=typeof r)throw new TypeError(r+"must be non-object");for(var n in r)r.hasOwnProperty(n)&&(e[n]=r[n])}}return e},r.shrinkBuf=function(e,t){return e.length===t?e:e.subarray?e.subarray(0,t):(e.length=t,e)};var i={arraySet:function(e,t,r,n,i){if(t.subarray&&e.subarray)e.set(t.subarray(r,r+n),i);else for(var s=0;s<n;s++)e[i+s]=t[r+s]},flattenChunks:function(e){var t,r,n,i,s,a;for(t=n=0,r=e.length;t<r;t++)n+=e[t].length;for(a=new Uint8Array(n),t=i=0,r=e.length;t<r;t++)s=e[t],a.set(s,i),i+=s.length;return a}},s={arraySet:function(e,t,r,n,i){for(var s=0;s<n;s++)e[i+s]=t[r+s]},flattenChunks:function(e){return[].concat.apply([],e)}};r.setTyped=function(e){e?(r.Buf8=Uint8Array,r.Buf16=Uint16Array,r.Buf32=Int32Array,r.assign(r,i)):(r.Buf8=Array,r.Buf16=Array,r.Buf32=Array,r.assign(r,s))},r.setTyped(n)},{}],42:[function(e,t,r){"use strict";var h=e("./common"),i=!0,s=!0;try{String.fromCharCode.apply(null,[0])}catch(e){i=!1}try{String.fromCharCode.apply(null,new Uint8Array(1))}catch(e){s=!1}for(var u=new h.Buf8(256),n=0;n<256;n++)u[n]=252<=n?6:248<=n?5:240<=n?4:224<=n?3:192<=n?2:1;function l(e,t){if(t<65537&&(e.subarray&&s||!e.subarray&&i))return String.fromCharCode.apply(null,h.shrinkBuf(e,t));for(var r="",n=0;n<t;n++)r+=String.fromCharCode(e[n]);return r}u[254]=u[254]=1,r.string2buf=function(e){var t,r,n,i,s,a=e.length,o=0;for(i=0;i<a;i++)55296==(64512&(r=e.charCodeAt(i)))&&i+1<a&&56320==(64512&(n=e.charCodeAt(i+1)))&&(r=65536+(r-55296<<10)+(n-56320),i++),o+=r<128?1:r<2048?2:r<65536?3:4;for(t=new h.Buf8(o),i=s=0;s<o;i++)55296==(64512&(r=e.charCodeAt(i)))&&i+1<a&&56320==(64512&(n=e.charCodeAt(i+1)))&&(r=65536+(r-55296<<10)+(n-56320),i++),r<128?t[s++]=r:(r<2048?t[s++]=192|r>>>6:(r<65536?t[s++]=224|r>>>12:(t[s++]=240|r>>>18,t[s++]=128|r>>>12&63),t[s++]=128|r>>>6&63),t[s++]=128|63&r);return t},r.buf2binstring=function(e){return l(e,e.length)},r.binstring2buf=function(e){for(var t=new h.Buf8(e.length),r=0,n=t.length;r<n;r++)t[r]=e.charCodeAt(r);return t},r.buf2string=function(e,t){var r,n,i,s,a=t||e.length,o=new Array(2*a);for(r=n=0;r<a;)if((i=e[r++])<128)o[n++]=i;else if(4<(s=u[i]))o[n++]=65533,r+=s-1;else{for(i&=2===s?31:3===s?15:7;1<s&&r<a;)i=i<<6|63&e[r++],s--;1<s?o[n++]=65533:i<65536?o[n++]=i:(i-=65536,o[n++]=55296|i>>10&1023,o[n++]=56320|1023&i)}return l(o,n)},r.utf8border=function(e,t){var r;for((t=t||e.length)>e.length&&(t=e.length),r=t-1;0<=r&&128==(192&e[r]);)r--;return r<0?t:0===r?t:r+u[e[r]]>t?r:t}},{"./common":41}],43:[function(e,t,r){"use strict";t.exports=function(e,t,r,n){for(var i=65535&e|0,s=e>>>16&65535|0,a=0;0!==r;){for(r-=a=2e3<r?2e3:r;s=s+(i=i+t[n++]|0)|0,--a;);i%=65521,s%=65521}return i|s<<16|0}},{}],44:[function(e,t,r){"use strict";t.exports={Z_NO_FLUSH:0,Z_PARTIAL_FLUSH:1,Z_SYNC_FLUSH:2,Z_FULL_FLUSH:3,Z_FINISH:4,Z_BLOCK:5,Z_TREES:6,Z_OK:0,Z_STREAM_END:1,Z_NEED_DICT:2,Z_ERRNO:-1,Z_STREAM_ERROR:-2,Z_DATA_ERROR:-3,Z_BUF_ERROR:-5,Z_NO_COMPRESSION:0,Z_BEST_SPEED:1,Z_BEST_COMPRESSION:9,Z_DEFAULT_COMPRESSION:-1,Z_FILTERED:1,Z_HUFFMAN_ONLY:2,Z_RLE:3,Z_FIXED:4,Z_DEFAULT_STRATEGY:0,Z_BINARY:0,Z_TEXT:1,Z_UNKNOWN:2,Z_DEFLATED:8}},{}],45:[function(e,t,r){"use strict";var o=function(){for(var e,t=[],r=0;r<256;r++){e=r;for(var n=0;n<8;n++)e=1&e?3988292384^e>>>1:e>>>1;t[r]=e}return t}();t.exports=function(e,t,r,n){var i=o,s=n+r;e^=-1;for(var a=n;a<s;a++)e=e>>>8^i[255&(e^t[a])];return-1^e}},{}],46:[function(e,t,r){"use strict";var h,c=e("../utils/common"),u=e("./trees"),d=e("./adler32"),p=e("./crc32"),n=e("./messages"),l=0,f=4,m=0,_=-2,g=-1,b=4,i=2,v=8,y=9,s=286,a=30,o=19,w=2*s+1,k=15,x=3,S=258,z=S+x+1,C=42,E=113,A=1,I=2,O=3,B=4;function R(e,t){return e.msg=n[t],t}function T(e){return(e<<1)-(4<e?9:0)}function D(e){for(var t=e.length;0<=--t;)e[t]=0}function F(e){var t=e.state,r=t.pending;r>e.avail_out&&(r=e.avail_out),0!==r&&(c.arraySet(e.output,t.pending_buf,t.pending_out,r,e.next_out),e.next_out+=r,t.pending_out+=r,e.total_out+=r,e.avail_out-=r,t.pending-=r,0===t.pending&&(t.pending_out=0))}function N(e,t){u._tr_flush_block(e,0<=e.block_start?e.block_start:-1,e.strstart-e.block_start,t),e.block_start=e.strstart,F(e.strm)}function U(e,t){e.pending_buf[e.pending++]=t}function P(e,t){e.pending_buf[e.pending++]=t>>>8&255,e.pending_buf[e.pending++]=255&t}function L(e,t){var r,n,i=e.max_chain_length,s=e.strstart,a=e.prev_length,o=e.nice_match,h=e.strstart>e.w_size-z?e.strstart-(e.w_size-z):0,u=e.window,l=e.w_mask,f=e.prev,c=e.strstart+S,d=u[s+a-1],p=u[s+a];e.prev_length>=e.good_match&&(i>>=2),o>e.lookahead&&(o=e.lookahead);do{if(u[(r=t)+a]===p&&u[r+a-1]===d&&u[r]===u[s]&&u[++r]===u[s+1]){s+=2,r++;do{}while(u[++s]===u[++r]&&u[++s]===u[++r]&&u[++s]===u[++r]&&u[++s]===u[++r]&&u[++s]===u[++r]&&u[++s]===u[++r]&&u[++s]===u[++r]&&u[++s]===u[++r]&&s<c);if(n=S-(c-s),s=c-S,a<n){if(e.match_start=t,o<=(a=n))break;d=u[s+a-1],p=u[s+a]}}}while((t=f[t&l])>h&&0!=--i);return a<=e.lookahead?a:e.lookahead}function j(e){var t,r,n,i,s,a,o,h,u,l,f=e.w_size;do{if(i=e.window_size-e.lookahead-e.strstart,e.strstart>=f+(f-z)){for(c.arraySet(e.window,e.window,f,f,0),e.match_start-=f,e.strstart-=f,e.block_start-=f,t=r=e.hash_size;n=e.head[--t],e.head[t]=f<=n?n-f:0,--r;);for(t=r=f;n=e.prev[--t],e.prev[t]=f<=n?n-f:0,--r;);i+=f}if(0===e.strm.avail_in)break;if(a=e.strm,o=e.window,h=e.strstart+e.lookahead,u=i,l=void 0,l=a.avail_in,u<l&&(l=u),r=0===l?0:(a.avail_in-=l,c.arraySet(o,a.input,a.next_in,l,h),1===a.state.wrap?a.adler=d(a.adler,o,l,h):2===a.state.wrap&&(a.adler=p(a.adler,o,l,h)),a.next_in+=l,a.total_in+=l,l),e.lookahead+=r,e.lookahead+e.insert>=x)for(s=e.strstart-e.insert,e.ins_h=e.window[s],e.ins_h=(e.ins_h<<e.hash_shift^e.window[s+1])&e.hash_mask;e.insert&&(e.ins_h=(e.ins_h<<e.hash_shift^e.window[s+x-1])&e.hash_mask,e.prev[s&e.w_mask]=e.head[e.ins_h],e.head[e.ins_h]=s,s++,e.insert--,!(e.lookahead+e.insert<x)););}while(e.lookahead<z&&0!==e.strm.avail_in)}function Z(e,t){for(var r,n;;){if(e.lookahead<z){if(j(e),e.lookahead<z&&t===l)return A;if(0===e.lookahead)break}if(r=0,e.lookahead>=x&&(e.ins_h=(e.ins_h<<e.hash_shift^e.window[e.strstart+x-1])&e.hash_mask,r=e.prev[e.strstart&e.w_mask]=e.head[e.ins_h],e.head[e.ins_h]=e.strstart),0!==r&&e.strstart-r<=e.w_size-z&&(e.match_length=L(e,r)),e.match_length>=x)if(n=u._tr_tally(e,e.strstart-e.match_start,e.match_length-x),e.lookahead-=e.match_length,e.match_length<=e.max_lazy_match&&e.lookahead>=x){for(e.match_length--;e.strstart++,e.ins_h=(e.ins_h<<e.hash_shift^e.window[e.strstart+x-1])&e.hash_mask,r=e.prev[e.strstart&e.w_mask]=e.head[e.ins_h],e.head[e.ins_h]=e.strstart,0!=--e.match_length;);e.strstart++}else e.strstart+=e.match_length,e.match_length=0,e.ins_h=e.window[e.strstart],e.ins_h=(e.ins_h<<e.hash_shift^e.window[e.strstart+1])&e.hash_mask;else n=u._tr_tally(e,0,e.window[e.strstart]),e.lookahead--,e.strstart++;if(n&&(N(e,!1),0===e.strm.avail_out))return A}return e.insert=e.strstart<x-1?e.strstart:x-1,t===f?(N(e,!0),0===e.strm.avail_out?O:B):e.last_lit&&(N(e,!1),0===e.strm.avail_out)?A:I}function W(e,t){for(var r,n,i;;){if(e.lookahead<z){if(j(e),e.lookahead<z&&t===l)return A;if(0===e.lookahead)break}if(r=0,e.lookahead>=x&&(e.ins_h=(e.ins_h<<e.hash_shift^e.window[e.strstart+x-1])&e.hash_mask,r=e.prev[e.strstart&e.w_mask]=e.head[e.ins_h],e.head[e.ins_h]=e.strstart),e.prev_length=e.match_length,e.prev_match=e.match_start,e.match_length=x-1,0!==r&&e.prev_length<e.max_lazy_match&&e.strstart-r<=e.w_size-z&&(e.match_length=L(e,r),e.match_length<=5&&(1===e.strategy||e.match_length===x&&4096<e.strstart-e.match_start)&&(e.match_length=x-1)),e.prev_length>=x&&e.match_length<=e.prev_length){for(i=e.strstart+e.lookahead-x,n=u._tr_tally(e,e.strstart-1-e.prev_match,e.prev_length-x),e.lookahead-=e.prev_length-1,e.prev_length-=2;++e.strstart<=i&&(e.ins_h=(e.ins_h<<e.hash_shift^e.window[e.strstart+x-1])&e.hash_mask,r=e.prev[e.strstart&e.w_mask]=e.head[e.ins_h],e.head[e.ins_h]=e.strstart),0!=--e.prev_length;);if(e.match_available=0,e.match_length=x-1,e.strstart++,n&&(N(e,!1),0===e.strm.avail_out))return A}else if(e.match_available){if((n=u._tr_tally(e,0,e.window[e.strstart-1]))&&N(e,!1),e.strstart++,e.lookahead--,0===e.strm.avail_out)return A}else e.match_available=1,e.strstart++,e.lookahead--}return e.match_available&&(n=u._tr_tally(e,0,e.window[e.strstart-1]),e.match_available=0),e.insert=e.strstart<x-1?e.strstart:x-1,t===f?(N(e,!0),0===e.strm.avail_out?O:B):e.last_lit&&(N(e,!1),0===e.strm.avail_out)?A:I}function M(e,t,r,n,i){this.good_length=e,this.max_lazy=t,this.nice_length=r,this.max_chain=n,this.func=i}function H(){this.strm=null,this.status=0,this.pending_buf=null,this.pending_buf_size=0,this.pending_out=0,this.pending=0,this.wrap=0,this.gzhead=null,this.gzindex=0,this.method=v,this.last_flush=-1,this.w_size=0,this.w_bits=0,this.w_mask=0,this.window=null,this.window_size=0,this.prev=null,this.head=null,this.ins_h=0,this.hash_size=0,this.hash_bits=0,this.hash_mask=0,this.hash_shift=0,this.block_start=0,this.match_length=0,this.prev_match=0,this.match_available=0,this.strstart=0,this.match_start=0,this.lookahead=0,this.prev_length=0,this.max_chain_length=0,this.max_lazy_match=0,this.level=0,this.strategy=0,this.good_match=0,this.nice_match=0,this.dyn_ltree=new c.Buf16(2*w),this.dyn_dtree=new c.Buf16(2*(2*a+1)),this.bl_tree=new c.Buf16(2*(2*o+1)),D(this.dyn_ltree),D(this.dyn_dtree),D(this.bl_tree),this.l_desc=null,this.d_desc=null,this.bl_desc=null,this.bl_count=new c.Buf16(k+1),this.heap=new c.Buf16(2*s+1),D(this.heap),this.heap_len=0,this.heap_max=0,this.depth=new c.Buf16(2*s+1),D(this.depth),this.l_buf=0,this.lit_bufsize=0,this.last_lit=0,this.d_buf=0,this.opt_len=0,this.static_len=0,this.matches=0,this.insert=0,this.bi_buf=0,this.bi_valid=0}function G(e){var t;return e&&e.state?(e.total_in=e.total_out=0,e.data_type=i,(t=e.state).pending=0,t.pending_out=0,t.wrap<0&&(t.wrap=-t.wrap),t.status=t.wrap?C:E,e.adler=2===t.wrap?0:1,t.last_flush=l,u._tr_init(t),m):R(e,_)}function K(e){var t=G(e);return t===m&&function(e){e.window_size=2*e.w_size,D(e.head),e.max_lazy_match=h[e.level].max_lazy,e.good_match=h[e.level].good_length,e.nice_match=h[e.level].nice_length,e.max_chain_length=h[e.level].max_chain,e.strstart=0,e.block_start=0,e.lookahead=0,e.insert=0,e.match_length=e.prev_length=x-1,e.match_available=0,e.ins_h=0}(e.state),t}function Y(e,t,r,n,i,s){if(!e)return _;var a=1;if(t===g&&(t=6),n<0?(a=0,n=-n):15<n&&(a=2,n-=16),i<1||y<i||r!==v||n<8||15<n||t<0||9<t||s<0||b<s)return R(e,_);8===n&&(n=9);var o=new H;return(e.state=o).strm=e,o.wrap=a,o.gzhead=null,o.w_bits=n,o.w_size=1<<o.w_bits,o.w_mask=o.w_size-1,o.hash_bits=i+7,o.hash_size=1<<o.hash_bits,o.hash_mask=o.hash_size-1,o.hash_shift=~~((o.hash_bits+x-1)/x),o.window=new c.Buf8(2*o.w_size),o.head=new c.Buf16(o.hash_size),o.prev=new c.Buf16(o.w_size),o.lit_bufsize=1<<i+6,o.pending_buf_size=4*o.lit_bufsize,o.pending_buf=new c.Buf8(o.pending_buf_size),o.d_buf=1*o.lit_bufsize,o.l_buf=3*o.lit_bufsize,o.level=t,o.strategy=s,o.method=r,K(e)}h=[new M(0,0,0,0,function(e,t){var r=65535;for(r>e.pending_buf_size-5&&(r=e.pending_buf_size-5);;){if(e.lookahead<=1){if(j(e),0===e.lookahead&&t===l)return A;if(0===e.lookahead)break}e.strstart+=e.lookahead,e.lookahead=0;var n=e.block_start+r;if((0===e.strstart||e.strstart>=n)&&(e.lookahead=e.strstart-n,e.strstart=n,N(e,!1),0===e.strm.avail_out))return A;if(e.strstart-e.block_start>=e.w_size-z&&(N(e,!1),0===e.strm.avail_out))return A}return e.insert=0,t===f?(N(e,!0),0===e.strm.avail_out?O:B):(e.strstart>e.block_start&&(N(e,!1),e.strm.avail_out),A)}),new M(4,4,8,4,Z),new M(4,5,16,8,Z),new M(4,6,32,32,Z),new M(4,4,16,16,W),new M(8,16,32,32,W),new M(8,16,128,128,W),new M(8,32,128,256,W),new M(32,128,258,1024,W),new M(32,258,258,4096,W)],r.deflateInit=function(e,t){return Y(e,t,v,15,8,0)},r.deflateInit2=Y,r.deflateReset=K,r.deflateResetKeep=G,r.deflateSetHeader=function(e,t){return e&&e.state?2!==e.state.wrap?_:(e.state.gzhead=t,m):_},r.deflate=function(e,t){var r,n,i,s;if(!e||!e.state||5<t||t<0)return e?R(e,_):_;if(n=e.state,!e.output||!e.input&&0!==e.avail_in||666===n.status&&t!==f)return R(e,0===e.avail_out?-5:_);if(n.strm=e,r=n.last_flush,n.last_flush=t,n.status===C)if(2===n.wrap)e.adler=0,U(n,31),U(n,139),U(n,8),n.gzhead?(U(n,(n.gzhead.text?1:0)+(n.gzhead.hcrc?2:0)+(n.gzhead.extra?4:0)+(n.gzhead.name?8:0)+(n.gzhead.comment?16:0)),U(n,255&n.gzhead.time),U(n,n.gzhead.time>>8&255),U(n,n.gzhead.time>>16&255),U(n,n.gzhead.time>>24&255),U(n,9===n.level?2:2<=n.strategy||n.level<2?4:0),U(n,255&n.gzhead.os),n.gzhead.extra&&n.gzhead.extra.length&&(U(n,255&n.gzhead.extra.length),U(n,n.gzhead.extra.length>>8&255)),n.gzhead.hcrc&&(e.adler=p(e.adler,n.pending_buf,n.pending,0)),n.gzindex=0,n.status=69):(U(n,0),U(n,0),U(n,0),U(n,0),U(n,0),U(n,9===n.level?2:2<=n.strategy||n.level<2?4:0),U(n,3),n.status=E);else{var a=v+(n.w_bits-8<<4)<<8;a|=(2<=n.strategy||n.level<2?0:n.level<6?1:6===n.level?2:3)<<6,0!==n.strstart&&(a|=32),a+=31-a%31,n.status=E,P(n,a),0!==n.strstart&&(P(n,e.adler>>>16),P(n,65535&e.adler)),e.adler=1}if(69===n.status)if(n.gzhead.extra){for(i=n.pending;n.gzindex<(65535&n.gzhead.extra.length)&&(n.pending!==n.pending_buf_size||(n.gzhead.hcrc&&n.pending>i&&(e.adler=p(e.adler,n.pending_buf,n.pending-i,i)),F(e),i=n.pending,n.pending!==n.pending_buf_size));)U(n,255&n.gzhead.extra[n.gzindex]),n.gzindex++;n.gzhead.hcrc&&n.pending>i&&(e.adler=p(e.adler,n.pending_buf,n.pending-i,i)),n.gzindex===n.gzhead.extra.length&&(n.gzindex=0,n.status=73)}else n.status=73;if(73===n.status)if(n.gzhead.name){i=n.pending;do{if(n.pending===n.pending_buf_size&&(n.gzhead.hcrc&&n.pending>i&&(e.adler=p(e.adler,n.pending_buf,n.pending-i,i)),F(e),i=n.pending,n.pending===n.pending_buf_size)){s=1;break}s=n.gzindex<n.gzhead.name.length?255&n.gzhead.name.charCodeAt(n.gzindex++):0,U(n,s)}while(0!==s);n.gzhead.hcrc&&n.pending>i&&(e.adler=p(e.adler,n.pending_buf,n.pending-i,i)),0===s&&(n.gzindex=0,n.status=91)}else n.status=91;if(91===n.status)if(n.gzhead.comment){i=n.pending;do{if(n.pending===n.pending_buf_size&&(n.gzhead.hcrc&&n.pending>i&&(e.adler=p(e.adler,n.pending_buf,n.pending-i,i)),F(e),i=n.pending,n.pending===n.pending_buf_size)){s=1;break}s=n.gzindex<n.gzhead.comment.length?255&n.gzhead.comment.charCodeAt(n.gzindex++):0,U(n,s)}while(0!==s);n.gzhead.hcrc&&n.pending>i&&(e.adler=p(e.adler,n.pending_buf,n.pending-i,i)),0===s&&(n.status=103)}else n.status=103;if(103===n.status&&(n.gzhead.hcrc?(n.pending+2>n.pending_buf_size&&F(e),n.pending+2<=n.pending_buf_size&&(U(n,255&e.adler),U(n,e.adler>>8&255),e.adler=0,n.status=E)):n.status=E),0!==n.pending){if(F(e),0===e.avail_out)return n.last_flush=-1,m}else if(0===e.avail_in&&T(t)<=T(r)&&t!==f)return R(e,-5);if(666===n.status&&0!==e.avail_in)return R(e,-5);if(0!==e.avail_in||0!==n.lookahead||t!==l&&666!==n.status){var o=2===n.strategy?function(e,t){for(var r;;){if(0===e.lookahead&&(j(e),0===e.lookahead)){if(t===l)return A;break}if(e.match_length=0,r=u._tr_tally(e,0,e.window[e.strstart]),e.lookahead--,e.strstart++,r&&(N(e,!1),0===e.strm.avail_out))return A}return e.insert=0,t===f?(N(e,!0),0===e.strm.avail_out?O:B):e.last_lit&&(N(e,!1),0===e.strm.avail_out)?A:I}(n,t):3===n.strategy?function(e,t){for(var r,n,i,s,a=e.window;;){if(e.lookahead<=S){if(j(e),e.lookahead<=S&&t===l)return A;if(0===e.lookahead)break}if(e.match_length=0,e.lookahead>=x&&0<e.strstart&&(n=a[i=e.strstart-1])===a[++i]&&n===a[++i]&&n===a[++i]){s=e.strstart+S;do{}while(n===a[++i]&&n===a[++i]&&n===a[++i]&&n===a[++i]&&n===a[++i]&&n===a[++i]&&n===a[++i]&&n===a[++i]&&i<s);e.match_length=S-(s-i),e.match_length>e.lookahead&&(e.match_length=e.lookahead)}if(e.match_length>=x?(r=u._tr_tally(e,1,e.match_length-x),e.lookahead-=e.match_length,e.strstart+=e.match_length,e.match_length=0):(r=u._tr_tally(e,0,e.window[e.strstart]),e.lookahead--,e.strstart++),r&&(N(e,!1),0===e.strm.avail_out))return A}return e.insert=0,t===f?(N(e,!0),0===e.strm.avail_out?O:B):e.last_lit&&(N(e,!1),0===e.strm.avail_out)?A:I}(n,t):h[n.level].func(n,t);if(o!==O&&o!==B||(n.status=666),o===A||o===O)return 0===e.avail_out&&(n.last_flush=-1),m;if(o===I&&(1===t?u._tr_align(n):5!==t&&(u._tr_stored_block(n,0,0,!1),3===t&&(D(n.head),0===n.lookahead&&(n.strstart=0,n.block_start=0,n.insert=0))),F(e),0===e.avail_out))return n.last_flush=-1,m}return t!==f?m:n.wrap<=0?1:(2===n.wrap?(U(n,255&e.adler),U(n,e.adler>>8&255),U(n,e.adler>>16&255),U(n,e.adler>>24&255),U(n,255&e.total_in),U(n,e.total_in>>8&255),U(n,e.total_in>>16&255),U(n,e.total_in>>24&255)):(P(n,e.adler>>>16),P(n,65535&e.adler)),F(e),0<n.wrap&&(n.wrap=-n.wrap),0!==n.pending?m:1)},r.deflateEnd=function(e){var t;return e&&e.state?(t=e.state.status)!==C&&69!==t&&73!==t&&91!==t&&103!==t&&t!==E&&666!==t?R(e,_):(e.state=null,t===E?R(e,-3):m):_},r.deflateSetDictionary=function(e,t){var r,n,i,s,a,o,h,u,l=t.length;if(!e||!e.state)return _;if(2===(s=(r=e.state).wrap)||1===s&&r.status!==C||r.lookahead)return _;for(1===s&&(e.adler=d(e.adler,t,l,0)),r.wrap=0,l>=r.w_size&&(0===s&&(D(r.head),r.strstart=0,r.block_start=0,r.insert=0),u=new c.Buf8(r.w_size),c.arraySet(u,t,l-r.w_size,r.w_size,0),t=u,l=r.w_size),a=e.avail_in,o=e.next_in,h=e.input,e.avail_in=l,e.next_in=0,e.input=t,j(r);r.lookahead>=x;){for(n=r.strstart,i=r.lookahead-(x-1);r.ins_h=(r.ins_h<<r.hash_shift^r.window[n+x-1])&r.hash_mask,r.prev[n&r.w_mask]=r.head[r.ins_h],r.head[r.ins_h]=n,n++,--i;);r.strstart=n,r.lookahead=x-1,j(r)}return r.strstart+=r.lookahead,r.block_start=r.strstart,r.insert=r.lookahead,r.lookahead=0,r.match_length=r.prev_length=x-1,r.match_available=0,e.next_in=o,e.input=h,e.avail_in=a,r.wrap=s,m},r.deflateInfo="pako deflate (from Nodeca project)"},{"../utils/common":41,"./adler32":43,"./crc32":45,"./messages":51,"./trees":52}],47:[function(e,t,r){"use strict";t.exports=function(){this.text=0,this.time=0,this.xflags=0,this.os=0,this.extra=null,this.extra_len=0,this.name="",this.comment="",this.hcrc=0,this.done=!1}},{}],48:[function(e,t,r){"use strict";t.exports=function(e,t){var r,n,i,s,a,o,h,u,l,f,c,d,p,m,_,g,b,v,y,w,k,x,S,z,C;r=e.state,n=e.next_in,z=e.input,i=n+(e.avail_in-5),s=e.next_out,C=e.output,a=s-(t-e.avail_out),o=s+(e.avail_out-257),h=r.dmax,u=r.wsize,l=r.whave,f=r.wnext,c=r.window,d=r.hold,p=r.bits,m=r.lencode,_=r.distcode,g=(1<<r.lenbits)-1,b=(1<<r.distbits)-1;e:do{p<15&&(d+=z[n++]<<p,p+=8,d+=z[n++]<<p,p+=8),v=m[d&g];t:for(;;){if(d>>>=y=v>>>24,p-=y,0===(y=v>>>16&255))C[s++]=65535&v;else{if(!(16&y)){if(0==(64&y)){v=m[(65535&v)+(d&(1<<y)-1)];continue t}if(32&y){r.mode=12;break e}e.msg="invalid literal/length code",r.mode=30;break e}w=65535&v,(y&=15)&&(p<y&&(d+=z[n++]<<p,p+=8),w+=d&(1<<y)-1,d>>>=y,p-=y),p<15&&(d+=z[n++]<<p,p+=8,d+=z[n++]<<p,p+=8),v=_[d&b];r:for(;;){if(d>>>=y=v>>>24,p-=y,!(16&(y=v>>>16&255))){if(0==(64&y)){v=_[(65535&v)+(d&(1<<y)-1)];continue r}e.msg="invalid distance code",r.mode=30;break e}if(k=65535&v,p<(y&=15)&&(d+=z[n++]<<p,(p+=8)<y&&(d+=z[n++]<<p,p+=8)),h<(k+=d&(1<<y)-1)){e.msg="invalid distance too far back",r.mode=30;break e}if(d>>>=y,p-=y,(y=s-a)<k){if(l<(y=k-y)&&r.sane){e.msg="invalid distance too far back",r.mode=30;break e}if(S=c,(x=0)===f){if(x+=u-y,y<w){for(w-=y;C[s++]=c[x++],--y;);x=s-k,S=C}}else if(f<y){if(x+=u+f-y,(y-=f)<w){for(w-=y;C[s++]=c[x++],--y;);if(x=0,f<w){for(w-=y=f;C[s++]=c[x++],--y;);x=s-k,S=C}}}else if(x+=f-y,y<w){for(w-=y;C[s++]=c[x++],--y;);x=s-k,S=C}for(;2<w;)C[s++]=S[x++],C[s++]=S[x++],C[s++]=S[x++],w-=3;w&&(C[s++]=S[x++],1<w&&(C[s++]=S[x++]))}else{for(x=s-k;C[s++]=C[x++],C[s++]=C[x++],C[s++]=C[x++],2<(w-=3););w&&(C[s++]=C[x++],1<w&&(C[s++]=C[x++]))}break}}break}}while(n<i&&s<o);n-=w=p>>3,d&=(1<<(p-=w<<3))-1,e.next_in=n,e.next_out=s,e.avail_in=n<i?i-n+5:5-(n-i),e.avail_out=s<o?o-s+257:257-(s-o),r.hold=d,r.bits=p}},{}],49:[function(e,t,r){"use strict";var I=e("../utils/common"),O=e("./adler32"),B=e("./crc32"),R=e("./inffast"),T=e("./inftrees"),D=1,F=2,N=0,U=-2,P=1,n=852,i=592;function L(e){return(e>>>24&255)+(e>>>8&65280)+((65280&e)<<8)+((255&e)<<24)}function s(){this.mode=0,this.last=!1,this.wrap=0,this.havedict=!1,this.flags=0,this.dmax=0,this.check=0,this.total=0,this.head=null,this.wbits=0,this.wsize=0,this.whave=0,this.wnext=0,this.window=null,this.hold=0,this.bits=0,this.length=0,this.offset=0,this.extra=0,this.lencode=null,this.distcode=null,this.lenbits=0,this.distbits=0,this.ncode=0,this.nlen=0,this.ndist=0,this.have=0,this.next=null,this.lens=new I.Buf16(320),this.work=new I.Buf16(288),this.lendyn=null,this.distdyn=null,this.sane=0,this.back=0,this.was=0}function a(e){var t;return e&&e.state?(t=e.state,e.total_in=e.total_out=t.total=0,e.msg="",t.wrap&&(e.adler=1&t.wrap),t.mode=P,t.last=0,t.havedict=0,t.dmax=32768,t.head=null,t.hold=0,t.bits=0,t.lencode=t.lendyn=new I.Buf32(n),t.distcode=t.distdyn=new I.Buf32(i),t.sane=1,t.back=-1,N):U}function o(e){var t;return e&&e.state?((t=e.state).wsize=0,t.whave=0,t.wnext=0,a(e)):U}function h(e,t){var r,n;return e&&e.state?(n=e.state,t<0?(r=0,t=-t):(r=1+(t>>4),t<48&&(t&=15)),t&&(t<8||15<t)?U:(null!==n.window&&n.wbits!==t&&(n.window=null),n.wrap=r,n.wbits=t,o(e))):U}function u(e,t){var r,n;return e?(n=new s,(e.state=n).window=null,(r=h(e,t))!==N&&(e.state=null),r):U}var l,f,c=!0;function j(e){if(c){var t;for(l=new I.Buf32(512),f=new I.Buf32(32),t=0;t<144;)e.lens[t++]=8;for(;t<256;)e.lens[t++]=9;for(;t<280;)e.lens[t++]=7;for(;t<288;)e.lens[t++]=8;for(T(D,e.lens,0,288,l,0,e.work,{bits:9}),t=0;t<32;)e.lens[t++]=5;T(F,e.lens,0,32,f,0,e.work,{bits:5}),c=!1}e.lencode=l,e.lenbits=9,e.distcode=f,e.distbits=5}function Z(e,t,r,n){var i,s=e.state;return null===s.window&&(s.wsize=1<<s.wbits,s.wnext=0,s.whave=0,s.window=new I.Buf8(s.wsize)),n>=s.wsize?(I.arraySet(s.window,t,r-s.wsize,s.wsize,0),s.wnext=0,s.whave=s.wsize):(n<(i=s.wsize-s.wnext)&&(i=n),I.arraySet(s.window,t,r-n,i,s.wnext),(n-=i)?(I.arraySet(s.window,t,r-n,n,0),s.wnext=n,s.whave=s.wsize):(s.wnext+=i,s.wnext===s.wsize&&(s.wnext=0),s.whave<s.wsize&&(s.whave+=i))),0}r.inflateReset=o,r.inflateReset2=h,r.inflateResetKeep=a,r.inflateInit=function(e){return u(e,15)},r.inflateInit2=u,r.inflate=function(e,t){var r,n,i,s,a,o,h,u,l,f,c,d,p,m,_,g,b,v,y,w,k,x,S,z,C=0,E=new I.Buf8(4),A=[16,17,18,0,8,7,9,6,10,5,11,4,12,3,13,2,14,1,15];if(!e||!e.state||!e.output||!e.input&&0!==e.avail_in)return U;12===(r=e.state).mode&&(r.mode=13),a=e.next_out,i=e.output,h=e.avail_out,s=e.next_in,n=e.input,o=e.avail_in,u=r.hold,l=r.bits,f=o,c=h,x=N;e:for(;;)switch(r.mode){case P:if(0===r.wrap){r.mode=13;break}for(;l<16;){if(0===o)break e;o--,u+=n[s++]<<l,l+=8}if(2&r.wrap&&35615===u){E[r.check=0]=255&u,E[1]=u>>>8&255,r.check=B(r.check,E,2,0),l=u=0,r.mode=2;break}if(r.flags=0,r.head&&(r.head.done=!1),!(1&r.wrap)||(((255&u)<<8)+(u>>8))%31){e.msg="incorrect header check",r.mode=30;break}if(8!=(15&u)){e.msg="unknown compression method",r.mode=30;break}if(l-=4,k=8+(15&(u>>>=4)),0===r.wbits)r.wbits=k;else if(k>r.wbits){e.msg="invalid window size",r.mode=30;break}r.dmax=1<<k,e.adler=r.check=1,r.mode=512&u?10:12,l=u=0;break;case 2:for(;l<16;){if(0===o)break e;o--,u+=n[s++]<<l,l+=8}if(r.flags=u,8!=(255&r.flags)){e.msg="unknown compression method",r.mode=30;break}if(57344&r.flags){e.msg="unknown header flags set",r.mode=30;break}r.head&&(r.head.text=u>>8&1),512&r.flags&&(E[0]=255&u,E[1]=u>>>8&255,r.check=B(r.check,E,2,0)),l=u=0,r.mode=3;case 3:for(;l<32;){if(0===o)break e;o--,u+=n[s++]<<l,l+=8}r.head&&(r.head.time=u),512&r.flags&&(E[0]=255&u,E[1]=u>>>8&255,E[2]=u>>>16&255,E[3]=u>>>24&255,r.check=B(r.check,E,4,0)),l=u=0,r.mode=4;case 4:for(;l<16;){if(0===o)break e;o--,u+=n[s++]<<l,l+=8}r.head&&(r.head.xflags=255&u,r.head.os=u>>8),512&r.flags&&(E[0]=255&u,E[1]=u>>>8&255,r.check=B(r.check,E,2,0)),l=u=0,r.mode=5;case 5:if(1024&r.flags){for(;l<16;){if(0===o)break e;o--,u+=n[s++]<<l,l+=8}r.length=u,r.head&&(r.head.extra_len=u),512&r.flags&&(E[0]=255&u,E[1]=u>>>8&255,r.check=B(r.check,E,2,0)),l=u=0}else r.head&&(r.head.extra=null);r.mode=6;case 6:if(1024&r.flags&&(o<(d=r.length)&&(d=o),d&&(r.head&&(k=r.head.extra_len-r.length,r.head.extra||(r.head.extra=new Array(r.head.extra_len)),I.arraySet(r.head.extra,n,s,d,k)),512&r.flags&&(r.check=B(r.check,n,d,s)),o-=d,s+=d,r.length-=d),r.length))break e;r.length=0,r.mode=7;case 7:if(2048&r.flags){if(0===o)break e;for(d=0;k=n[s+d++],r.head&&k&&r.length<65536&&(r.head.name+=String.fromCharCode(k)),k&&d<o;);if(512&r.flags&&(r.check=B(r.check,n,d,s)),o-=d,s+=d,k)break e}else r.head&&(r.head.name=null);r.length=0,r.mode=8;case 8:if(4096&r.flags){if(0===o)break e;for(d=0;k=n[s+d++],r.head&&k&&r.length<65536&&(r.head.comment+=String.fromCharCode(k)),k&&d<o;);if(512&r.flags&&(r.check=B(r.check,n,d,s)),o-=d,s+=d,k)break e}else r.head&&(r.head.comment=null);r.mode=9;case 9:if(512&r.flags){for(;l<16;){if(0===o)break e;o--,u+=n[s++]<<l,l+=8}if(u!==(65535&r.check)){e.msg="header crc mismatch",r.mode=30;break}l=u=0}r.head&&(r.head.hcrc=r.flags>>9&1,r.head.done=!0),e.adler=r.check=0,r.mode=12;break;case 10:for(;l<32;){if(0===o)break e;o--,u+=n[s++]<<l,l+=8}e.adler=r.check=L(u),l=u=0,r.mode=11;case 11:if(0===r.havedict)return e.next_out=a,e.avail_out=h,e.next_in=s,e.avail_in=o,r.hold=u,r.bits=l,2;e.adler=r.check=1,r.mode=12;case 12:if(5===t||6===t)break e;case 13:if(r.last){u>>>=7&l,l-=7&l,r.mode=27;break}for(;l<3;){if(0===o)break e;o--,u+=n[s++]<<l,l+=8}switch(r.last=1&u,l-=1,3&(u>>>=1)){case 0:r.mode=14;break;case 1:if(j(r),r.mode=20,6!==t)break;u>>>=2,l-=2;break e;case 2:r.mode=17;break;case 3:e.msg="invalid block type",r.mode=30}u>>>=2,l-=2;break;case 14:for(u>>>=7&l,l-=7&l;l<32;){if(0===o)break e;o--,u+=n[s++]<<l,l+=8}if((65535&u)!=(u>>>16^65535)){e.msg="invalid stored block lengths",r.mode=30;break}if(r.length=65535&u,l=u=0,r.mode=15,6===t)break e;case 15:r.mode=16;case 16:if(d=r.length){if(o<d&&(d=o),h<d&&(d=h),0===d)break e;I.arraySet(i,n,s,d,a),o-=d,s+=d,h-=d,a+=d,r.length-=d;break}r.mode=12;break;case 17:for(;l<14;){if(0===o)break e;o--,u+=n[s++]<<l,l+=8}if(r.nlen=257+(31&u),u>>>=5,l-=5,r.ndist=1+(31&u),u>>>=5,l-=5,r.ncode=4+(15&u),u>>>=4,l-=4,286<r.nlen||30<r.ndist){e.msg="too many length or distance symbols",r.mode=30;break}r.have=0,r.mode=18;case 18:for(;r.have<r.ncode;){for(;l<3;){if(0===o)break e;o--,u+=n[s++]<<l,l+=8}r.lens[A[r.have++]]=7&u,u>>>=3,l-=3}for(;r.have<19;)r.lens[A[r.have++]]=0;if(r.lencode=r.lendyn,r.lenbits=7,S={bits:r.lenbits},x=T(0,r.lens,0,19,r.lencode,0,r.work,S),r.lenbits=S.bits,x){e.msg="invalid code lengths set",r.mode=30;break}r.have=0,r.mode=19;case 19:for(;r.have<r.nlen+r.ndist;){for(;g=(C=r.lencode[u&(1<<r.lenbits)-1])>>>16&255,b=65535&C,!((_=C>>>24)<=l);){if(0===o)break e;o--,u+=n[s++]<<l,l+=8}if(b<16)u>>>=_,l-=_,r.lens[r.have++]=b;else{if(16===b){for(z=_+2;l<z;){if(0===o)break e;o--,u+=n[s++]<<l,l+=8}if(u>>>=_,l-=_,0===r.have){e.msg="invalid bit length repeat",r.mode=30;break}k=r.lens[r.have-1],d=3+(3&u),u>>>=2,l-=2}else if(17===b){for(z=_+3;l<z;){if(0===o)break e;o--,u+=n[s++]<<l,l+=8}l-=_,k=0,d=3+(7&(u>>>=_)),u>>>=3,l-=3}else{for(z=_+7;l<z;){if(0===o)break e;o--,u+=n[s++]<<l,l+=8}l-=_,k=0,d=11+(127&(u>>>=_)),u>>>=7,l-=7}if(r.have+d>r.nlen+r.ndist){e.msg="invalid bit length repeat",r.mode=30;break}for(;d--;)r.lens[r.have++]=k}}if(30===r.mode)break;if(0===r.lens[256]){e.msg="invalid code -- missing end-of-block",r.mode=30;break}if(r.lenbits=9,S={bits:r.lenbits},x=T(D,r.lens,0,r.nlen,r.lencode,0,r.work,S),r.lenbits=S.bits,x){e.msg="invalid literal/lengths set",r.mode=30;break}if(r.distbits=6,r.distcode=r.distdyn,S={bits:r.distbits},x=T(F,r.lens,r.nlen,r.ndist,r.distcode,0,r.work,S),r.distbits=S.bits,x){e.msg="invalid distances set",r.mode=30;break}if(r.mode=20,6===t)break e;case 20:r.mode=21;case 21:if(6<=o&&258<=h){e.next_out=a,e.avail_out=h,e.next_in=s,e.avail_in=o,r.hold=u,r.bits=l,R(e,c),a=e.next_out,i=e.output,h=e.avail_out,s=e.next_in,n=e.input,o=e.avail_in,u=r.hold,l=r.bits,12===r.mode&&(r.back=-1);break}for(r.back=0;g=(C=r.lencode[u&(1<<r.lenbits)-1])>>>16&255,b=65535&C,!((_=C>>>24)<=l);){if(0===o)break e;o--,u+=n[s++]<<l,l+=8}if(g&&0==(240&g)){for(v=_,y=g,w=b;g=(C=r.lencode[w+((u&(1<<v+y)-1)>>v)])>>>16&255,b=65535&C,!(v+(_=C>>>24)<=l);){if(0===o)break e;o--,u+=n[s++]<<l,l+=8}u>>>=v,l-=v,r.back+=v}if(u>>>=_,l-=_,r.back+=_,r.length=b,0===g){r.mode=26;break}if(32&g){r.back=-1,r.mode=12;break}if(64&g){e.msg="invalid literal/length code",r.mode=30;break}r.extra=15&g,r.mode=22;case 22:if(r.extra){for(z=r.extra;l<z;){if(0===o)break e;o--,u+=n[s++]<<l,l+=8}r.length+=u&(1<<r.extra)-1,u>>>=r.extra,l-=r.extra,r.back+=r.extra}r.was=r.length,r.mode=23;case 23:for(;g=(C=r.distcode[u&(1<<r.distbits)-1])>>>16&255,b=65535&C,!((_=C>>>24)<=l);){if(0===o)break e;o--,u+=n[s++]<<l,l+=8}if(0==(240&g)){for(v=_,y=g,w=b;g=(C=r.distcode[w+((u&(1<<v+y)-1)>>v)])>>>16&255,b=65535&C,!(v+(_=C>>>24)<=l);){if(0===o)break e;o--,u+=n[s++]<<l,l+=8}u>>>=v,l-=v,r.back+=v}if(u>>>=_,l-=_,r.back+=_,64&g){e.msg="invalid distance code",r.mode=30;break}r.offset=b,r.extra=15&g,r.mode=24;case 24:if(r.extra){for(z=r.extra;l<z;){if(0===o)break e;o--,u+=n[s++]<<l,l+=8}r.offset+=u&(1<<r.extra)-1,u>>>=r.extra,l-=r.extra,r.back+=r.extra}if(r.offset>r.dmax){e.msg="invalid distance too far back",r.mode=30;break}r.mode=25;case 25:if(0===h)break e;if(d=c-h,r.offset>d){if((d=r.offset-d)>r.whave&&r.sane){e.msg="invalid distance too far back",r.mode=30;break}p=d>r.wnext?(d-=r.wnext,r.wsize-d):r.wnext-d,d>r.length&&(d=r.length),m=r.window}else m=i,p=a-r.offset,d=r.length;for(h<d&&(d=h),h-=d,r.length-=d;i[a++]=m[p++],--d;);0===r.length&&(r.mode=21);break;case 26:if(0===h)break e;i[a++]=r.length,h--,r.mode=21;break;case 27:if(r.wrap){for(;l<32;){if(0===o)break e;o--,u|=n[s++]<<l,l+=8}if(c-=h,e.total_out+=c,r.total+=c,c&&(e.adler=r.check=r.flags?B(r.check,i,c,a-c):O(r.check,i,c,a-c)),c=h,(r.flags?u:L(u))!==r.check){e.msg="incorrect data check",r.mode=30;break}l=u=0}r.mode=28;case 28:if(r.wrap&&r.flags){for(;l<32;){if(0===o)break e;o--,u+=n[s++]<<l,l+=8}if(u!==(4294967295&r.total)){e.msg="incorrect length check",r.mode=30;break}l=u=0}r.mode=29;case 29:x=1;break e;case 30:x=-3;break e;case 31:return-4;case 32:default:return U}return e.next_out=a,e.avail_out=h,e.next_in=s,e.avail_in=o,r.hold=u,r.bits=l,(r.wsize||c!==e.avail_out&&r.mode<30&&(r.mode<27||4!==t))&&Z(e,e.output,e.next_out,c-e.avail_out)?(r.mode=31,-4):(f-=e.avail_in,c-=e.avail_out,e.total_in+=f,e.total_out+=c,r.total+=c,r.wrap&&c&&(e.adler=r.check=r.flags?B(r.check,i,c,e.next_out-c):O(r.check,i,c,e.next_out-c)),e.data_type=r.bits+(r.last?64:0)+(12===r.mode?128:0)+(20===r.mode||15===r.mode?256:0),(0==f&&0===c||4===t)&&x===N&&(x=-5),x)},r.inflateEnd=function(e){if(!e||!e.state)return U;var t=e.state;return t.window&&(t.window=null),e.state=null,N},r.inflateGetHeader=function(e,t){var r;return e&&e.state?0==(2&(r=e.state).wrap)?U:((r.head=t).done=!1,N):U},r.inflateSetDictionary=function(e,t){var r,n=t.length;return e&&e.state?0!==(r=e.state).wrap&&11!==r.mode?U:11===r.mode&&O(1,t,n,0)!==r.check?-3:Z(e,t,n,n)?(r.mode=31,-4):(r.havedict=1,N):U},r.inflateInfo="pako inflate (from Nodeca project)"},{"../utils/common":41,"./adler32":43,"./crc32":45,"./inffast":48,"./inftrees":50}],50:[function(e,t,r){"use strict";var D=e("../utils/common"),F=[3,4,5,6,7,8,9,10,11,13,15,17,19,23,27,31,35,43,51,59,67,83,99,115,131,163,195,227,258,0,0],N=[16,16,16,16,16,16,16,16,17,17,17,17,18,18,18,18,19,19,19,19,20,20,20,20,21,21,21,21,16,72,78],U=[1,2,3,4,5,7,9,13,17,25,33,49,65,97,129,193,257,385,513,769,1025,1537,2049,3073,4097,6145,8193,12289,16385,24577,0,0],P=[16,16,16,16,17,17,18,18,19,19,20,20,21,21,22,22,23,23,24,24,25,25,26,26,27,27,28,28,29,29,64,64];t.exports=function(e,t,r,n,i,s,a,o){var h,u,l,f,c,d,p,m,_,g=o.bits,b=0,v=0,y=0,w=0,k=0,x=0,S=0,z=0,C=0,E=0,A=null,I=0,O=new D.Buf16(16),B=new D.Buf16(16),R=null,T=0;for(b=0;b<=15;b++)O[b]=0;for(v=0;v<n;v++)O[t[r+v]]++;for(k=g,w=15;1<=w&&0===O[w];w--);if(w<k&&(k=w),0===w)return i[s++]=20971520,i[s++]=20971520,o.bits=1,0;for(y=1;y<w&&0===O[y];y++);for(k<y&&(k=y),b=z=1;b<=15;b++)if(z<<=1,(z-=O[b])<0)return-1;if(0<z&&(0===e||1!==w))return-1;for(B[1]=0,b=1;b<15;b++)B[b+1]=B[b]+O[b];for(v=0;v<n;v++)0!==t[r+v]&&(a[B[t[r+v]]++]=v);if(d=0===e?(A=R=a,19):1===e?(A=F,I-=257,R=N,T-=257,256):(A=U,R=P,-1),b=y,c=s,S=v=E=0,l=-1,f=(C=1<<(x=k))-1,1===e&&852<C||2===e&&592<C)return 1;for(;;){for(p=b-S,_=a[v]<d?(m=0,a[v]):a[v]>d?(m=R[T+a[v]],A[I+a[v]]):(m=96,0),h=1<<b-S,y=u=1<<x;i[c+(E>>S)+(u-=h)]=p<<24|m<<16|_|0,0!==u;);for(h=1<<b-1;E&h;)h>>=1;if(0!==h?(E&=h-1,E+=h):E=0,v++,0==--O[b]){if(b===w)break;b=t[r+a[v]]}if(k<b&&(E&f)!==l){for(0===S&&(S=k),c+=y,z=1<<(x=b-S);x+S<w&&!((z-=O[x+S])<=0);)x++,z<<=1;if(C+=1<<x,1===e&&852<C||2===e&&592<C)return 1;i[l=E&f]=k<<24|x<<16|c-s|0}}return 0!==E&&(i[c+E]=b-S<<24|64<<16|0),o.bits=k,0}},{"../utils/common":41}],51:[function(e,t,r){"use strict";t.exports={2:"need dictionary",1:"stream end",0:"","-1":"file error","-2":"stream error","-3":"data error","-4":"insufficient memory","-5":"buffer error","-6":"incompatible version"}},{}],52:[function(e,t,r){"use strict";var i=e("../utils/common"),o=0,h=1;function n(e){for(var t=e.length;0<=--t;)e[t]=0}var s=0,a=29,u=256,l=u+1+a,f=30,c=19,_=2*l+1,g=15,d=16,p=7,m=256,b=16,v=17,y=18,w=[0,0,0,0,0,0,0,0,1,1,1,1,2,2,2,2,3,3,3,3,4,4,4,4,5,5,5,5,0],k=[0,0,0,0,1,1,2,2,3,3,4,4,5,5,6,6,7,7,8,8,9,9,10,10,11,11,12,12,13,13],x=[0,0,0,0,0,0,0,0,0,0,0,0,0,0,0,0,2,3,7],S=[16,17,18,0,8,7,9,6,10,5,11,4,12,3,13,2,14,1,15],z=new Array(2*(l+2));n(z);var C=new Array(2*f);n(C);var E=new Array(512);n(E);var A=new Array(256);n(A);var I=new Array(a);n(I);var O,B,R,T=new Array(f);function D(e,t,r,n,i){this.static_tree=e,this.extra_bits=t,this.extra_base=r,this.elems=n,this.max_length=i,this.has_stree=e&&e.length}function F(e,t){this.dyn_tree=e,this.max_code=0,this.stat_desc=t}function N(e){return e<256?E[e]:E[256+(e>>>7)]}function U(e,t){e.pending_buf[e.pending++]=255&t,e.pending_buf[e.pending++]=t>>>8&255}function P(e,t,r){e.bi_valid>d-r?(e.bi_buf|=t<<e.bi_valid&65535,U(e,e.bi_buf),e.bi_buf=t>>d-e.bi_valid,e.bi_valid+=r-d):(e.bi_buf|=t<<e.bi_valid&65535,e.bi_valid+=r)}function L(e,t,r){P(e,r[2*t],r[2*t+1])}function j(e,t){for(var r=0;r|=1&e,e>>>=1,r<<=1,0<--t;);return r>>>1}function Z(e,t,r){var n,i,s=new Array(g+1),a=0;for(n=1;n<=g;n++)s[n]=a=a+r[n-1]<<1;for(i=0;i<=t;i++){var o=e[2*i+1];0!==o&&(e[2*i]=j(s[o]++,o))}}function W(e){var t;for(t=0;t<l;t++)e.dyn_ltree[2*t]=0;for(t=0;t<f;t++)e.dyn_dtree[2*t]=0;for(t=0;t<c;t++)e.bl_tree[2*t]=0;e.dyn_ltree[2*m]=1,e.opt_len=e.static_len=0,e.last_lit=e.matches=0}function M(e){8<e.bi_valid?U(e,e.bi_buf):0<e.bi_valid&&(e.pending_buf[e.pending++]=e.bi_buf),e.bi_buf=0,e.bi_valid=0}function H(e,t,r,n){var i=2*t,s=2*r;return e[i]<e[s]||e[i]===e[s]&&n[t]<=n[r]}function G(e,t,r){for(var n=e.heap[r],i=r<<1;i<=e.heap_len&&(i<e.heap_len&&H(t,e.heap[i+1],e.heap[i],e.depth)&&i++,!H(t,n,e.heap[i],e.depth));)e.heap[r]=e.heap[i],r=i,i<<=1;e.heap[r]=n}function K(e,t,r){var n,i,s,a,o=0;if(0!==e.last_lit)for(;n=e.pending_buf[e.d_buf+2*o]<<8|e.pending_buf[e.d_buf+2*o+1],i=e.pending_buf[e.l_buf+o],o++,0===n?L(e,i,t):(L(e,(s=A[i])+u+1,t),0!==(a=w[s])&&P(e,i-=I[s],a),L(e,s=N(--n),r),0!==(a=k[s])&&P(e,n-=T[s],a)),o<e.last_lit;);L(e,m,t)}function Y(e,t){var r,n,i,s=t.dyn_tree,a=t.stat_desc.static_tree,o=t.stat_desc.has_stree,h=t.stat_desc.elems,u=-1;for(e.heap_len=0,e.heap_max=_,r=0;r<h;r++)0!==s[2*r]?(e.heap[++e.heap_len]=u=r,e.depth[r]=0):s[2*r+1]=0;for(;e.heap_len<2;)s[2*(i=e.heap[++e.heap_len]=u<2?++u:0)]=1,e.depth[i]=0,e.opt_len--,o&&(e.static_len-=a[2*i+1]);for(t.max_code=u,r=e.heap_len>>1;1<=r;r--)G(e,s,r);for(i=h;r=e.heap[1],e.heap[1]=e.heap[e.heap_len--],G(e,s,1),n=e.heap[1],e.heap[--e.heap_max]=r,e.heap[--e.heap_max]=n,s[2*i]=s[2*r]+s[2*n],e.depth[i]=(e.depth[r]>=e.depth[n]?e.depth[r]:e.depth[n])+1,s[2*r+1]=s[2*n+1]=i,e.heap[1]=i++,G(e,s,1),2<=e.heap_len;);e.heap[--e.heap_max]=e.heap[1],function(e,t){var r,n,i,s,a,o,h=t.dyn_tree,u=t.max_code,l=t.stat_desc.static_tree,f=t.stat_desc.has_stree,c=t.stat_desc.extra_bits,d=t.stat_desc.extra_base,p=t.stat_desc.max_length,m=0;for(s=0;s<=g;s++)e.bl_count[s]=0;for(h[2*e.heap[e.heap_max]+1]=0,r=e.heap_max+1;r<_;r++)p<(s=h[2*h[2*(n=e.heap[r])+1]+1]+1)&&(s=p,m++),h[2*n+1]=s,u<n||(e.bl_count[s]++,a=0,d<=n&&(a=c[n-d]),o=h[2*n],e.opt_len+=o*(s+a),f&&(e.static_len+=o*(l[2*n+1]+a)));if(0!==m){do{for(s=p-1;0===e.bl_count[s];)s--;e.bl_count[s]--,e.bl_count[s+1]+=2,e.bl_count[p]--,m-=2}while(0<m);for(s=p;0!==s;s--)for(n=e.bl_count[s];0!==n;)u<(i=e.heap[--r])||(h[2*i+1]!==s&&(e.opt_len+=(s-h[2*i+1])*h[2*i],h[2*i+1]=s),n--)}}(e,t),Z(s,u,e.bl_count)}function X(e,t,r){var n,i,s=-1,a=t[1],o=0,h=7,u=4;for(0===a&&(h=138,u=3),t[2*(r+1)+1]=65535,n=0;n<=r;n++)i=a,a=t[2*(n+1)+1],++o<h&&i===a||(o<u?e.bl_tree[2*i]+=o:0!==i?(i!==s&&e.bl_tree[2*i]++,e.bl_tree[2*b]++):o<=10?e.bl_tree[2*v]++:e.bl_tree[2*y]++,s=i,u=(o=0)===a?(h=138,3):i===a?(h=6,3):(h=7,4))}function V(e,t,r){var n,i,s=-1,a=t[1],o=0,h=7,u=4;for(0===a&&(h=138,u=3),n=0;n<=r;n++)if(i=a,a=t[2*(n+1)+1],!(++o<h&&i===a)){if(o<u)for(;L(e,i,e.bl_tree),0!=--o;);else 0!==i?(i!==s&&(L(e,i,e.bl_tree),o--),L(e,b,e.bl_tree),P(e,o-3,2)):o<=10?(L(e,v,e.bl_tree),P(e,o-3,3)):(L(e,y,e.bl_tree),P(e,o-11,7));s=i,u=(o=0)===a?(h=138,3):i===a?(h=6,3):(h=7,4)}}n(T);var q=!1;function J(e,t,r,n){P(e,(s<<1)+(n?1:0),3),function(e,t,r,n){M(e),n&&(U(e,r),U(e,~r)),i.arraySet(e.pending_buf,e.window,t,r,e.pending),e.pending+=r}(e,t,r,!0)}r._tr_init=function(e){q||(function(){var e,t,r,n,i,s=new Array(g+1);for(n=r=0;n<a-1;n++)for(I[n]=r,e=0;e<1<<w[n];e++)A[r++]=n;for(A[r-1]=n,n=i=0;n<16;n++)for(T[n]=i,e=0;e<1<<k[n];e++)E[i++]=n;for(i>>=7;n<f;n++)for(T[n]=i<<7,e=0;e<1<<k[n]-7;e++)E[256+i++]=n;for(t=0;t<=g;t++)s[t]=0;for(e=0;e<=143;)z[2*e+1]=8,e++,s[8]++;for(;e<=255;)z[2*e+1]=9,e++,s[9]++;for(;e<=279;)z[2*e+1]=7,e++,s[7]++;for(;e<=287;)z[2*e+1]=8,e++,s[8]++;for(Z(z,l+1,s),e=0;e<f;e++)C[2*e+1]=5,C[2*e]=j(e,5);O=new D(z,w,u+1,l,g),B=new D(C,k,0,f,g),R=new D(new Array(0),x,0,c,p)}(),q=!0),e.l_desc=new F(e.dyn_ltree,O),e.d_desc=new F(e.dyn_dtree,B),e.bl_desc=new F(e.bl_tree,R),e.bi_buf=0,e.bi_valid=0,W(e)},r._tr_stored_block=J,r._tr_flush_block=function(e,t,r,n){var i,s,a=0;0<e.level?(2===e.strm.data_type&&(e.strm.data_type=function(e){var t,r=4093624447;for(t=0;t<=31;t++,r>>>=1)if(1&r&&0!==e.dyn_ltree[2*t])return o;if(0!==e.dyn_ltree[18]||0!==e.dyn_ltree[20]||0!==e.dyn_ltree[26])return h;for(t=32;t<u;t++)if(0!==e.dyn_ltree[2*t])return h;return o}(e)),Y(e,e.l_desc),Y(e,e.d_desc),a=function(e){var t;for(X(e,e.dyn_ltree,e.l_desc.max_code),X(e,e.dyn_dtree,e.d_desc.max_code),Y(e,e.bl_desc),t=c-1;3<=t&&0===e.bl_tree[2*S[t]+1];t--);return e.opt_len+=3*(t+1)+5+5+4,t}(e),i=e.opt_len+3+7>>>3,(s=e.static_len+3+7>>>3)<=i&&(i=s)):i=s=r+5,r+4<=i&&-1!==t?J(e,t,r,n):4===e.strategy||s===i?(P(e,2+(n?1:0),3),K(e,z,C)):(P(e,4+(n?1:0),3),function(e,t,r,n){var i;for(P(e,t-257,5),P(e,r-1,5),P(e,n-4,4),i=0;i<n;i++)P(e,e.bl_tree[2*S[i]+1],3);V(e,e.dyn_ltree,t-1),V(e,e.dyn_dtree,r-1)}(e,e.l_desc.max_code+1,e.d_desc.max_code+1,a+1),K(e,e.dyn_ltree,e.dyn_dtree)),W(e),n&&M(e)},r._tr_tally=function(e,t,r){return e.pending_buf[e.d_buf+2*e.last_lit]=t>>>8&255,e.pending_buf[e.d_buf+2*e.last_lit+1]=255&t,e.pending_buf[e.l_buf+e.last_lit]=255&r,e.last_lit++,0===t?e.dyn_ltree[2*r]++:(e.matches++,t--,e.dyn_ltree[2*(A[r]+u+1)]++,e.dyn_dtree[2*N(t)]++),e.last_lit===e.lit_bufsize-1},r._tr_align=function(e){P(e,2,3),L(e,m,z),function(e){16===e.bi_valid?(U(e,e.bi_buf),e.bi_buf=0,e.bi_valid=0):8<=e.bi_valid&&(e.pending_buf[e.pending++]=255&e.bi_buf,e.bi_buf>>=8,e.bi_valid-=8)}(e)}},{"../utils/common":41}],53:[function(e,t,r){"use strict";t.exports=function(){this.input=null,this.next_in=0,this.avail_in=0,this.total_in=0,this.output=null,this.next_out=0,this.avail_out=0,this.total_out=0,this.msg="",this.state=null,this.data_type=2,this.adler=0}},{}],54:[function(e,t,r){(function(e){!function(r,n){"use strict";if(!r.setImmediate){var i,s,t,a,o=1,h={},u=!1,l=r.document,e=Object.getPrototypeOf&&Object.getPrototypeOf(r);e=e&&e.setTimeout?e:r,i="[object process]"==={}.toString.call(r.process)?function(e){process.nextTick(function(){c(e)})}:function(){if(r.postMessage&&!r.importScripts){var e=!0,t=r.onmessage;return r.onmessage=function(){e=!1},r.postMessage("","*"),r.onmessage=t,e}}()?(a="setImmediate$"+Math.random()+"$",r.addEventListener?r.addEventListener("message",d,!1):r.attachEvent("onmessage",d),function(e){r.postMessage(a+e,"*")}):r.MessageChannel?((t=new MessageChannel).port1.onmessage=function(e){c(e.data)},function(e){t.port2.postMessage(e)}):l&&"onreadystatechange"in l.createElement("script")?(s=l.documentElement,function(e){var t=l.createElement("script");t.onreadystatechange=function(){c(e),t.onreadystatechange=null,s.removeChild(t),t=null},s.appendChild(t)}):function(e){setTimeout(c,0,e)},e.setImmediate=function(e){"function"!=typeof e&&(e=new Function(""+e));for(var t=new Array(arguments.length-1),r=0;r<t.length;r++)t[r]=arguments[r+1];var n={callback:e,args:t};return h[o]=n,i(o),o++},e.clearImmediate=f}function f(e){delete h[e]}function c(e){if(u)setTimeout(c,0,e);else{var t=h[e];if(t){u=!0;try{!function(e){var t=e.callback,r=e.args;switch(r.length){case 0:t();break;case 1:t(r[0]);break;case 2:t(r[0],r[1]);break;case 3:t(r[0],r[1],r[2]);break;default:t.apply(n,r)}}(t)}finally{f(e),u=!1}}}}function d(e){e.source===r&&"string"==typeof e.data&&0===e.data.indexOf(a)&&c(+e.data.slice(a.length))}}("undefined"==typeof self?void 0===e?this:e:self)}).call(this,"undefined"!=typeof global?global:"undefined"!=typeof self?self:"undefined"!=typeof window?window:{})},{}]},{},[10])(10)});
```
