## supplement_file1 for "GENETH’OFF: a flexible workflow for genome-wide profiling of CRISPR/Cas off-targets": jquery.nouislider.min.js

```
/*! noUiSlider - 7.0.10 - 2015-03-17 17:30:22 */

!function(a){"use strict";function b(a,b){return Math.round(a/b)*b}function c(a){return"number"==typeof a&&!isNaN(a)&&isFinite(a)}function d(a){var b=Math.pow(10,7);return Number((Math.round(a*b)/b).toFixed(7))}function e(a,b,c){a.addClass(b),setTimeout(function(){a.removeClass(b)},c)}function f(a){return Math.max(Math.min(a,100),0)}function g(b){return a.isArray(b)?b:[b]}function h(a){var b=a.split(".");return b.length>1?b[1].length:0}function i(a,b){return 100/(b-a)}function j(a,b){return 100*b/(a[1]-a[0])}function k(a,b){return j(a,a[0]<0?b+Math.abs(a[0]):b-a[0])}function l(a,b){return b*(a[1]-a[0])/100+a[0]}function m(a,b){for(var c=1;a>=b[c];)c+=1;return c}function n(a,b,c){if(c>=a.slice(-1)[0])return 100;var d,e,f,g,h=m(c,a);return d=a[h-1],e=a[h],f=b[h-1],g=b[h],f+k([d,e],c)/i(f,g)}function o(a,b,c){if(c>=100)return a.slice(-1)[0];var d,e,f,g,h=m(c,b);return d=a[h-1],e=a[h],f=b[h-1],g=b[h],l([d,e],(c-f)*i(f,g))}function p(a,c,d,e){if(100===e)return e;var f,g,h=m(e,a);return d?(f=a[h-1],g=a[h],e-f>(g-f)/2?g:f):c[h-1]?a[h-1]+b(e-a[h-1],c[h-1]):e}function q(a,b,d){var e;if("number"==typeof b&&(b=[b]),"[object Array]"!==Object.prototype.toString.call(b))throw new Error("noUiSlider: 'range' contains invalid value.");if(e="min"===a?0:"max"===a?100:parseFloat(a),!c(e)||!c(b[0]))throw new Error("noUiSlider: 'range' value isn't numeric.");d.xPct.push(e),d.xVal.push(b[0]),e?d.xSteps.push(isNaN(b[1])?!1:b[1]):isNaN(b[1])||(d.xSteps[0]=b[1])}function r(a,b,c){return b?void(c.xSteps[a]=j([c.xVal[a],c.xVal[a+1]],b)/i(c.xPct[a],c.xPct[a+1])):!0}function s(a,b,c,d){this.xPct=[],this.xVal=[],this.xSteps=[d||!1],this.xNumSteps=[!1],this.snap=b,this.direction=c;var e,f=[];for(e in a)a.hasOwnProperty(e)&&f.push([a[e],e]);for(f.sort(function(a,b){return a[0]-b[0]}),e=0;e<f.length;e++)q(f[e][1],f[e][0],this);for(this.xNumSteps=this.xSteps.slice(0),e=0;e<this.xNumSteps.length;e++)r(e,this.xNumSteps[e],this)}function t(a,b){if(!c(b))throw new Error("noUiSlider: 'step' is not numeric.");a.singleStep=b}function u(b,c){if("object"!=typeof c||a.isArray(c))throw new Error("noUiSlider: 'range' is not an object.");if(void 0===c.min||void 0===c.max)throw new Error("noUiSlider: Missing 'min' or 'max' in 'range'.");b.spectrum=new s(c,b.snap,b.dir,b.singleStep)}function v(b,c){if(c=g(c),!a.isArray(c)||!c.length||c.length>2)throw new Error("noUiSlider: 'start' option is incorrect.");b.handles=c.length,b.start=c}function w(a,b){if(a.snap=b,"boolean"!=typeof b)throw new Error("noUiSlider: 'snap' option must be a boolean.")}function x(a,b){if(a.animate=b,"boolean"!=typeof b)throw new Error("noUiSlider: 'animate' option must be a boolean.")}function y(a,b){if("lower"===b&&1===a.handles)a.connect=1;else if("upper"===b&&1===a.handles)a.connect=2;else if(b===!0&&2===a.handles)a.connect=3;else{if(b!==!1)throw new Error("noUiSlider: 'connect' option doesn't match handle count.");a.connect=0}}function z(a,b){switch(b){case"horizontal":a.ort=0;break;case"vertical":a.ort=1;break;default:throw new Error("noUiSlider: 'orientation' option is invalid.")}}function A(a,b){if(!c(b))throw new Error("noUiSlider: 'margin' option must be numeric.");if(a.margin=a.spectrum.getMargin(b),!a.margin)throw new Error("noUiSlider: 'margin' option is only supported on linear sliders.")}function B(a,b){if(!c(b))throw new Error("noUiSlider: 'limit' option must be numeric.");if(a.limit=a.spectrum.getMargin(b),!a.limit)throw new Error("noUiSlider: 'limit' option is only supported on linear sliders.")}function C(a,b){switch(b){case"ltr":a.dir=0;break;case"rtl":a.dir=1,a.connect=[0,2,1,3][a.connect];break;default:throw new Error("noUiSlider: 'direction' option was not recognized.")}}function D(a,b){if("string"!=typeof b)throw new Error("noUiSlider: 'behaviour' must be a string containing options.");var c=b.indexOf("tap")>=0,d=b.indexOf("drag")>=0,e=b.indexOf("fixed")>=0,f=b.indexOf("snap")>=0;a.events={tap:c||f,drag:d,fixed:e,snap:f}}function E(a,b){if(a.format=b,"function"==typeof b.to&&"function"==typeof b.from)return!0;throw new Error("noUiSlider: 'format' requires 'to' and 'from' methods.")}function F(b){var c,d={margin:0,limit:0,animate:!0,format:V};return c={step:{r:!1,t:t},start:{r:!0,t:v},connect:{r:!0,t:y},direction:{r:!0,t:C},snap:{r:!1,t:w},animate:{r:!1,t:x},range:{r:!0,t:u},orientation:{r:!1,t:z},margin:{r:!1,t:A},limit:{r:!1,t:B},behaviour:{r:!0,t:D},format:{r:!1,t:E}},b=a.extend({connect:!1,direction:"ltr",behaviour:"tap",orientation:"horizontal"},b),a.each(c,function(a,c){if(void 0===b[a]){if(c.r)throw new Error("noUiSlider: '"+a+"' is required.");return!0}c.t(d,b[a])}),d.style=d.ort?"top":"left",d}function G(a,b,c){var d=a+b[0],e=a+b[1];return c?(0>d&&(e+=Math.abs(d)),e>100&&(d-=e-100),[f(d),f(e)]):[d,e]}function H(a){a.preventDefault();var b,c,d=0===a.type.indexOf("touch"),e=0===a.type.indexOf("mouse"),f=0===a.type.indexOf("pointer"),g=a;return 0===a.type.indexOf("MSPointer")&&(f=!0),a.originalEvent&&(a=a.originalEvent),d&&(b=a.changedTouches[0].pageX,c=a.changedTouches[0].pageY),(e||f)&&(f||void 0!==window.pageXOffset||(window.pageXOffset=document.documentElement.scrollLeft,window.pageYOffset=document.documentElement.scrollTop),b=a.clientX+window.pageXOffset,c=a.clientY+window.pageYOffset),g.points=[b,c],g.cursor=e,g}function I(b,c){var d=a("<div><div/></div>").addClass(U[2]),e=["-lower","-upper"];return b&&e.reverse(),d.children().addClass(U[3]+" "+U[3]+e[c]),d}function J(a,b,c){switch(a){case 1:b.addClass(U[7]),c[0].addClass(U[6]);break;case 3:c[1].addClass(U[6]);case 2:c[0].addClass(U[7]);case 0:b.addClass(U[6])}}function K(a,b,c){var d,e=[];for(d=0;a>d;d+=1)e.push(I(b,d).appendTo(c));return e}function L(b,c,d){return d.addClass([U[0],U[8+b],U[4+c]].join(" ")),a("<div/>").appendTo(d).addClass(U[1])}function M(b,c,d){function i(){return C[["width","height"][c.ort]]()}function j(a){var b,c=[E.val()];for(b=0;b<a.length;b+=1)E.trigger(a[b],c)}function k(a){return 1===a.length?a[0]:c.dir?a.reverse():a}function l(a){return function(b,c){E.val([a?null:c,a?c:null],!0)}}function m(b){var c=a.inArray(b,N);E[0].linkAPI&&E[0].linkAPI[b]&&E[0].linkAPI[b].change(M[c],D[c].children(),E)}function n(b,d){var e=a.inArray(b,N);return d&&d.appendTo(D[e].children()),c.dir&&c.handles>1&&(e=1===e?0:1),l(e)}function o(){var a,b;for(a=0;a<N.length;a+=1)this.linkAPI&&this.linkAPI[b=N[a]]&&this.linkAPI[b].reconfirm(b)}function p(a,b,d,e){return a=a.replace(/\s/g,S+" ")+S,b.on(a,function(a){return E.attr("disabled")?!1:E.hasClass(U[14])?!1:(a=H(a),a.calcPoint=a.points[c.ort],void d(a,e))})}function q(a,b){var c,d=b.handles||D,e=!1,f=100*(a.calcPoint-b.start)/i(),g=d[0][0]!==D[0][0]?1:0;c=G(f,b.positions,d.length>1),e=v(d[0],c[g],1===d.length),d.length>1&&(e=v(d[1],c[g?0:1],!1)||e),e&&j(["slide"])}function r(b){a("."+U[15]).removeClass(U[15]),b.cursor&&a("body").css("cursor","").off(S),Q.off(S),E.removeClass(U[12]),j(["set","change"])}function s(b,c){1===c.handles.length&&c.handles[0].children().addClass(U[15]),b.stopPropagation(),p(T.move,Q,q,{start:b.calcPoint,handles:c.handles,positions:[F[0],F[D.length-1]]}),p(T.end,Q,r,null),b.cursor&&(a("body").css("cursor",a(b.target).css("cursor")),D.length>1&&E.addClass(U[12]),a("body").on("selectstart"+S,!1))}function t(b){var d,f=b.calcPoint,g=0;b.stopPropagation(),a.each(D,function(){g+=this.offset()[c.style]}),g=g/2>f||1===D.length?0:1,f-=C.offset()[c.style],d=100*f/i(),c.events.snap||e(E,U[14],300),v(D[g],d),j(["slide","set","change"]),c.events.snap&&s(b,{handles:[D[g]]})}function u(a){var b,c;if(!a.fixed)for(b=0;b<D.length;b+=1)p(T.start,D[b].children(),s,{handles:[D[b]]});a.tap&&p(T.start,C,t,{handles:D}),a.drag&&(c=C.find("."+U[7]).addClass(U[10]),a.fixed&&(c=c.add(C.children().not(c).children())),p(T.start,c,s,{handles:D}))}function v(a,b,d){var e=a[0]!==D[0][0]?1:0,g=F[0]+c.margin,h=F[1]-c.margin,i=F[0]+c.limit,j=F[1]-c.limit;return D.length>1&&(b=e?Math.max(b,g):Math.min(b,h)),d!==!1&&c.limit&&D.length>1&&(b=e?Math.min(b,i):Math.max(b,j)),b=I.getStep(b),b=f(parseFloat(b.toFixed(7))),b===F[e]?!1:(a.css(c.style,b+"%"),a.is(":first-child")&&a.toggleClass(U[17],b>50),F[e]=b,M[e]=I.fromStepping(b),m(N[e]),!0)}function w(a,b){var d,e,f;for(c.limit&&(a+=1),d=0;a>d;d+=1)e=d%2,f=b[e],null!==f&&f!==!1&&("number"==typeof f&&(f=String(f)),f=c.format.from(f),(f===!1||isNaN(f)||v(D[e],I.toStepping(f),d===3-c.dir)===!1)&&m(N[e]))}function x(a){if(E[0].LinkIsEmitting)return this;var b,d=g(a);return c.dir&&c.handles>1&&d.reverse(),c.animate&&-1!==F[0]&&e(E,U[14],300),b=D.length>1?3:1,1===d.length&&(b=1),w(b,d),j(["set"]),this}function y(){var a,b=[];for(a=0;a<c.handles;a+=1)b[a]=c.format.to(M[a]);return k(b)}function z(){return a(this).off(S).removeClass(U.join(" ")).empty(),delete this.LinkUpdate,delete this.LinkConfirm,delete this.LinkDefaultFormatter,delete this.LinkDefaultFlag,delete this.reappend,delete this.vGet,delete this.vSet,delete this.getCurrentStep,delete this.getInfo,delete this.destroy,d}function A(){var b=a.map(F,function(a,b){var c=I.getApplicableStep(a),d=h(String(c[2])),e=M[b],f=100===a?null:c[2],g=Number((e-c[2]).toFixed(d)),i=0===a?null:g>=c[1]?c[2]:c[0]||!1;return[[i,f]]});return k(b)}function B(){return d}var C,D,E=a(b),F=[-1,-1],I=c.spectrum,M=[],N=["lower","upper"].slice(0,c.handles);if(c.dir&&N.reverse(),b.LinkUpdate=m,b.LinkConfirm=n,b.LinkDefaultFormatter=c.format,b.LinkDefaultFlag="lower",b.reappend=o,E.hasClass(U[0]))throw new Error("Slider was already initialized.");C=L(c.dir,c.ort,E),D=K(c.handles,c.dir,C),J(c.connect,E,D),u(c.events),b.vSet=x,b.vGet=y,b.destroy=z,b.getCurrentStep=A,b.getOriginalOptions=B,b.getInfo=function(){return[I,c.style,c.ort]},E.val(c.start)}function N(a){var b=F(a,this);return this.each(function(){M(this,b,a)})}function O(b){return this.each(function(){if(!this.destroy)return void a(this).noUiSlider(b);var c=a(this).val(),d=this.destroy(),e=a.extend({},d,b);a(this).noUiSlider(e),this.reappend(),d.start===e.start&&a(this).val(c)})}function P(){return this[0][arguments.length?"vSet":"vGet"].apply(this[0],arguments)}var Q=a(document),R=a.fn.val,S=".nui",T=window.navigator.pointerEnabled?{start:"pointerdown",move:"pointermove",end:"pointerup"}:window.navigator.msPointerEnabled?{start:"MSPointerDown",move:"MSPointerMove",end:"MSPointerUp"}:{start:"mousedown touchstart",move:"mousemove touchmove",end:"mouseup touchend"},U=["noUi-target","noUi-base","noUi-origin","noUi-handle","noUi-horizontal","noUi-vertical","noUi-background","noUi-connect","noUi-ltr","noUi-rtl","noUi-dragable","","noUi-state-drag","","noUi-state-tap","noUi-active","","noUi-stacking"];s.prototype.getMargin=function(a){return 2===this.xPct.length?j(this.xVal,a):!1},s.prototype.toStepping=function(a){return a=n(this.xVal,this.xPct,a),this.direction&&(a=100-a),a},s.prototype.fromStepping=function(a){return this.direction&&(a=100-a),d(o(this.xVal,this.xPct,a))},s.prototype.getStep=function(a){return this.direction&&(a=100-a),a=p(this.xPct,this.xSteps,this.snap,a),this.direction&&(a=100-a),a},s.prototype.getApplicableStep=function(a){var b=m(a,this.xPct),c=100===a?2:1;return[this.xNumSteps[b-2],this.xVal[b-c],this.xNumSteps[b-c]]},s.prototype.convert=function(a){return this.getStep(this.toStepping(a))};var V={to:function(a){return a.toFixed(2)},from:Number};a.fn.val=function(b){function c(a){return a.hasClass(U[0])?P:R}if(!arguments.length){var d=a(this[0]);return c(d).call(d)}var e=a.isFunction(b);return this.each(function(d){var f=b,g=a(this);e&&(f=b.call(this,d,g.val())),c(g).call(g,f)})},a.fn.noUiSlider=function(a,b){switch(a){case"step":return this[0].getCurrentStep();case"options":return this[0].getOriginalOptions()}return(b?O:N).call(this,a)}}(window.jQuery||window.Zepto);
```
