## supplement_file1 for "GENETH’OFF: a flexible workflow for genome-wide profiling of CRISPR/Cas off-targets": selectize.min.js

```
/*! selectize.js - v0.12.1 | https://github.com/brianreavis/selectize.js | Apache License (v2) */
!function(a,b){"function"==typeof define&&define.amd?define("sifter",b):"object"==typeof exports?module.exports=b():a.Sifter=b()}(this,function(){var a=function(a,b){this.items=a,this.settings=b||{diacritics:!0}};a.prototype.tokenize=function(a){if(a=d(String(a||"").toLowerCase()),!a||!a.length)return[];var b,c,f,h,i=[],j=a.split(/ +/);for(b=0,c=j.length;c>b;b++){if(f=e(j[b]),this.settings.diacritics)for(h in g)g.hasOwnProperty(h)&&(f=f.replace(new RegExp(h,"g"),g[h]));i.push({string:j[b],regex:new RegExp(f,"i")})}return i},a.prototype.iterator=function(a,b){var c;c=f(a)?Array.prototype.forEach||function(a){for(var b=0,c=this.length;c>b;b++)a(this[b],b,this)}:function(a){for(var b in this)this.hasOwnProperty(b)&&a(this[b],b,this)},c.apply(a,[b])},a.prototype.getScoreFunction=function(a,b){var c,d,e,f;c=this,a=c.prepareSearch(a,b),e=a.tokens,d=a.options.fields,f=e.length;var g=function(a,b){var c,d;return a?(a=String(a||""),d=a.search(b.regex),-1===d?0:(c=b.string.length/a.length,0===d&&(c+=.5),c)):0},h=function(){var a=d.length;return a?1===a?function(a,b){return g(b[d[0]],a)}:function(b,c){for(var e=0,f=0;a>e;e++)f+=g(c[d[e]],b);return f/a}:function(){return 0}}();return f?1===f?function(a){return h(e[0],a)}:"and"===a.options.conjunction?function(a){for(var b,c=0,d=0;f>c;c++){if(b=h(e[c],a),0>=b)return 0;d+=b}return d/f}:function(a){for(var b=0,c=0;f>b;b++)c+=h(e[b],a);return c/f}:function(){return 0}},a.prototype.getSortFunction=function(a,c){var d,e,f,g,h,i,j,k,l,m,n;if(f=this,a=f.prepareSearch(a,c),n=!a.query&&c.sort_empty||c.sort,l=function(a,b){return"$score"===a?b.score:f.items[b.id][a]},h=[],n)for(d=0,e=n.length;e>d;d++)(a.query||"$score"!==n[d].field)&&h.push(n[d]);if(a.query){for(m=!0,d=0,e=h.length;e>d;d++)if("$score"===h[d].field){m=!1;break}m&&h.unshift({field:"$score",direction:"desc"})}else for(d=0,e=h.length;e>d;d++)if("$score"===h[d].field){h.splice(d,1);break}for(k=[],d=0,e=h.length;e>d;d++)k.push("desc"===h[d].direction?-1:1);return i=h.length,i?1===i?(g=h[0].field,j=k[0],function(a,c){return j*b(l(g,a),l(g,c))}):function(a,c){var d,e,f;for(d=0;i>d;d++)if(f=h[d].field,e=k[d]*b(l(f,a),l(f,c)))return e;return 0}:null},a.prototype.prepareSearch=function(a,b){if("object"==typeof a)return a;b=c({},b);var d=b.fields,e=b.sort,g=b.sort_empty;return d&&!f(d)&&(b.fields=[d]),e&&!f(e)&&(b.sort=[e]),g&&!f(g)&&(b.sort_empty=[g]),{options:b,query:String(a||"").toLowerCase(),tokens:this.tokenize(a),total:0,items:[]}},a.prototype.search=function(a,b){var c,d,e,f,g=this;return d=this.prepareSearch(a,b),b=d.options,a=d.query,f=b.score||g.getScoreFunction(d),a.length?g.iterator(g.items,function(a,e){c=f(a),(b.filter===!1||c>0)&&d.items.push({score:c,id:e})}):g.iterator(g.items,function(a,b){d.items.push({score:1,id:b})}),e=g.getSortFunction(d,b),e&&d.items.sort(e),d.total=d.items.length,"number"==typeof b.limit&&(d.items=d.items.slice(0,b.limit)),d};var b=function(a,b){return"number"==typeof a&&"number"==typeof b?a>b?1:b>a?-1:0:(a=h(String(a||"")),b=h(String(b||"")),a>b?1:b>a?-1:0)},c=function(a){var b,c,d,e;for(b=1,c=arguments.length;c>b;b++)if(e=arguments[b])for(d in e)e.hasOwnProperty(d)&&(a[d]=e[d]);return a},d=function(a){return(a+"").replace(/^\s+|\s+$|/g,"")},e=function(a){return(a+"").replace(/([.?*+^$[\]\\(){}|-])/g,"\\$1")},f=Array.isArray||$&&$.isArray||function(a){return"[object Array]"===Object.prototype.toString.call(a)},g={a:"[aÀÁÂÃÄÅàáâãäåĀāąĄ]",c:"[cÇçćĆčČ]",d:"[dđĐďĎ]",e:"[eÈÉÊËèéêëěĚĒēęĘ]",i:"[iÌÍÎÏìíîïĪī]",l:"[lłŁ]",n:"[nÑñňŇńŃ]",o:"[oÒÓÔÕÕÖØòóôõöøŌō]",r:"[rřŘ]",s:"[sŠšśŚ]",t:"[tťŤ]",u:"[uÙÚÛÜùúûüůŮŪū]",y:"[yŸÿýÝ]",z:"[zŽžżŻźŹ]"},h=function(){var a,b,c,d,e="",f={};for(c in g)if(g.hasOwnProperty(c))for(d=g[c].substring(2,g[c].length-1),e+=d,a=0,b=d.length;b>a;a++)f[d.charAt(a)]=c;var h=new RegExp("["+e+"]","g");return function(a){return a.replace(h,function(a){return f[a]}).toLowerCase()}}();return a}),function(a,b){"function"==typeof define&&define.amd?define("microplugin",b):"object"==typeof exports?module.exports=b():a.MicroPlugin=b()}(this,function(){var a={};a.mixin=function(a){a.plugins={},a.prototype.initializePlugins=function(a){var c,d,e,f=this,g=[];if(f.plugins={names:[],settings:{},requested:{},loaded:{}},b.isArray(a))for(c=0,d=a.length;d>c;c++)"string"==typeof a[c]?g.push(a[c]):(f.plugins.settings[a[c].name]=a[c].options,g.push(a[c].name));else if(a)for(e in a)a.hasOwnProperty(e)&&(f.plugins.settings[e]=a[e],g.push(e));for(;g.length;)f.require(g.shift())},a.prototype.loadPlugin=function(b){var c=this,d=c.plugins,e=a.plugins[b];if(!a.plugins.hasOwnProperty(b))throw new Error('Unable to find "'+b+'" plugin');d.requested[b]=!0,d.loaded[b]=e.fn.apply(c,[c.plugins.settings[b]||{}]),d.names.push(b)},a.prototype.require=function(a){var b=this,c=b.plugins;if(!b.plugins.loaded.hasOwnProperty(a)){if(c.requested[a])throw new Error('Plugin has circular dependency ("'+a+'")');b.loadPlugin(a)}return c.loaded[a]},a.define=function(b,c){a.plugins[b]={name:b,fn:c}}};var b={isArray:Array.isArray||function(a){return"[object Array]"===Object.prototype.toString.call(a)}};return a}),function(a,b){"function"==typeof define&&define.amd?define("selectize",["jquery","sifter","microplugin"],b):"object"==typeof exports?module.exports=b(require("jquery"),require("sifter"),require("microplugin")):a.Selectize=b(a.jQuery,a.Sifter,a.MicroPlugin)}(this,function(a,b,c){"use strict";var d=function(a,b){if("string"!=typeof b||b.length){var c="string"==typeof b?new RegExp(b,"i"):b,d=function(a){var b=0;if(3===a.nodeType){var e=a.data.search(c);if(e>=0&&a.data.length>0){var f=a.data.match(c),g=document.createElement("span");g.className="highlight";var h=a.splitText(e),i=(h.splitText(f[0].length),h.cloneNode(!0));g.appendChild(i),h.parentNode.replaceChild(g,h),b=1}}else if(1===a.nodeType&&a.childNodes&&!/(script|style)/i.test(a.tagName))for(var j=0;j<a.childNodes.length;++j)j+=d(a.childNodes[j]);return b};return a.each(function(){d(this)})}},e=function(){};e.prototype={on:function(a,b){this._events=this._events||{},this._events[a]=this._events[a]||[],this._events[a].push(b)},off:function(a,b){var c=arguments.length;return 0===c?delete this._events:1===c?delete this._events[a]:(this._events=this._events||{},void(a in this._events!=!1&&this._events[a].splice(this._events[a].indexOf(b),1)))},trigger:function(a){if(this._events=this._events||{},a in this._events!=!1)for(var b=0;b<this._events[a].length;b++)this._events[a][b].apply(this,Array.prototype.slice.call(arguments,1))}},e.mixin=function(a){for(var b=["on","off","trigger"],c=0;c<b.length;c++)a.prototype[b[c]]=e.prototype[b[c]]};var f=/Mac/.test(navigator.userAgent),g=65,h=13,i=27,j=37,k=38,l=80,m=39,n=40,o=78,p=8,q=46,r=16,s=f?91:17,t=f?18:17,u=9,v=1,w=2,x=!/android/i.test(window.navigator.userAgent)&&!!document.createElement("form").validity,y=function(a){return"undefined"!=typeof a},z=function(a){return"undefined"==typeof a||null===a?null:"boolean"==typeof a?a?"1":"0":a+""},A=function(a){return(a+"").replace(/&/g,"&amp;").replace(/</g,"&lt;").replace(/>/g,"&gt;").replace(/"/g,"&quot;")},B=function(a){return(a+"").replace(/\$/g,"$$$$")},C={};C.before=function(a,b,c){var d=a[b];a[b]=function(){return c.apply(a,arguments),d.apply(a,arguments)}},C.after=function(a,b,c){var d=a[b];a[b]=function(){var b=d.apply(a,arguments);return c.apply(a,arguments),b}};var D=function(a){var b=!1;return function(){b||(b=!0,a.apply(this,arguments))}},E=function(a,b){var c;return function(){var d=this,e=arguments;window.clearTimeout(c),c=window.setTimeout(function(){a.apply(d,e)},b)}},F=function(a,b,c){var d,e=a.trigger,f={};a.trigger=function(){var c=arguments[0];return-1===b.indexOf(c)?e.apply(a,arguments):void(f[c]=arguments)},c.apply(a,[]),a.trigger=e;for(d in f)f.hasOwnProperty(d)&&e.apply(a,f[d])},G=function(a,b,c,d){a.on(b,c,function(b){for(var c=b.target;c&&c.parentNode!==a[0];)c=c.parentNode;return b.currentTarget=c,d.apply(this,[b])})},H=function(a){var b={};if("selectionStart"in a)b.start=a.selectionStart,b.length=a.selectionEnd-b.start;else if(document.selection){a.focus();var c=document.selection.createRange(),d=document.selection.createRange().text.length;c.moveStart("character",-a.value.length),b.start=c.text.length-d,b.length=d}return b},I=function(a,b,c){var d,e,f={};if(c)for(d=0,e=c.length;e>d;d++)f[c[d]]=a.css(c[d]);else f=a.css();b.css(f)},J=function(b,c){if(!b)return 0;var d=a("<test>").css({position:"absolute",top:-99999,left:-99999,width:"auto",padding:0,whiteSpace:"pre"}).text(b).appendTo("body");I(c,d,["letterSpacing","fontSize","fontFamily","fontWeight","textTransform"]);var e=d.width();return d.remove(),e},K=function(a){var b=null,c=function(c,d){var e,f,g,h,i,j,k,l;c=c||window.event||{},d=d||{},c.metaKey||c.altKey||(d.force||a.data("grow")!==!1)&&(e=a.val(),c.type&&"keydown"===c.type.toLowerCase()&&(f=c.keyCode,g=f>=97&&122>=f||f>=65&&90>=f||f>=48&&57>=f||32===f,f===q||f===p?(l=H(a[0]),l.length?e=e.substring(0,l.start)+e.substring(l.start+l.length):f===p&&l.start?e=e.substring(0,l.start-1)+e.substring(l.start+1):f===q&&"undefined"!=typeof l.start&&(e=e.substring(0,l.start)+e.substring(l.start+1))):g&&(j=c.shiftKey,k=String.fromCharCode(c.keyCode),k=j?k.toUpperCase():k.toLowerCase(),e+=k)),h=a.attr("placeholder"),!e&&h&&(e=h),i=J(e,a)+4,i!==b&&(b=i,a.width(i),a.triggerHandler("resize")))};a.on("keydown keyup update blur",c),c()},L=function(c,d){var e,f,g,h,i=this;h=c[0],h.selectize=i;var j=window.getComputedStyle&&window.getComputedStyle(h,null);if(g=j?j.getPropertyValue("direction"):h.currentStyle&&h.currentStyle.direction,g=g||c.parents("[dir]:first").attr("dir")||"",a.extend(i,{order:0,settings:d,$input:c,tabIndex:c.attr("tabindex")||"",tagType:"select"===h.tagName.toLowerCase()?v:w,rtl:/rtl/i.test(g),eventNS:".selectize"+ ++L.count,highlightedValue:null,isOpen:!1,isDisabled:!1,isRequired:c.is("[required]"),isInvalid:!1,isLocked:!1,isFocused:!1,isInputHidden:!1,isSetup:!1,isShiftDown:!1,isCmdDown:!1,isCtrlDown:!1,ignoreFocus:!1,ignoreBlur:!1,ignoreHover:!1,hasOptions:!1,currentResults:null,lastValue:"",caretPos:0,loading:0,loadedSearches:{},$activeOption:null,$activeItems:[],optgroups:{},options:{},userOptions:{},items:[],renderCache:{},onSearchChange:null===d.loadThrottle?i.onSearchChange:E(i.onSearchChange,d.loadThrottle)}),i.sifter=new b(this.options,{diacritics:d.diacritics}),i.settings.options){for(e=0,f=i.settings.options.length;f>e;e++)i.registerOption(i.settings.options[e]);delete i.settings.options}if(i.settings.optgroups){for(e=0,f=i.settings.optgroups.length;f>e;e++)i.registerOptionGroup(i.settings.optgroups[e]);delete i.settings.optgroups}i.settings.mode=i.settings.mode||(1===i.settings.maxItems?"single":"multi"),"boolean"!=typeof i.settings.hideSelected&&(i.settings.hideSelected="multi"===i.settings.mode),i.initializePlugins(i.settings.plugins),i.setupCallbacks(),i.setupTemplates(),i.setup()};return e.mixin(L),c.mixin(L),a.extend(L.prototype,{setup:function(){var b,c,d,e,g,h,i,j,k,l=this,m=l.settings,n=l.eventNS,o=a(window),p=a(document),q=l.$input;if(i=l.settings.mode,j=q.attr("class")||"",b=a("<div>").addClass(m.wrapperClass).addClass(j).addClass(i),c=a("<div>").addClass(m.inputClass).addClass("items").appendTo(b),d=a('<input type="text" autocomplete="off" />').appendTo(c).attr("tabindex",q.is(":disabled")?"-1":l.tabIndex),h=a(m.dropdownParent||b),e=a("<div>").addClass(m.dropdownClass).addClass(i).hide().appendTo(h),g=a("<div>").addClass(m.dropdownContentClass).appendTo(e),l.settings.copyClassesToDropdown&&e.addClass(j),b.css({width:q[0].style.width}),l.plugins.names.length&&(k="plugin-"+l.plugins.names.join(" plugin-"),b.addClass(k),e.addClass(k)),(null===m.maxItems||m.maxItems>1)&&l.tagType===v&&q.attr("multiple","multiple"),l.settings.placeholder&&d.attr("placeholder",m.placeholder),!l.settings.splitOn&&l.settings.delimiter){var u=l.settings.delimiter.replace(/[-\/\\^$*+?.()|[\]{}]/g,"\\$&");l.settings.splitOn=new RegExp("\\s*"+u+"+\\s*")}q.attr("autocorrect")&&d.attr("autocorrect",q.attr("autocorrect")),q.attr("autocapitalize")&&d.attr("autocapitalize",q.attr("autocapitalize")),l.$wrapper=b,l.$control=c,l.$control_input=d,l.$dropdown=e,l.$dropdown_content=g,e.on("mouseenter","[data-selectable]",function(){return l.onOptionHover.apply(l,arguments)}),e.on("mousedown click","[data-selectable]",function(){return l.onOptionSelect.apply(l,arguments)}),G(c,"mousedown","*:not(input)",function(){return l.onItemSelect.apply(l,arguments)}),K(d),c.on({mousedown:function(){return l.onMouseDown.apply(l,arguments)},click:function(){return l.onClick.apply(l,arguments)}}),d.on({mousedown:function(a){a.stopPropagation()},keydown:function(){return l.onKeyDown.apply(l,arguments)},keyup:function(){return l.onKeyUp.apply(l,arguments)},keypress:function(){return l.onKeyPress.apply(l,arguments)},resize:function(){l.positionDropdown.apply(l,[])},blur:function(){return l.onBlur.apply(l,arguments)},focus:function(){return l.ignoreBlur=!1,l.onFocus.apply(l,arguments)},paste:function(){return l.onPaste.apply(l,arguments)}}),p.on("keydown"+n,function(a){l.isCmdDown=a[f?"metaKey":"ctrlKey"],l.isCtrlDown=a[f?"altKey":"ctrlKey"],l.isShiftDown=a.shiftKey}),p.on("keyup"+n,function(a){a.keyCode===t&&(l.isCtrlDown=!1),a.keyCode===r&&(l.isShiftDown=!1),a.keyCode===s&&(l.isCmdDown=!1)}),p.on("mousedown"+n,function(a){if(l.isFocused){if(a.target===l.$dropdown[0]||a.target.parentNode===l.$dropdown[0])return!1;l.$control.has(a.target).length||a.target===l.$control[0]||l.blur(a.target)}}),o.on(["scroll"+n,"resize"+n].join(" "),function(){l.isOpen&&l.positionDropdown.apply(l,arguments)}),o.on("mousemove"+n,function(){l.ignoreHover=!1}),this.revertSettings={$children:q.children().detach(),tabindex:q.attr("tabindex")},q.attr("tabindex",-1).hide().after(l.$wrapper),a.isArray(m.items)&&(l.setValue(m.items),delete m.items),x&&q.on("invalid"+n,function(a){a.preventDefault(),l.isInvalid=!0,l.refreshState()}),l.updateOriginalInput(),l.refreshItems(),l.refreshState(),l.updatePlaceholder(),l.isSetup=!0,q.is(":disabled")&&l.disable(),l.on("change",this.onChange),q.data("selectize",l),q.addClass("selectized"),l.trigger("initialize"),m.preload===!0&&l.onSearchChange("")},setupTemplates:function(){var b=this,c=b.settings.labelField,d=b.settings.optgroupLabelField,e={optgroup:function(a){return'<div class="optgroup">'+a.html+"</div>"},optgroup_header:function(a,b){return'<div class="optgroup-header">'+b(a[d])+"</div>"},option:function(a,b){return'<div class="option">'+b(a[c])+"</div>"},item:function(a,b){return'<div class="item">'+b(a[c])+"</div>"},option_create:function(a,b){return'<div class="create">Add <strong>'+b(a.input)+"</strong>&hellip;</div>"}};b.settings.render=a.extend({},e,b.settings.render)},setupCallbacks:function(){var a,b,c={initialize:"onInitialize",change:"onChange",item_add:"onItemAdd",item_remove:"onItemRemove",clear:"onClear",option_add:"onOptionAdd",option_remove:"onOptionRemove",option_clear:"onOptionClear",optgroup_add:"onOptionGroupAdd",optgroup_remove:"onOptionGroupRemove",optgroup_clear:"onOptionGroupClear",dropdown_open:"onDropdownOpen",dropdown_close:"onDropdownClose",type:"onType",load:"onLoad",focus:"onFocus",blur:"onBlur"};for(a in c)c.hasOwnProperty(a)&&(b=this.settings[c[a]],b&&this.on(a,b))},onClick:function(a){var b=this;b.isFocused||(b.focus(),a.preventDefault())},onMouseDown:function(b){{var c=this,d=b.isDefaultPrevented();a(b.target)}if(c.isFocused){if(b.target!==c.$control_input[0])return"single"===c.settings.mode?c.isOpen?c.close():c.open():d||c.setActiveItem(null),!1}else d||window.setTimeout(function(){c.focus()},0)},onChange:function(){this.$input.trigger("change")},onPaste:function(b){var c=this;c.isFull()||c.isInputHidden||c.isLocked?b.preventDefault():c.settings.splitOn&&setTimeout(function(){for(var b=a.trim(c.$control_input.val()||"").split(c.settings.splitOn),d=0,e=b.length;e>d;d++)c.createItem(b[d])},0)},onKeyPress:function(a){if(this.isLocked)return a&&a.preventDefault();var b=String.fromCharCode(a.keyCode||a.which);return this.settings.create&&"multi"===this.settings.mode&&b===this.settings.delimiter?(this.createItem(),a.preventDefault(),!1):void 0},onKeyDown:function(a){var b=(a.target===this.$control_input[0],this);if(b.isLocked)return void(a.keyCode!==u&&a.preventDefault());switch(a.keyCode){case g:if(b.isCmdDown)return void b.selectAll();break;case i:return void(b.isOpen&&(a.preventDefault(),a.stopPropagation(),b.close()));case o:if(!a.ctrlKey||a.altKey)break;case n:if(!b.isOpen&&b.hasOptions)b.open();else if(b.$activeOption){b.ignoreHover=!0;var c=b.getAdjacentOption(b.$activeOption,1);c.length&&b.setActiveOption(c,!0,!0)}return void a.preventDefault();case l:if(!a.ctrlKey||a.altKey)break;case k:if(b.$activeOption){b.ignoreHover=!0;var d=b.getAdjacentOption(b.$activeOption,-1);d.length&&b.setActiveOption(d,!0,!0)}return void a.preventDefault();case h:return void(b.isOpen&&b.$activeOption&&(b.onOptionSelect({currentTarget:b.$activeOption}),a.preventDefault()));case j:return void b.advanceSelection(-1,a);case m:return void b.advanceSelection(1,a);case u:return b.settings.selectOnTab&&b.isOpen&&b.$activeOption&&(b.onOptionSelect({currentTarget:b.$activeOption}),b.isFull()||a.preventDefault()),void(b.settings.create&&b.createItem()&&a.preventDefault());case p:case q:return void b.deleteSelection(a)}return!b.isFull()&&!b.isInputHidden||(f?a.metaKey:a.ctrlKey)?void 0:void a.preventDefault()},onKeyUp:function(a){var b=this;if(b.isLocked)return a&&a.preventDefault();var c=b.$control_input.val()||"";b.lastValue!==c&&(b.lastValue=c,b.onSearchChange(c),b.refreshOptions(),b.trigger("type",c))},onSearchChange:function(a){var b=this,c=b.settings.load;c&&(b.loadedSearches.hasOwnProperty(a)||(b.loadedSearches[a]=!0,b.load(function(d){c.apply(b,[a,d])})))},onFocus:function(a){var b=this,c=b.isFocused;return b.isDisabled?(b.blur(),a&&a.preventDefault(),!1):void(b.ignoreFocus||(b.isFocused=!0,"focus"===b.settings.preload&&b.onSearchChange(""),c||b.trigger("focus"),b.$activeItems.length||(b.showInput(),b.setActiveItem(null),b.refreshOptions(!!b.settings.openOnFocus)),b.refreshState()))},onBlur:function(a,b){var c=this;if(c.isFocused&&(c.isFocused=!1,!c.ignoreFocus)){if(!c.ignoreBlur&&document.activeElement===c.$dropdown_content[0])return c.ignoreBlur=!0,void c.onFocus(a);var d=function(){c.close(),c.setTextboxValue(""),c.setActiveItem(null),c.setActiveOption(null),c.setCaret(c.items.length),c.refreshState(),(b||document.body).focus(),c.ignoreFocus=!1,c.trigger("blur")};c.ignoreFocus=!0,c.settings.create&&c.settings.createOnBlur?c.createItem(null,!1,d):d()}},onOptionHover:function(a){this.ignoreHover||this.setActiveOption(a.currentTarget,!1)},onOptionSelect:function(b){var c,d,e=this;b.preventDefault&&(b.preventDefault(),b.stopPropagation()),d=a(b.currentTarget),d.hasClass("create")?e.createItem(null,function(){e.settings.closeAfterSelect&&e.close()}):(c=d.attr("data-value"),"undefined"!=typeof c&&(e.lastQuery=null,e.setTextboxValue(""),e.addItem(c),e.settings.closeAfterSelect?e.close():!e.settings.hideSelected&&b.type&&/mouse/.test(b.type)&&e.setActiveOption(e.getOption(c))))},onItemSelect:function(a){var b=this;b.isLocked||"multi"===b.settings.mode&&(a.preventDefault(),b.setActiveItem(a.currentTarget,a))},load:function(a){var b=this,c=b.$wrapper.addClass(b.settings.loadingClass);b.loading++,a.apply(b,[function(a){b.loading=Math.max(b.loading-1,0),a&&a.length&&(b.addOption(a),b.refreshOptions(b.isFocused&&!b.isInputHidden)),b.loading||c.removeClass(b.settings.loadingClass),b.trigger("load",a)}])},setTextboxValue:function(a){var b=this.$control_input,c=b.val()!==a;c&&(b.val(a).triggerHandler("update"),this.lastValue=a)},getValue:function(){return this.tagType===v&&this.$input.attr("multiple")?this.items:this.items.join(this.settings.delimiter)},setValue:function(a,b){var c=b?[]:["change"];F(this,c,function(){this.clear(b),this.addItems(a,b)})},setActiveItem:function(b,c){var d,e,f,g,h,i,j,k,l=this;if("single"!==l.settings.mode){if(b=a(b),!b.length)return a(l.$activeItems).removeClass("active"),l.$activeItems=[],void(l.isFocused&&l.showInput());if(d=c&&c.type.toLowerCase(),"mousedown"===d&&l.isShiftDown&&l.$activeItems.length){for(k=l.$control.children(".active:last"),g=Array.prototype.indexOf.apply(l.$control[0].childNodes,[k[0]]),h=Array.prototype.indexOf.apply(l.$control[0].childNodes,[b[0]]),g>h&&(j=g,g=h,h=j),e=g;h>=e;e++)i=l.$control[0].childNodes[e],-1===l.$activeItems.indexOf(i)&&(a(i).addClass("active"),l.$activeItems.push(i));c.preventDefault()}else"mousedown"===d&&l.isCtrlDown||"keydown"===d&&this.isShiftDown?b.hasClass("active")?(f=l.$activeItems.indexOf(b[0]),l.$activeItems.splice(f,1),b.removeClass("active")):l.$activeItems.push(b.addClass("active")[0]):(a(l.$activeItems).removeClass("active"),l.$activeItems=[b.addClass("active")[0]]);l.hideInput(),this.isFocused||l.focus()}},setActiveOption:function(b,c,d){var e,f,g,h,i,j=this;j.$activeOption&&j.$activeOption.removeClass("active"),j.$activeOption=null,b=a(b),b.length&&(j.$activeOption=b.addClass("active"),(c||!y(c))&&(e=j.$dropdown_content.height(),f=j.$activeOption.outerHeight(!0),c=j.$dropdown_content.scrollTop()||0,g=j.$activeOption.offset().top-j.$dropdown_content.offset().top+c,h=g,i=g-e+f,g+f>e+c?j.$dropdown_content.stop().animate({scrollTop:i},d?j.settings.scrollDuration:0):c>g&&j.$dropdown_content.stop().animate({scrollTop:h},d?j.settings.scrollDuration:0)))},selectAll:function(){var a=this;"single"!==a.settings.mode&&(a.$activeItems=Array.prototype.slice.apply(a.$control.children(":not(input)").addClass("active")),a.$activeItems.length&&(a.hideInput(),a.close()),a.focus())},hideInput:function(){var a=this;a.setTextboxValue(""),a.$control_input.css({opacity:0,position:"absolute",left:a.rtl?1e4:-1e4}),a.isInputHidden=!0},showInput:function(){this.$control_input.css({opacity:1,position:"relative",left:0}),this.isInputHidden=!1},focus:function(){var a=this;a.isDisabled||(a.ignoreFocus=!0,a.$control_input[0].focus(),window.setTimeout(function(){a.ignoreFocus=!1,a.onFocus()},0))},blur:function(a){this.$control_input[0].blur(),this.onBlur(null,a)},getScoreFunction:function(a){return this.sifter.getScoreFunction(a,this.getSearchOptions())},getSearchOptions:function(){var a=this.settings,b=a.sortField;return"string"==typeof b&&(b=[{field:b}]),{fields:a.searchField,conjunction:a.searchConjunction,sort:b}},search:function(b){var c,d,e,f=this,g=f.settings,h=this.getSearchOptions();if(g.score&&(e=f.settings.score.apply(this,[b]),"function"!=typeof e))throw new Error('Selectize "score" setting must be a function that returns a function');if(b!==f.lastQuery?(f.lastQuery=b,d=f.sifter.search(b,a.extend(h,{score:e})),f.currentResults=d):d=a.extend(!0,{},f.currentResults),g.hideSelected)for(c=d.items.length-1;c>=0;c--)-1!==f.items.indexOf(z(d.items[c].id))&&d.items.splice(c,1);return d},refreshOptions:function(b){var c,e,f,g,h,i,j,k,l,m,n,o,p,q,r,s;"undefined"==typeof b&&(b=!0);var t=this,u=a.trim(t.$control_input.val()),v=t.search(u),w=t.$dropdown_content,x=t.$activeOption&&z(t.$activeOption.attr("data-value"));for(g=v.items.length,"number"==typeof t.settings.maxOptions&&(g=Math.min(g,t.settings.maxOptions)),h={},i=[],c=0;g>c;c++)for(j=t.options[v.items[c].id],k=t.render("option",j),l=j[t.settings.optgroupField]||"",m=a.isArray(l)?l:[l],e=0,f=m&&m.length;f>e;e++)l=m[e],t.optgroups.hasOwnProperty(l)||(l=""),h.hasOwnProperty(l)||(h[l]=[],i.push(l)),h[l].push(k);for(this.settings.lockOptgroupOrder&&i.sort(function(a,b){var c=t.optgroups[a].$order||0,d=t.optgroups[b].$order||0;return c-d}),n=[],c=0,g=i.length;g>c;c++)l=i[c],t.optgroups.hasOwnProperty(l)&&h[l].length?(o=t.render("optgroup_header",t.optgroups[l])||"",o+=h[l].join(""),n.push(t.render("optgroup",a.extend({},t.optgroups[l],{html:o})))):n.push(h[l].join(""));if(w.html(n.join("")),t.settings.highlight&&v.query.length&&v.tokens.length)for(c=0,g=v.tokens.length;g>c;c++)d(w,v.tokens[c].regex);if(!t.settings.hideSelected)for(c=0,g=t.items.length;g>c;c++)t.getOption(t.items[c]).addClass("selected");p=t.canCreate(u),p&&(w.prepend(t.render("option_create",{input:u})),s=a(w[0].childNodes[0])),t.hasOptions=v.items.length>0||p,t.hasOptions?(v.items.length>0?(r=x&&t.getOption(x),r&&r.length?q=r:"single"===t.settings.mode&&t.items.length&&(q=t.getOption(t.items[0])),q&&q.length||(q=s&&!t.settings.addPrecedence?t.getAdjacentOption(s,1):w.find("[data-selectable]:first"))):q=s,t.setActiveOption(q),b&&!t.isOpen&&t.open()):(t.setActiveOption(null),b&&t.isOpen&&t.close())},addOption:function(b){var c,d,e,f=this;if(a.isArray(b))for(c=0,d=b.length;d>c;c++)f.addOption(b[c]);else(e=f.registerOption(b))&&(f.userOptions[e]=!0,f.lastQuery=null,f.trigger("option_add",e,b))},registerOption:function(a){var b=z(a[this.settings.valueField]);return!b||this.options.hasOwnProperty(b)?!1:(a.$order=a.$order||++this.order,this.options[b]=a,b)},registerOptionGroup:function(a){var b=z(a[this.settings.optgroupValueField]);return b?(a.$order=a.$order||++this.order,this.optgroups[b]=a,b):!1},addOptionGroup:function(a,b){b[this.settings.optgroupValueField]=a,(a=this.registerOptionGroup(b))&&this.trigger("optgroup_add",a,b)},removeOptionGroup:function(a){this.optgroups.hasOwnProperty(a)&&(delete this.optgroups[a],this.renderCache={},this.trigger("optgroup_remove",a))},clearOptionGroups:function(){this.optgroups={},this.renderCache={},this.trigger("optgroup_clear")},updateOption:function(b,c){var d,e,f,g,h,i,j,k=this;if(b=z(b),f=z(c[k.settings.valueField]),null!==b&&k.options.hasOwnProperty(b)){if("string"!=typeof f)throw new Error("Value must be set in option data");j=k.options[b].$order,f!==b&&(delete k.options[b],g=k.items.indexOf(b),-1!==g&&k.items.splice(g,1,f)),c.$order=c.$order||j,k.options[f]=c,h=k.renderCache.item,i=k.renderCache.option,h&&(delete h[b],delete h[f]),i&&(delete i[b],delete i[f]),-1!==k.items.indexOf(f)&&(d=k.getItem(b),e=a(k.render("item",c)),d.hasClass("active")&&e.addClass("active"),d.replaceWith(e)),k.lastQuery=null,k.isOpen&&k.refreshOptions(!1)}},removeOption:function(a,b){var c=this;a=z(a);var d=c.renderCache.item,e=c.renderCache.option;d&&delete d[a],e&&delete e[a],delete c.userOptions[a],delete c.options[a],c.lastQuery=null,c.trigger("option_remove",a),c.removeItem(a,b)},clearOptions:function(){var a=this;a.loadedSearches={},a.userOptions={},a.renderCache={},a.options=a.sifter.items={},a.lastQuery=null,a.trigger("option_clear"),a.clear()},getOption:function(a){return this.getElementWithValue(a,this.$dropdown_content.find("[data-selectable]"))},getAdjacentOption:function(b,c){var d=this.$dropdown.find("[data-selectable]"),e=d.index(b)+c;return e>=0&&e<d.length?d.eq(e):a()},getElementWithValue:function(b,c){if(b=z(b),"undefined"!=typeof b&&null!==b)for(var d=0,e=c.length;e>d;d++)if(c[d].getAttribute("data-value")===b)return a(c[d]);return a()},getItem:function(a){return this.getElementWithValue(a,this.$control.children())},addItems:function(b,c){for(var d=a.isArray(b)?b:[b],e=0,f=d.length;f>e;e++)this.isPending=f-1>e,this.addItem(d[e],c)},addItem:function(b,c){var d=c?[]:["change"];F(this,d,function(){var d,e,f,g,h,i=this,j=i.settings.mode;return b=z(b),-1!==i.items.indexOf(b)?void("single"===j&&i.close()):void(i.options.hasOwnProperty(b)&&("single"===j&&i.clear(c),"multi"===j&&i.isFull()||(d=a(i.render("item",i.options[b])),h=i.isFull(),i.items.splice(i.caretPos,0,b),i.insertAtCaret(d),(!i.isPending||!h&&i.isFull())&&i.refreshState(),i.isSetup&&(f=i.$dropdown_content.find("[data-selectable]"),i.isPending||(e=i.getOption(b),g=i.getAdjacentOption(e,1).attr("data-value"),i.refreshOptions(i.isFocused&&"single"!==j),g&&i.setActiveOption(i.getOption(g))),!f.length||i.isFull()?i.close():i.positionDropdown(),i.updatePlaceholder(),i.trigger("item_add",b,d),i.updateOriginalInput({silent:c})))))})},removeItem:function(a,b){var c,d,e,f=this;c="object"==typeof a?a:f.getItem(a),a=z(c.attr("data-value")),d=f.items.indexOf(a),-1!==d&&(c.remove(),c.hasClass("active")&&(e=f.$activeItems.indexOf(c[0]),f.$activeItems.splice(e,1)),f.items.splice(d,1),f.lastQuery=null,!f.settings.persist&&f.userOptions.hasOwnProperty(a)&&f.removeOption(a,b),d<f.caretPos&&f.setCaret(f.caretPos-1),f.refreshState(),f.updatePlaceholder(),f.updateOriginalInput({silent:b}),f.positionDropdown(),f.trigger("item_remove",a,c))},createItem:function(b,c){var d=this,e=d.caretPos;b=b||a.trim(d.$control_input.val()||"");var f=arguments[arguments.length-1];if("function"!=typeof f&&(f=function(){}),"boolean"!=typeof c&&(c=!0),!d.canCreate(b))return f(),!1;d.lock();var g="function"==typeof d.settings.create?this.settings.create:function(a){var b={};return b[d.settings.labelField]=a,b[d.settings.valueField]=a,b},h=D(function(a){if(d.unlock(),!a||"object"!=typeof a)return f();var b=z(a[d.settings.valueField]);return"string"!=typeof b?f():(d.setTextboxValue(""),d.addOption(a),d.setCaret(e),d.addItem(b),d.refreshOptions(c&&"single"!==d.settings.mode),void f(a))}),i=g.apply(this,[b,h]);return"undefined"!=typeof i&&h(i),!0},refreshItems:function(){this.lastQuery=null,this.isSetup&&this.addItem(this.items),this.refreshState(),this.updateOriginalInput()},refreshState:function(){var a,b=this;b.isRequired&&(b.items.length&&(b.isInvalid=!1),b.$control_input.prop("required",a)),b.refreshClasses()},refreshClasses:function(){var b=this,c=b.isFull(),d=b.isLocked;b.$wrapper.toggleClass("rtl",b.rtl),b.$control.toggleClass("focus",b.isFocused).toggleClass("disabled",b.isDisabled).toggleClass("required",b.isRequired).toggleClass("invalid",b.isInvalid).toggleClass("locked",d).toggleClass("full",c).toggleClass("not-full",!c).toggleClass("input-active",b.isFocused&&!b.isInputHidden).toggleClass("dropdown-active",b.isOpen).toggleClass("has-options",!a.isEmptyObject(b.options)).toggleClass("has-items",b.items.length>0),b.$control_input.data("grow",!c&&!d)},isFull:function(){return null!==this.settings.maxItems&&this.items.length>=this.settings.maxItems},updateOriginalInput:function(a){var b,c,d,e,f=this;if(a=a||{},f.tagType===v){for(d=[],b=0,c=f.items.length;c>b;b++)e=f.options[f.items[b]][f.settings.labelField]||"",d.push('<option value="'+A(f.items[b])+'" selected="selected">'+A(e)+"</option>");d.length||this.$input.attr("multiple")||d.push('<option value="" selected="selected"></option>'),f.$input.html(d.join(""))}else f.$input.val(f.getValue()),f.$input.attr("value",f.$input.val());f.isSetup&&(a.silent||f.trigger("change",f.$input.val()))},updatePlaceholder:function(){if(this.settings.placeholder){var a=this.$control_input;this.items.length?a.removeAttr("placeholder"):a.attr("placeholder",this.settings.placeholder),a.triggerHandler("update",{force:!0})}},open:function(){var a=this;a.isLocked||a.isOpen||"multi"===a.settings.mode&&a.isFull()||(a.focus(),a.isOpen=!0,a.refreshState(),a.$dropdown.css({visibility:"hidden",display:"block"}),a.positionDropdown(),a.$dropdown.css({visibility:"visible"}),a.trigger("dropdown_open",a.$dropdown))},close:function(){var a=this,b=a.isOpen;"single"===a.settings.mode&&a.items.length&&a.hideInput(),a.isOpen=!1,a.$dropdown.hide(),a.setActiveOption(null),a.refreshState(),b&&a.trigger("dropdown_close",a.$dropdown)},positionDropdown:function(){var a=this.$control,b="body"===this.settings.dropdownParent?a.offset():a.position();b.top+=a.outerHeight(!0),this.$dropdown.css({width:a.outerWidth(),top:b.top,left:b.left})},clear:function(a){var b=this;b.items.length&&(b.$control.children(":not(input)").remove(),b.items=[],b.lastQuery=null,b.setCaret(0),b.setActiveItem(null),b.updatePlaceholder(),b.updateOriginalInput({silent:a}),b.refreshState(),b.showInput(),b.trigger("clear"))},insertAtCaret:function(b){var c=Math.min(this.caretPos,this.items.length);0===c?this.$control.prepend(b):a(this.$control[0].childNodes[c]).before(b),this.setCaret(c+1)},deleteSelection:function(b){var c,d,e,f,g,h,i,j,k,l=this;if(e=b&&b.keyCode===p?-1:1,f=H(l.$control_input[0]),l.$activeOption&&!l.settings.hideSelected&&(i=l.getAdjacentOption(l.$activeOption,-1).attr("data-value")),g=[],l.$activeItems.length){for(k=l.$control.children(".active:"+(e>0?"last":"first")),h=l.$control.children(":not(input)").index(k),e>0&&h++,c=0,d=l.$activeItems.length;d>c;c++)g.push(a(l.$activeItems[c]).attr("data-value"));
b&&(b.preventDefault(),b.stopPropagation())}else(l.isFocused||"single"===l.settings.mode)&&l.items.length&&(0>e&&0===f.start&&0===f.length?g.push(l.items[l.caretPos-1]):e>0&&f.start===l.$control_input.val().length&&g.push(l.items[l.caretPos]));if(!g.length||"function"==typeof l.settings.onDelete&&l.settings.onDelete.apply(l,[g])===!1)return!1;for("undefined"!=typeof h&&l.setCaret(h);g.length;)l.removeItem(g.pop());return l.showInput(),l.positionDropdown(),l.refreshOptions(!0),i&&(j=l.getOption(i),j.length&&l.setActiveOption(j)),!0},advanceSelection:function(a,b){var c,d,e,f,g,h,i=this;0!==a&&(i.rtl&&(a*=-1),c=a>0?"last":"first",d=H(i.$control_input[0]),i.isFocused&&!i.isInputHidden?(f=i.$control_input.val().length,g=0>a?0===d.start&&0===d.length:d.start===f,g&&!f&&i.advanceCaret(a,b)):(h=i.$control.children(".active:"+c),h.length&&(e=i.$control.children(":not(input)").index(h),i.setActiveItem(null),i.setCaret(a>0?e+1:e))))},advanceCaret:function(a,b){var c,d,e=this;0!==a&&(c=a>0?"next":"prev",e.isShiftDown?(d=e.$control_input[c](),d.length&&(e.hideInput(),e.setActiveItem(d),b&&b.preventDefault())):e.setCaret(e.caretPos+a))},setCaret:function(b){var c=this;if(b="single"===c.settings.mode?c.items.length:Math.max(0,Math.min(c.items.length,b)),!c.isPending){var d,e,f,g;for(f=c.$control.children(":not(input)"),d=0,e=f.length;e>d;d++)g=a(f[d]).detach(),b>d?c.$control_input.before(g):c.$control.append(g)}c.caretPos=b},lock:function(){this.close(),this.isLocked=!0,this.refreshState()},unlock:function(){this.isLocked=!1,this.refreshState()},disable:function(){var a=this;a.$input.prop("disabled",!0),a.$control_input.prop("disabled",!0).prop("tabindex",-1),a.isDisabled=!0,a.lock()},enable:function(){var a=this;a.$input.prop("disabled",!1),a.$control_input.prop("disabled",!1).prop("tabindex",a.tabIndex),a.isDisabled=!1,a.unlock()},destroy:function(){var b=this,c=b.eventNS,d=b.revertSettings;b.trigger("destroy"),b.off(),b.$wrapper.remove(),b.$dropdown.remove(),b.$input.html("").append(d.$children).removeAttr("tabindex").removeClass("selectized").attr({tabindex:d.tabindex}).show(),b.$control_input.removeData("grow"),b.$input.removeData("selectize"),a(window).off(c),a(document).off(c),a(document.body).off(c),delete b.$input[0].selectize},render:function(a,b){var c,d,e="",f=!1,g=this,h=/^[\t \r\n]*<([a-z][a-z0-9\-_]*(?:\:[a-z][a-z0-9\-_]*)?)/i;return("option"===a||"item"===a)&&(c=z(b[g.settings.valueField]),f=!!c),f&&(y(g.renderCache[a])||(g.renderCache[a]={}),g.renderCache[a].hasOwnProperty(c))?g.renderCache[a][c]:(e=g.settings.render[a].apply(this,[b,A]),("option"===a||"option_create"===a)&&(e=e.replace(h,"<$1 data-selectable")),"optgroup"===a&&(d=b[g.settings.optgroupValueField]||"",e=e.replace(h,'<$1 data-group="'+B(A(d))+'"')),("option"===a||"item"===a)&&(e=e.replace(h,'<$1 data-value="'+B(A(c||""))+'"')),f&&(g.renderCache[a][c]=e),e)},clearCache:function(a){var b=this;"undefined"==typeof a?b.renderCache={}:delete b.renderCache[a]},canCreate:function(a){var b=this;if(!b.settings.create)return!1;var c=b.settings.createFilter;return!(!a.length||"function"==typeof c&&!c.apply(b,[a])||"string"==typeof c&&!new RegExp(c).test(a)||c instanceof RegExp&&!c.test(a))}}),L.count=0,L.defaults={options:[],optgroups:[],plugins:[],delimiter:",",splitOn:null,persist:!0,diacritics:!0,create:!1,createOnBlur:!1,createFilter:null,highlight:!0,openOnFocus:!0,maxOptions:1e3,maxItems:null,hideSelected:null,addPrecedence:!1,selectOnTab:!1,preload:!1,allowEmptyOption:!1,closeAfterSelect:!1,scrollDuration:60,loadThrottle:300,loadingClass:"loading",dataAttr:"data-data",optgroupField:"optgroup",valueField:"value",labelField:"text",optgroupLabelField:"label",optgroupValueField:"value",lockOptgroupOrder:!1,sortField:"$order",searchField:["text"],searchConjunction:"and",mode:null,wrapperClass:"selectize-control",inputClass:"selectize-input",dropdownClass:"selectize-dropdown",dropdownContentClass:"selectize-dropdown-content",dropdownParent:null,copyClassesToDropdown:!0,render:{}},a.fn.selectize=function(b){var c=a.fn.selectize.defaults,d=a.extend({},c,b),e=d.dataAttr,f=d.labelField,g=d.valueField,h=d.optgroupField,i=d.optgroupLabelField,j=d.optgroupValueField,k=function(b,c){var h,i,j,k,l=b.attr(e);if(l)for(c.options=JSON.parse(l),h=0,i=c.options.length;i>h;h++)c.items.push(c.options[h][g]);else{var m=a.trim(b.val()||"");if(!d.allowEmptyOption&&!m.length)return;for(j=m.split(d.delimiter),h=0,i=j.length;i>h;h++)k={},k[f]=j[h],k[g]=j[h],c.options.push(k);c.items=j}},l=function(b,c){var k,l,m,n,o=c.options,p={},q=function(a){var b=e&&a.attr(e);return"string"==typeof b&&b.length?JSON.parse(b):null},r=function(b,e){b=a(b);var i=z(b.attr("value"));if(i||d.allowEmptyOption)if(p.hasOwnProperty(i)){if(e){var j=p[i][h];j?a.isArray(j)?j.push(e):p[i][h]=[j,e]:p[i][h]=e}}else{var k=q(b)||{};k[f]=k[f]||b.text(),k[g]=k[g]||i,k[h]=k[h]||e,p[i]=k,o.push(k),b.is(":selected")&&c.items.push(i)}},s=function(b){var d,e,f,g,h;for(b=a(b),f=b.attr("label"),f&&(g=q(b)||{},g[i]=f,g[j]=f,c.optgroups.push(g)),h=a("option",b),d=0,e=h.length;e>d;d++)r(h[d],f)};for(c.maxItems=b.attr("multiple")?null:1,n=b.children(),k=0,l=n.length;l>k;k++)m=n[k].tagName.toLowerCase(),"optgroup"===m?s(n[k]):"option"===m&&r(n[k])};return this.each(function(){if(!this.selectize){var e,f=a(this),g=this.tagName.toLowerCase(),h=f.attr("placeholder")||f.attr("data-placeholder");h||d.allowEmptyOption||(h=f.children('option[value=""]').text());var i={placeholder:h,options:[],optgroups:[],items:[]};"select"===g?l(f,i):k(f,i),e=new L(f,a.extend(!0,{},c,i,b))}})},a.fn.selectize.defaults=L.defaults,a.fn.selectize.support={validity:x},L.define("drag_drop",function(){if(!a.fn.sortable)throw new Error('The "drag_drop" plugin requires jQuery UI "sortable".');if("multi"===this.settings.mode){var b=this;b.lock=function(){var a=b.lock;return function(){var c=b.$control.data("sortable");return c&&c.disable(),a.apply(b,arguments)}}(),b.unlock=function(){var a=b.unlock;return function(){var c=b.$control.data("sortable");return c&&c.enable(),a.apply(b,arguments)}}(),b.setup=function(){var c=b.setup;return function(){c.apply(this,arguments);var d=b.$control.sortable({items:"[data-value]",forcePlaceholderSize:!0,disabled:b.isLocked,start:function(a,b){b.placeholder.css("width",b.helper.css("width")),d.css({overflow:"visible"})},stop:function(){d.css({overflow:"hidden"});var c=b.$activeItems?b.$activeItems.slice():null,e=[];d.children("[data-value]").each(function(){e.push(a(this).attr("data-value"))}),b.setValue(e),b.setActiveItem(c)}})}}()}}),L.define("dropdown_header",function(b){var c=this;b=a.extend({title:"Untitled",headerClass:"selectize-dropdown-header",titleRowClass:"selectize-dropdown-header-title",labelClass:"selectize-dropdown-header-label",closeClass:"selectize-dropdown-header-close",html:function(a){return'<div class="'+a.headerClass+'"><div class="'+a.titleRowClass+'"><span class="'+a.labelClass+'">'+a.title+'</span><a href="javascript:void(0)" class="'+a.closeClass+'">&times;</a></div></div>'}},b),c.setup=function(){var d=c.setup;return function(){d.apply(c,arguments),c.$dropdown_header=a(b.html(b)),c.$dropdown.prepend(c.$dropdown_header)}}()}),L.define("optgroup_columns",function(b){var c=this;b=a.extend({equalizeWidth:!0,equalizeHeight:!0},b),this.getAdjacentOption=function(b,c){var d=b.closest("[data-group]").find("[data-selectable]"),e=d.index(b)+c;return e>=0&&e<d.length?d.eq(e):a()},this.onKeyDown=function(){var a=c.onKeyDown;return function(b){var d,e,f,g;return!this.isOpen||b.keyCode!==j&&b.keyCode!==m?a.apply(this,arguments):(c.ignoreHover=!0,g=this.$activeOption.closest("[data-group]"),d=g.find("[data-selectable]").index(this.$activeOption),g=b.keyCode===j?g.prev("[data-group]"):g.next("[data-group]"),f=g.find("[data-selectable]"),e=f.eq(Math.min(f.length-1,d)),void(e.length&&this.setActiveOption(e)))}}();var d=function(){var a,b=d.width,c=document;return"undefined"==typeof b&&(a=c.createElement("div"),a.innerHTML='<div style="width:50px;height:50px;position:absolute;left:-50px;top:-50px;overflow:auto;"><div style="width:1px;height:100px;"></div></div>',a=a.firstChild,c.body.appendChild(a),b=d.width=a.offsetWidth-a.clientWidth,c.body.removeChild(a)),b},e=function(){var e,f,g,h,i,j,k;if(k=a("[data-group]",c.$dropdown_content),f=k.length,f&&c.$dropdown_content.width()){if(b.equalizeHeight){for(g=0,e=0;f>e;e++)g=Math.max(g,k.eq(e).height());k.css({height:g})}b.equalizeWidth&&(j=c.$dropdown_content.innerWidth()-d(),h=Math.round(j/f),k.css({width:h}),f>1&&(i=j-h*(f-1),k.eq(f-1).css({width:i})))}};(b.equalizeHeight||b.equalizeWidth)&&(C.after(this,"positionDropdown",e),C.after(this,"refreshOptions",e))}),L.define("remove_button",function(b){if("single"!==this.settings.mode){b=a.extend({label:"&times;",title:"Remove",className:"remove",append:!0},b);var c=this,d='<a href="javascript:void(0)" class="'+b.className+'" tabindex="-1" title="'+A(b.title)+'">'+b.label+"</a>",e=function(a,b){var c=a.search(/(<\/[^>]+>\s*)$/);return a.substring(0,c)+b+a.substring(c)};this.setup=function(){var f=c.setup;return function(){if(b.append){var g=c.settings.render.item;c.settings.render.item=function(){return e(g.apply(this,arguments),d)}}f.apply(this,arguments),this.$control.on("click","."+b.className,function(b){if(b.preventDefault(),!c.isLocked){var d=a(b.currentTarget).parent();c.setActiveItem(d),c.deleteSelection()&&c.setCaret(c.items.length)}})}}()}}),L.define("restore_on_backspace",function(a){var b=this;a.text=a.text||function(a){return a[this.settings.labelField]},this.onKeyDown=function(){var c=b.onKeyDown;return function(b){var d,e;return b.keyCode===p&&""===this.$control_input.val()&&!this.$activeItems.length&&(d=this.caretPos-1,d>=0&&d<this.items.length)?(e=this.options[this.items[d]],this.deleteSelection(b)&&(this.setTextboxValue(a.text.apply(this,[e])),this.refreshOptions(!0)),void b.preventDefault()):c.apply(this,arguments)}}()}),L});
```
