## supplement_file1 for "GENETH’OFF: a flexible workflow for genome-wide profiling of CRISPR/Cas off-targets": VEGFA_s1_K562_pos_offtargets.html

table output


| Position | cut offset | Alignment | UMIs | UMIs (%) | Reads | Edits crRNA | Edits pam | Symbol | Feature | Predicted | PCR | Oncogene |
| --- | --- | --- | --- | --- | --- | --- | --- | --- | --- | --- | --- | --- |
| chr6:43769560 | -2 | gRNA: GGGTGGGGGGAGTTTGCTCC NGG  gDNA: .................... T.. | 156 | 20.6 | 3097 | 0 | 0 |  |  | yes | + |  |
| chr15:65345199 | -1 | gRNA: GGGTGGGGGGAGTTTGCTCC NGG  gDNA: ..A...A............. T.. | 50 | 6.6 | 941 | 2 | 0 | IGDCC3 | intron | yes | + |  |
| chr17:41640076 | -1 | gRNA: GGGTGGGGGGAGTTTGCTCC NGG  gDNA: TA....A.....C....... T.. | 40 | 5.3 | 793 | 4 | 0 | KRT42P | exon | yes | + |  |
| chr12:131205653 | 0 | gRNA: GGGTGGGGGGAGTTTGCTCC NGG  gDNA: ...A...T............ T.. | 15 | 2.0 | 153 | 2 | 0 | LINC01257 | intron | yes | + |  |
| chr10:13976192 | -21 | gRNA: GGGTGGGGGGAGTTTGCTCC NGG  gDNA: A.A.....T.....-...GA GT. | 11 | 1.4 | 214 | 6 | 1 | FRMD4A | intron | no | + |  |
| chr1:14157983 | 9 | gRNA: GGGTGGGGGGAGTTTG-CTCC NGG  gDNA: T......A..T.A...G...A T.A | 10 | 1.3 | 175 | 6 | 1 | KAZN | intron | no | + |  |
| chr22:45277485 | 16 | gRNA: GGGTGGGGGGAGTTTGCTCC NGG  gDNA: A......A...--.....TG A.. | 10 | 1.3 | 162 | 6 | 0 | ENSG00000301739 | intron | yes | + |  |
| chr1:30825017 | 28 | gRNA: GGGTGGGGGGAGTTTGCTCC NGG  gDNA: ..........C-....TGAG G.C | 9 | 1.2 | 187 | 6 | 1 | ENSG00000229607   LINC01778 | intron  intron | yes | + |  |
| chr9:89687061 | -25 | gRNA: GGGTGGGGGGAGTTTGCTCC NGG  gDNA: ...GCC..A......A..T. CA. | 9 | 1.2 | 52 | 6 | 1 | LINC03062 | intron | no | + |  |
| chr9:122375678 | 18 | gRNA: GGGTGGGGGGAGTTTGCTCC NGG  gDNA: ...A..A......G..GA.A TTT | 9 | 1.2 | 265 | 6 | 2 | PTGS1 | intron | no | + |  |
| chr10:71907038 | 27 | gRNA: GGGTGGGGGGA--GTTTGCTCC NGG  gDNA: A......A...TT.....AG.. TT. | 7 | 0.9 | 70 | 6 | 1 |  |  | no | + |  |
| chr16:9930141 | -27 | gRNA: GGGTGGGGGGAGTTTGCTCC NGG  gDNA: .A.....CT...C.....GG GT. | 7 | 0.9 | 44 | 6 | 1 | GRIN2A | intron | yes | + |  |
| chr17:35942717 | 0 | gRNA: GGGTGGGGGGAGTTTGCTCC NGG  gDNA: ..A......A...---.... T.. | 7 | 0.9 | 130 | 5 | 0 | LYZL6   ENSG00000270240 | intron  intron | yes | + |  |
| chr18:13409285 | -7 | gRNA: GGGTGGGGGGAGTTTGCTCC NGG  gDNA: .......--T..GC....G. CTC | 7 | 0.9 | 433 | 6 | 2 | LDLRAD4 | intron | no | + |  |
| chr21:29349589 | -47 | gRNA: GGGTGGGGGGAGTTTGCTCC NGG  gDNA: .T....A....A...-..G. ... | 7 | 0.9 | 336 | 5 | 3 | BACH1 | intron | no | + |  |
| chr21:46302287 | -20 | gRNA: GGGTGGGGGGAGTTTGCTCC NGG  gDNA: ...GC.......GAGC.... TCT | 7 | 0.9 | 273 | 6 | 2 | C21orf58 | intron | no | + |  |
| chr5:57735044 | 0 | gRNA: GGGTGGGGGGAGTTTGCTCC NGG  gDNA: CTC..A.............T G.. | 7 | 0.9 | 44 | 5 | 0 | ENSG00000287709 | intron | yes | + |  |
| chr6:44218666 | -43 | gRNA: GGGTGGGGGGAGTTTGCTCC NGG  gDNA: ....T...T.....GC..TA ... | 7 | 0.9 | 292 | 6 | 3 |  |  | no | + |  |
| chr9:19810161 | 31 | gRNA: GGGTGGGGGGAGTTTGCTCC NGG  gDNA: AA......A...G.G.T... A.. | 7 | 0.9 | 489 | 6 | 0 | ENSG00000286685 | intron | yes | + |  |
| chr1:24302880 | 34 | gRNA: GGGTGGGGGGAGTTTGCTCC NGG  gDNA: .....T..T.-..GG..G.. T.T | 6 | 0.8 | 35 | 6 | 1 | GRHL3 | intron | no | + |  |
| chr10:124743674 | -31 | gRNA: GGGTGGGGGGAGTT-TGCTCC NGG  gDNA: ...C.C.....C.GC...G.. C.. | 6 | 0.8 | 145 | 6 | 0 | ENSG00000258539   FAM53B | intron  intron | yes | + |  |
| chr11:47715138 | -2 | gRNA: GGGTGGGGGGAGTTTGCTCC NGG  gDNA: T....A....-....C..AA G.. | 6 | 0.8 | 89 | 6 | 0 | AGBL2 | exon | yes | + |  |
| chr11:81737472 | 14 | gRNA: GGGTGGGGGGAGTTTGCTCC NGG  gDNA: ..A.....T.GC.G....G. A.. | 6 | 0.8 | 38 | 6 | 0 |  |  | yes | + |  |
| chr16:22265092 | 45 | gRNA: GGGTGGGGG-GAGTTTGCTCC NGG  gDNA: ..A.....CT...C.-....T ... | 6 | 0.8 | 78 | 6 | 3 | EEF2K   ENSG00000305400 | exon  intron | no | + |  |
| chr4:876621 | -21 | gRNA: GGGTGGGGGGAGTTTGCTCC NGG  gDNA: .....--TC.T........T CTT | 6 | 0.8 | 105 | 6 | 2 | GAK | exon | no | + |  |
| chr7:73377346 | -42 | gRNA: GGGTGGGGGGAGTTTGCTCC NGG  gDNA: ...GT......A..A.GG.. ... | 6 | 0.8 | 308 | 6 | 3 |  |  | no | + |  |
| chr1:21627095 | -3 | gRNA: GGGTGGGGGGAGTTTGCTCC NGG  gDNA: .C.A...T.A...CA..... TAT | 5 | 0.7 | 167 | 6 | 2 | RAP1GAP | intron | no | + |  |
| chr1:166973608 | -12 | gRNA: GGGTGGGGGGAGTTTGCTCC NGG  gDNA: ...G........G--.G... CT. | 5 | 0.7 | 36 | 5 | 1 | ILDR2 | intron | no | + |  |
| chr1:189931239 | 7 | gRNA: GGGTGGGGGGAGTTTGCTCC NGG  gDNA: ..C....T.C...G-....A T.C | 5 | 0.7 | 89 | 6 | 1 |  |  | no | + |  |
| chr1:231287016 | -16 | gRNA: GGGTGGGGGGAGTTTGCTCC NGG  gDNA: .....T..T.-.CA...A.. T.T | 5 | 0.7 | 47 | 6 | 1 |  |  | no | + |  |
| chr10:55019722 | 19 | gRNA: GGGTGGGGGGAGTTTGCTCC NGG  gDNA: ....AT.A-...G......T ACT | 5 | 0.7 | 123 | 6 | 2 | PCDH15 | intron | no | + |  |
| chr17:49240177 | -1 | gRNA: GGGTGGGGGGAGTTTGCTCC NGG  gDNA: T......-....C....... A.. | 5 | 0.7 | 116 | 3 | 0 | FLJ40194 | intron | yes | + |  |
| chr17:76001112 | 18 | gRNA: GGGTGGGGGGAGTTTGCTCC NGG  gDNA: .......A.-...G..GG.. CT. | 5 | 0.7 | 217 | 5 | 1 | TEN1-CDK3   CDK3 | exon  intron | no | + |  |
| chr2:32778459 | 41 | gRNA: GGGTGGGGGGAGTTTGCTCC NGG  gDNA: ....ACA...-....-...T ... | 5 | 0.7 | 37 | 6 | 3 | TTC27 | intron | no | + |  |
| chr2:156799811 | 8 | gRNA: GGGTGGGGG-GAGTTTGCTCC NGG  gDNA: .....ACA.A.........TT TCA | 5 | 0.7 | 36 | 6 | 2 | ENSG00000299347   ENSG00000287048 | intron  intron | no | + |  |
| chr22:34956016 | -12 | gRNA: GGGTGGGGGGAGTTTGCTCC NGG  gDNA: .T.CC.T..T........G. ACC | 5 | 0.7 | 35 | 6 | 2 | LINC02885 | intron | no | + |  |
| chr3:13014624 | -45 | gRNA: GGGTGGGGGGAGTTTGCTCC NGG  gDNA: .........AG.C...G--. ... | 5 | 0.7 | 123 | 6 | 3 | IQSEC1 | intron | yes | + |  |
| chr6:43717957 | -14 | gRNA: GGGTGGGGGGAGTTTGCTCC NGG  gDNA: .......T....GC.AT..T A.C | 5 | 0.7 | 183 | 6 | 1 |  |  | no | + |  |
| chr9:123540964 | 7 | gRNA: GGGTGGGGGGAGTTTGCTCC NGG  gDNA: .....A..T...G.A.G.G. CAA | 5 | 0.7 | 112 | 6 | 2 | DENND1A | intron | no | + |  |
| chr1:6210723 | -46 | gRNA: GGGTGGG--GGGAGTTTGCTCC NGG  gDNA: ..A....AT...G.....-... ... | 4 | 0.5 | 339 | 5 | 3 | RNF207 | exon | yes | + |  |
| chr1:84984911 | -7 | gRNA: GGGTGGGGGGAGTTTGCTCC NGG  gDNA: A......A...--......A A.T | 4 | 0.5 | 39 | 5 | 1 | MCOLN2 | intron | no | + |  |
| chr1:228246166 | 33 | gRNA: GGGTGGGGGGAGTTTGCTCC NGG  gDNA: ...A.T.......A-...G. TAA | 4 | 0.5 | 185 | 5 | 2 | ENSG00000269934   OBSCN | intron  intron | no | + |  |
| chr1:231648798 | 25 | gRNA: GGGTGGGGGGAGTTTGCTCC NGG  gDNA: ...A...T.T.-.G..-... A.. | 4 | 0.5 | 12 | 6 | 0 | TSNAX-DISC1   DISC1 | intron  intron | yes | + |  |
| chr10:49829330 | -28 | gRNA: GGGTGGGGGGA--GTTTGCTCC NGG  gDNA: A......A...AT.....AG.. CT. | 4 | 0.5 | 149 | 6 | 1 | PARG | intron | no | + |  |
| chr12:1878910 | 0 | gRNA: GGGTGGGGGGAGTTTGCTCC NGG  gDNA: C..G..A............. T.. | 4 | 0.5 | 212 | 3 | 0 | CACNA2D4 | exon | yes | + |  |
| chr12:100996418 | -46 | gRNA: GGGTGGGGGGAGTTTGCTCC NGG  gDNA: ..C....T.C...G-....A ... | 4 | 0.5 | 181 | 6 | 3 | ANO4 | intron | no | + |  |
| chr16:85445325 | 42 | gRNA: GGGTGGGGGGAGTTTGCTCC NGG  gDNA: .A........-...C...GG ... | 4 | 0.5 | 178 | 5 | 3 | GSE1 | intron | no | + |  |
| chr2:25610994 | -21 | gRNA: GGGTGGGGGGAGTTTGCTCC NGG  gDNA: ..C....T.C...G-....A C.C | 4 | 0.5 | 225 | 6 | 1 | DTNB | intron | no | + |  |
| chr2:115410380 | -33 | gRNA: GGGTGGGGGGAGTTTGCTCC NGG  gDNA: .....CTT..TT....T... T.C | 4 | 0.5 | 33 | 6 | 1 | DPP10 | intron | no | + |  |
| chr2:241362558 | 32 | gRNA: GGGTGGGGGGAGTTTGCTCC NGG  gDNA: CA...A..C...A....G.. ACT | 4 | 0.5 | 261 | 6 | 2 | FARP2   ENSG00000288080 | intron  intron | no | + |  |
| chr4:135865150 | -16 | gRNA: GGGTGGGGGGAGTTTGCTCC NGG  gDNA: ...A..TT.........--. A.T | 4 | 0.5 | 30 | 5 | 1 |  |  | no | + |  |
| chr5:178094677 | 33 | gRNA: GGGTGGGGGGAGTTTGCTCC NGG  gDNA: ..A....A..G..G..A.A. GA. | 4 | 0.5 | 111 | 6 | 1 | ENSG00000307551 | intron | no | + |  |
| chr6:2191835 | -3 | gRNA: GGGTGGGGGGAGTTTGCTCC NGG  gDNA: ......AA..G.G.A.G... CT. | 4 | 0.5 | 25 | 6 | 1 | GMDS | intron | no | + |  |
| chr6:43455921 | -26 | gRNA: GGGTGGGGGGAGTTTGC-TCC NGG  gDNA: ...C....A.C.A....C.G. GT. | 4 | 0.5 | 14 | 6 | 1 | DLK2 | intron | no | + |  |
| chr6:133253569 | 7 | gRNA: GGGTGGGGGGAGTTTGCTCC NGG  gDNA: ......T.A..A.A...--. A.. | 4 | 0.5 | 13 | 6 | 0 | EYA4 | intron | yes | + |  |
| chr6:134296851 | 28 | gRNA: GGGTGGGGGGA-GTTTGCTCC NGG  gDNA: AC...C.....A....A..G. TT. | 4 | 0.5 | 112 | 6 | 1 | SGK1   ENSG00000286887 | intron  exon | no | + |  |
| chr6:165187045 | -45 | gRNA: GGGTGGGGGGAGTTTGCTCC NGG  gDNA: ...A.CA......GA..A.. ... | 4 | 0.5 | 59 | 6 | 3 |  |  | no | + |  |
| chr7:158261576 | 45 | gRNA: GGGTGGGGGGAGTTTGCTCC NGG  gDNA: ......CT.....---.C.. ... | 4 | 0.5 | 42 | 6 | 3 | PTPRN2 | intron | yes | + |  |
| chr9:96304692 | -29 | gRNA: GGGTGGGGGGAGTTTGCTCC NGG  gDNA: T......T....A.G..--. A.A | 4 | 0.5 | 66 | 6 | 1 | ENSG00000285269 | intron | no | + |  |
| chr1:175394052 | 33 | gRNA: GGGTGGGGGGAGTTTGCTCC NGG  gDNA: .......TT..T.C...A.. T.. | 3 | 0.4 | 49 | 5 | 0 | TNR | intron | yes | + |  |
| chr1:192419044 | -4 | gRNA: GGGTGGGGGGAGTTTGCTCC NGG  gDNA: ..T...TCA..-....G... ACA | 3 | 0.4 | 10 | 6 | 2 | ENSG00000285280 | intron | no | + |  |
| chr1:230845299 | 16 | gRNA: GGGTGGGGGGAGTTTGCTCC NGG  gDNA: A......A...--.....TG A.C | 3 | 0.4 | 118 | 6 | 1 | C1orf198 | intron | no | + |  |
| chr10:132763089 | -3 | gRNA: GGGTGG-GGGGAGTT-TGCTCC NGG  gDNA: T....CA......A.C..A... CCT | 3 | 0.4 | 11 | 6 | 2 | INPP5A | intron | no | + |  |
| chr11:78257246 | -9 | gRNA: GGGTGGGGGGAGTTTGCTCC NGG  gDNA: ..T...C........-..GG AAA | 3 | 0.4 | 21 | 5 | 2 | GAB2 | intron | no | + |  |
| chr12:103969796 | 10 | gRNA: GGGTGGGGGGAGTTTG-CTCC NGG  gDNA: ....TCT.C-......T.... AAA | 3 | 0.4 | 19 | 6 | 2 | TDG | intron | no | + |  |
| chr12:112708586 | -38 | gRNA: GGGTGGGGGGAG--TTTGCTCC NGG  gDNA: ....T.T...TCCA........ TCC | 3 | 0.4 | 31 | 6 | 2 | RPH3A | intron | no | + |  |
| chr13:52053034 | -6 | gRNA: GGGTGGGGGGAGTTTGCTCC NGG  gDNA: ..C....T.C...G-....A CAC | 3 | 0.4 | 122 | 6 | 2 | NEK5   ALG11 | intron  intron | no | + |  |
| chr14:100213968 | 22 | gRNA: GGGTGGGGGGAGTTTGCTCC NGG  gDNA: ..C.....C...G--..G.. GT. | 3 | 0.4 | 185 | 6 | 1 | YY1-DT | exon | no | + |  |
| chr15:89009207 | -26 | gRNA: GGGTGGGGGGAGTTTGCTCC NGG  gDNA: .....A......CCCT.... T.C | 3 | 0.4 | 97 | 5 | 1 | ENSG00000303391 | intron | no | + |  |
| chr17:49160228 | 20 | gRNA: GGGTGGGGGGAGTTTGCTCC NGG  gDNA: .......--T.T...TA... T.. | 3 | 0.4 | 73 | 6 | 0 | B4GALNT2 | intron | yes | + |  |
| chr17:63981528 | -1 | gRNA: GGGTG-GGGGGAGTTTGCTCC NGG  gDNA: ..T..CA.CC...A....... ATT | 3 | 0.4 | 22 | 6 | 2 | ENSG00000263489 | intron | no | + |  |
| chr19:47261428 | -14 | gRNA: GGGTGGGGGGAGTTTGCTCC NGG  gDNA: .......---...C..-..T A.C | 3 | 0.4 | 123 | 6 | 1 | CCDC9 | intron | no | + |  |
| chr2:173990948 | 29 | gRNA: GGGTGGGGGGAGTTTGCTCC NGG  gDNA: A......A...--.....TG A.C | 3 | 0.4 | 22 | 6 | 1 |  |  | no | + |  |
| chr22:24231972 | -37 | gRNA: GGGTGGGGGGAGTTTGCTCC NGG  gDNA: ..T..A...C..---..... TTC | 3 | 0.4 | 52 | 6 | 2 | GGT5 | intron | no | + |  |
| chr22:41077858 | -36 | gRNA: GGGTGGGGGGAGTTTGCTCC NGG  gDNA: T.......T.T.G.G....A T.C | 3 | 0.4 | 9 | 6 | 1 |  |  | no | + |  |
| chr3:50624658 | 19 | gRNA: GGGTGGGGGGAGTTTGCTCC NGG  gDNA: ...GAA.T......A.A... CTC | 3 | 0.4 | 26 | 6 | 2 | MAPKAPK3 | intron | no | + |  |
| chr3:124860650 | 12 | gRNA: GGGTGGGGGGAGT-TTGCTCC NGG  gDNA: .A.........A.C...T.TT TCA | 3 | 0.4 | 32 | 6 | 2 | ITGB5 | intron | no | + |  |
| chr4:24586152 | 26 | gRNA: GGGTGGGGGGAGTTTGCTCC NGG  gDNA: ..C....T.C...G-....A C.C | 3 | 0.4 | 50 | 6 | 1 |  |  | no | + |  |
| chr4:36142732 | -4 | gRNA: GGGTGGGGGGAGTTTGCTCC NGG  gDNA: ....TA.....A....-..A G.. | 3 | 0.4 | 22 | 5 | 0 | ARAP2 | intron | yes | + |  |
| chr4:54580406 | 2 | gRNA: GGGTGGGGGGAGTTTGCTCC NGG  gDNA: .T...T....--..G.G... AT. | 3 | 0.4 | 19 | 6 | 1 |  |  | no | + |  |
| chr4:64093257 | 21 | gRNA: GGGTGGGGGGAGTTTGCTCC NGG  gDNA: ...CC...C.C...G....A C.C | 3 | 0.4 | 18 | 6 | 1 |  |  | no | + |  |
| chr5:3311124 | -18 | gRNA: GGGTGGGGGGAGTTTGCTCC NGG  gDNA: ..C.....T..C.C...CA. T.C | 3 | 0.4 | 187 | 6 | 1 |  |  | no | + |  |
| chr5:32945168 | 0 | gRNA: GGGTGGGGGGAGTTTGCTCC NGG  gDNA: .C........T......... C.. | 3 | 0.4 | 56 | 2 | 0 | ENSG00000250697 | intron | yes | + |  |
| chr5:177566619 | 18 | gRNA: GGGTGGGGGGAGTTTGCTCC NGG  gDNA: .....T..T.-..G...A.. T.T | 3 | 0.4 | 31 | 5 | 1 | FAM193B-DT | intron | no | + |  |
| chr6:5629157 | 5 | gRNA: GGGTGGGGGGAGTTTGCTCC NGG  gDNA: .....A..C-.......--. ATC | 3 | 0.4 | 108 | 5 | 2 | FARS2 | intron | no | + |  |
| chr6:37206684 | 22 | gRNA: GGGTGGGGGGAGTTTGCTCC NGG  gDNA: ..C....T.C...G-....A C.C | 3 | 0.4 | 375 | 6 | 1 |  |  | no | + |  |
| chr7:130031281 | -14 | gRNA: GGGTGGGGGGAGTTTGCTCC NGG  gDNA: T....A.CT...A....G.. ATT | 3 | 0.4 | 84 | 6 | 2 | ZC3HC1 | intron | no | + |  |
| chrX:94070434 | -19 | gRNA: GGGTGGGGGGAGTTTGCTCC NGG  gDNA: ..A.....T.-....T..TA TTA | 3 | 0.4 | 14 | 6 | 2 |  |  | no | + |  |
| chrX:130567144 | 19 | gRNA: GGGTGGGGGGAGTTTGCTCC NGG  gDNA: TA.A..T....C....-... TTA | 3 | 0.4 | 17 | 6 | 2 |  |  | no | + |  |
| chr1:185742 | -7 | gRNA: GGGTGGG-GGGAGTTTGCTCC NGG  gDNA: A.....CT.....G.G..... TCA | 2 | 0.3 | 11 | 5 | 2 | WASH9P   WASH9P   ENSG00000310528 | intron  exon  intron | no | + |  |
| chr1:10412927 | -10 | gRNA: GGGTGGGGGGAGTT-TGCTCC NGG  gDNA: .CC.....--.C..C...... T.C | 2 | 0.3 | 82 | 6 | 1 | PGD | exon | no | + |  |
| chr1:21372929 | 2 | gRNA: GGGTGGGGGGAGTTTGCTCC NGG  gDNA: ..C...A.C...G.G....A T.C | 2 | 0.3 | 21 | 6 | 1 |  |  | no | + |  |
| chr1:170284678 | -17 | gRNA: GGGTGGGGG-GAGTTTGCTCC NGG  gDNA: CA.......ACG.......T. CTC | 2 | 0.3 | 78 | 6 | 2 | LINC01681 | intron | no | + |  |
| chr1:175462887 | -13 | gRNA: GGGTGGGGGGA--GTTTGCTCC NGG  gDNA: ...A....CA.AA......A.. CT. | 2 | 0.3 | 14 | 6 | 1 | TNR | intron | no | + |  |
| chr11:71072298 | 1 | gRNA: GGGTGGGGGGAGTTTGCTCC NGG  gDNA: .C........CTG...G... C.C | 2 | 0.3 | 74 | 5 | 1 | SHANK2 | intron | no | + |  |
| chr11:107273019 | -16 | gRNA: GGGTGGGGGGAGTTTGCTCC NGG  gDNA: .....C....-.C.AA.G.. T.T | 2 | 0.3 | 10 | 6 | 1 |  |  | no | + |  |
| chr11:124533762 | -19 | gRNA: GGGTGGGGGGAGTTTGCTCC NGG  gDNA: ...A...CA..T...C...A C.A | 2 | 0.3 | 7 | 6 | 1 |  |  | no | + |  |
| chr11:129914287 | -17 | gRNA: GGGTGGGGGGAGTTTGCTCC NGG  gDNA: ..C....C.C...G-....A T.C | 2 | 0.3 | 7 | 6 | 1 | PRDM10 | intron | no | + |  |
| chr13:51800484 | 35 | gRNA: GGGTGGGGGGAGTTTGCTCC NGG  gDNA: .C.....CA....C...C.T CT. | 2 | 0.3 | 55 | 6 | 1 | ENSG00000285444   DHRS12 | intron  intron | yes | + |  |
| chr14:23425565 | 4 | gRNA: GGGTGGGGGGAGT-TTGCTCC NGG  gDNA: ..A..A.......GG....AG AT. | 2 | 0.3 | 39 | 6 | 1 | MYH7 | intron | no | + |  |
