## supplement_file1 for "GENETH’OFF: a flexible workflow for genome-wide profiling of CRISPR/Cas off-targets": test_dataset_report.html

Off-targets analysis report


### Off-targets analysis report

### Therapeutic Gene Editing - GENETHON, INSERM U951, Paris-Saclay University, Evry, France

#### Off-targets analysis report

- Description of
  libraries
- Statistics of reads
  processing
- Reads
  alignment, cut sites calling and clustering
- Best
  match(es)
  - Rank-abundance curve
  - Distribution
    of cut sites around best candidates position
- Genome distribution of
  clusters
- Configuration settings
- R session informations

Guillaume
CORRE, PhD

2026-09-02

The GUIDE-Seq (Genome-wide Unbiased Identification of Double-strand
breaks Enabled by Sequencing) method enables comprehensive detection and
quantification of off-target activity by capturing double-strand breaks
across the genome through oligonucleotide tagging, followed by
sequencing and bioinformatic analysis.

---

Working directory :
**/media/Data/gcorre/GENETHOFF/test\_dataset**

Pipeline version: **V1.3**

The analysis was performed **with** bulge tolerance.

The quantification is based on **UMI counts** .

All parameters can be found in Configuration settings at the end of
this document.

---

### Description of libraries

Table 1

| library | Cells | Genome | gRNA | gRNA sequence | PAM | Cas | type | orientation |
| --- | --- | --- | --- | --- | --- | --- | --- | --- |
| VEGFA\_s1\_K562\_neg | K562 | GRCh38 | VEGFA\_s1 | GGGTGGGGGGAGTTTGCTCC | NGG | Cas9 | guideseq | negative |
| VEGFA\_s1\_K562\_pos |  |  |  |  |  |  |  | positive |

### Statistics of reads processing

Table 2

| library | Demultiplexed | ODN\_match | Filtered |
| --- | --- | --- | --- |
| VEGFA\_s1\_K562\_neg | 2,297,044 (100%) | 2,098,470 (91.36%) | 1,878,519 (81.78%) |
| VEGFA\_s1\_K562\_pos | 1,175,378 (100%) | 1,082,221 (92.07%) | 1,067,562 (90.83%) |

> Read-pairs with length greater than **25** bp (both of
> the pair) were considered for analysis.
>
> All percentages represents % of demultiplexed reads

### Reads alignment, cut sites calling and clustering

Summary of the alignment step, calling step and clustering of cutting
sites including :

- The number of reads aligned on the genome
- The number of UMIs detected (estimation of total number of
  events)
- The number of unique ODN insertion sites
- The number of clusters.

Table 3

|  | | | | Clusters count | | | |
| --- | --- | --- | --- | --- | --- | --- | --- |
| library | Reads | UMIs | Insertions | Clusters | With gRNA match .. | And .. | count |
| VEGFA\_s1\_K562\_neg | 861,040 | 36,604 | 10,149 | 10,041 | 192 | .. 2 PCR orientations | 0 |
|  |  |  |  |  |  | .. 2 ODN orientations | 10 |
|  |  |  |  |  |  | .. In Oncogene | 0 |
| VEGFA\_s1\_K562\_pos | 513,659 | 22,460 | 5,960 | 5,937 | 139 | .. 2 PCR orientations | 0 |
|  |  |  |  |  |  | .. 2 ODN orientations | 4 |
|  |  |  |  |  |  | .. In Oncogene | 0 |

> Cut sites were identified from the alignment start position of R2
> reads.
>
> - Reads were aggregated if they share the exact same start position
>   and the same UMI sequence.
> - UMI were corrected using the **directional** method
>   with a Hamming distance tolerance of **1**.
> - Positions/UMI with more than **3** reads were
>   considered for next step.
>
> Clusters are defined as a group of ODN insertion sites within a
> distance smaller than **100** bp and characterized by :
>
> - Match of the crRNA sequence with less than **6**
>   edits withing the cluster boundaries +/- **50** bp (“With
>   gRNA match . . .” in table below)
> - Presence of reads aligning in both directions, indicating both
>   ODN orientations (“2 ODN orientations” in table below)
> - Presence of reads from 2 PCR orientations if available (“2 PCR
>   orientations” in table below)
> - Number of clusters overlapping an oncogene.
>
> Clusters with more than **2** total UMIs were
> considered.

### Best match(es)

For each library, the best match(es) are clusters with the minimal
number of edits in the gRNA and PAM sequences ( which may not
necessarily be 0 ).

Table 4

| library | Position | Cut offset | Edits\_gRNA | Edits\_PAM | N\_UMI\_cluster | Relative\_abundance | Rank |
| --- | --- | --- | --- | --- | --- | --- | --- |
| VEGFA\_s1\_K562\_neg | chr6:43769560 | -1 | 0 | 0 | 445 | 25.72 | 1 |
| VEGFA\_s1\_K562\_pos | chr6:43769560 | -2 | 0 | 0 | 156 | 20.55 | 1 |

> - “Position” : Theoretical cutting site based on gRNA alignment to
>   gDNA and nuclease offset.
> - “Cut offset” : Difference between theoretical cutting site and
>   most frequent cutting site in the cluster.
> - Edits\* : Number of INDEL+mismatches
> - “Relative Abundance” : Contribution of cluster abundance in % of
>   total UMIs count. Only clusters with gRNA match with ≤
>   **6** edits and ≥ **2** UMIs are
>   considered.
> - Rank: Rank of the best candidate(s) based on UMI count.

#### Rank-abundance curve

The rank-abundance curve provides insights into relative cluster
abundance - a steep curve indicates dominance by a few clusters, while a
shallow curve suggests more even distribution among different
clusters.

Figure 1

> - Only clusters with gRNA match and more than **1**
>   total UMIs are plotted.
> - `Red` dots correspond to clusters with
>   `minimal edits` (see table 4).
> - Top3 most abundant clusters (UMI counts) are labeled.

#### Distribution of cut sites around best candidates position

For each best candidate cluster, plot the UMI count detected around
the gRNA theoretical cut site (dashed line) for each cut site, in the
forward or reverse ODN orientation (blue and red bars respectively).

Figure 2

### Genome distribution of clusters

This figure represents the distribution of unique clusters
`with gRNA` match per chromosome, colored by
`prediction` status. Only clusters with number of edits
(INDELs and substitutions) smaller or equal to **6** are
considered.

figure 3

This figure represents the total number of UMIs (cells) per
chromosome from clusters with gRNA match, colored by
`prediction` status. Only clusters with number of edits
(INDELs and substitutions) smaller or equal to **6** are
considered.

figure 4

### Configuration settings

```
author: "Guillaume CORRE, PhD"
affiliation: "Therapeutic Gene Editing - GENETHON, INSERM U951, Paris-Saclay University, Evry, France"
contact: ""
version: 'V1.3'

clean_intermediates_files: 'FALSE'
skip_demultiplexing: 'FALSE'
quantification: "umi"                # use "umi" or "fragment" for the quantification (only if rescue_R2 is 'no")
rescue_R2: "TRUE"                    # Rescue R2 reads if R1 is too short or the pair doesn't align properly.
stop_after_umi_correction: "FALSE"    # Wether to stop just after the UMI correction step (no annotation ,no reporting, no gRNA match)


## Library informations
sampleInfo_path: "sampleInfo.csv"

read_path:  ""
R1: "Undetermined_S0_L001_R1_001.fastq.gz"
R2: "Undetermined_S0_L001_R2_001.fastq.gz"
I1: "Undetermined_S0_L001_I1_001.fastq.gz"
I2: "Undetermined_S0_L001_I2_001.fastq.gz"


################################################
## path to references
genome:
  GRCh38:
    fasta: "/media/References/Human/ensembl/GRCh38/Sequences/Homo_sapiens.GRCh38.dna.primary_assembly_ChrOnly.fa"
    index: "/media/References/Human/ensembl/GRCh38/Indexes/Bowtie2/Homo_sapiens.GRCh38.dna.primary_assembly_ChrOnly"
    annotation: "/media/References/Human/ensembl/GRCh38/Annotations/Homo_sapiens.GRCh38.113.chr.gtf.gz"
    oncogene_list: "../02-ressources/OncoList_OncoKB_GRCh38_2025-07-04.tsv"
  mouse:
    fasta: ""
    index: ""
    annotation: ""
################################################

################################################
demux_error_rate: 0.1  # max error rate for barecode during demultiplexing (this is applied to I1 and I2 concatenated)
minLength: 25          # Minimal read length after trimming, before alignment
odn_error_rate: 0      # max error rate for ODN trimming [0:1]
trim_error_rate: 0.1   # max error rate for adaptors trimming [0:1] -> set to 0.1 as quality is lower in 3' of reads
################################################

## Alignement 
################################################
aligner: "bowtie2"         # Aligner to use (bowtie2 or bwa)
minFragLength: 100         # Minimal fragment length after alignment
maxFragLength: 1500        # Maximal fragment length after alignment 

################################################

################################################
# post alignment
minMAPQ: 20                       # Min MAPQ score to keep alignments 
UMI_hamming_distance: 1           # min distance to cluster UMI using network-based deduplication, use [0] to keep raw UMIs
UMI_deduplication: "directional"  # method to correct UMI (cluster or Adjacency)
UMI_side: "3"                     # position of the UMI in the I2 read sequence: 5 = UMI+BC, 3 = BC+UMI
UMI_error_rate: 0                 # use by cutadapt to remove reads without a correct UMI pattern. this is a proportion of UMI pattern length [0.0 - 1.0].IUPAC symbol are correctly handled.
################################################


################################################
## Off targets calling
tolerate_bulges: "TRUE"           # whether to include gaps in the gRNA alignment (this will change the gap penalty during SW pairwise alignment)
max_edits_crRNA: 6                # filter clusters with less or equal than n edits in the crRNA sequence (edits = substitutions + INDELs)
ISbinWindow: 100                  # insertion sites closer than 'ISbinWindow' will be clustered together
minReadsPerUMI: 3                 # Min number of reads per UMIs (>=)
minUMIPerIS: 2                    # Min number of UMI per IS (>=)
slopSize: 50                      # window size (bp) around IS (both directions) to identify gRNA sequence (ie 50bp = -50bp to +50bp)
min_predicted_distance: 100       # distance between cut site and predicted cut site to consider as predicted
################################################

################################################
# reporting
max_clusters: 100                 # max number of cluster alignments to report
minUMI_alignments_figure: 1       # filter clusters with more than n UMI in the report alignment figure (set to 0 to keep all clusters -> can be slow)
################################################

# Prediction
################################################
SWoffFinder:
  path: "/opt/SWOffinder_1.0.1/" ## Path to SWoffinder on your server (downloaded from https://github.com/OrensteinLab/SWOffinder)
  maxE: 6                 # Max edits allowed (integer).
  maxM: 6                 # Max mismatches allowed without bulges (integer).
  maxMB: 6                # Max mismatches allowed with bulges (integer).
  maxB: 3                 # Max bulges allowed (integer).
  window_size: 100
################################################


################################################
# Sequences for the trimming steps

guideseq:
  positive:
    R1_trailing: "GTTTAATTGAGTTGTCATATGT"
    R2_leading: "ACATATGACAACTCAATTAAAC"
    R2_trailing: "AGATCGGAAGAGCGTCGTGT"
  negative:
    R1_trailing: "ATACCGTTATTAACATATGACAACTCAA"
    R2_leading: "TTGAGTTGTCATATGTTAATAACGGTAT"
    R2_trailing: "AGATCGGAAGAGCGTCGTGT"


iguideseq:
  positive:
    R2_leading: "ACATATGACAACTCAATTAAACGCGAGC"
    R2_trailing: "AGATCGGAAGAGCGTCGTGT"
    R1_trailing: "GCTCGCGTTTAATTGAGTTGTCATATGT"
  negative:
    R1_trailing: "TCGCGTATACCGTTATTAACATATGACAACTCAA"
    R2_leading: "TTGAGTTGTCATATGTTAATAACGGTATACGCGA"
    R2_trailing: "AGATCGGAAGAGCGTCGTGT"

olitagseq:
  positive:
    R1_trailing: "GGGGTTTAATTGAGTTGTCATATGTT"
    R2_leading: "AACATATGACAACTCAATTAAACCCC"
    R2_trailing: "TCCGCTCCCTCG"
  negative:
    R1_trailing: "CCCATACCGTTATTAACATATGAC"
    R2_leading: "GTCATATGTTAATAACGGTATGGG"
    R2_trailing: "TCCGCTCCCTCG"

tagseq:
  positive:
    R1_trailing: "TGCGATAACACGCATTTCGCATAA"
    R2_leading: "CTTATGCGAAATGCGTGTTATCGCA"
    R2_trailing: "AGATCGGAAGAGCGTCGTGT"
  negative:
    R1_trailing: "ATCTCTGAGCCTTATGCGAAATGC"
    R2_leading: "CGCATTTCGCATAAGGCTCAGAGAT"
    R2_trailing: "AGATCGGAAGAGCGTCGTGT"
################################################
```

### R session informations

```
## R version 4.4.3 (2025-02-28)
## Platform: x86_64-conda-linux-gnu
## Running under: Debian GNU/Linux 12 (bookworm)
## 
## Matrix products: default
## BLAS/LAPACK: /opt/miniconda/envs/GENETHOFF/lib/libopenblasp-r0.3.33.so;  LAPACK version 3.12.0
## 
## locale:
##  [1] LC_CTYPE=fr_FR.UTF-8       LC_NUMERIC=C              
##  [3] LC_TIME=fr_FR.UTF-8        LC_COLLATE=fr_FR.UTF-8    
##  [5] LC_MONETARY=fr_FR.UTF-8    LC_MESSAGES=fr_FR.UTF-8   
##  [7] LC_PAPER=fr_FR.UTF-8       LC_NAME=C                 
##  [9] LC_ADDRESS=C               LC_TELEPHONE=C            
## [11] LC_MEASUREMENT=fr_FR.UTF-8 LC_IDENTIFICATION=C       
## 
## time zone: Europe/Paris
## tzcode source: system (glibc)
## 
## attached base packages:
## [1] stats     graphics  grDevices utils     datasets  methods   base     
## 
## other attached packages:
##  [1] UpSetR_1.4.1     yaml_2.3.12      kableExtra_1.4.1 lubridate_1.9.5 
##  [5] forcats_1.0.1    stringr_1.6.0    dplyr_1.2.1      purrr_1.2.2     
##  [9] readr_2.2.0      tidyr_1.3.2      tibble_3.3.1     ggplot2_4.0.3   
## [13] tidyverse_2.0.0  rmdformats_1.0.4
## 
## loaded via a namespace (and not attached):
##  [1] sass_0.4.10        generics_0.1.4     xml2_1.6.0         stringi_1.8.7     
##  [5] hms_1.1.4          digest_0.6.39      magrittr_2.0.5     evaluate_1.0.5    
##  [9] grid_4.4.3         timechange_0.4.0   RColorBrewer_1.1-3 bookdown_0.47     
## [13] fastmap_1.2.0      plyr_1.8.9         jsonlite_2.0.0     ggrepel_0.9.8     
## [17] gridExtra_2.3.1    viridisLite_0.4.3  scales_1.4.0       textshaping_1.0.5 
## [21] jquerylib_0.1.4    cli_3.6.6          rlang_1.3.0        withr_3.0.3       
## [25] cachem_1.1.0       otel_0.2.0         tools_4.4.3        tzdb_0.5.0        
## [29] vctrs_0.7.3        R6_2.6.1           lifecycle_1.0.5    pkgconfig_2.0.3   
## [33] pillar_1.11.1      bslib_0.11.0       gtable_0.3.6       glue_1.8.1        
## [37] Rcpp_1.1.2         systemfonts_1.3.2  xfun_0.60          tidyselect_1.2.1  
## [41] rstudioapi_0.19.0  knitr_1.51         farver_2.1.2       htmltools_0.5.9   
## [45] labeling_0.4.3     svglite_2.2.2      rmarkdown_2.31     compiler_4.4.3    
## [49] S7_0.2.2
```
